# Saturation mutagenesis of disease-associated regulatory elements

**DOI:** 10.1101/505362

**Authors:** Martin Kircher, Chenling Xiong, Beth Martin, Max Schubach, Fumitaka Inoue, Robert J.A. Bell, Joseph F. Costello, Jay Shendure, Nadav Ahituv

## Abstract

The majority of common variants associated with common diseases, as well as an unknown proportion of causal mutations for rare diseases, fall in noncoding regions of the genome. Although catalogs of noncoding regulatory elements are steadily improving, we have a limited understanding of the functional effects of mutations within them. Here, we performed saturation mutagenesis in conjunction with massively parallel reporter assays on 20 disease-associated gene promoters and enhancers, generating functional measurements for over 30,000 single nucleotide substitution and deletion mutations. We find that the density of putative transcription factor binding sites varies widely between regulatory elements, as does the extent to which evolutionary conservation or various integrative scores predict functional effects. These data provide a powerful resource for interpreting the pathogenicity of clinically observed mutations in these disease-associated regulatory elements, and also comprise a gold-standard dataset for the further development of algorithms that aim to predict the regulatory effects of noncoding mutations.

## Main

The vast majority of the human genome is noncoding. Nonetheless, even as the cost of DNA sequencing plummeted over the past decade, the primary focus of sequencing-based studies of human disease has been on the ∼1% that is protein-coding, *i.e.* the exome. However, it is clear that disease-contributory variation can and does occur within the noncoding regions of the genome^1,2^. For example, there are many Mendelian diseases for which specific mutations in promoters and enhancers are unequivocally causal^3^. Furthermore, for most Mendelian diseases, not all cases are explained by coding mutations, suggesting that regulatory mutations may explain some proportion of the remainder. For common diseases, although coding regions may be the most enriched subset of the genome, the vast majority of signal maps to the noncoding genome, and in particular to accessible chromatin in disease-relevant cell types^4,5^.

Nonetheless, the pinpointing of disease-contributory noncoding variants among the millions of variants present in any single individual^6^, or the hundreds of millions of variants observed in human populations^7^, remains a daunting challenge. To advance our understanding of disease as well as the clinical utility of genetic information, it is clear that we need to develop scalable methods for accurately assessing the functional consequences of noncoding variants.

While our mechanistic understanding of regulatory sequences remains limited^8^, several groups, including us, have developed tools that summarize large amounts of functional genomic data (*e.g.* evolutionary conservation, gene model information, histone or TF ChIP-seq peaks, transcription factor binding site predictions) into scores that can be used to predict noncoding variant effects (*e.g.* CADD^9^, DeepSEA^10^, Eigen^11^, FATHMM-MKL^12^, FunSeq2^13^, GWAVA^14^, LINSIGHT^15^, ReMM^16^), segment annotations (*e.g.* chromHMM^17^, Segway^18^, fitCons^19^), or sequence-based models (deltaSVM^20^). However, although these scores are clearly informative, it remains unclear how well they work.

A major bottleneck in the development of any interpretive method for noncoding variants is the assessment of prediction quality, as there are relatively few known pathogenic noncoding variants, nor consistently ascertained sets of functional measurements of noncoding variants. A recent study by Smedley *et al.^16^* cataloged a total of 453 known disease-associated noncoding single nucleotide variants (SNVs) and used those to derive a score (ReMM). However, many of these variants fall within a small number of promoters that have been extensively studied. This leaves in question how generalizable the resulting scores are. Furthermore, catalogs of disease-associated variants like the one used by Smedley *et al.,* or available from ClinVar^21^ or HGMD^22^, provide only qualitative labels for SNVs (*e.g.* “likely pathogenic”), rather than quantitative information on the magnitude of the effect. In sum, the qualitative nature, possible ascertainment biases, and relative paucity of “known” functional noncoding variants severely limit the assessment of available methods.

To address this gap, we set out to generate variant-specific activity maps for 20 disease-associated regulatory elements, including ten promoters (of *TERT, LDLR, HBB, HBG, HNF4A, MSMB, PKLR, F9, FOXE1* and *GP1BB*) and ten enhancers (of *SORT1, ZRS, BCL11A, IRF4, IRF6, MYC (2x), RET, TCF7L2* and *ZFAND3*), together with one ultraconserved enhancer (*UC88*)^23,24^. Specifically, we used massively parallel reporter assays (MPRAs) to perform saturation mutagenesis on each of these regulatory elements^25,26^. Altogether, we empirically measured the functional effects of over 30,000 SNVs or single nucleotide deletions. We observe that the density of putative transcription factor binding site (TFBS) varies widely across the elements tested, as does the performance of various predictive strategies. These data comprise a much-needed resource for the benchmarking and further development of noncoding variant effect scores, as well as an empirical database for the interpretation of the disease-causing potential of nearly any possible SNV in these regulatory elements.

## Results

### Selection of disease-associated promoters and enhancers

We selected 21 regulatory elements, including 20 commonly studied, disease-relevant promoter and enhancer sequences from the literature (Supplementary Tables 1, 2), and one ultraconserved enhancer (*UC88*). For the former, we focused primarily on regulatory sequences in which specific mutations are known to cause disease, both for their clinical relevance and to provide for positive control variants. Selected elements were limited to 600 base pairs (bp) for technical reasons related to the mapping of variants to barcodes by subassembly^27^. In addition, we selected only sequences where cell line-based reporter assays were previously established.

For example, we selected the low-density lipoprotein receptor (*LDLR*) promoter, where mutations have been shown to cause familial hypercholesterolemia (FH), a disorder that results in accelerated atherosclerosis and increased risk for coronary heart disease^28–30^. We also tested the core promoter region (−200 to +57) of the Telomerase Reverse Transcriptase (*TERT*) gene which is associated with oncogenic mutations^31^. In particular, NM_198253.2:c.-124C>T or c.-146C>T are frequently found in several cancer types, including glioblastoma^31–34^.

We also selected a sortilin 1 (*SORT1*) enhancer. A series of genome-wide association studies showed that the minor allele of a common noncoding polymorphism at the 1p13 locus (rs12740374) creates a CCAAT/enhancer binding protein (C/EBP) TFBS and increases the hepatic expression of the *SORT1* gene, reducing LDL-C levels and risk for myocardial infarction in humans^35^. We cloned a ∼600 bp region that includes rs12740374 as well as most nearby annotated TFBS according to ENCODE data (wgEncodeRegTfbsClusteredV3 track, UCSC Genome Browser), to identify additional functional variants in the enhancer and surrounding region. For this enhancer, we also conducted MPRA experiments in both forward and reverse orientations, with the goal of testing for any directionality-dependence of variant effects.

All 21 selected promoter and enhancer regions were individually validated for functional activity in the appropriate cell lines (Supplementary Figures 1, 2; Supplementary Tables 1, 2). This initial validation allowed us to optimize reporter assay conditions and to confirm that the cloned subsequences of the candidate regulatory elements were sufficient for measurable activities in the appropriate cell types. The validated luciferase expression levels ranged from 2- to 200-fold over empty vector (Supplementary Tables 1, 2).

### Saturation mutagenesis and construction of MPRA libraries

In order to test the functional effects of thousands of mutations in these selected disease-associated regulatory elements, we first developed a scalable protocol for saturation mutagenesis-based massively parallel reporter assays (MPRAs)^25,26^ (Figure 1). For each of the 21 regulatory elements (Supplementary Tables 1, 2), we used error-prone PCR to introduce sequence variation at a frequency of less than 1 change per 100 bp. While error-prone PCR is known to be biased in the types of mutations that are generated (*e.g.* a preference for transitions and T/A transversions)^36^, high library complexities (50k-2M constructs per target) allowed us to capture nearly all possible SNVs as well as many single base pair deletions with multiple independent constructs per variant (Supplementary Table 3). To distinguish the individual amplification products, we incorporated 15 or 20 bp random sequence tags 3’ of the target region using overhanging primers during the error-prone PCR.

**Figure 1.**
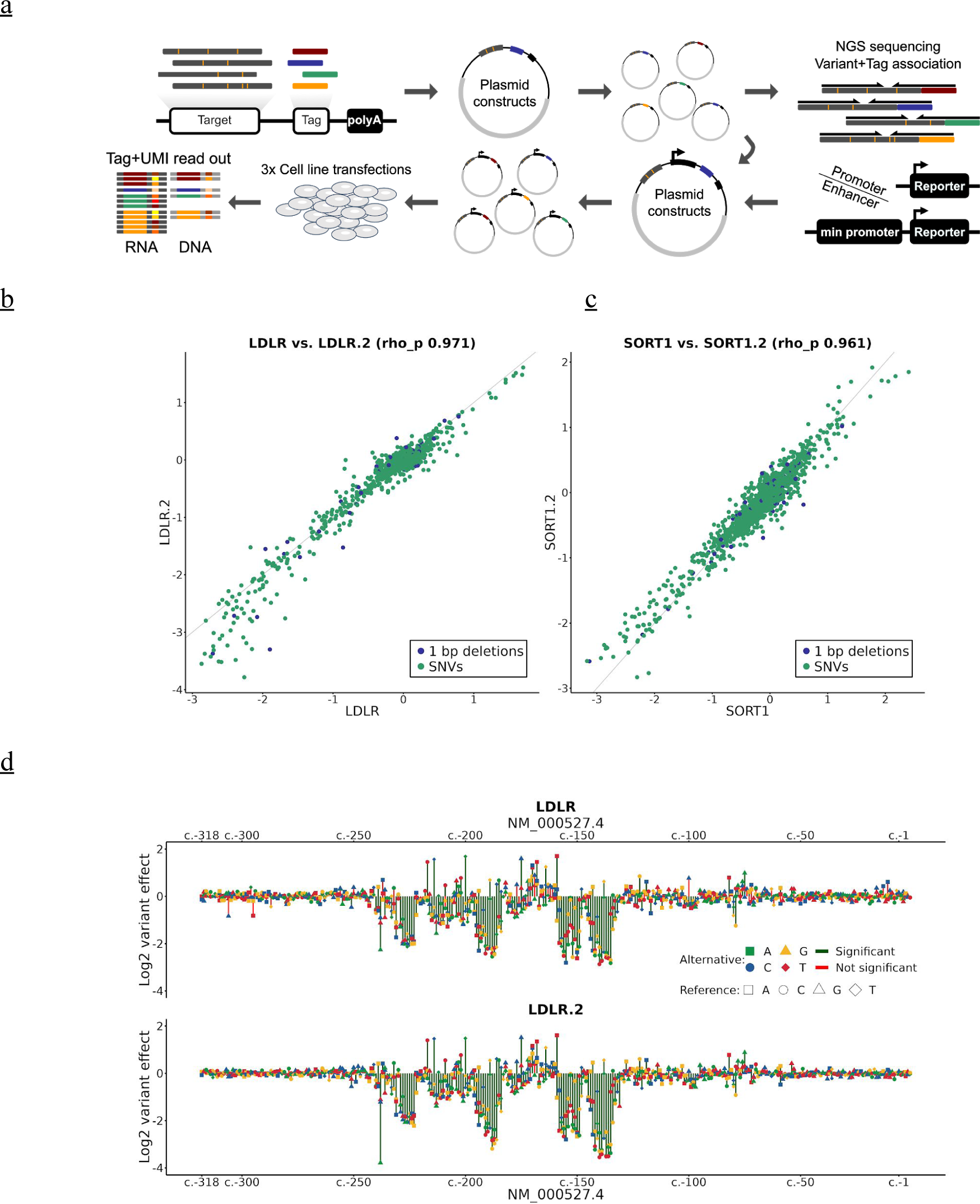
Saturation mutagenesis MPRA of disease-associated regulatory elements including the hypercholesterolemia-associated *LDLR* promoter. **(a)** Saturation mutagenesis MPRA. Error-prone PCR is used to generate sequence variants in a regulatory region of interest. The resulting PCR products with approximately 1/100 changes compared to the template region are integrated in a plasmid library containing random tag sequences in the 3’ UTR of a reporter gene. Associations between tags and sequence variants are learned through high-throughput sequencing. High complexity MPRA libraries (50k-2M) are transfected as plasmids into cell lines of interest. RNA and DNA is collected and sequence tags are used as a readout. Variant expression correlation (min. 10 tags required) between full replicates of **(b)** *LDLR* (LDLR; LDLR.2) and **(c)** *SORT1* (SORT1; SORT1.2). **(d)** Log^2^ variant effect of all SNVs (min. required tags 10) ordered by their RefSeq transcript position in NM_000527.4 of *LDLR*. Upper part shows the LDLR experiment, lower the full replicate LDLR.2. Significance level is 10^−5^ in both expression profiles.

We then cloned promoters and all but two enhancers into the backbones of slightly modified pGL4.11 (Promega, promoter) or pGL4.23 (Promega, enhancer) vectors (see Supplementary Tables 1, 2), respectively, from which the reporter gene (as well as the minimal promoter in the case of enhancers) had been removed. For each of the 21 regulatory elements, we determined which variants were linked to which random tag sequences by deeply sequencing the corresponding library (see Methods). In the final step, we inserted the luciferase reporter gene (as well as the minimal promoter in the case of enhancers) in between the regulatory element and the tag sequence, and transformed the MPRA library into *E.coli*. With this insertion of the luciferase reporter gene, the above-introduced random tag sequence becomes part of its 3’ untranslated region (3’ UTR).

We obtained tag assignments (*i.e.* variant-tag associations) for a total of 24 saturation mutagenesis libraries (see Methods). This included the 21 selected regions listed in Supplementary Tables 1, 2, as well as an additional full replicate for the *LDLR* and *SORT1* enhancer libraries, and an additional *SORT1* library with reversed sequence orientation of the enhancer. Supplementary Figure 3 plots the number of tags associated with substitutions and 1 bp deletions along the target sequences for each library. The representation of tags associated with specific variants follows previously characterized biases in error-prone PCR using Taq polymerase^37^, with a preference of transitions over transversions and T-A preference among transversions. Insertions were rare, while short deletions occur at rates similar to those of the rare transversions. For all libraries, we observed complete or near-complete coverage of all potential SNVs (Supplementary Table 3) as well as partial 1 bp deletion coverage. On average, 99.9% (min. 99.1%, max. 100%) of all potential SNVs in the targeted regions are associated with at least one tag, while on average 55.4% (min. 31.4%, max. 71.1%) of 1 bp deletions are associated with at least one tag.

### Saturation mutagenesis MPRAs of disease-associated promoters and enhancers

For each MPRA experiment, around 5 million cells (Supplementary Tables 1, 2) were plated and incubated for 24 hours before transfection with the libraries. In each experiment, three independent cultures (replicates) were transfected with the same library. In addition, for *LDLR* and *SORT1*, independent MPRA libraries were created, as outlined above, and cells were transfected from a different culture and on a different day. In one case (*TERT*), the same MPRA library was used for experiments in two different cell-types (HEK293T and a glioblastoma cell line).

We then used our published protocol for quantifying effects from RNA and DNA tag-sequencing readouts, including the previously suggested modification of using unique molecular identifiers (UMIs) during targeted amplification^38^ (see Methods). More specifically, the relative abundance of reporter gene transcripts driven by each promoter or enhancer variant was measured by counting associated 3’ UTR tags in amplicons derived from RNA (obtained by targeted RT-PCR), and normalized to its relative abundance in plasmid DNA (obtained by targeted PCR). For all experiments, we excluded tags not matching the assignment and determined the frequency of a tag in RNA or DNA from high-throughput sequencing experiments based on the number of unique UMIs. We only considered tag sequences observed in both RNA and DNA. Supplementary Table 4 summarizes the number of RNA and DNA counts obtained in each experiment. From individual tag counts in RNA and DNA, we fit a multiple linear regression model to infer individual variant effects (see Methods).

For data quality reasons, we introduced a minimum threshold on the number of associated tags per variant used in model fitting (various quality measures for fitted variant effects versus the number of tags are plotted in Supplementary Figure 4). We picked this threshold based on the correlation of variant effects obtained when comparing between the independent libraries of *LDLR* (Figure 1b, Supplementary Figure 5) and *SORT1* (Figure 1c, Supplementary Figure 6). Using all SNVs and 1 bp deletions with at least one associated tag in each transfection replicate, variant effects show a Pearson correlation of 0.93 (*LDLR*) and 0.94 (*SORT1*). When requiring a minimum of 10 tag measurements after combining all three transfection replicates, correlations increase to 0.97 (*LDLR*) and 0.96 (*SORT1*). Requiring even higher thresholds (Supplementary Table 5) further improves replicate correlation up to 0.98 for both experiments (min. 50 tags), but reduces SNV coverage to 86.3% and 1 bp deletion coverage to 15.6% across all datasets (Supplementary Table 6). We therefore used a minimum of 10 tags, reducing average coverage from 99.8% to 98.4% [range 93.0%, 100.0%] for all putative SNVs and from 44.4% to 25.3% [range 10.5%, 41.7%] for 1 bp deletions.

To assure high quality of our complete dataset, we evaluated the Pearson correlation of variant effects divided by their standard deviation among pairs of transfection replicates (Supplementary Table 7). We observed the lowest replicate correlation for *BCL11A, FOXE1* and one of the *MYC* elements (rs1198622). In contrast, experiments for *HBG1, IRF4, LDLR, SORT1* and *TERT* exhibited high reproducibility among transfection replicates (Pearson correlation > 0.9). Exploring differences in the proportion of alleles with significant regulatory activity, we observed a wide range of values across elements (3-52% of variants using a lenient p-value threshold of < 0.01; average 22%; Supplementary Table 7). We find that this proportion is strongly correlated with the performance of transfection replicates (Pearson correlation of 0.78), but we also note a circular relation for the significance of results, low experimental noise and high reproducibility.

We sought to explore whether factors like target length, wild-type activity in the luciferase assay (Supplementary Figure 1, Supplementary Tables 1, 2), measures of assignment complexity (Supplementary Table 3), as well as DNA and RNA sequencing depth (Supplementary Table 4) contribute to technical reproducibility. Linear models of up to three features fit in a leave-one-out setup explain up to 33% of variance (luciferase wild-type activity, average number of tags per SNVs and the proportion of wild-type haplotypes) or 29% of the variance (luciferase wild-type activity, average number of variants per haplotype, average number of DNA counts obtained) in reproducibility between transfection replicates. Overall, these analyses emphasize the baseline activity of a regulatory element as the largest factor (*i.e.* highly active elements are associated with greater technical reproducibility).

### General properties of observed effects of regulatory mutations

Altogether, our MPRAs quantified the regulatory effects of 31,243 potential mutations (min. 10 tags) at 9,834 unique positions (Supplementary Table 8). Of these mutations, 4,830 (15%) were identified as causing significant changes relative to the wild-type promoter or enhancer sequence (p-value < 10^−5^). Of those causing significant changes, 1,791 (37%) increased expression (by a median of 20%) and 3,039 (63%) decreased expression (by a median of 24%). The majority of significant effects were of small magnitude. If we require a minimum 2-fold change, we identify a total of 84 activating and 561 repressing mutations. The significant shift towards repressing mutations (binomial test, p-value < 10^−42^) is consistent with a model where most transcriptional regulators are activators and binding is more easily lost than gained with single nucleotide changes.

Out of the 31,243 successfully assayed mutations, 2,306 are 1 bp deletions, of which 229 meet the significance threshold (p-value < 10^−5^). This is a lower proportion than observed for SNVs (10% vs. 16%), most likely due to the lower rates at which 1 bp deletions are created by error-prone PCR, resulting in representation by fewer tags. Supporting this notion, 1 bp deletions tend to be associated with larger absolute effect sizes than SNVs (Wilcoxon Rank Sum test with continuity correction, p-value < 0.05, location shift 0.04). Similarly, we observe a large shift towards repressive effects with 1 bp deletions (significant effects: 33% activating [27%, 40%], 76 activating and 153 repressing; min. 2-fold change 5% activating [1%, 17%], 2 activating and 37 repressing), but due to the low number of observations, this shift is not significantly different than that observed for SNVs (significant effects: 37% activating [36%, 39%], 1,715 activating and 2,886 repressing; min. 2-fold change 14% activating [11%, 17%], 82 activating and 524 repressing).

We have greater power to detect significant effects for transitions than transversions, likely consequent to the higher sampling by error-prone PCR (Supplementary Table 9; binomial test comparing the proportion of significant transitions (2,190/9,824) vs. transversions (2,411/19,113); p-value < 10^−16^). This is supported by specific transversions (A-T, T-A) that are also created more frequently by error-prone PCR (Supplementary Figure 3) and represent a higher proportion of significant observations (A-T 392/2,334 and T-A 404/2,327, combined binomial test vs. all transversions, p-value < 10^−16^). Despite our greater power for assaying transitions, transversions had larger absolute effect sizes (Wilcoxon Rank Sum test with continuity correction, p-value < 10^−16^, location shift 0.14). This observation supports a model were regulatory elements evolved some level of robustness to the more frequent transitional changes (as for coding sequences^39^), and is consistent with previous research showing that transversions have a larger impact on TF motifs and allele-specific TF binding^40^.

Our increased power to measure the effects of transitions resulted in an artifactual enrichment for significant effects among SNVs previously observed in gnomAD r2.1^7^ (binomial test; overlap of tested SNVs with gnomAD, n=707/28,937; of those with significant effects, n=146/707; p-value < 0.001), where 64% of SNVs are transitions compared to 34% of mutations created in our libraries. However, testing separately for transitions and transversions, there is no enrichment of significant effects among SNVs previously observed in gnomAD. In fact, we observed a significantly smaller absolute effect size for previously observed SNVs (p-value < 0.05, location shift 0.04; Wilcoxon Rank Sum test with continuity correction). This is intensified if we exclude singletons (excl. 82/146 significant variants; p-value < 0.01, location shift 0.07; Wilcoxon Rank Sum test with continuity correction), consistent with purifying selection acting on standing regulatory variation.

The most obvious pattern upon visual inspection of the data is a strong clustering of positions associated with significant mutations (*e.g.* Figure 1d). This clustering was non-random for all but the *F9* and *FOXE1* experiments (p-value < 0.01; for 16/21 elements, p-value < 10^−5^; Wilcoxon Rank Sum tests with continuity correction vs. 1000 data shuffles), as determined from comparing run lengths for significant changes including directionality of the change. While *FOXE1* is one of the experiments mentioned above with low experimental reproducibility, a non-random clustering (p-value < 10^−9^) of significant regulatory changes was observed in *F9* when additionally requiring a minimum effect size of 20%. These results are consistent with expectations for TFBS (specific examples are discussed below).

In the below sections, we describe the results of saturation mutagenesis of three of the regulatory elements in greater detail: the *LDLR* promoter, the *TERT* promoter, a *SORT1*-associated enhancer. Similar expositions on the remaining 18 elements are provided as Supplementary Note S1. Finally, we compare the relative performance of various computational tools for predicting these empirical measurements of regulatory effects.

### Low density lipoprotein receptor (LDLR) promoter

Familial hypercholesterolemia (FH) is an autosomal dominant disorder of low-density lipoprotein (LDL) metabolism, which results in accelerated atherosclerosis and increased risk of coronary heart disease^41^. With a prevalence of about 1 in 500 individuals, FH is the most common monogenic disorder of lipoprotein metabolism. It is mainly due to mutations in the LDL receptor (*LDLR*) gene that lead to the accumulation of LDL particles in the plasma^42^. Several studies have shown that variants in the *LDLR* promoter can alter the transcriptional activity of the gene and also cause FH^28–30^ (full reference list in Supplementary Table 10). While in some cases, mutations in the promoter were identified in patients, a functional follow-up, like testing the regulatory effect of the variants by means of a luciferase assay, was not always conducted^43–45^. To decipher the functional activity of these previously identified, as well as essentially all potential SNVs in the *LDLR* (NM_000527.4) promoter, we performed saturation mutagenesis MPRA in HepG2 cells, a commonly used cell line for *LDLR* functional studies^30^. Experiments for this promoter were performed using two full replicates (*i.e.* two independently constructed saturation mutagenesis libraries), referred to below as LDLR and LDLR.2 (Figure 1b).

We observed strong concordance between our MPRA-based measurements and variants with previous luciferase activity results (Supplementary Table 10). For example, a c.-152C>T mutation was previously reported to reduce promoter activity (to 40% of normal activity), while a c.-217C>T variant was shown to increase transcription (to 160% of normal activity)^28^. We observe a reduction of 32%/39% (LDLR/LDLR.2) and activation of 273%/263% (LDLR/LDLR.2) for these variants, respectively. c.-142C>T reduced promoter activity (to 20% of normal activity) in transient transfection assays in HepG2 cells^29^, and we observe 20%/11% (LDLR/LDLR.2) residual activity. Mutations located in regulatory elements R2 and R3 (c.-136C>G, c.-140C>G, and c.-140C>T) resulted in 6 to 15% residual activity^30^; our MPRA results confirm these findings in both replicates (10-22% residual activity). We also observed no significant changes in promoter activity for c.-36T>G and c.-88A, consistent with a previous study of these variants^30^.

Overall, we observe that variants located in close proximity and overlapping the same TFBS tend to show similar deactivating effects (*e.g.* SP1 and SREBP1/SREBP2 sites in Supplementary Table 10 and Figure 1d). Previously reported variants located in the 5’ UTR of *LDLR* generally did not affect promoter activity. The high concordance between full replicates (Pearson correlation of 0.97) as well as the agreement with previous studies give us confidence in the potential of our MPRA results to be useful for the clinical interpretation of *LDLR* promoter mutations. It also reinforces the value of functional assays covering all possible variants of a regulatory sequence of interest, as this provides consistent and comparable readouts together with a distribution of effect sizes.

### Telomerase reverse transcriptase (TERT) promoter

Mutations in the *TERT* promoter (NM_198253.2), in particular c.-124C>T or c.-146C>T, increase telomerase activity and are among the most common somatic mutations observed in cancer^31–34^. Previous luciferase assay studies showed that these mutations increase promoter activity in human embryonic kidney (HEK) 293 cells, glioblastoma, melanoma, bladder cancer and hepatocellular carcinoma (HepG2) cells^31,46–48^. In glioblastoma cells, c.-124C>T or c.-146C>T mutations result in a 2-4 log^2^ fold increase in promoter activity^48^.

Here, we tested the *TERT* promoter MPRA library in two different cell types, HEK293T and glioblastoma SF7996 cells^49^, referred to here as GBM (Figure 2a), observing a log^2^ fold increase in promoter activity of 2.00/2.86-fold for c.-124C>T and 1.42/2.42-fold for c.-146C>T in HEK293T and GBM cells, respectively. We also identified additional activating mutations, some of which were previously identified in cancer studies^46,50,51^ and are annotated in COSMIC^52^. These include c.-45G>T and c.-54C>A, previously identified as somatic mutations in bladder cancers^50,51^, and c.-57A>C, previously associated with both melanoma^46^ and bladder cancer^51^. We observed activating effects for c.-45G>T and c.-54C>A with a log^2^ increase of 0.81/1.65-fold and 0.45/1.03-fold for HEK293T and GBM cells, respectively. For c.-57A>C, we observed a 0.65/1.14-fold log^2^ increase in HEK293T and GBM cells, respectively, similar to previous reporter assays that obtained increased expression of 0.6-fold (152%) and 0.3-fold (123%) on a log^2^-scale over the wild-type construct in Ma-Mel-86a and HEK293T cells^46^, respectively.

**Figure 2.**
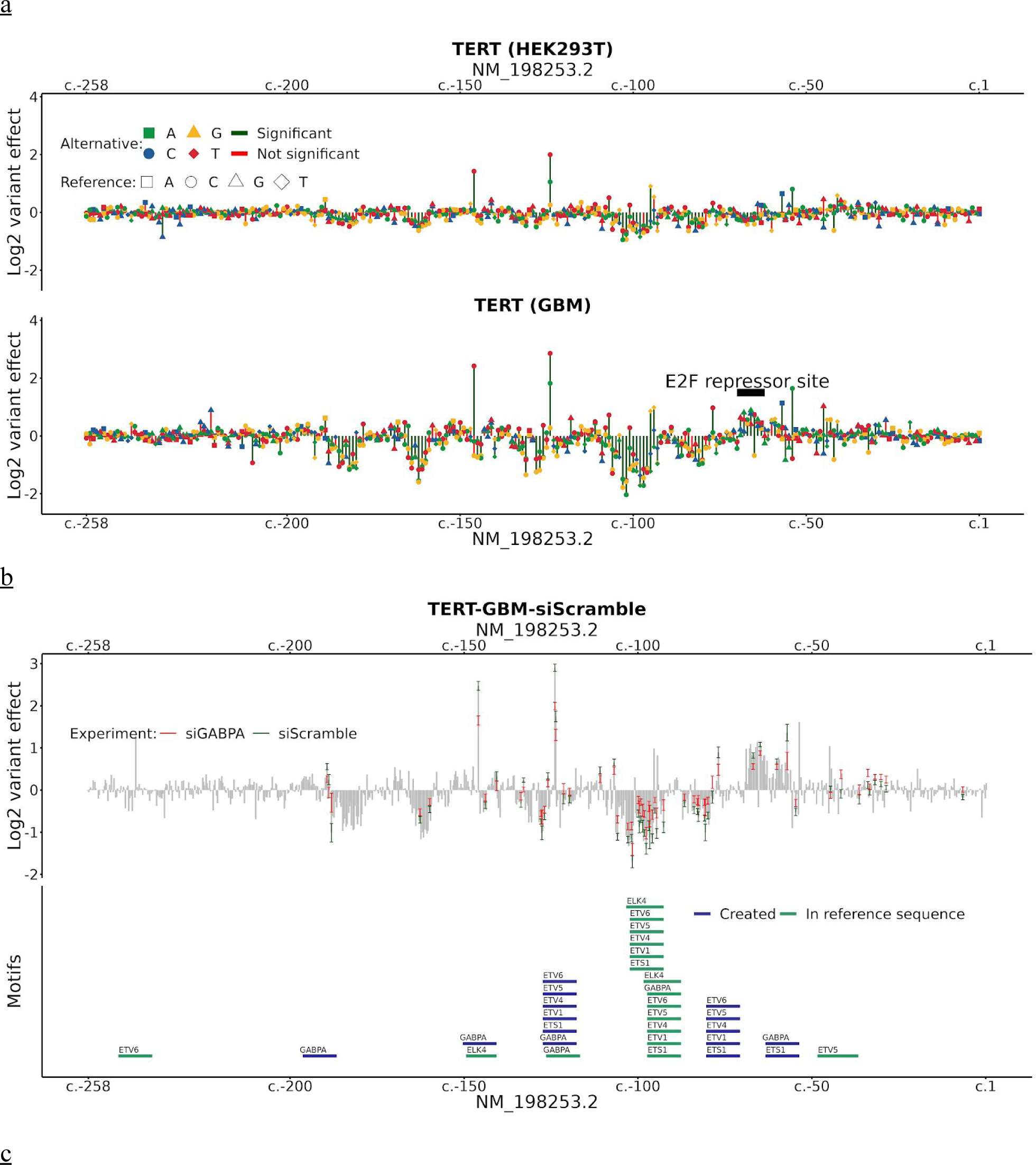
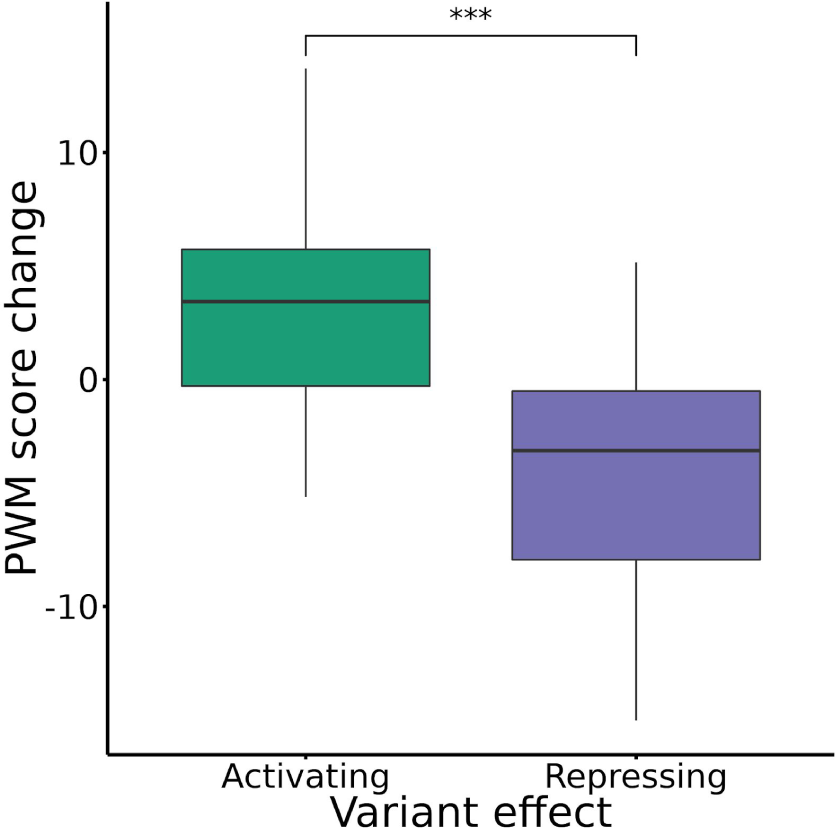
Saturation mutagenesis MPRA of the cancer-associated *TERT* promoter. **(a)** Log^2^ variant effect of all SNVs (min. 10 tags required) ordered by their RefSeq transcript position in NM_198253.2 of *TERT*. Upper panel shows the *TERT* experiment in HEK293T cells and the lower in GBM (SF7996) cells, where the E2F repressor site is marked. Significance level is 10^−5^ in both expression profiles. **(b)** Expression profile of TERT-GBM-siScramble (grey). 95%-confidence intervals of variants from TERT-GBM-siScramble (green) and TERT-GBM-siGABPA (red), that were significantly different between the two experiments, are overlaid. In addition, predicted Ets-related motifs in the reference sequence (green) or variant induced Ets-related motifs (blue) are marked. **(c)** Motif score change of variants that show a significant difference between siGABPA and scramble siRNA experiment. Motif scores are ploted as boxplots with median center line, upper and lower quartiles box limits and 1.5x interquartile range whiskers. Variants were only used if they overlapped an Ets-related factor motif (GABPA, ETS1, ELK4, ETV1, ETV4-6) with a score (reference or alternative sequence) larger than the 80th percentile of the best possible motif match to the position weight matrix (PWM). TERT-GBM-siGABPA variants effects where divided by the effect measured in the siRNA scramble experiment. Three asterisks (* * *) mark a significance level of 10^−9^ by the two sided Wilcoxon-rank sum test.

A common single nucleotide polymorphism (SNP) in the *TERT* promoter, rs2853669 (c.-245A>G), previously studied in several cell lines and cancer cohorts, has been suggested to alter promoter-mediated *TERT* expression by impacting E2F1 or Ets/TCF binding^53,54^. However, studies in both breast cancer^55^ and glioblastomas^56^ failed to find any impact on risk or prognosis of this polymorphism. The epidemiologic findings are in line with our results, as we did not observe a significant effect of this variant on promoter activity in either cell type.

We next sought to assess whether there are differences in mutational effects on the *TERT* promoter between HEK293T and GBM cells that could be driven by the *trans* environment. Overall, variant effects were highly concordant between the two cell types (Figure 2a). However, we did observe significant differences at several specific positions. In particular, variants between c.-62 to c.-70, which corresponds to an E2F repressor site, were found to increase promoter activity in GBM cells, likely due to disruption of this motif (Figure 2a). None of these effects were observed in HEK293T cells, suggesting that different E2F family protein abundances could be driving the differences in promoter activity between these cell types, and potentially between the corresponding cancer types.

Previous work has shown that the commonly observed cancer-associated activating somatic mutations, c.-146C>T and c.-124C>T, lead to the formation of an ETS binding site that is bound by the multimeric GABP transcription factor in GBM cells^48^. To evaluate the relevance of GABP binding on TERT promoter activity more globally, we re-tested our *TERT* MPRA library in GBM cells with an siRNA targeting GABPA. We first optimized GABPA knockdown conditions using qPCR, such that it reduced GABPA expression by 68% ± 7% and TERT promoter activity by 58% ± 12%, compared to a scrambled siRNA control (Supplementary Figure 7). We then tested our MPRA library in GBM cells using either the GABPA siRNA or the scrambled control. A total of 63 variants were identified as significantly different (see Online Methods and Supplementary Table 11), 59 leading to a reduction and 4 to an increase in activity. Both c.-146C>T and c.-124C>T, previously reported to create GABP binding sites, showed significantly reduced activity in the siGABPA knockdown compared to the scrambled control (Figure 2b). Variants with significantly different activity were 2.8-fold enriched for Ets-related factor motifs (ETS1, ELK4, ETV1, ETV4-6, GAPBA) annotated from JASPAR 2018^57^ as compared with all 908 other variants present in both experiments (p-value < 0.001; one-sided Fisher’s exact test).

Apart from the two variants (c.-146C>T and c.-124C>T) known to create GABPA binding sites, we identified 9 additional variants with the potential to create new ETS family motifs from the siGABPA knockdown (Supplementary Table 11). To include such instances as part of a global analysis, we computed score differences of the corresponding position weight matrices from activating and repressing variants that overlap either a reference or newly created ETS family motif (reference or alternative sequence larger than the 80^th^ percentile of the motifs’ best matches; 12 activating and 26 repressive variants). Figure 2c shows that activating variants create new ETS-related factor motifs and repressing variants disrupt them. The PWM score changes are highly significant between activating and repressing variants (one-sided Wilcoxon Rank Sum test, p-value < 10^−11^). Overall, the average expression reduction for the motif-disruptive allele in cases where variants disrupt or create ETS family motifs is 29.5%±14%, which is in concordance with the 58%±12% reduced qPCR expression of *TERT* (Supplementary Figure 7).

### Sortilin 1 (SORT1) associated enhancer

The Sortilin 1 (*SORT1*)-associated enhancer was identified via a common SNP, rs12740374 (GRCh37 chr1:109,817,590G>T), which is associated with myocardial infarction^58^. The minor allele T creates a potential CCAAT/enhancer binding protein (C/EBP) binding site, leading to ∼4-fold greater luciferase activity compared to the major allele G in a reporter assay^35^. This is thought to alter the expression of the *SORT1* and proline and serine rich coiled-coil 1 (*PSRC1*) genes, leading to changes in LDL and VLDL plasma levels^35^. The major allele is thus associated with higher LDL-C levels and increased risk for myocardial infarction. These results are also consistent with prior human lipoprotein QTL analyses^59,60^.

We carried out saturation-based MPRA in HepG2 cells using a 600 bp region encompassing rs12740374 with two full replicates (*i.e.* two independently constructed saturation mutagenesis libraries transfected at different days with three technical replicates each, SORT1 and SORT1.2, Figure 1c). Consistent with the literature, our MPRA results show a significant effect for rs12740374, leading to a 2.92/2.74 log^2^-fold increase in expression. Furthermore, we observe many other substitutions of large effect, with a disproportionate number of >2-fold expression changes in our overall dataset (144/645) occuring in the SORT1 enhancer (Supplementary Table 8). The locations of these variants are strongly clustered (Figure 3), indicative of several TFBS in this region.

**Figure 3.**
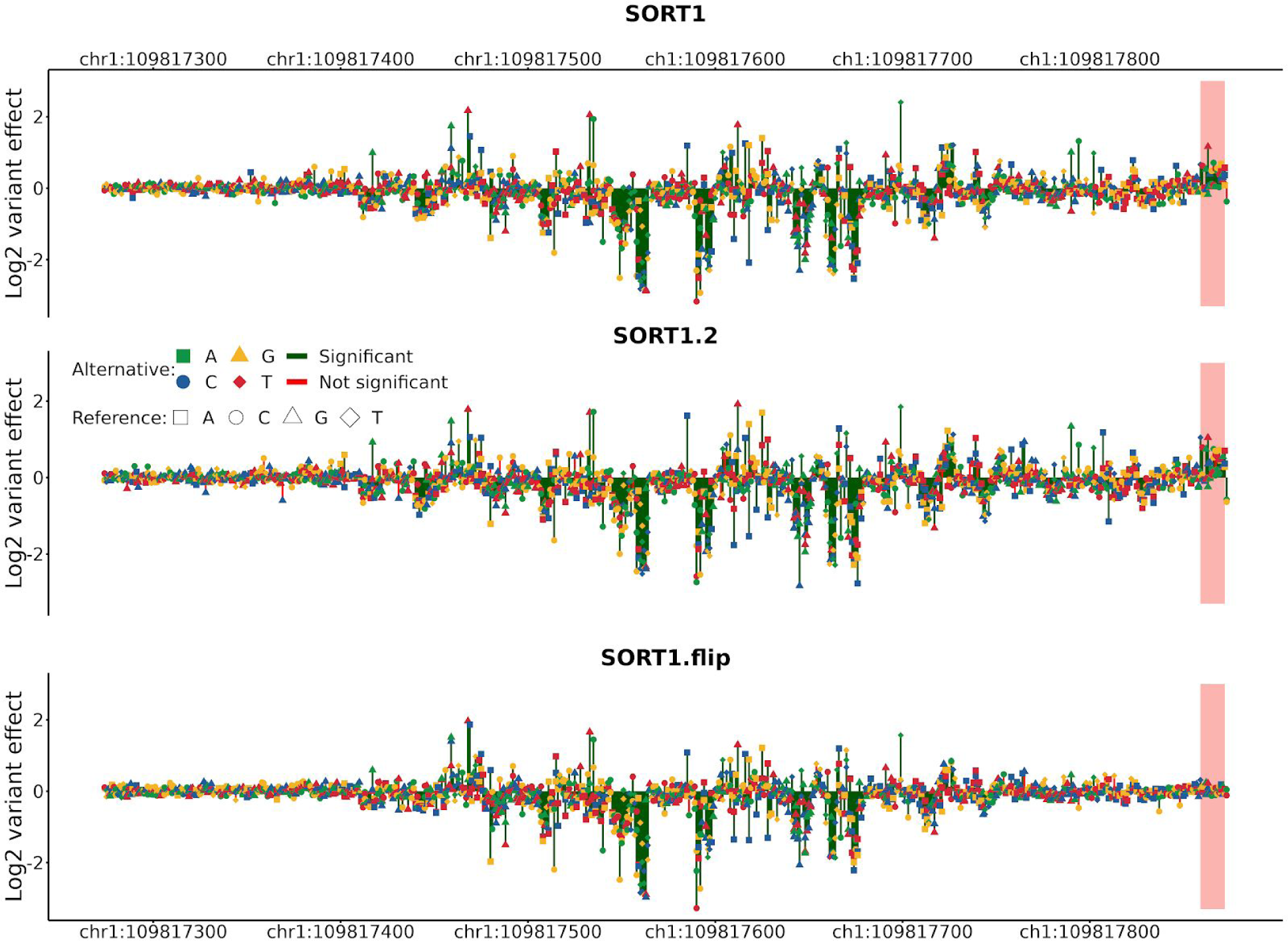
Saturation mutagenesis MPRA of a myocardial infarction-associated *SORT1* enhancer. Expression effects of SNVs from experiments SORT1, SORT1.2 and SORT1.flip. Direction of SORT1 and SORT1.2 was from left to right in the experiments. In the SORT1.flip experiment, the direction was reversed (right to left in the figure). Highlighted area in red, close to the experimental promoter site in SORT1 and SORT1.2, is different between the SORT1/SORT1.2 and SORT1.flip experiments. In this region, JASPAR annotates an EBF1 motif (MA0154.3). Significance level of variants is 10^−5^.

The directionality-independence of enhancers is inherent to their definition but is not often tested. To evaluate whether the orientation of an enhancer could bias the effects of mutations within it, we generated a third *SORT1* enhancer library where the orientation of the enhancer was flipped, termed SORT1.flip. Comparison of all variants showing a difference in activity compared to the reference sequence (p-value < 10^−5^) in the SORT1.flip with the other two libraries in the opposite orientation (SORT1 and SORT1.2), we observe a very strong correlation (0.97 and 0.96 for SORT1 and SORT1.2 respectively; Pearson’s correlation), on par with the two biological replicates in the same orientation, SORT1 and SORT1.2 (0.98; Pearson’s correlation). This result supports the directionality-independence of enhancers^61^ as well as of the effects of variants within them.

However, we did observe a few significant differences between SORT1/SORT1.2 and SORT1.flip near the 3’ region of the forward orientation (GRCh37 chr1:109,817,859-109,817,872), *i.e.* adjacent to the minimal promoter on the reporter construct. This block of variants led to a significant increase in activity in the forward orientation (SORT1/SORT1.2) but not in the opposite orientation (SORT1.flip) (Figure 3). Analysis of this region for TFBS using JASPAR 2018^57^ found a potential EBF1 motif (MA0154.3), suggesting that this factor might lead to the orientation activity differences observed in our assays. EBF1 is known to act as an activator or repressor of gene expression^62,63^ and mutations to its core motif sequences (GGG and CCC) had the strongest effect in our SORT1 assay. Nonetheless, its 3’ location and orientation-dependence suggests that it is likely contributing to minimal promoter rather than enhancer activity.

### Current computational tools are poor predictors of expression effects

Altogether, our MPRAs analyzed over 30,000 different mutations for their effects on regulatory function. We next set out to assess the performance of available computational tools and annotations for predicting the regulatory effects of individual variants. For this purpose, we examined various measures of conservation (PhyloP^64^, PhastCons^65^, GERP++^66^) as well as a number of computational tools that integrate large sets of functional genomics data into combined scores (CADD^9^, DeepSEA^10^, Eigen^11^, FATHMM-MKL^12^, FunSeq2^13^, GWAVA^14^, LINSIGHT^15^, ReMM^16^). In addition, we previously identified the number of overlapping TFBS as a significant predictive measure of the activity of a specific region^38^. We therefore also analyzed TFBS annotations resulting from motif predictions overlayed with biochemical evidence from ChIP-seq experiments (available as Ensembl Regulatory Build (ERB)^67^ and ENCODE^68^ annotations) as well as pure motif predictions from JASPAR 2018^57^. Using JASPAR predictions, we extended this analysis to individual positions and explored different score thresholds or just the factors predicted most frequently across the region (see Methods). All these annotations and scores are agnostic to the cell type(s) in which we studied each sequences. Therefore, we also compared our results to sequence-based models (deltaSVM^20^) for 10 of 21 MPRAs, *i.e.* where a model was publically available for the corresponding cell type (HEK293T, HeLa S3, HepG2, K562, and LNCaP). In cases where an annotation is based on positions rather than alleles, we assumed the same value for all substitutions at each position. We did not include the 1 bp deletions in this analysis, as most annotations are not defined for deletions.

Supplementary Tables 12 and 13 report the Pearson and Spearman correlation of the obtained expression effect readouts with conservation metrics, combined annotation scores and overlapping TF predictions, respectively. Figure 4 and Supplementary Figure 8, 9 visually contrast expression effects as well as a subset of these annotations (including ENCODE^68^ and Ensembl^67^ motif annotations). By correlating absolute expression effects with functional scores, we identified species conservation as a major driver of the currently available combined scores. For example, the correlation results of CADD, Eigen, FATHMM-MKL and LINSIGHT show more than 90% Pearson correlation across elements with the results of PhastCons scores calculated from the alignment of mammalian genome sequences. However, conservation seems only informative for a subset of the studied noncoding regulatory elements (*e.g. LDLR, ZFAND3*, and *IRF4*, but not *SORT1, F9*, or *GP1BB*). We observe that repressive effects can at least be partially aligned with available motif data (*e.g. F9, GP1BB, IRF4, LDLR, SORT1*). However, experimentally supported motif annotations are frequently incomplete (for an example see Figure 4a, around c.-215 where motifs are predicted in JASPAR but absent from ENCODE and ERB). In several cases, motif annotation is also missing completely. For 13 of the 21 elements studied here, no motif annotation was available from ERB; for 2 of the 21 elements no motif annotation was available from ENCODE. Furthermore, the gain-of-binding motifs, *e.g.* the motifs underlying activating mutations in *TERT*, are currently not at all or insufficiently modeled in available scores, as these motifs are frequently missed by scans of the reference genome.

**Figure 4.**
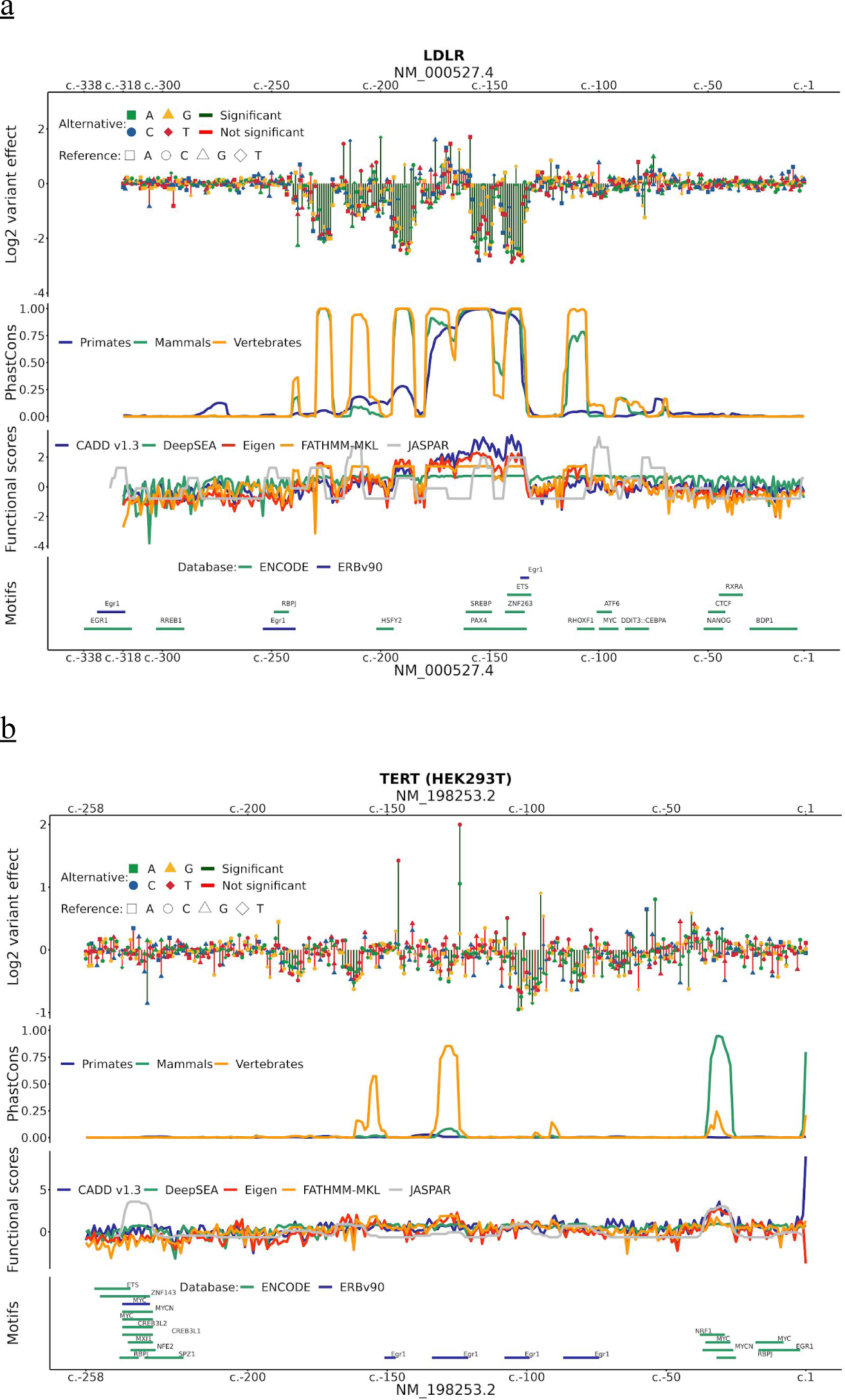
Current computational tools are poor predictors of expression effects. Expression effects of **(a)** *LDLR* and **(b)** *TERT* (significance level 10^−5^) compared to PhastCons^65^ conservation scores, combined scores of functional genomics data (CADD^9^ v1.3, DeepSEA^10^, Eigen^11^, FATHMM-MKL^12^, and number of overlapping 10th percentile scoring JASPAR^12,57^ motifs) and annotated motifs by ENCODE^68^ and Ensemble Regulatory Build (ERB) v90^67^.

Looking at average Spearman correlations across our 21 regions (Table 1, Supplementary Table 13), DeepSEA (0.22) performed best, followed by FunSeq2 (0.14), high-scoring (top 10^th^ percentile) JASPAR predictions (0.14) and Eigen (0.14). However, the average is a poor measure here. Spearman correlations for absolute expression effects of some elements were reasonably high for several methods (0.3-0.6), while for other elements no or negative correlations were detected for most or all methods. We saw the best performance in predicting an individual element (*LDLR)* for FATHMM-MKL (0.59), followed by PhastCons (0.58) and Eigen (0.58). Besides *LDLR*, the next best agreement between annotations and absolute expression effects was observed for *F9* (top 10^th^ percentile JASPAR 0.52), *IRF4* (Eigen 0.48), *ZFAND3* (DeepSEA 0.44) and *PKLR* (LINSIGHT 0.41). The lowest agreement was observed for *FOXE1* (DeepSEA 0.05) and *BCL11A* (DeepSEA 0.03); the two elements observed with the lowest replicate correlation (Supplementary Table 7). However, high replicate correlations did also not indicate high correlations with existing scores or annotations. For example, *SORT1* (replicate correlations of 0.99) and *TERT* (replicate correlations of >0.96 for GBM experiment) have highest correlations of 0.29 (top 10^th^ percentile JASPAR) and 0.37 (DeepSEA), respectively.

**Table 1.** Spearman correlation of computational scores with measured expression effects. The table reports Spearman correlation coefficients (in percent) of the absolute expression effect for all SNVs with at least 10 tags in each region with various measures agnostic to the cell-type, like conservation (mammalian PhyloP, mammalian PhastCons, GERP++), overlapping TFBS as predicted in JASPAR 2018 (counting all or those in the top 10^th^ percentile of motif scores across all elements, additional percentiles available in Supplementary Table 13) and computational tools that integrate large sets of functional genomics data in combined scores (CADD, DeepSEA, Eigen, FATHMM-MKL, FunSeq2, GWAVA region model, LINSIGHT, ReMM). In addition, we compared a subset of experiments (10/21) to absolute deltaSVM scores available for specific cell-types (HEK293T, HeLa S3, HepG2, K562, and LNCaP). In cases where an annotation is based on positions rather than alleles, we assumed the same value for all substitutions at each position. The column “Type” assigns each region as either enhancer (E), promoter or ultra conserved element (C). MYC (rs11986220) and MYC (rs6983267) are abbreviated to MYCs1 and MYCs2, respectively.

| Name | T<br>y<br>p<br>e | C<br>A<br>D<br>D | Fun<br>Seq2 | Eigen | FAT<br>HMM<br>-<br>MKL | Deep<br>SEA | LINS<br>IGHT | R<br>e<br>M<br>M | GW<br>AVA<br>[R] | delta<br>SVM | JASP<br>AR<br>[all] | JASP<br>AR<br>[10th<br>P] | Phast<br>Cons | Phylo<br>P | G<br>e<br>r<br>p<br>N |
| --- | --- | --- | --- | --- | --- | --- | --- | --- | --- | --- | --- | --- | --- | --- | --- |
| BCL11A | E | -7 | 0 | -3 | 2 | 3 | -9 | -2 | -2 | - | -1 | 1 | -1 | -1 | -1 |
| F9 | P | -5 | -2 | -3 | 3 | 23 | -2 | 4 | -4 | 20 | 11 | 52 | -4 | 3 | -2 |
| FOXE1 | P | -7 | 4 | 2 | 4 | 5 | 3 | -1 | -8 | 1 | 1 | 0 | -1 | 3 | 0 |
| GP1BB | P | -5 | 13 | 0 | 10 | 25 | 9 | 7 | 28 | - | 28 | 33 | 4 | 5 | 1 |
| HBB | P | 3 | 11 | 15 | 11 | 29 | 12 | 11 | -5 | - | -1 | 5 | 10 | 19 | 5 |
| HBG1 | P | 0 | 11 | 16 | 15 | 26 | 13 | 0 | -3 | - | -2 | 1 | 21 | 12 | -3 |
| HNF4a | P | 15 | 10 | 22 | 18 | 25 | 22 | 14 | 9 | 3 | 11 | 10 | 20 | 15 | 22 |
| IRF4 | E | 43 | 40 | 48 | 29 | 47 | 44 | 30 | 24 | - | 14 | 26 | 31 | 24 | 47 |
| IRF6 | E | 14 | 12 | 19 | 22 | 28 | 14 | 19 | -1 | - | 6 | 2 | 21 | 17 | 4 |
| LDLR.2 | E | 41 | 40 | 58 | 59 | 56 | 56 | 52 | 4 | 15 | -2 | 39 | 57 | 44 | 59 |
| MSMB | P | -1 | 24 | 10 | 14 | 20 | 15 | -9 | 27 | 11 | 23 | 17 | 7 | 0 | 31 |
| MYCs1 | E | -7 | 2 | -2 | 0 | 4 | -9 | -4 | -2 | 6 | 6 | 2 | -1 | -1 | 0 |
| MYCs2 | E | -10 | 2 | -4 | -9 | 12 | -11 | -1 | 2 | 8 | -9 | 7 | -4 | 1 | -10 |
| PKLR 48h | P | 17 | 39 | 25 | 26 | 34 | 41 | 19 | 32 | 21 | 3 | 15 | 23 | 17 | 28 |
| RET | E | 11 | 7 | 13 | 11 | 14 | 5 | 14 | 10 | - | 13 | 17 | 13 | 11 | 11 |
| SORT1 | E | -21 | 28 | -13 | 2 | 6 | -16 | -16 | 16 | 28 | 20 | 29 | -13 | -11 | -28 |
| TCF7L2 | E | -1 | 12 | 9 | 4 | 6 | 5 | -6 | 5 | - | 13 | 15 | 0 | 3 | 17 |
| TERT-GBM | P | 2 | 21 | 20 | 20 | 37 | 11 | 0 | -4 | 12 | 5 | 7 | 16 | 14 | 13 |
| UC88 | C | -4 | 6 | -4 | -3 | 11 | -3 | -7 | 4 | - | 4 | 1 | -2 | -2 | -10 |
| ZFAND3 | E | 37 | 23 | 40 | 40 | 44 | 38 | 30 | 9 | - | 15 | 4 | 41 | 27 | 37 |
| ZRS h13 | E | 12 | 0 | 7 | 7 | 7 | 3 | 5 | -15 | - | 0 | 5 | 1 | 6 | 7 |

Among the ten elements for which sequence-based models were available from deltaSVM, DeepSEA, the top 10th percentile JASPAR predictions and FunSeq2 scores still performed best in predicting absolute effect size (using absolute scores). The deltaSVM models ranked 6^th^ out of 25 measures and were most similar to the number of overlapping top 25^th^ percentile JASPAR predicted motifs (0.73 Pearson correlation). deltaSVM models showed improved performance when correlating expression effects including their directionality (0.55 Spearman correlation for *SORT1*, 0.44 Spearman correlation for *F9*, Supplementary Table 13). This illustrates how sequence-based models can overcome missing gain-of-motif annotations from reference sequence-based predictions.

In summary, we observe that even the best performing computational tools or annotations only explain a small proportion of the expression effects observed in our data; the highest variance explained based on Pearson R^2^ (for *LDLR*) is 0.40, the average across all elements is just 0.02. In these comparisons, we are including a large number of close-to-zero effect estimates affected by experimental noise. Therefore, the analysis is conservative for estimating the predictive power, but including potential functionally neutral effects is critical for assessing how well annotations and scores discriminate between functional and non-functional variants.

## Discussion

Although limited in their naturalness, MPRAs enable a rigorous, quantitative ascertainment of the regulatory consequences of genetic variants. Previous studies applied MPRAs to study common genetic variants in various regions, with the goal of fine-mapping the causal regulatory determinants of GWAS or eQTL associations^69,70^. In contrast, here we selected sequences with known regulatory potential -- and moreover, sequences previously implicated in human disease -- and sought to quantify the consequences of all possible single nucleotide variants on that potential.

Saturating MPRAs uniquely facilitates several kinds of analysis. First, we are able to formally evaluate the distribution of effect sizes in regulatory elements. For example, what proportion of variants are inert, activating or repressing? How atypical is a regulatory variant that results in a two-fold expression change? Although the answers to such questions undoubtedly differs between regulatory elements, the number of elements and variants that were studied allows us to begin to make generalizations. Second, saturating MPRAs facilitate the fine-scale identification of transcription factor binding sites, including ones that may correspond to transcriptional regulators for which ChIP-seq data is not available or that are not well-represented in motif databases. Importantly, it also allows the discovery of binding sites that are created by genetic variants, *i.e.* through activating mutations. Third, we show that the intersection of saturation mutagenesis MPRAs and TF perturbation, *i.e.* our siRNA-based knockdown of GABPA, enables confirmation that a particular TF binds to a particular TFBS. Larger-scale implementations of this approach may facilitate the routine identification of the specific *trans-*acting factors that are responsible for the regulatory potential of each *cis-*acting regulatory element.

Our study and these data have several limitations that merit highlighting. First, we are limited with respect to context, both *cis* and *trans.* To address the former, we used longer sequences than are typical for MPRAs, up to 600 bp, but it remains the case that these are studied on episomal vectors rather than in their native locations. To address the latter, we selected cell lines in which these elements were previously shown to be active, and moreover relevant to the diseases in which these elements were implicated. Nonetheless, previous studies have demonstrated that some regulatory polymorphisms do not always reflect their *in vivo* effects in cell culture-based assays^71^, particularly for developmental genes that show temporal and tissue-specific expression patterns (for example, see results for the IRF6 and ZRS in Supplementary Note 1). A second limitation relates to the reproducibility of measurements for some of the elements studied, and in particular for those with lower basal activity (which we found to be the largest factor impacting reproducibility). Potential approaches to address this in future work include using a stronger minimal promoter (for enhancers) or simply using more complex libraries to further reduce noise (for all elements).

A clear result of our analyses is that although myriad annotations and integrative scores are available, and although some annotations/scores are surprisingly successful in specific cases, no current score consistently performs well in predicting the regulatory consequences of SNVs in the human genome. It is our hope that this dataset of functional measurements for over 30,000 single nucleotide substitution and deletion regulatory mutations in disease-associated regulatory elements will be useful for the field for studying the shortcomings of current tools, and hopefully inspiring their improvement.

In summary, we successfully scaled saturation mutagenesis-based MPRAs to measure the regulatory consequences of tens-of-thousands of sequence variants in promoter and enhancer sequences previously associated with clinically relevant phenotypes. We believe that our experiments provide a rich dataset for benchmarking predictive models of variant effects, an unprecedented database for the interpretation of potentially disease-causing regulatory mutations, and the potential for critical insight for the development of improved computational tools.

## Online Methods

### Selection of target sequences and luciferase assays

Promoter and enhancer sequences of interest (Supplementary Table 1, 2) were amplified from human genomic DNA (Roche 11691112001). All promoters were cloned into pGL4.11b vector [modified from pGL4.11 (Promega) by Dr. Richard M. Myers lab] and most of the enhancers were cloned into the pGL4.23 vector (Promega) that contains a minimal promoter followed by the luciferase reporter gene (Supplementary Table 1, 2). All inserts were confirmed by Sanger sequencing. We measured the relative luciferase activity of the selected promoters and enhancers for the wild-type as well as the saturation mutagenesis library, normalized to the empty vector. For this purpose, HepG2, HEK293T, HeLa, HaCaT, Min6, Neuro-2a, LNCap, SK-MEL-28 were obtained from American Type Culture Collection, and primary gliobastoma (GBM) cell line SF7996^49^ was obtained from UCSF (Dr. Costello’s lab). 2.0 x10^5^ cells were cultured in 96-well plates overnight using standard protocols and were transfected with 100 ng of plasmid bearing the promoter/enhancer sequence (Supplementary Table 1, 2) (along with 10 ng of the Renilla vector to facilitate normalization for transfection efficiency) using X-tremeGENE HP DNA transfection reagent (Roche 06366236001) according to the manufacturer’s protocol. K562, HEL92.1.7, and NIH/3T3 cells were cultured in 24-well plates and transfected using X-tremeGENE HP with 500 ng of the constructed plasmid, along with 50 ng of the Renilla vector. The promoter/enhancer activity was measured using the Dual-Luciferase reporter assay (Promega E1910) on a Synergy 2 microplate reader (BioTek Instruments) following a post-transfection interval that varied by experiment (Supplementary Table 1, 2).

### Construction of MPRA libraries

The cloned enhancer or promoter sequences were amplified in two rounds of PCR. After amplifying once to append universal adaptors for subsequent cloning, a second error-prone PCR with Mutazyme II (GeneMorph II Random Mutagenesis Kit, Agilent 200550) was used to introduce sequence variation. This second round also added a 15 or 20 bp random tag (*i.e.* barcode) contained within an overhanging primer oligo to each construct.

The resulting PCR products were cloned into the respective vector backbone (Supplementary Table 1, 2) without the luciferase reporter via NEBuilder HiFi DNA Assembly (NEB E2621) and transformed into 10-Beta Electrocompetent cells (NEB C3020K). As needed, multiple transformations were pooled and midi-prepped together (Chargeswitch Pro Filter Plasmid Midi Kit, Invitrogen CS31104). Using the vector backbone without the luciferase gene allowed for association of each sequence variant with its newly added tag via sequencing. Once this interim library was determined to have a sufficient complexity and representation of variants, the luciferase gene was inserted between the enhancer/promoter and its tag via a sticky end ligation and transformed again to make the final library for transfection.

### Assignment of tags to sequence variants

The associations between tags and sequence variants created by error-prone PCR were learned by amplifying and deep sequencing of this region of the plasmid library before the luciferase gene was added, *i.e.* while the enhancer/promoters and tags are in close proximity. For short enhancers/promoters, the libraries were amplified with sequencing adaptor primers that captured the cloned sequence with its tag and added the P5/P7 Illumina flow cell sequences. For long enhancers/promoters, a custom subassembly sequencing approach^27^ was used to obtain associated tags and sequences along the targets. Here, libraries were first amplified as before with sequencing adaptors. Some of the full-length product was then subjected to tagmentation via a Nextera library prep (Illumina FC-121-1031). The tagmented products were amplified with a Nextera-specific primer on the P5 end, and a primer containing only the P7 flow cell sequence on the other end. These PCR fragments were size-selected on a 1%-agarose gel into two size bins. Full-length and fragmented libraries were quantitated with a Kapa Library Quantification Kit (Roche 07960140001). Products were run on either an Illumina MiSeq or NextSeq instrument. Full-length and large-size fragment bins were loaded with increased DNA concentration, as these are less efficiently amplified during the cluster generation process. Sequence reads were aligned using BWA-mem v0.7.10-r789^72^ with an increased penalty against local alignments (-L 80) to the Sanger determined references. A minimum coverage of three reads along the whole target was required to include variant calls from bcftools v1.2^73^ for each identified tag. Summary statistics for these assignments are available in Supplementary Table 3.

### Expression of libraries and nucleic acid extraction

For each experiment, about 5 million cells were plated in 15-cm plates and incubated for 24 hours before transfection. Each of three independent cultures (replicates) were transfected with 15 μg of the constructed MPRA libraries using X-tremeGENE HP (Roche 06366236001). After indicated hours (Supplementary Tables 1, 2), cells were harvested, genomic DNA and total RNA were extracted using AllPrep DNA/RNA mini kit (Qiagen 80204). Total RNA was subjected to mRNA selection (Oligotex, Qiagen 72022) and treated with Turbo DNase (Thermo Fisher Scientific AM2238) to remove contaminating DNA.

### RNA interference (TERT promoter)

Following the protocol outlined in Bell et al.^48^, short interfering RNAs (siRNAs) were transfected into GBM cells (SF7996) using DharmaFECT 1 following the manufacturer’s protocol. Briefly, cells were seeded at a density of 30,000 cells/mL in a 96-well plate and 5 million cells in 15-cm plates in parallel. 24-hours post seeding, cells were transfected with 50 nM of siRNA and 0.3 μL of DharmaFECT 1 reagent (Dharmacon T-2001). At 48 hours post-transfection with siRNA, cells were transfected again with the TERT saturation mutagenesis library for another 24 hours before harvesting for genomic DNA and total RNA as described above. To measure siRNA knockdown efficiency, cDNA was generated from the 96-well plate, and qPCR performed (Power SYBR Green Cells-to-Ct kit, Ambion 4402953) to measure mRNA abundance of GABPA and TERT with primer sequences previously used (GABPA: 5’-AAGAACGCCTTGGGATACCCT-3’; 5’-GTGAGGTCTATATCGGTCATGCT-3’; TERT: 5’-TCACGGAGACCACGTTTCAAA-3’, 5’-TTCAAGTGCTGTCTGATTCCAAT-3’; GUSB: 5’-CTCATTTGGAATTTTGCCGATT-3’, 5’-CCGAGTGAAGATCCCCTTTTTA-3’). Relative expression levels (Supplementary Figure 7) were calculated using the deltaCT method against housekeeping gene GUSB as previously described^48^.

### RNA and DNA library preparation

For each replicate, RNA was reverse transcribed with Superscript II (Invitrogen 18064-014) using a primer downstream of the tag, which contained a sample index and a P7 Illumina adaptor sequence. The resulting cDNA was first pooled and then split into multiple reactions to reduce PCR jackpotting effects. Amplification was performed with Kapa Robust polymerase (Roche KK5024) for three cycles, incorporating unique molecular identifiers (UMIs) 10 bp in length. PCR products were cleaned up with AMPure XP beads (Beckman Coulter A63880) to remove the primers and concentrate the products. These products underwent a second round of amplification in eight reactions per replicate for 15 cycles, with a reverse primer containing only P7. All reactions were pooled, run on an agarose gel for size-selection, and then sequenced. For the DNA, each replicate was amplified for three cycles with UMI-incorporating primers, just as the RNA. First round products were then cleaned up with AMPure XP beads, and amplified in split reactions, each for 20 cycles. Reactions were pooled, gel-purified, and sequenced.

### Sequencing and primary processing

RNA and DNA for each of the three replicates was sequenced on an Illumina NextSeq instrument (2×15 or 2×20 bp + 10 bp UMI + 10 bp sample index). Paired-end reads each sequenced the tags from the forward and reverse direction and allowed for adapter trimming and consensus calling of tags^74^. Tag or UMI reads containing unresolved bases (N) or those not matching the designed length were excluded. In the downstream analysis, each tag x UMI pair is counted only once and only tags matching the above obtained assignment were considered (Supplementary Table 4).

### Inferring single nucleotide variant effects

RNA and DNA counts for each replicate were combined by tag sequence, excluding tags not observed in both RNA and DNA of the same experimental replicate. All tags (T) not associated with insertions or multiple base-pair deletions were included in a matrix of RNA count, DNA count, and *N* binary columns indicating whether a specific sequences variant was associated with the tag. We then fit a multiple linear regression model of log2(RNA^j^) ∼ log2(DNA^j^) + N + offset (j ∈ T) and report the coefficients of N as effects for each variant. Further, we fit a combined model across all three experimental replicates (*i* ∈ {1, 2, 3}), where we combine all RNA measures in one column and keep the DNA readouts separated by replicate (i.e. filling missing values with 0):

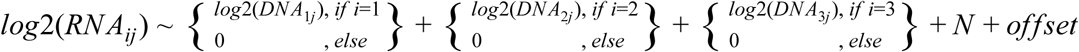

We required a minimum of 10 tags per variant, before considering variant effects in downstream analyses (Supplementary Table 5, 6). While statistically inflated (assumption of independence between tags, correlated counts due to saturation mutagenesis process and no modeling of epistasis effects), we used the p-value for the obtained coefficients as a proxy for the support for each individual non-zero variant effect.

We used the correlation between transfection replication of fitted variant effects divided by the inferred standard deviation as a measure of reproducibility (Supplementary Table 7). We explored major contributors to the reproducibility by Leave One Out Regression models using putative predictors from Supplementary Tables 1-4 (target length, Luciferase fold-change, number of associated tags, proportion wild-type in the library, average mutation rate, number of variants per insert, tags per SNV, tags per 1 bp deletion, assigned DNA and RNA counts, number of wild-type tags, number of variants across tags, average number of variants per tag).

To identify significantly different variants in the siRNA knockdown experiment 95%-confidence intervals were generated from the generalized linear model using the confint.lm method of the stats package in R. Variants with non-overlapping confidence intervals between the experiments TERT-GBM-SiGABPA and TERT-GBM-SiScramble were considered significantly different.

### Annotation and downstream analysis

Base pair positions and single nucleotide variants along the targeted regions were annotated based on GRCh37/hg19 coordinates. CADD scores as well as phastCons, phyloP and GERP++ scores were downloaded from CADD v1.3 [http://cadd.gs.washington.edu/download]. Functional effect scores for FunSeq v2.16 [http://archive.gersteinlab.org/funseq2.1.2/hg19_NCscore_funseq216.tsv.bgz], LINSIGHT [http://compgen.cshl.edu/∼yihuang/tracks/LINSIGHT.bw], and ReMM v0.3.1 [https://charite.github.io/software-remm-score.html] were downloaded for the whole genome and regions of interest extracted. GWAVA scores for the unknown, region and TSS models were calculated for all positions along the targeted regions using available software [ftp://ftp.sanger.ac.uk/pub/resources/software/gwava/v1.0/]. DeepSEA [http://deepsea.princeton.edu/job/analysis/create/] and fathmm-MKL [http://fathmm.biocompute.org.uk/fathmmMKL.htm] scores were retrieved using the respective online web interfaces.

DeltaSVM scores were computed from the precomputed k-mer weights [http://www.beerlab.org/deltasvm] of the initial deltaSVM publication^20^. For each variant the average of all possible k-mer scores of the alternative allele is subtracted from the average of all possible k-mer scores of the reference allele. Not available k-mers in the files are treated as zero. Only precomputed k-mer weights of HEK293T, HeLa S3, HepG2, K562, and LNCaP were available. The model “DukeDnase” was used for experiments in HEK293T (HNF4a, MSMB, MYC rs6983267, TERT). For cell-types with multiple available k-mer models, we selected the best performing model based on our data. This was “DHS_H3Kme1” for HepG2 (SORT1, F9, LDLR) and K562 (PKLR), “UwDnase” for LNCaP (MYC rs11986220) and HeLa S3 (FOXE1). We applied the HEK293T model to our TERT experimental results for the matching cell-line as well as for GBM cells, based on the high correlation observed for these experiments. We obtained higher correlations in GBM cells, probably due to better experimental performance and are using these values in the comparison with other functional scores described above.

We used predicted TFBS available from JASPAR 2018 [http://expdata.cmmt.ubc.ca/JASPAR/downloads/UCSC_tracks/2018/hg19/JASPAR2018_hg19_all_chr.bed.gz]^57^. Scores reported for each motif match were divided by the length of the match. These motif scores of all binding site predictions were combined across the 21 genomic regions to identify thresholds for the 90th (38), 75th (32.5), 50th (27.8182), 25th (24.4167) and 10th (21.4) percentile. To identify factors with the highest number of motifs in each region, we identified the 5 most frequent factors and included additional factors with the same number of motif matches in the region. For visualization, overlapping matches of the same motif were combined and matches on both strands considered only once.

TFBS predictions overlapping respective ChIP-peaks in ENCODE experiments were downloaded from http://compbio.mit.edu/encode-motifs/. TFBSs annotated in the Ensembl Regulatory Build v90 were downloaded from ftp://ftp.ensembl.org/pub/release-90/regulation/homo_sapiens and coordinate converted to GRCh37 using the UCSC liftover program.

## Data access

The sequencing data, obtained tag-to-variant assignments, processed RNA/DNA data including annotations have been submitted to the NCBI Gene Expression Omnibus (GEO; http://www.ncbi.nlm.nih.gov/geo/) under accession number XXXX. The expression effect estimates and further information is available at DOI <u>10.17605/OSF.IO/75B2M.</u>

## Acknowledgements and funding

We thank current and previous members of the Ahituv, Kircher and Shendure laboratories, as well as Daniela Witten, Greg Cooper and Franklin Huang, for valuable discussions and feedback. Our work was supported by the National Human Genome Research Institute (NHGRI) grant numbers 1R01HG006768 (NA & JS) and 1R01HG009136 (JS), NHGRI grant number 1UM1HG009408 (NA & JS), NHGRI and National Cancer Institute grant number 1R01CA197139 (NA & JS), National Institute of Mental Health grant numbers 1R01MH109907 and 1U01MH116438 (NA) and the Berlin Institute of Health (MK & MS). JS is an investigator of the Howard Hughes Medical Institute.

## Author’s contributions

All authors designed the experiments; B.M., C.X., and F.I. performed all wet lab experiments; C.X., J.S., M.K., M.S., and N.A. outlined data analysis; M.K. processed and analysed the high-throughput sequencing data, C.X., M.K., and M.S. performed data analysis; All authors interpreted the experimental results and wrote the manuscript.

## Supplementary information

Supplementary Note 1, Supplementary Figures 1-9, and Supplementary Tables 1-15 are available in the separate Supplementary Information file.

## Competing interests

The authors declare no competing interests.

## References

1. Shendure, J. & Akey, J. M. The origins, determinants, and consequences of human mutations. Science 349, 1478–1483 (2015).

2. Li, X. et al. The impact of rare variation on gene expression across tissues. Nature 550, 239–243 (2017).

3. Chatterjee, S. & Ahituv, N. Gene Regulatory Elements, Major Drivers of Human Disease. Annu. Rev. Genomics Hum. Genet. 18, 45–63 (2017).

4. Maurano, M. T. et al. Systematic localization of common disease-associated variation in regulatory DNA. Science 337, 1190–1195 (2012).

5. Cusanovich, D. A. et al. A Single-Cell Atlas of In Vivo Mammalian Chromatin Accessibility. Cell 174, 1309–1324.e18 (2018).

6. 1000 Genomes Project Consortium et al. A map of human genome variation from population-scale sequencing. Nature 467, 1061–1073 (2010).

7. Lek, M. et al. Analysis of protein-coding genetic variation in 60,706 humans. Nature 536, 285–291 (2016).

8. Levo, M. & Segal, E. In pursuit of design principles of regulatory sequences. Nat. Rev. Genet. 15, 453–468 (2014).

9. Kircher, M. et al. A general framework for estimating the relative pathogenicity of human genetic variants. Nat. Genet. 46, 310–315 (2014).

10. Zhou, J. & Troyanskaya, O. G. Predicting effects of noncoding variants with deep learning-based sequence model. Nat. Methods 12, 931–934 (2015).

11. Ionita-Laza, I., McCallum, K., Xu, B. & Buxbaum, J. D. A spectral approach integrating functional genomic annotations for coding and noncoding variants. Nat. Genet. 48, 214–220 (2016).

12. Shihab, H. A. et al. An integrative approach to predicting the functional effects of non-coding and coding sequence variation. Bioinformatics 31, 1536–1543 (2015).

13. Fu, Y. et al. FunSeq2: a framework for prioritizing noncoding regulatory variants in cancer. Genome Biol. 15, 480 (2014).

14. Ritchie, G. R. S., Dunham, I., Zeggini, E. & Flicek, P. Functional annotation of noncoding sequence variants. Nat. Methods 11, 294–296 (2014).

15. Huang, Y.-F., Gulko, B. & Siepel, A. Fast, scalable prediction of deleterious noncoding variants from functional and population genomic data. Nat. Genet. 49, 618–624 (2017).

16. Smedley, D. et al. A Whole-Genome Analysis Framework for Effective Identification of Pathogenic Regulatory Variants in Mendelian Disease. Am. J. Hum. Genet. 99, 595–606 (2016).

17. Ernst, J. & Kellis, M. Chromatin-state discovery and genome annotation with ChromHMM. Nat. Protoc. 12, 2478–2492 (2017).

18. Chan, R. C. W. et al. Segway 2.0: Gaussian mixture models and minibatch training. Bioinformatics 34, 669–671 (2018).

19. Gulko, B., Hubisz, M. J., Gronau, I. & Siepel, A. A method for calculating probabilities of fitness consequences for point mutations across the human genome. Nat. Genet. 47, 276–283 (2015).

20. Lee, D. et al. A method to predict the impact of regulatory variants from DNA sequence. Nat. Genet. 47, 955–961 (2015).

21. Landrum, M. J. et al. ClinVar: public archive of interpretations of clinically relevant variants. Nucleic Acids Res. 44, D862–8 (2016).

22. Stenson, P. D. et al. The Human Gene Mutation Database: towards a comprehensive repository of inherited mutation data for medical research, genetic diagnosis and next-generation sequencing studies. Hum. Genet. 136, 665–677 (2017).

23. Bejerano, G. et al. Ultraconserved elements in the human genome. Science 304, 1321–1325 (2004).

24. Visel, A. et al. Ultraconservation identifies a small subset of extremely constrained developmental enhancers. Nat. Genet. 40, 158–160 (2008).

25. Patwardhan, R. P. et al. High-resolution analysis of DNA regulatory elements by synthetic saturation mutagenesis. Nat. Biotechnol. 27, 1173–1175 (2009).

26. Patwardhan, R. P. et al. Massively parallel functional dissection of mammalian enhancers in vivo. Nat. Biotechnol. 30, 265–270 (2012).

27. Hiatt, J. B., Patwardhan, R. P., Turner, E. H., Lee, C. & Shendure, J. Parallel, tag-directed assembly of locally derived short sequence reads. Nat. Methods 7, 119–122 (2010).

28. Scholtz, C. L. et al. Mutation -59c-->t in repeat 2 of the LDL receptor promoter: reduction in transcriptional activity and possible allelic interaction in a South African family with familial hypercholesterolaemia. Hum. Mol. Genet. 8, 2025–2030 (1999).

29. Mozas, P. et al. A mutation (−49C>T) in the promoter of the low density lipoprotein receptor gene associated with familial hypercholesterolemia. J. Lipid Res. 43, 13–18 (2002).

30. De Castro-Orós, I. et al. Functional analysis of LDLR promoter and 5’ UTR mutations in subjects with clinical diagnosis of familial hypercholesterolemia. Hum. Mutat. 32, 868–872 (2011).

31. Huang, F. W. et al. Highly recurrent TERT promoter mutations in human melanoma. Science 339, 957–959 (2013).

32. Liu, T., Yuan, X. & Xu, D. Cancer-Specific Telomerase Reverse Transcriptase (TERT) Promoter Mutations: Biological and Clinical Implications. Genes 7, (2016).

33. Killela, P. J. et al. TERT promoter mutations occur frequently in gliomas and a subset of tumors derived from cells with low rates of self-renewal. Proc. Natl. Acad. Sci. U. S. A. 110, 6021–6026 (2013).

34. Vinagre, J. et al. Frequency of TERT promoter mutations in human cancers. Nat. Commun. 4, 2185 (2013).

35. Musunuru, K. et al. From noncoding variant to phenotype via SORT1 at the 1p13 cholesterol locus. Nature 466, 714–719 (2010).

36. Copp, J. N., Hanson-Manful, P., Ackerley, D. F. & Patrick, W. M. Error-prone PCR and effective generation of gene variant libraries for directed evolution. Methods Mol. Biol. 1179, 3–22 (2014).

37. McCullum, E. O., Williams, B. A. R., Zhang, J. & Chaput, J. C. Random mutagenesis by error-prone PCR. Methods Mol. Biol. 634, 103–109 (2010).

38. Inoue, F. et al. A systematic comparison reveals substantial differences in chromosomal versus episomal encoding of enhancer activity. Genome Res. 27, 38–52 (2017).

39. Li, W. H., Wu, C. I. & Luo, C. C. A new method for estimating synonymous and nonsynonymous rates of nucleotide substitution considering the relative likelihood of nucleotide and codon changes. Mol. Biol. Evol. 2, 150–174 (1985).

40. Guo, C. et al. Transversions have larger regulatory effects than transitions. BMC Genomics 18, 394 (2017).

41. Leigh, S. E. A., Foster, A. H., Whittall, R. A., Hubbart, C. S. & Humphries, S. E. Update and analysis of the University College London low density lipoprotein receptor familial hypercholesterolemia database. Ann. Hum. Genet. 72, 485–498 (2008).

42. Liyanage, K. E., Burnett, J. R., Hooper, A. J. & van Bockxmeer, F. M. Familial hypercholesterolemia: epidemiology, Neolithic origins and modern geographic distribution. Crit. Rev. Clin. Lab. Sci. 48, 1–18 (2011).

43. Lind, S. et al. Genetic characterization of Swedish patients with familial hypercholesterolemia: a heterogeneous pattern of mutations in the LDL receptor gene. Atherosclerosis 163, 399–407 (2002).

44. Fouchier, S. W., Kastelein, J. J. P. & Defesche, J. C. Update of the molecular basis of familial hypercholesterolemia in The Netherlands. Hum. Mutat. 26, 550–556 (2005).

45. Dedoussis, G. V. Z. et al. Molecular characterization of familial hypercholesterolemia in German and Greek patients. Hum. Mutat. 23, 285–286 (2004).

46. Horn, S. et al. TERT promoter mutations in familial and sporadic melanoma. Science 339, 959–961 (2013).

47. Rachakonda, P. S. et al. TERT promoter mutations in bladder cancer affect patient survival and disease recurrence through modification by a common polymorphism. Proc. Natl. Acad. Sci. U. S. A. 110, 17426–17431 (2013).

48. Bell, R. J. A. et al. Cancer. The transcription factor GABP selectively binds and activates the mutant TERT promoter in cancer. Science 348, 1036–1039 (2015).

49. Mancini, A. et al. Disruption of the β1L Isoform of GABP Reverses Glioblastoma Replicative Immortality in a TERT Promoter Mutation-Dependent Manner. Cancer Cell 34, 513–528.e8 (2018).

50. Huang, D.-S. et al. Recurrent TERT promoter mutations identified in a large-scale study of multiple tumour types are associated with increased TERT expression and telomerase activation. Eur. J. Cancer 51, 969–976 (2015).

51. Hurst, C. D., Platt, F. M. & Knowles, M. A. Comprehensive mutation analysis of the TERT promoter in bladder cancer and detection of mutations in voided urine. Eur. Urol. 65, 367–369 (2014).

52. Forbes, S. A. et al. COSMIC: somatic cancer genetics at high-resolution. Nucleic Acids Res. 45, D777–D783 (2017).

53. Labussière, M. et al. TERT promoter mutations in gliomas, genetic associations and clinico-pathological correlations. Br. J. Cancer 111, 2024–2032 (2014).

54. Spiegl-Kreinecker, S. et al. Prognostic quality of activating TERT promoter mutations in glioblastoma: interaction with the rs2853669 polymorphism and patient age at diagnosis. Neuro. Oncol. 17, 1231–1240 (2015).

55. Varadi, V. et al. A functional promoter polymorphism in the TERT gene does not affect inherited susceptibility to breast cancer. Cancer Genet. Cytogenet. 190, 71–74 (2009).

56. Nencha, U. et al. TERT promoter mutations and rs2853669 polymorphism: prognostic impact and interactions with common alterations in glioblastomas. J. Neurooncol. 126, 441–446 (2016).

57. Khan, A. et al. JASPAR 2018: update of the open-access database of transcription factor binding profiles and its web framework. Nucleic Acids Res. 46, D260–D266 (2018).

58. Myocardial Infarction Genetics Consortium et al. Genome-wide association of early-onset myocardial infarction with single nucleotide polymorphisms and copy number variants. Nat. Genet. 41, 334–341 (2009).

59. Kathiresan, S. et al. Six new loci associated with blood low-density lipoprotein cholesterol, high-density lipoprotein cholesterol or triglycerides in humans. Nat. Genet. 40, 189–197 (2008).

60. Teslovich, T. M. et al. Biological, clinical and population relevance of 95 loci for blood lipids. Nature 466, 707–713 (2010).

61. Melgar, M. F., Collins, F. S. & Sethupathy, P. Discovery of active enhancers through bidirectional expression of short transcripts. Genome Biol. 12, R113 (2011).

62. Treiber, T. et al. Early B cell factor 1 regulates B cell gene networks by activation, repression, and transcription- independent poising of chromatin. Immunity 32, 714–725 (2010).

63. Li, R. et al. Dynamic EBF1 occupancy directs sequential epigenetic and transcriptional events in B-cell programming. Genes Dev. 32, 96–111 (2018).

64. Pollard, K. S., Hubisz, M. J., Rosenbloom, K. R. & Siepel, A. Detection of nonneutral substitution rates on mammalian phylogenies. Genome Res. 20, 110–121 (2010).

65. Siepel, A. et al. Evolutionarily conserved elements in vertebrate, insect, worm, and yeast genomes. Genome Res. 15, 1034–1050 (2005).

66. Davydov, E. V. et al. Identifying a high fraction of the human genome to be under selective constraint using GERP++. PLoS Comput. Biol. 6, e1001025 (2010).

67. Zerbino, D. R., Wilder, S. P., Johnson, N., Juettemann, T. & Flicek, P. R. The ensembl regulatory build. Genome Biol. 16, 56 (2015).

68. Kheradpour, P. & Kellis, M. Systematic discovery and characterization of regulatory motifs in ENCODE TF binding experiments. Nucleic Acids Res. 42, 2976–2987 (2014).

69. Ulirsch, J. C. et al. Systematic Functional Dissection of Common Genetic Variation Affecting Red Blood Cell Traits. Cell 165, 1530–1545 (2016).

70. Tewhey, R. et al. Direct Identification of Hundreds of Expression-Modulating Variants using a Multiplexed Reporter Assay. Cell 172, 1132–1134 (2018).

71. Cirulli, E. T. & Goldstein, D. B. In vitro assays fail to predict in vivo effects of regulatory polymorphisms. Hum. Mol. Genet. 16, 1931–1939 (2007).

72. Li, H. & Durbin, R. Fast and accurate long-read alignment with Burrows-Wheeler transform. Bioinformatics 26, 589–595 (2010).

73. Li, H. A statistical framework for SNP calling, mutation discovery, association mapping and population genetical parameter estimation from sequencing data. Bioinformatics 27, 2987–2993 (2011).

74. Kircher, M. Analysis of high-throughput ancient DNA sequencing data. Methods Mol. Biol. 840, 197–228 (2012).

