## Supplementary Information for "Saturation mutagenesis of disease-associated regulatory elements"

#### Supplementary Note 1

All graphs showing inferred variant effects for SNVs from promoters or enhancers with available annotation data are shown in Supplementary Figure 8 or Supplementary Figure 9, respectively.

All correlations between replicates are shown in Supplementary Table 7. An overview of the promoter elements is in Supplementary Table 1 and Supplementary Table 2 contains all enhancer elements.

All MPRA results for disease-associated promoter or enhancer mutations referenced in the text are provided in Supplementary Table 14 or Supplementary Table 15, respectively.

##### *Promoters (in alphabetical order)*

##### Factor IX (F9)

Hemophilia B Leyden is an X-linked bleeding disorder characterized by low plasma levels of blood coagulation Factor IX (F9) during childhood. Single point mutations in the F9 promoter at positions NM_000133.3:c.-55G>C, c.-50T>G, c.-49T>A, c.-35G>A/C, c.-34A>T, c.-22 T>C and c.-17A>G/C have been found to be associated with this disease (Supplementary Table 14)[1,2]. The hepatocyte nuclear factor 4 alpha (HNF4A) is thought to bind to this promoter at positions -63 to -39 to activate F9[1,2] and both c.-55G>C and c.-49T>A are thought to reduce promoter activity due to aberrant HNF4A binding. In our MPRA results, carried out in HepG2 cells, we did not observe a significant reduction of promoter activity due to mutations in the HNF4A binding region (-63 to -39). We believe that this is mostly due to low barcode coverage at this region. However, we did observe that variants (c.-51T>A, c.-51T>C, c.-50T>A, c.-49T>A) in the core CTTTG sequence of the HNF4A motif (JASPAR ID MA0114.2) led to reduced promoter activity. We also found another region, positions -235 to -222, where mutations led to a significant reduction in promoter activity. Analysis of ENCODE motifs[3] of this region suggests that it is associated with Ets-related factors. In addition to the repressing variants, we observed activating variants in F9, several of which increase the transcription level up to 200%. These include mutations c.-25T>A and c.-25T>G, which are located in another annotated HNF4A motif, with both variants leading to a stronger motif. Overall, our results showed a high correlation between replicates (0.88 Pearson correlation; Supplementary Table 7).

#### FOXE1

A SNP, rs1867277 (NM_004473.3:c.-283G>A), in the promoter of the Forkhead Box E1 (*FOXE1*) gene was found to be associated with papillary thyroid cancer susceptibility[4]. The cancer-associated allele is thought to recruit the upstream transcription factor (USF) 1 and 2. Transfection studies in HeLa cells co-transfected with USF1 and USF2 showed that the cancer-associated allele (A) led to a significant increase in promoter activity (~8 fold versus empty vector) compared to the unassociated allele (~3.5 fold versus empty vector)[4]. Our MPRA for this promoter was carried out in HeLa cells along with the co-transfection of USF1 and USF2. We did not observe a significant effect on promoter activity for rs1867277 (Supplementary Table 14). Overall, we only observed that 6 out of 1923 variants with a minimum of 10 associated tags showed a significant effect (p-value $< 10^{-5}$) on promoter activity, but with low expression effects. Our three technical replicates also had the lowest correlation of all MPRA experiments (0.16 Pearson correlation; Supplementary Table 7), precluding our ability to provide definitively interpretable results for this promoter.

## GP1BB

Bernard–Soulier syndrome (BSS) is a rare autosomal recessive bleeding disorder characterized by defects of the GPIb-IX-V complex, a platelet receptor for von Willebrand factor (VWF). Most of the mutations identified in the genes encoding for the GP1BA (GPIbα), GP1BB (GPIbβ), and GP9 (GPIX) subunits prevent expression of the complex at the platelet membrane or its interaction with VWF. A mutation at the *GP1BB* promoter, NM_000407.4:c.-160C>G, is thought to disrupt the second GATA consensus binding site, and was shown to decrease promoter activity by 84% (Supplementary Table 14)[5]. In our MPRA, which was carried out in HEL 92.1.7 cell lines (erythroblasts originating from human bone marrow), we also observed a reduction in promoter activity for this mutation (52% promoter activity compared to baseline). We also saw a significant reduction across the whole GATA binding site. Several additional regions led to reduced promoter activity. For example, chr22:19,710,886-19,710,900 (GRCh37), which is annotated by JASPAR to have binding sites for Ets-related factors[6], showed significant reduction of promoter activity due to various mutations. Additional regions such as chr22:19,710,912-19,710,923, chr22:19,710,940-19,710,949 or chr22:19,711,001-19,711,032 (all GRCh37) also encompassed several mutations that led to significant reduction in promoter activity. Overall, we observed a good correlation (0.88 Pearson correlation; Supplementary Table 7) between replicates for this MPRA.

#### HBB

Mutations in the the Hemoglobin Subunit Beta (*HBB*) gene cause beta thalassemias ($\beta_0$ or $\beta_+$), characterized by reduced hemoglobin levels that lead to lower oxygen levels in the body. The severity of the disease depends on the nature of the mutation. There are 31 clinical variants in the *HBB* promoter that were reported as disease causing, based on the Leiden Open Variation Database (https://lovd.bx.psu.edu/home.php?select_db=HBB)[7]. In our MPRA, which was carried out in in HEL 92.1.7 cells (erythroblasts originating from human bone marrow), 17 out of 31 clinical variants were found to have a significant effect on promoter activity (p-value < $10^{-5}$) (Supplementary Table 14). We observed several blocks where mutations led to reduction in activity, possibly due to transcription factor binding sites. For example, in positions c.-142 to c.-136 there is a known binding site for the erythroid Krüppel factor (EKLF), a zinc-finger transcription factor that plays a critical role in erythropoiesis and regulation of β-globin switching[8]. We observed a decrease in expression of about 12%-19% for previously reported nucleotide variants within this binding site (Supplementary Table 14). In contrast, the promoter mutation NM_000518.4:c.-18C>G, which has been previously shown to decrease steady state levels of mRNA to 85.2%[9] and is thought to disrupt the binding and transactivation of EKLF (causing mild β+ thalassemia[8]), did not show a significant effect in our study (Supplementary Table 14). We identified 12 promoter activating variants (showing greater than 20% upregulation in promoter activity), for example variant c.-99C>G showed an 128% increase in activity (Supplementary Table 14) and could potentially have a strong effect on *HBB* expression. Overall, we observed a good correlation between technical replicates (0.77 Pearson correlation; Supplementary Table 7). There could be several reasons why some of our MPRA results don't completely match previous results for these clinically associated variants. Our p-value minimum threshold might be too stringent to allow us to characterize these mutations (e.g. if we use a p-value threshold of 0.01, 23 out of the 31 variants show a significant effect on promoter activity), the clinical association might be incorrect and/or experimental differences could be contributing to these differences.

**HBG1**

The Hemoglobin Subunit Gamma 1 gene (*HBG1*) is a component of fetal hemoglobin, which is later replaced by adult hemoglobin at birth. Fourteen mutations in the promoter of this gene (NM_000559.2:c.-264C>T, c.-255C>T, c.-255C>G, c.-251T>C, c.-250C>T, c.-249C>T, c.-248C>G, c.-228T>C, c.-211C>T, c.-170G>A, c.-167C>T, c.-167C>G, c.-167C>A, c.-167A>C) have been identified in patients with hereditary persistence of fetal hemoglobin (HPFH; OMIM 142200; Supplementary Table 14). Work carried out in human erythroleukemia cells (K562 and HEL) has identified several proteins that may bind to the *HBG1* promoter and either activate or repress its activity[10]. For example, gamma CAAT is thought to bind to duplicated CCAAT box sequences (c.-168 to c.-164, c.-141 to c.-137) and activate activity. The c.-170G>A mutation is thought to increase the affinity of interaction between gamma CAAT and its cognate site. Two proteins, gamma CAC1 and gamma CAC2, bind a CACCC sequence located c.-197 to c.-194 from the TSS. Gamma OBP, binds an octamer sequence ATGCAAAT located c.-235 to c.-228 from the TSS. c.-228T>C mutates the eighth base of this octamer leading to a reduction of Gamma OBP binding and upregulation of *HBG1* expression, suggesting that OBP may act as a repressor in this context. The Sp1 transcription factor (SP1) is thought to function as a transcriptional activator, binding to positions c.-266 to c.-244. The c.-255C>G mutation is predicted to enhance SPI binding activity (5 to 10 fold-higher affinity for SP1) while c.-255C>T decreases the similarity to the SP1 recognition site and is associated with a more moderate increase in *HBG1* expression[10,11].

We carried out MPRA on the *HBG1* promoter in HEL 92.1.7 cells (erythroblasts originating from human bone marrow) obtaining high reproducibility between technical replicates (0.92 Pearson correlation; Supplementary Table 7). We detected multiple blocks where mutations lead to reduced promoter activity, suggesting they serve as binding sites for activating transcription factors. We saw a significant repressive activity for mutations within the CAAT, CCAAT and CACCC blocks. However, we did not observe a continuous block of significant variants in our MPRA for the ATGCAAAT octamer sequence (c.-235 to c.-228) and the SP1 element (c.-266 to c.-244). Two blocks, one unannotated as a GAAAATAA sequence from c.-84 to c.-77 and the other is the CCAAT box from c.-141 to c.-137, showed a strong correlation with PhastCons vertebrate conservation scores.

As for individual mutations, we observed a significant change in promoter activity (p-value $< 10^{-5}$) for 6 (c.-255C>T, c.-251T>C, c.-228T>C, c.-211C>T, c.-167C>T, c.-167C>A) of the 14 disease-associated mutations (Supplementary Table 14). For example, c.-228T>C showed a strong increase in expression of over 50% and c.-167C>T led to a 65% reduction in promoter activity. Similar to *HBB*, these results could be due to: 1) Our p-value minimum threshold being too stringent (if we use a p-value threshold of 0.01, 8 out of the 14 variants show a significant effect on promoter activity); 2) The clinical association might be incorrect; 3) Experimental differences could all be contributing to these differences 4) Another limitation of our results for this promoter could be due to its regulation being driven by a complex pattern of repressor and activator binding which are temporally expressed.

##### HNF4A P2 promoter

Mutations in the Hepatocyte Nuclear Factor 4 Alpha (*HNF4A*) gene lead to maturity-onset diabetes of the young (MODY), a monogenic autosomal dominant subtype of early-onset diabetes mellitus caused by defective insulin secretion of pancreatic beta cells[12,13]. *HNF4A* has two characterized promoters, P1 and P2. We focused on the distant upstream P2 promoter, which is 46 kb 5' to the P1 promoter and is thought to be the major promoter regulating *HNF4A* pancreatic beta and hepatic cell expression[12]. Mutations in this promoter (NM_175914.4:c.-136A>G, c.-146T>C, c.-169C>T, c.-181G>A, c.-192C>G) have been associated with MODY[80,81] and are thought to affect the binding of various transcription factors and reduce promoter activity. Transfection assays with various mutations have identified functional binding sites for *HNF1A*, *HNF1B*, pancreatic and duodenal homeobox 1 (PDX1) and other transcription factors known to be associated with MODY[12]. They also showed that mutations c.-136A>G and c.-169C>T lead to reduced promoter activity compared to the wild-type sequence in HEK293 cells[12,13].

We carried out MPRA in HEK293T cells and observed an average Pearson correlation of 0.89 across replicates (Supplemental Table 7). Out of the five MODY-associated mutations, we only observed a significant effect for c.-192C>G, which increased promoter activity by 20% (Supplementary Table 14). For c.-146T>C we observed a marginal reduction in activity (4%). Of interest were two variants, c.-6T>G and c.-138T>C, which increased activity up to 200% and might create strong new TF binding sites. Only c.-138T>C showed an overlap with motifs in ENCODE[3] (ONECUT1/2/3), with this variant causing a disruption in the binding site (position 7 of the MA0756.1 JASPAR[6] motif; compare with Supplementary Figure 8).

#### MSMB

A SNP, rs10993994, in the promoter (-57 bp from TSS) of the Microseminoprotein Beta (*MSMB*) gene was found to be associated with prostate cancer risk in two separate GWAS[14,15] and showed differential reporter activity[16,17]. The risk allele, T, of this SNP reduced promoter activity to 30% and 13% compared to the unassociated allele in HEK293T human embryonic kidney cells and prostate cancer cell lines (LNCaP), respectively[18,19]. We carried out MPRA for this promoter in HEK293T cells line, and observed a good correlation between replicates (0.88 Pearson correlation; Supplementary Table 7). For rs10993994, we obtained a 15% increase in promoter activity for the risk allele (T) instead of reduction in promoter activity as observed in LNCaP cells (Supplementary Table 14). This is likely due to the difference in cell lines used. The *MSMB* promoter has two potential androgen response element half sites (TGTTCT) located -113 to -118 bp 5' to the TSS and another is 23 bp upstream of SNP rs10993994[18] (chr10:51,549,467-51,549,472; GRCh37), which are likely to only get activated in LNCap cells but not in HEK293T cells. We observe multiple blocks that showed significant effects on promoter activity, which align with ENCODE motifs and conservation at the 3' half, but not on the other half (Supplementary Figure 8).

#### PKLR

Three different single nucleotide mutations (NM_000298.5:c.-324T>A, c.-83G>C, c.-72A>G) and a polymorphic deletion c.-248delT in the promoter of the Pyruvate Kinase L/R (*PKLR*) gene have been found in patients with pyruvate kinase (PK) deficiency[20,21], which leads to anemia. Functional analysis for both rat and human *PKLR* promoters revealed four conserved motifs (CAC/SP1, PKR-RE, GATA1) within the first 250 bp upstream region[21]. Both c.-72A>G and c.-83G>C significantly reduced promoter activity. c.-72A>G has been attributed to disruption of the consensus binding motif for GATA-1 at c.-69 to c.-74. c.-83G>C was shown to be a part of a core binding motif (CTCTG) of a novel regulatory element (PKR-RE1) in the erythroid-specific promoter of *PKLR*, in close proximity to a regulatory GATA-1 binding site. c.-324T>A did not show an effect on promoter activity, c.-248delT turned out to be a non-functional polymorphism[20,21].

We carried out MPRA for this promoter using two different post-transfection time points (24 and 48 hours) in K562 cells and observed a good correlation between replicates (0.86 and 0.91 Pearson correlation for 24hr and 48hr respectively; Supplementary Table 7). We observed a significant reduction in promoter activity for two of the disease causing mutations, c.-72A>G (24h=-1.77, 48h=-2.40) and c.-83G>C (24h=-1.25, 48h=-2.19). For c.-324T>A and c.-248delT we did not observe a significant effect which is consistent with the previous functional study[20,21]. Around 15% of all tested variants showed a significant effect, with the majority of these variants being located either near the 3' or 5' end, suggesting that these two regions are important for *PKLR* promoter activity. All positions at the GATA1 conserved site from c.-73 to c.-70 had significant repressing variants in both experiments. The same repressing effects can be seen for the two conserved CAC/SP1 sequences (c.-119 to c.-114 and c.-107 to c.-103) and the PKR-RE motif (c.-87 to c.-83). In general, annotated motifs align well with the significant variant effects on *PKLR* (Supplementary Figure 8). This experiment had one of the highest correlation for JASPAR motifs (Supplementary Table 12, 13), compared to all tested promoter elements.

##### *Enhancers (in alphabetical order)*

##### BCL11A

B cell CLL/lymphoma 11A (*BCL11A*) is a transcription factor that regulates hemoglobin switching and has a characterized erythroid enhancer where common genetic variation has been associated with fetal hemoglobin (HbF) levels[22]. Saturation mutagenesis of the *BCL11A* enhancer via CRISPR-Cas9 guide RNA libraries uncovered key functional regions in this enhancer and implicated DNase hypersensitive site (DHS) +58 to have the most significant effect on *BCL11A* expression and HbF levels[23]. We thus used DHS +58 for our MPRA, which was carried out in the HEL92.1.7 erythroblast cell line. For the *BCL11A* experiments, we did not observe high correlations between replicates (0.38 Pearson correlation; Supplementary Table 7), possibly due to low fold activation by this enhancer in our assay/cell line (2.5 fold compared to empty vector) and a high mutation rate from the error-prone PCR step (creating an average of 6 variants per construct and very few wild-type sequences; factors that negatively impact model fit). Only 3% of the tested variants had a significant effect (p-value < $10^{-5}$). The average $\log_2$ expression effect of all variants was only 0.064 (4.5%) and 0.21 (16%) for the subset of significant variants. The low activation we observed for this enhancer could be due to the use of different cells lines. HEL92.1.7 is an established erythoblast cell line collected from bone marrow. HUDEP-2[24], which was used for the CRISPR-Cas9 assays, are human erythroid progenitors derived from the umbilical cord and are thought to be more similar to adult erythroid cells. It could also be due to differences in experimental methodologies, i.e.

endogenous mutations via CRISPR-Cas9 that assay phenotypic changes versus MPRAs that test the ability of the sequence to drive regulatory activity.

#### IRF4

A SNP, rs12203592, within an intron of the gene encoding the Interferon Regulatory Factor 4 (*IRF4*), a transcription factor involved in pigmentation, is strongly associated with sensitivity of skin to sun exposure, freckles, blue eyes and brown hair color[25]. IRF4 is activated by the melanocyte master regulator melanocyte inducing transcription factor (MITF) along with the transcription factor AP-2 alpha (TFAP2A). The pigmentation-associated allele, rs12203592-T, is thought to disrupt a TFAP2A binding site and was shown to reduce enhancer activity[25]. We carried out saturation-based MPRA for this enhancer in human melanoma SK-MEL-28 cells and observed strong correlations between technical replicates (0.99 Pearson correlation; Supplementary Table 7). The rs12203592-T allele led to a significant reduction in enhancer activity, 36% compared to wild-type (Supplementary Table 15). Overall, we observed that 37% of the variants had a significant effect on enhancer activity. Conservation seems to be an informative measurement for activity and *IRF4* has one of the best agreement between different annotations and absolute expression effects (see Results and Supplementary Table 12, 13).

#### IRF6

A SNP, rs642961, in an Interferon Regulatory Factor (IRF6) associated-enhancer was shown to confer an 18% attributable risk for isolated cleft lip[26]. This variant is part of a common haplotype, rs76145088 (G>A), rs642961 (G>A) and rs77542756 (A>G) and the AAG associated-haplotype is thought to disrupt AP-2α binding sites and increase enhancer activity compared to the unassociated haplotypes in human foreskin keratinocyte HFK cells[26]. However, this difference did not reach statistical significance. Saturation based MPRA for this IRF6 enhancer was done in the human keratinocyte HaCaT cells and obtained a strong correlation between technical replicates (0.96 Pearson correlation; Supplementary Table 7). We observed that around 19% of all variants show a significant effect on enhancer activity. We did not observe a significant effect on enhancer activity for rs642961 and rs77542756 by themselves (Supplementary Table 15). However, the region where rs642961 and rs77542756 were located (chr1:209,989,152-209,989,314; GRCh37) had several activating mutations. For SNP rs76145088 our MPRA detected a significant repression by 18% (Supplementary Table 15). We could not detect an enrichment of significant variants at the AP-2α binding sites and because of the design of our MPRA, the associated-haplotype AAG is not covered. IRF6 had several conserved sites, but most of them do not align with blocks of significant expression effects (Supplementary Figure 9), leading to low correlation of conservation and functional scores (Supplementary Table 12, 13). Of note, although rs642961 shows minor effects on reporter gene expression in cell culture, it is possible that it has significantly larger or even opposing effects on IRF6 expression in vivo in its native chromosomal context and/or in the presence of trans-acting variants[26].

**MYC rs6983267 and rs11986220**

Activation of the MYC proto-oncogene, bHLH transcription factor (*MYC*) gene is a hallmark of cancer initiation and maintenance. Several GWAS reported association between the G-allele of rs6983267 and increased risk for various types of cancers such as colorectal and prostate cancer[27–29]. Studies have also shown that a region including rs6983267 has enhancer activity and interacts with the proto-oncogene *MYC* during early stages of colorectal cancer development[30,31]. rs11986220 resides ~100 kb away from rs6983267, and is associated with prostate cancer risk independent of rs6983267[32]. rs11986220 resides within a FOXA1 binding site, and the prostate cancer risk allele (A) is thought to facilitate both stronger FOXA1 binding and androgen responsiveness[32]. To study the individual variant effects of these SNPs and their surrounding sequences, we generated saturation mutagenesis libraries of 600 bp around rs6983267 and 464 bp around rs11986220. We tested the MYC rs6983267 MPRA library in human embryonic kidney cells, HEK293T, adding 20mM LiCl to activate the Wnt pathway and the MYC rs11986220 in human prostate adenocarcinoma LNCaP cells grown in a medium with 100nM Dihydrotestosterone to stimulate androgen activity.

For MYC rs6983267, we observed a strong correlation between replicates (0.75 Pearson correlation; Supplementary Table 7). We did not see a change in activity (p-value of coefficient 0.05; $\log_2$ expression effect -0.08 / 95% activity) due to the cancer-associated rs6983267 allele (T) (Supplementary Table 15). We observed a stretch of nucleotides that led to a reduction in enhancer activity at chr8:128,413,089-128,413,107 (GRCh37). This region correlates with an annotated CTCF binding site as determined by ERB v90[33]. Of note, two variants, chr8:128,413,279C>G and chr8:128,413,289G>T (GRCh37) led to a strong increase in enhancer activity by 86% and 84%.

For MYC rs11986220, we did not obtain a good correlation between replicates (0.31 Pearson correlation; Supplementary Table 7). rs11986220 showed a 14% repressing activity, but was not statistically significant (Supplementary Table 15). In total, only 11 out of 1685 variants had an expression effect on enhancer activity at a significance level of $10^{-5}$.

**RET**

A common non-coding variant, rs2435357, within an enhancer in intron 1 of the gene Rearranged During Transfection Proto-oncogene (*RET*) is significantly associated with Hirschsprung disease (HSCR) susceptibility[34]. The disease-associated variant leads to a significant reduction in reporter activity (6-8 fold) compared to the unassociated allele in

Neuro-2a cells[34]. Saturation mutagenesis based MPRA was carried out for this enhancer in Neuro-2a cells, with the library cloned into the pGL3 vector, similar to the experiment that originally tested this enhancer[34]. Overall, we observed a reasonable correlation between technical replicates (0.71 Pearson correlation; Supplementary Table 7). We did not detect a significant effect on enhancer activity for rs2435357 (Supplementary Table 15). However, this SNP is located within a sequence block that has multiple variants with significant activating and repressing effects (chr10:43,582,096-43,582,232; GRCh37). Both rs2506005 and rs2506004, which reside 76 bp upstream and 217 bp downstream of rs2435357 respectively, and are in linkage disequilibrium with this SNP, also showed no significant effect on enhancer activity. A region located towards the 3'-end of the sequence (chr10:43,582,390-43,582,526; GRCh37) is enriched in variants that led to differential enhancer activity.

##### TCF7L2

GWAS for type-2 diabetes (T2D) found a significant association with SNP rs7903146, located within an intron of the gene Transcription Factor 7 Like 2 (*TCF7L2*). The rs7903146 risk T allele is associated with impaired insulin secretion, incretin effects and enhanced rate of hepatic glucose production[35]. Luciferase assays of both alleles in mouse pancreatic beta MIN6 cells exhibited significant allele-specific differences with the T allele leading to a three fold stronger enhancer activity compared to the non-risk allele (Supplementary Table 15)[102]. We carried out mutation saturation based MPRA for this enhancer in MIN6 cells, which covered over 99% of all mutations in the tested region (Supplementary Table 3) and obtained high variant effect correlations between replicates (0.76 Pearson correlation; Supplementary Table 7). Functional and conservation scores do not correlate well with the observed variant effects, but we observed some correlation with JASPAR motifs (Supplementary Table 12, 13). We do observe a significant effect on enhancer activity for rs7903146, even though only a 19% expression increase.

### UC88

Ultraconserved elements (UCEs) are sequences greater than 200 bp in length that are identical between human, mouse and rat[36]. Half of the UCEs were shown to be functional enhancers in developing mouse embryos, the majority of which are active in the central nervous system[37]. UC88 is located on chr2:162,094,920-162,095,508 (GRCh37) and was shown to be a robust forebrain enhancer at mouse embryonic day E11.5 (hs416; VISTA enhancer browser[38]). UC88 was tested for enhancer activity in mouse neuroblastoma Neuro-2a cells and showed significant enhancer activity (9.3 fold compared to empty vector). Mutation saturation based MPRA was performed on UC88 in Neuro-2a cells. Replicates had a lower replicate correlation (0.64 average Pearson coefficient); however, this increased to 0.9 for significant variants (using a lenient p-value threshold of < 0.01) (Supplementary Table 7). This limited our interpretability to 10% (196 out of 1964, p-value

< 0.01) of all measured variants in UC88, not allowing us to see clear patterns regarding known motifs or other annotations (Supplementary Figure 9). Overall, we obtained a low correlation of functional, conservation and motif scores (Supplementary Table 12, 13).

##### ZFAND3

A Type 2 Diabetes GWAS found an associated with rs58692659, which is ~10 kb upstream to the zinc finger AN1-type containing 3 (*ZFAND3*) gene[39]. The rs58692659 SNP maps to an open chromatin region, identified previously by both islet FAIRE-seq and ChIP-seq for H2A.Z, and is thought to be bound by several beta cell transcription factors, including PDX1, FOXA2, NKX2.2 NKX6.1 and NEUROD1. Further TFBS analyses and EMSA suggest that the T allele alters the binding of NEUROD1[39]. Luciferase assays showed reduced enhancer activity (~5 fold) for the rs58692659 risk allele (T) compared to the C allele in mouse MIN6 beta cells[39]. We carried out mutation saturation based MPRA on this enhancer in MIN6 cells and obtained high variant effect correlations between replicates (0.89 Pearson correlation; Supplementary Table 7). For rs58692659, we observed a significant reduction in enhancer activity (70% residual activity) for the disease-associated allele (T) which is consistent with the previous reported luciferase results. Furthermore, we found that rs58692659 resides in a region (chr6:37,775,451-37,775,758; GRCh37) that with numerous mutations that lead to either a reduction or increase in enhancer activity (Supplementary Table 15). This region is highly conserved in vertebrates and contains predicted binding sites for the islet-enriched transcription factors RFX4, MAX and NHLH1[39] as annotated by JASPAR 2018[6]. Due to the high correlation of conservation, the tested functional scores result in high correlations as well (Spearman correlation ~0.4; Supplementary Table 12, 13).

##### ZRS

Mutations in the Sonic Hedgehog (*SHH*) limb enhancer, called the zone of polarizing activity regulatory sequence (ZRS), lead to a wide array of limb malformations[40]. These include polydactyly, triphalangeal thumb (TPT), syndactyly, tibial hypoplasia and Werner mesomelic syndrome (OMIM #188770)[41–46]. Several of these mutations were tested for enhancer activity using mouse transgenic assays, with the majority showing ectopic expression compared to the wild-type sequence[40]. *In vitro* assays were also done for a 1.7 kb version of this enhancer in NIH3T3 mouse fibroblasts which showed higher enhancer activity upon co-transfection of homeobox d13 (*Hoxd13*) and heart and neural crest derivatives expressed 2 (*Hand2*). As we were limited in length due to technical reasons, we first tested a 485 bp ZRS sequence that encompasses the majority of disease causing mutations in this *in vitro* system. We observed a significant fold change for co-transfections with Hoxd13 by itself (4.2 fold) or along with Hand2 (2.0 fold) compared to the empty vector. We thus set out to do saturation mutagenesis MPRA in NIH3T3 cells using both conditions (i.e. Hoxd13 or Hoxd13+Hand2).

We observed reasonable correlations between technical replicates (0.76 Pearson correlation for both Hoxd13 or Hoxd13+Hand2; Supplementary Table 7). Overall, we did not detect a high fraction of significant variant effects and those that were significant were distributed over the entire region with stronger effects towards the 3' end. For most disease-causing mutations, we did not observe a significant effect, likely due to technical reasons. These could be due to this enhancer having important spatial and temporal activities in the developing limb that are likely not represented in an *in vitro* setting that uses a single cell type along with the co-transfection of additional factors. However, we did observe an effect for some known mutations. These include chr7:156,584,153T>C (GRCh37; ZRS417), which resulted in 16% and 9% increase with Hoxd13 and Hoxd13+Hand2 respectively (Supplementary Table 11), and is known to cause a more severe limb phenotype which includes polydactyly of the four extremities and bilateral tibial deficiency[44]. Another interesting mutation is chr7:156,584,285C>A (GRCh37; ZRS285) that led to 17% and 13% higher enhancer activity with Hoxd13 and Hoxd13+Hand2 respectively (Supplementary Table 15), and is known to cause polydactyly in Silkie chickens[47].

#### Supplementary Figures

##### *Supplementary Figure 1*

**A**

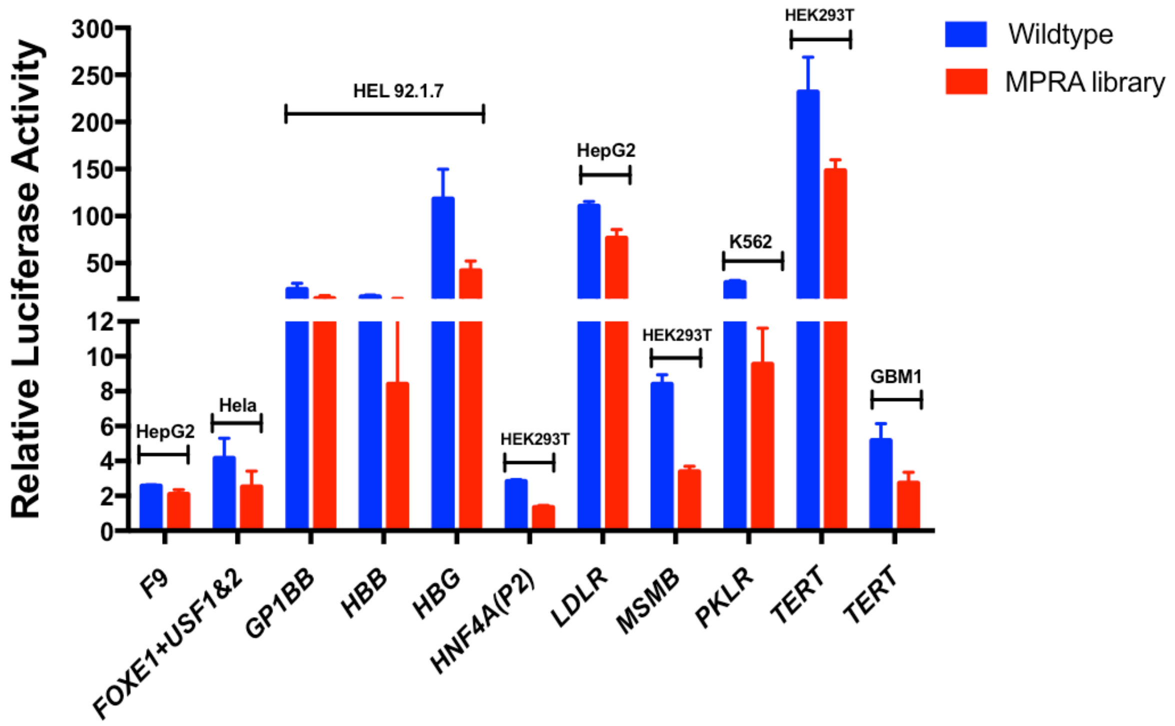

**B**

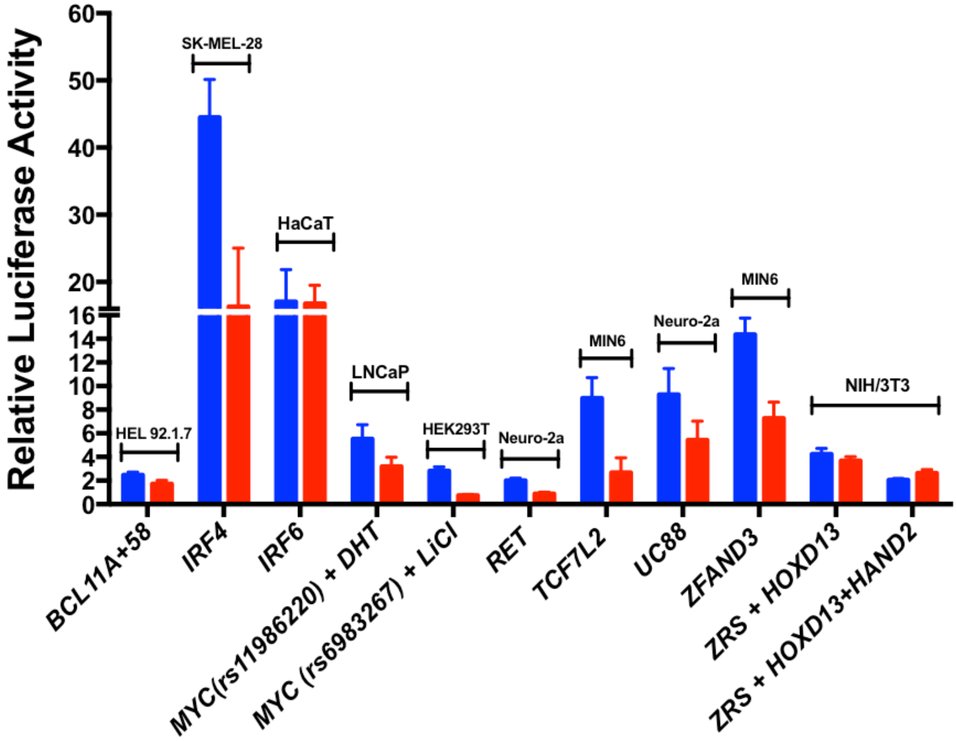

**Promoter and enhancer luciferase assays**. Bar charts representing the relative luciferase activity of the 10 selected promoters (A) and 10 selected enhancers (B) for either the wild-type (blue) or the saturation mutagenesis library (mutant library; red) normalized to the empty vector. In (A), for *FOXE1*, 7.5ug of upstream transcription factor (USF) 1 and 2 were co-transfected along with the vector/library. All promoters activities were measured after 24hr transfection except for *PKLR* which was measured after 48hr. In (B), for *ZRS,* 3.75ug *HOXD13* or *HOXD13* plus *HAND2,* were co-transfected along with the vector/library. For *MYC* (rs11986220), 10nM dihydrotestosterone (DHT) were also co-transfected. For *MYC* (rs6983267), 20mM of LiCl was added after 24hr to induce Wnt pathway activity and luciferase activity was measured at 32 hours. For all other enhancers, luciferase activity was measured 24 hours after transfection. All results are mean ± standard deviation of at least 3 independent experiments.

***Supplementary Figure 2***

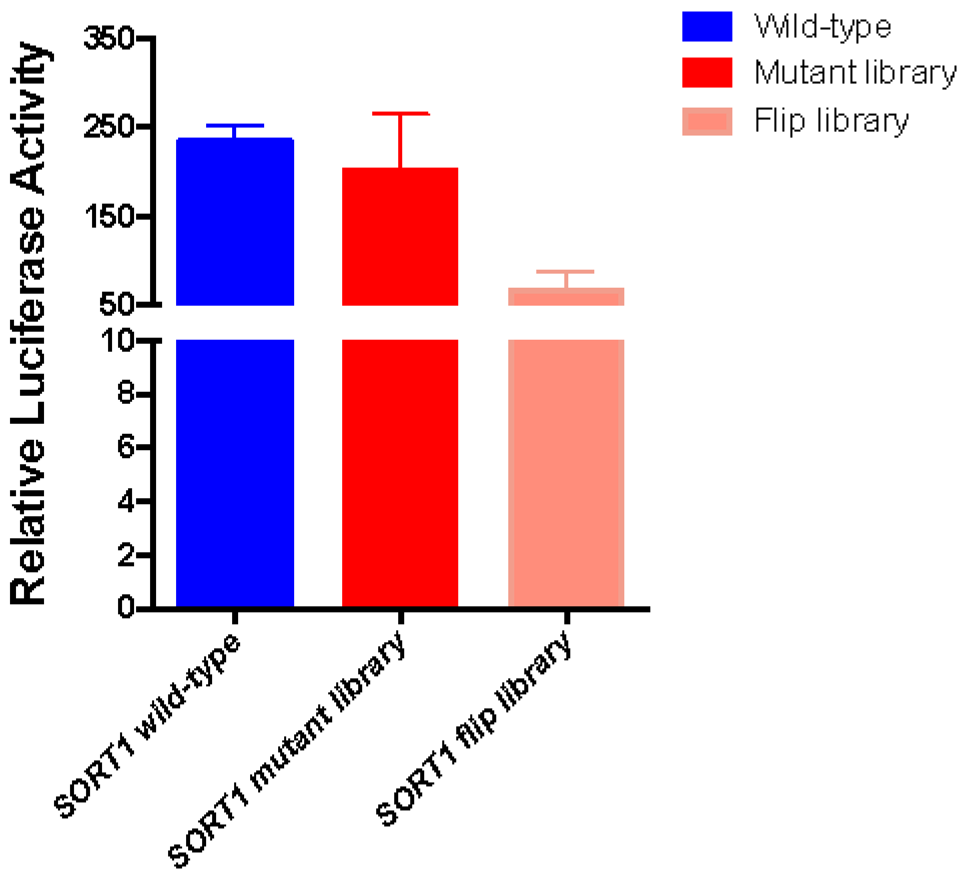

***SORT1* enhancer assays.** Bar charts representing the relative luciferase activity of the *SORT1* enhancer wild-type sequence (blue), mutant library (red) and flip library (light red). Luciferase levels were measured following 24 hour and results are plotted with mean ± standard deviation of at least 3 independent experiments.

*Supplementary Figure 3*

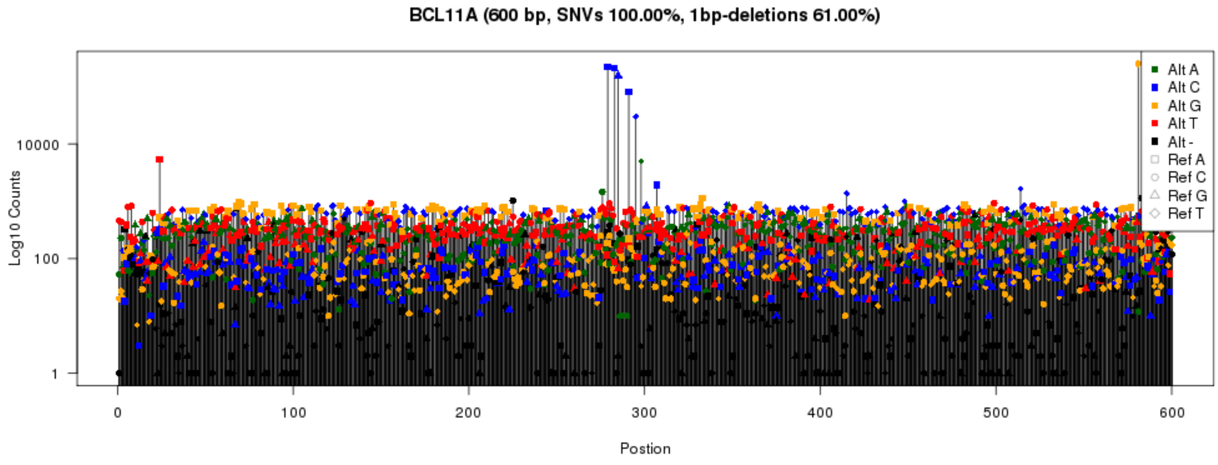

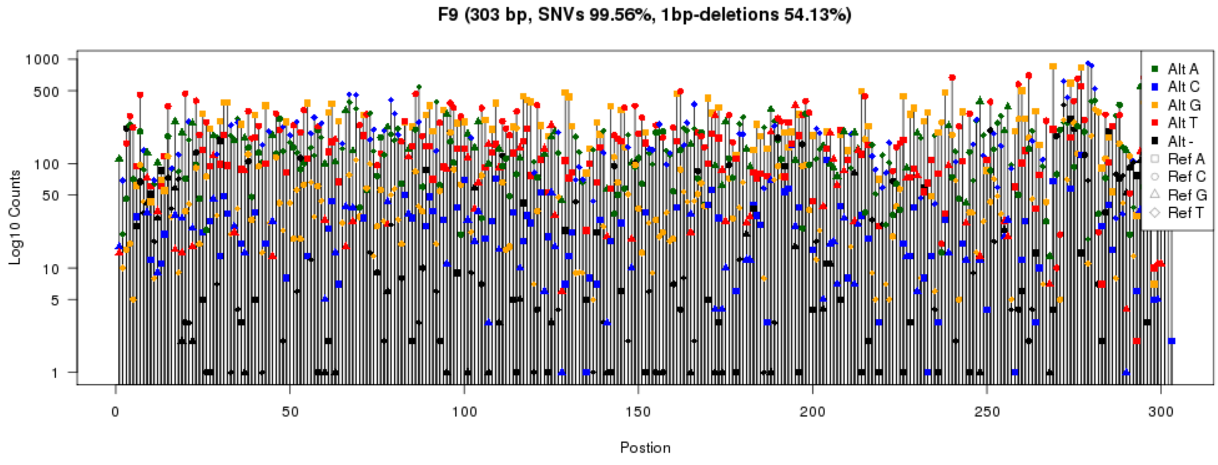

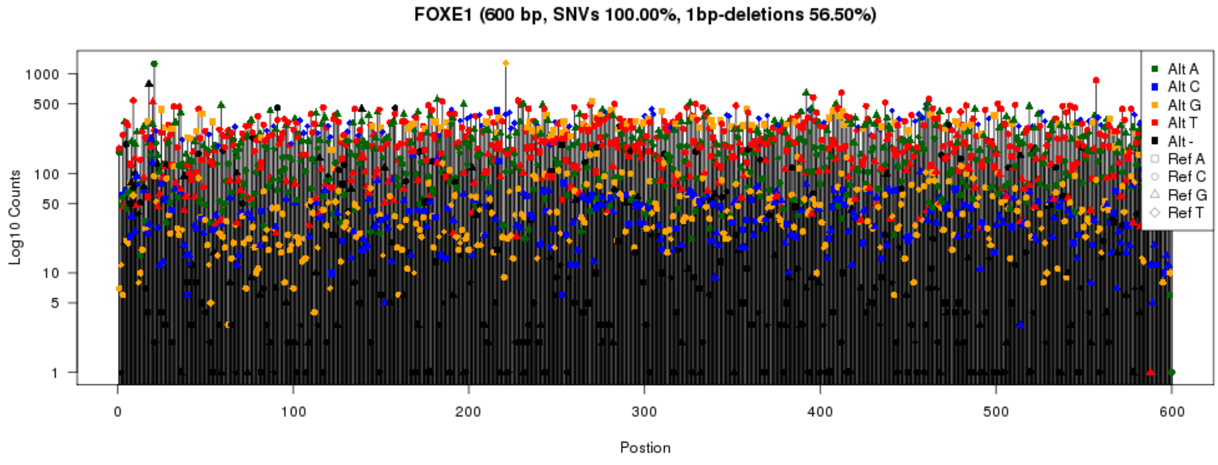

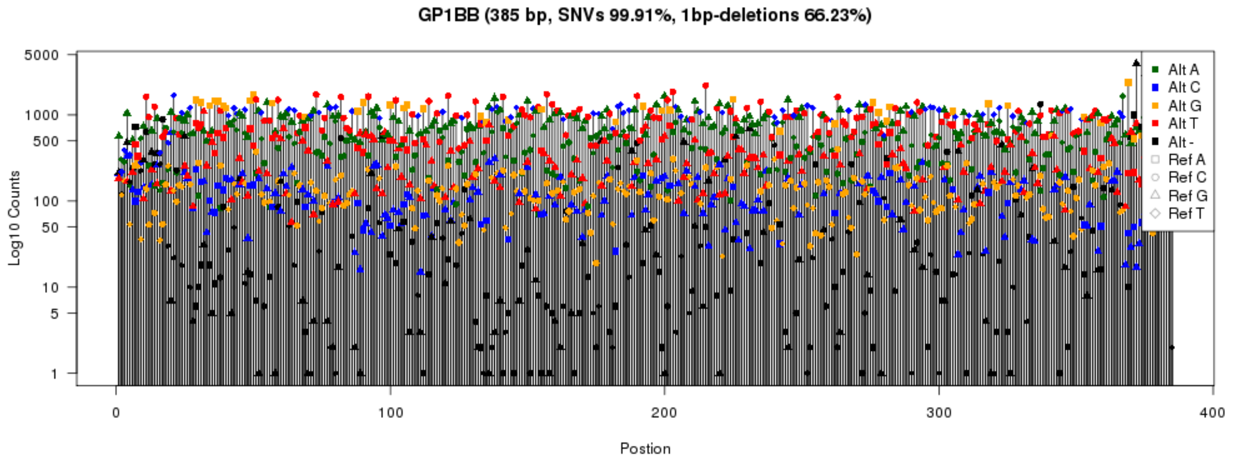
GP1BB (385 bp, SNVs 99.91%, 1bp-deletions 66.23%)
Log10 Counts
Postion
Alt A
Alt C
Alt G
Alt T
Alt -
Ref A
Ref C
Ref G
Ref T

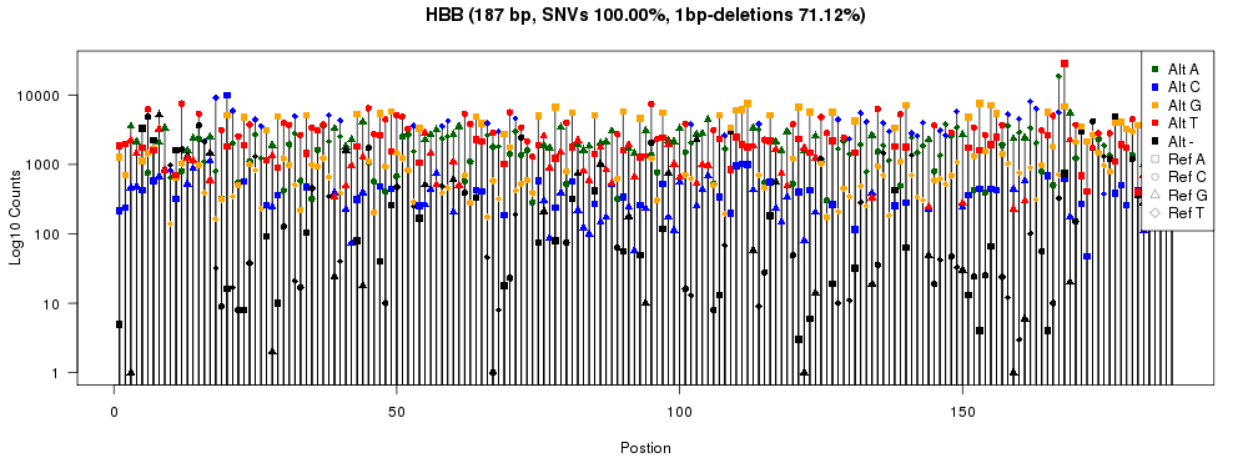
HBB (187 bp, SNVs 100.00%, 1bp-deletions 71.12%)
Log10 Counts
Postion
Alt A
Alt C
Alt G
Alt T
Alt -
Ref A
Ref C
Ref G
Ref T

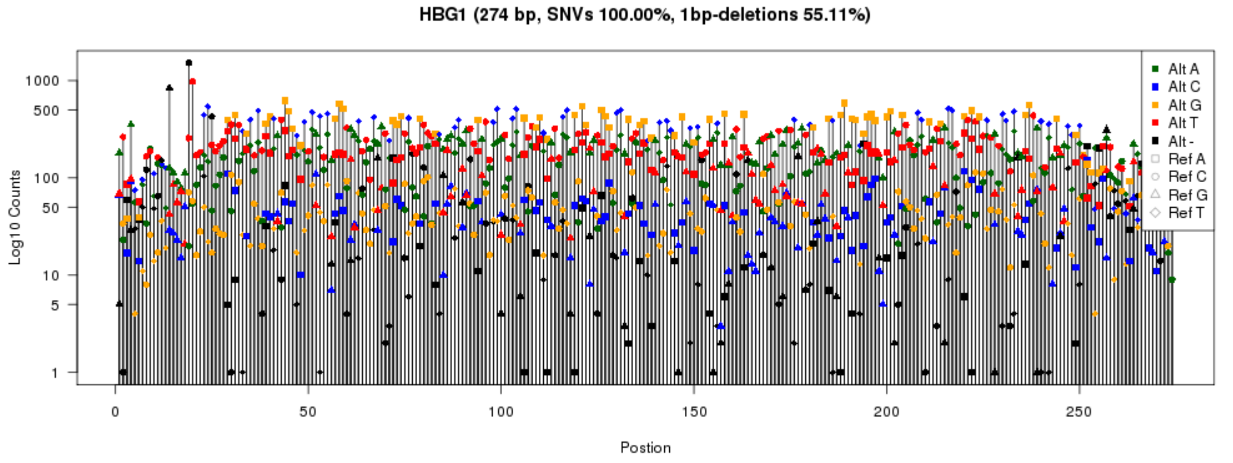
HBG1 (274 bp, SNVs 100.00%, 1bp-deletions 55.11%)
Log10 Counts
Postion
Alt A
Alt C
Alt G
Alt T
Alt -
Ref A
Ref C
Ref G
Ref T

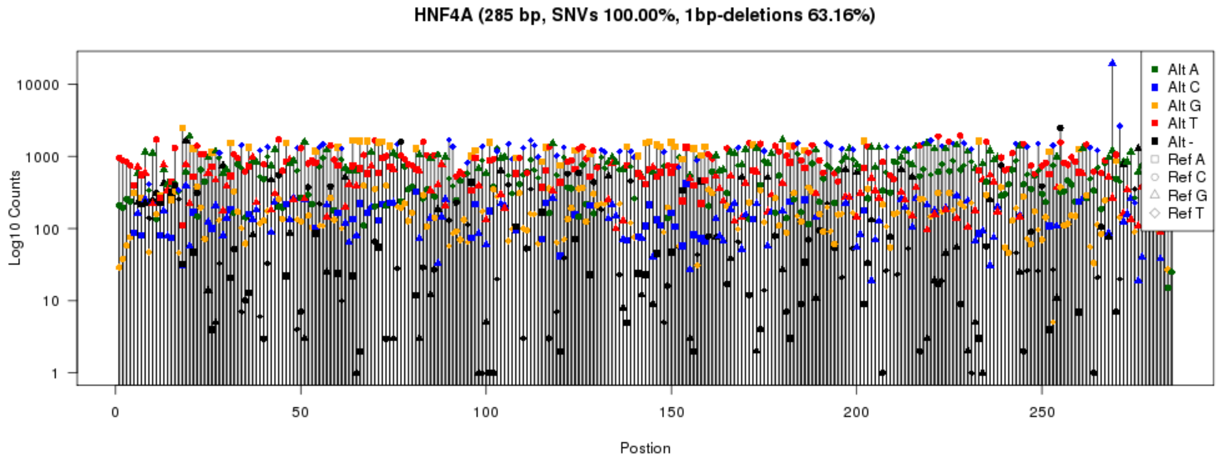
HNF4A (285 bp, SNVs 100.00%, 1bp-deletions 63.16%)
Log10 Counts
Postion
Alt A
Alt C
Alt G
Alt T
Alt -
Ref A
Ref C
Ref G
Ref T

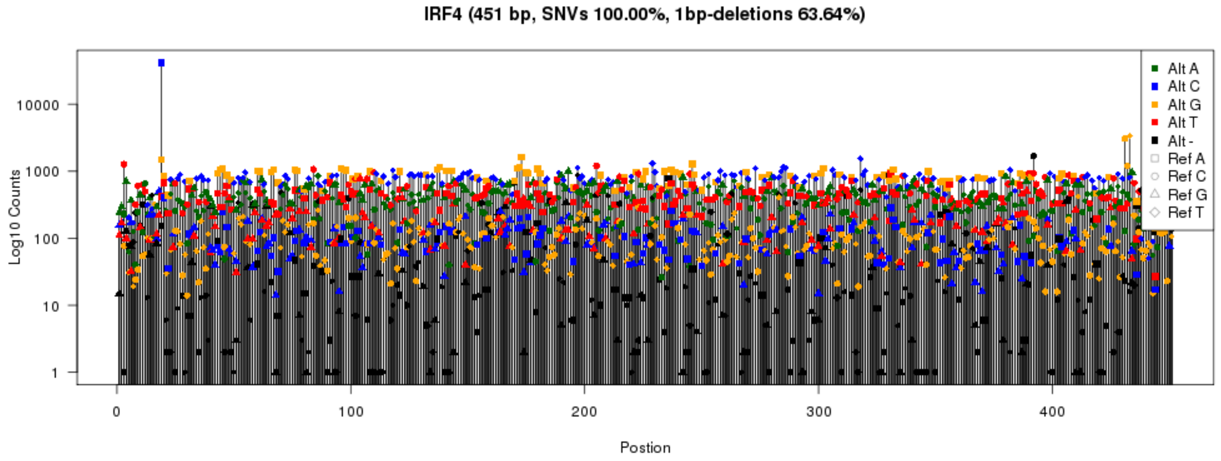
IRF4 (451 bp, SNVs 100.00%, 1bp-deletions 63.64%)
Log10 Counts
Postion
Alt A
Alt C
Alt G
Alt T
Alt -
Ref A
Ref C
Ref G
Ref T

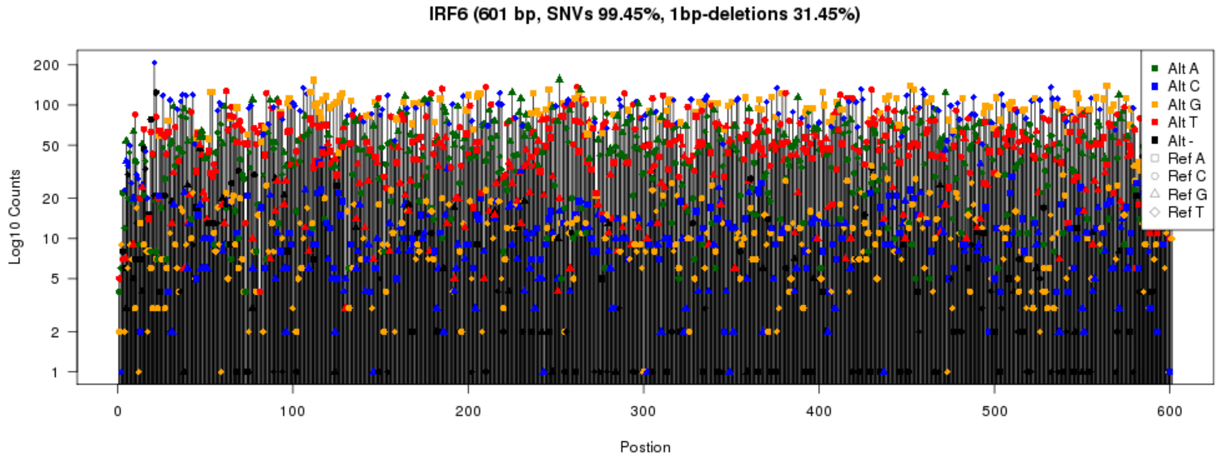
IRF6 (601 bp, SNVs 99.45%, 1bp-deletions 31.45%)
Log10 Counts
Postion
Alt A
Alt C
Alt G
Alt T
Alt -
Ref A
Ref C
Ref G
Ref T

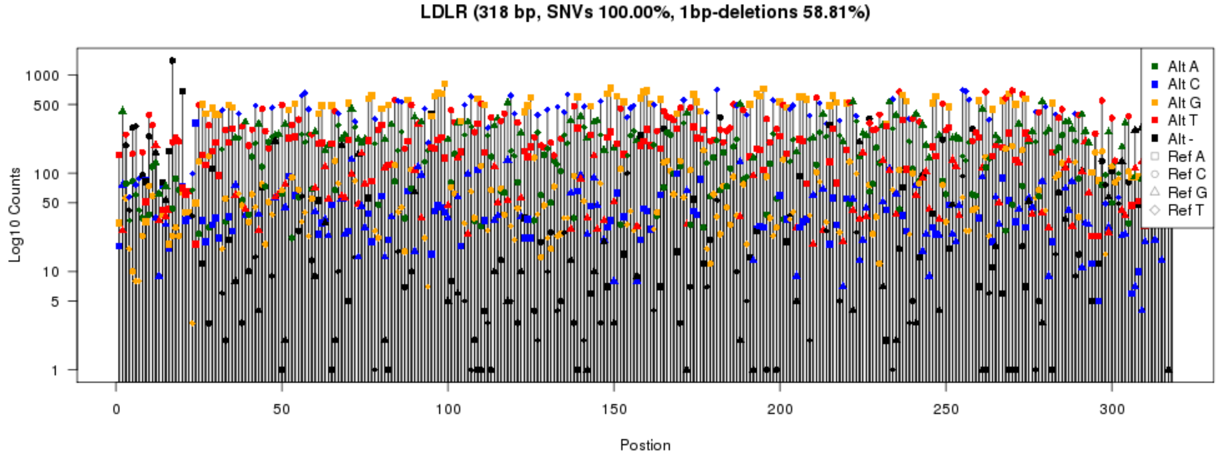
LDLR (318 bp, SNVs 100.00%, 1bp-deletions 58.81%)
Log10 Counts
1000
500
100
50
10
5
1
0
50
100
150
200
250
300
Postion
Alt A
Alt C
Alt G
Alt T
Alt -
Ref A
Ref C
Ref G
Ref T

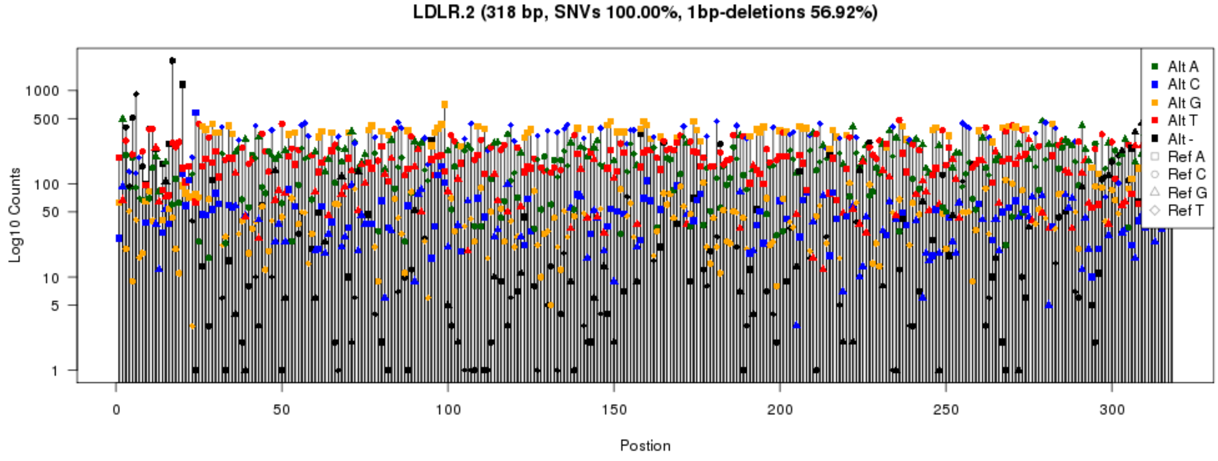
LDLR.2 (318 bp, SNVs 100.00%, 1bp-deletions 56.92%)
Log10 Counts
1000
500
100
50
10
5
1
0
50
100
150
200
250
300
Postion
Alt A
Alt C
Alt G
Alt T
Alt -
Ref A
Ref C
Ref G
Ref T

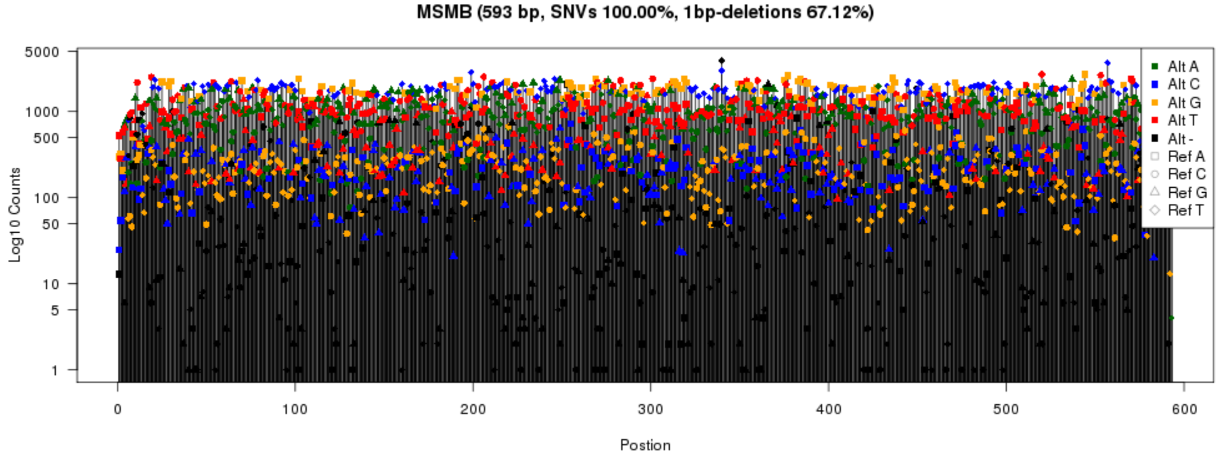
MSMB (593 bp, SNVs 100.00%, 1bp-deletions 67.12%)
Log10 Counts
5000
1000
500
100
50
10
5
1
0
100
200
300
400
500
600
Postion
Alt A
Alt C
Alt G
Alt T
Alt -
Ref A
Ref C
Ref G
Ref T

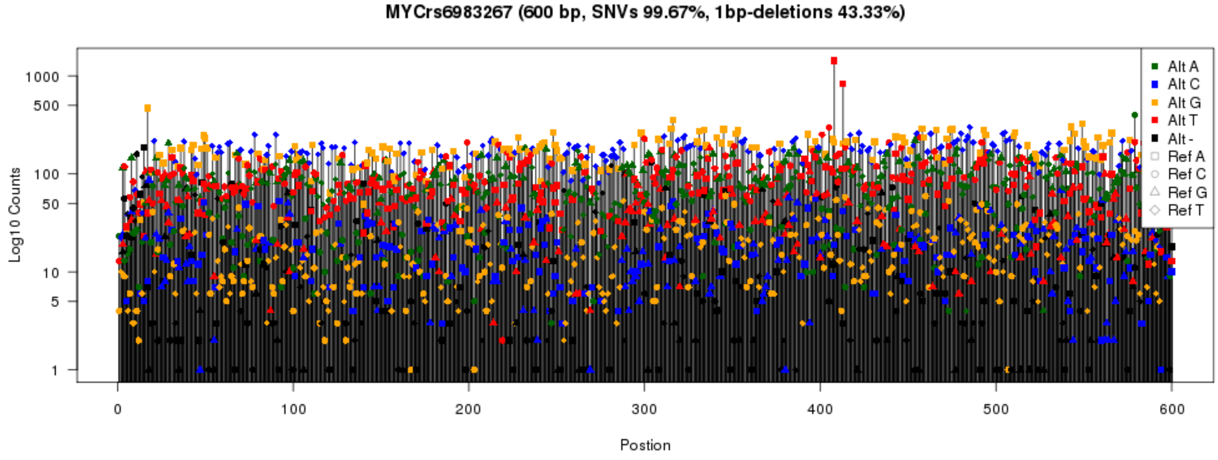
MYCrs6983267 (600 bp, SNVs 99.67%, 1bp-deletions 43.33%)
Log10 Counts
Postion
Alt A
Alt C
Alt G
Alt T
Alt -
Ref A
Ref C
Ref G
Ref T

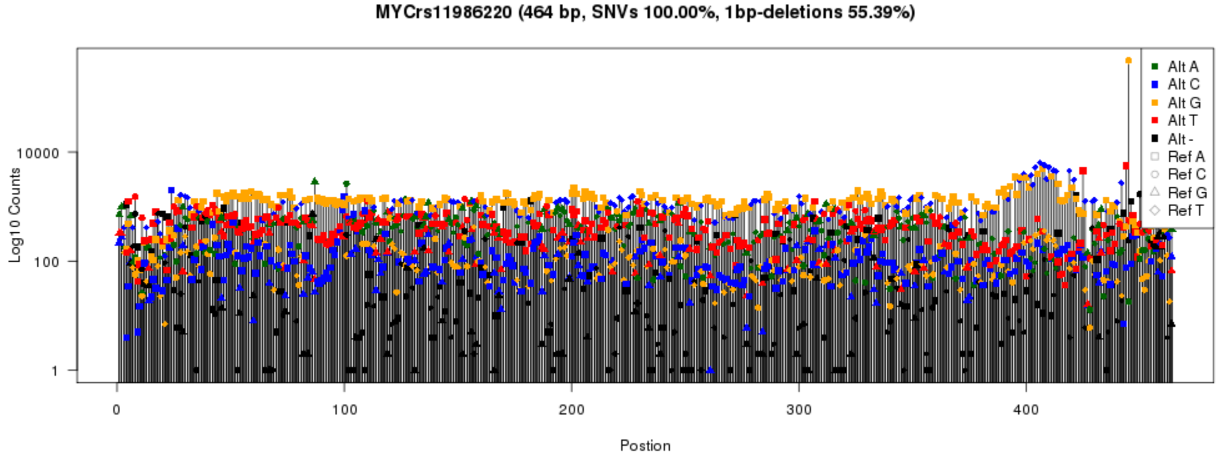
MYCrs11986220 (464 bp, SNVs 100.00%, 1bp-deletions 55.39%)
Log10 Counts
Postion
Alt A
Alt C
Alt G
Alt T
Alt -
Ref A
Ref C
Ref G
Ref T

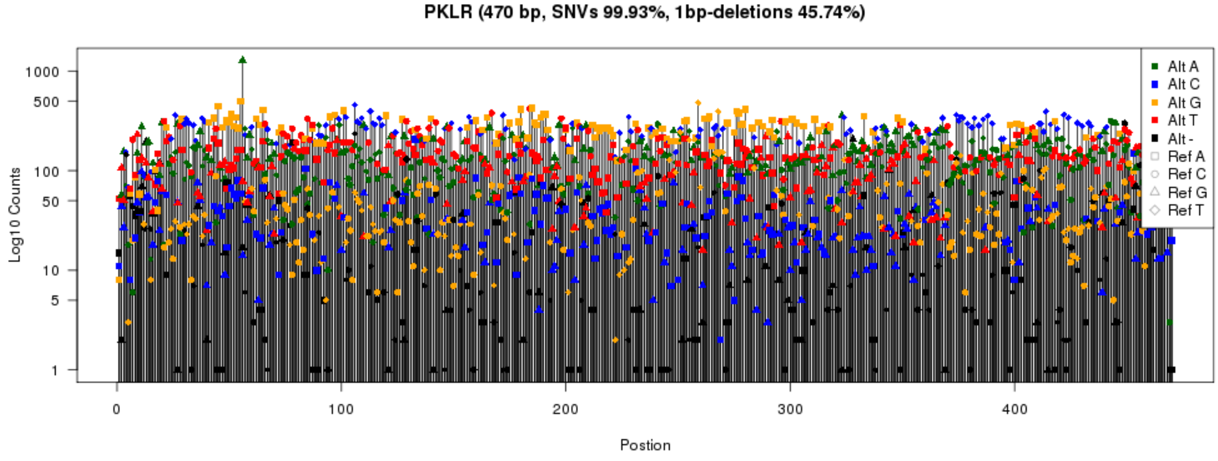
PKLR (470 bp, SNVs 99.93%, 1bp-deletions 45.74%)
Log10 Counts
Postion
Alt A
Alt C
Alt G
Alt T
Alt -
Ref A
Ref C
Ref G
Ref T

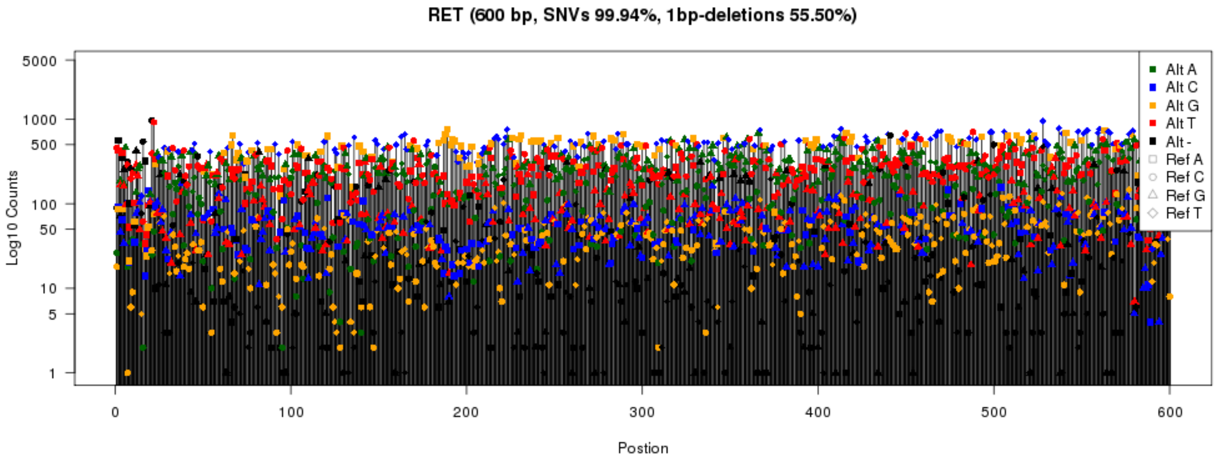
RET (600 bp, SNVs 99.94%, 1bp-deletions 55.50%)
Log10 Counts
Postion
Alt A
Alt C
Alt G
Alt T
Alt -
Ref A
Ref C
Ref G
Ref T

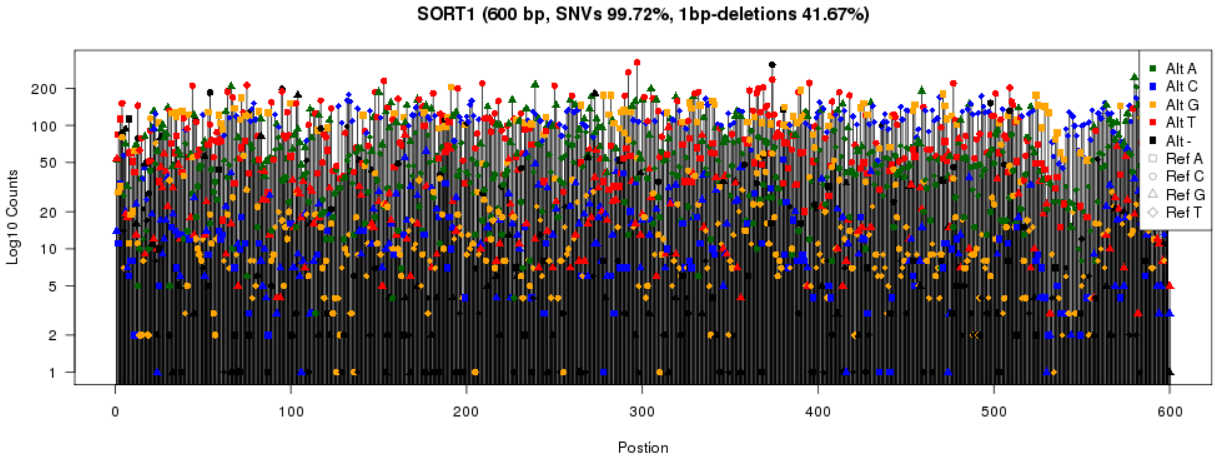
SORT1 (600 bp, SNVs 99.72%, 1bp-deletions 41.67%)
Log10 Counts
Postion
Alt A
Alt C
Alt G
Alt T
Alt -
Ref A
Ref C
Ref G
Ref T

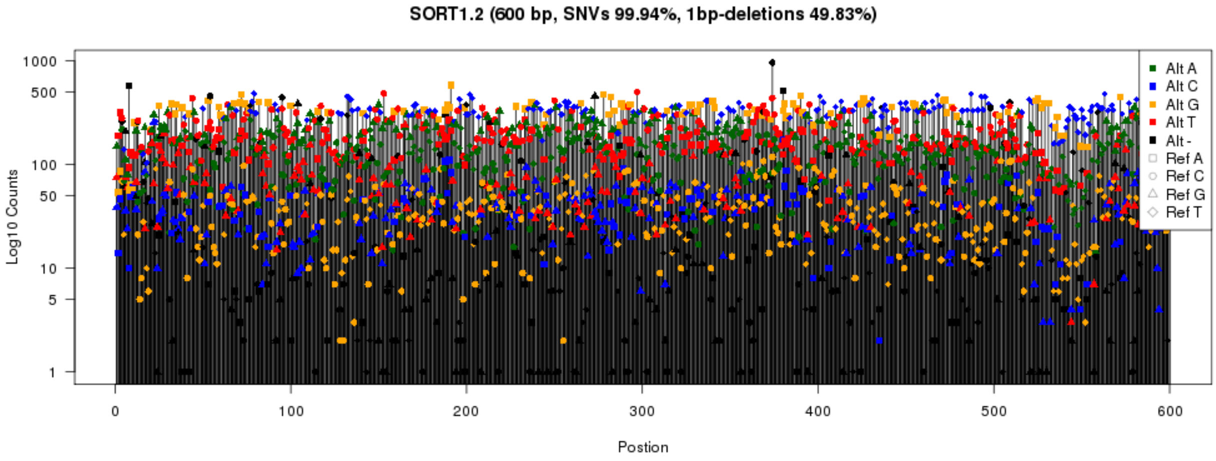
SORT1.2 (600 bp, SNVs 99.94%, 1bp-deletions 49.83%)
Log10 Counts
Postion
Alt A
Alt C
Alt G
Alt T
Alt -
Ref A
Ref C
Ref G
Ref T

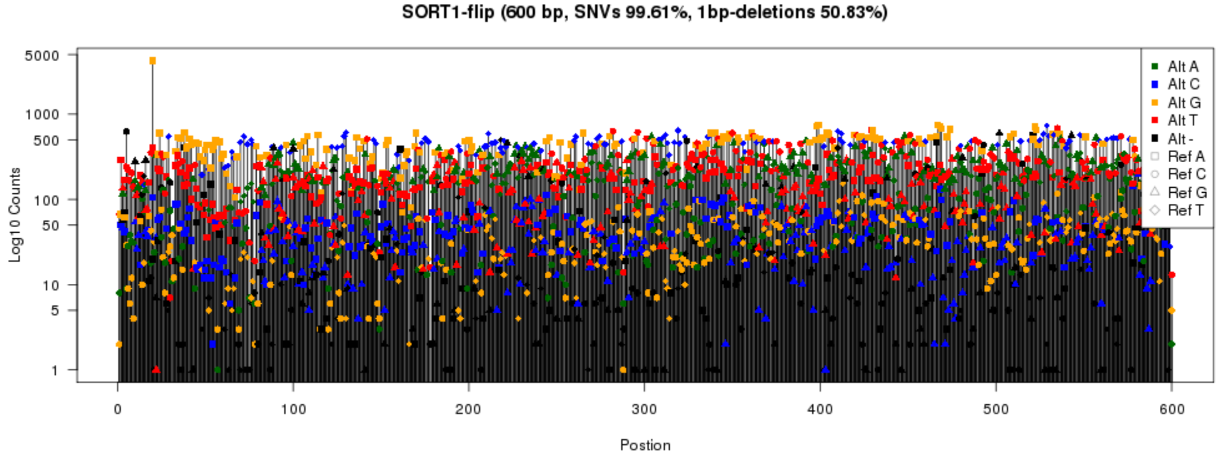
SORT1-flip (600 bp, SNVs 99.61%, 1bp-deletions 50.83%)
Log10 Counts
Postion
Alt A
Alt C
Alt G
Alt T
Alt -
Ref A
Ref C
Ref G
Ref T

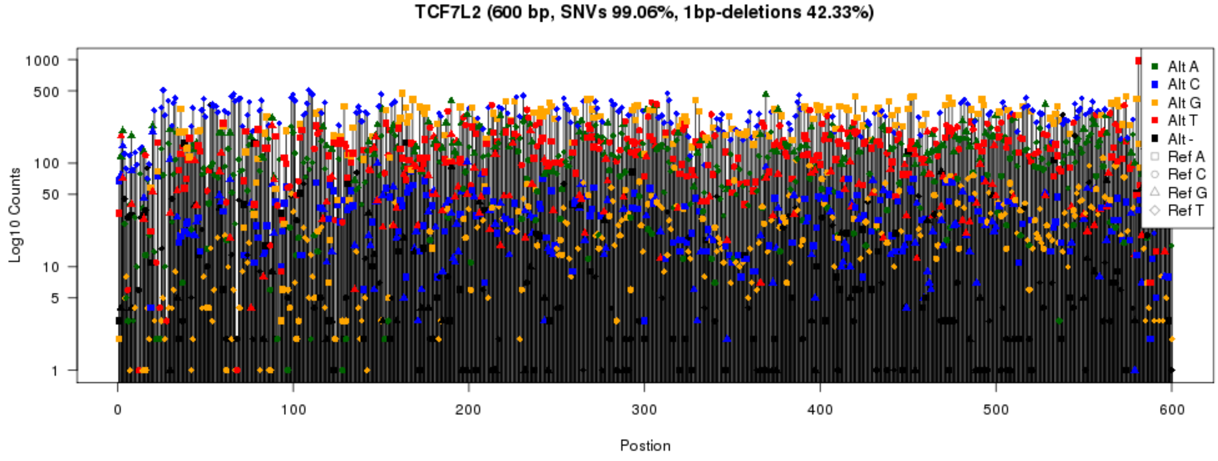
TCF7L2 (600 bp, SNVs 99.06%, 1bp-deletions 42.33%)
Log10 Counts
Postion
Alt A
Alt C
Alt G
Alt T
Alt -
Ref A
Ref C
Ref G
Ref T

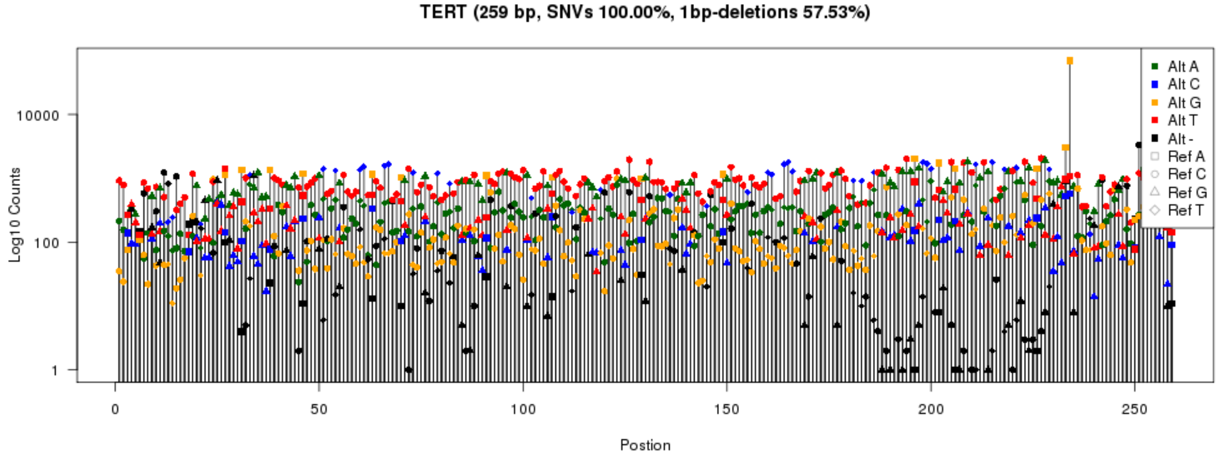
TERT (259 bp, SNVs 100.00%, 1bp-deletions 57.53%)
Log10 Counts
Postion
Alt A
Alt C
Alt G
Alt T
Alt -
Ref A
Ref C
Ref G
Ref T

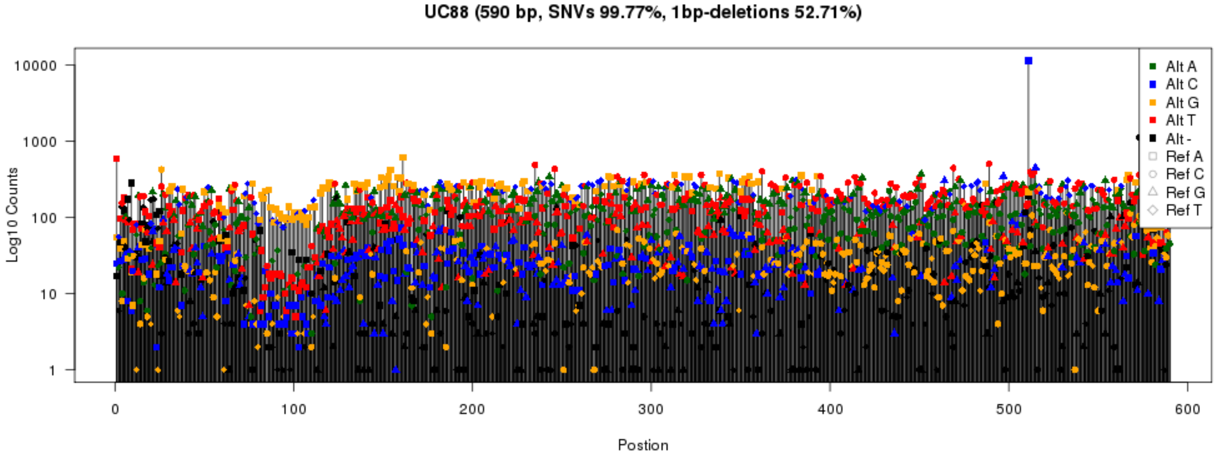

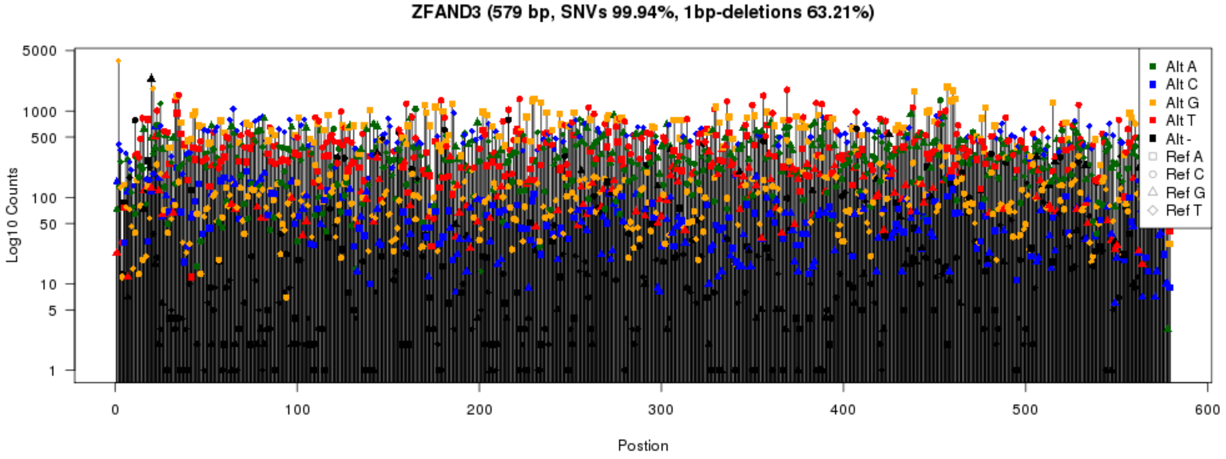

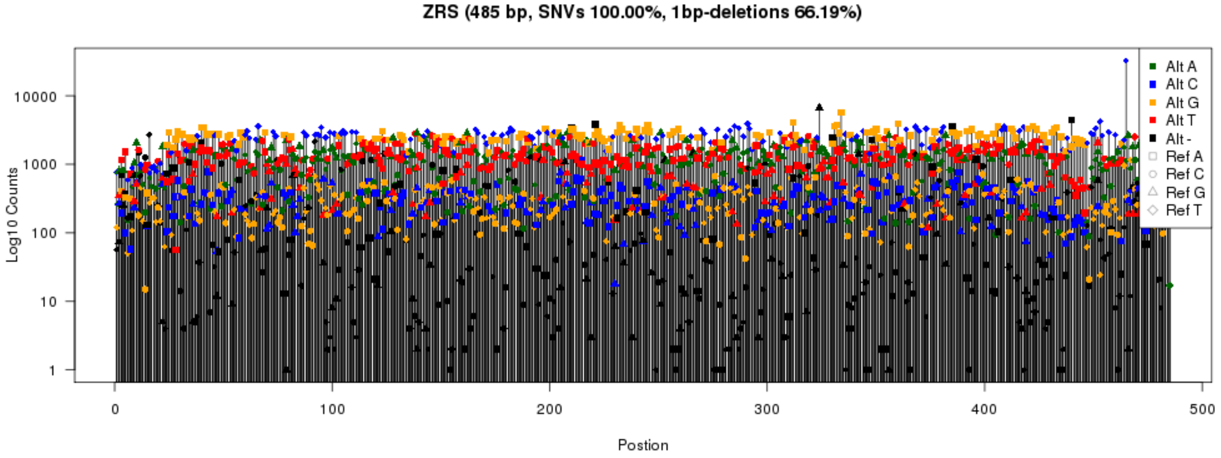

***Tags assigned from saturation mutagenesis libraries.*** Plots for all 24 assignments described in Supplementary Table 3. Variant are sampled evenly across all the regions. The number of tags per variant follows previously described substitution patterns for mutagenesis using Taq polymerases.

#### *Supplementary Figure 4*

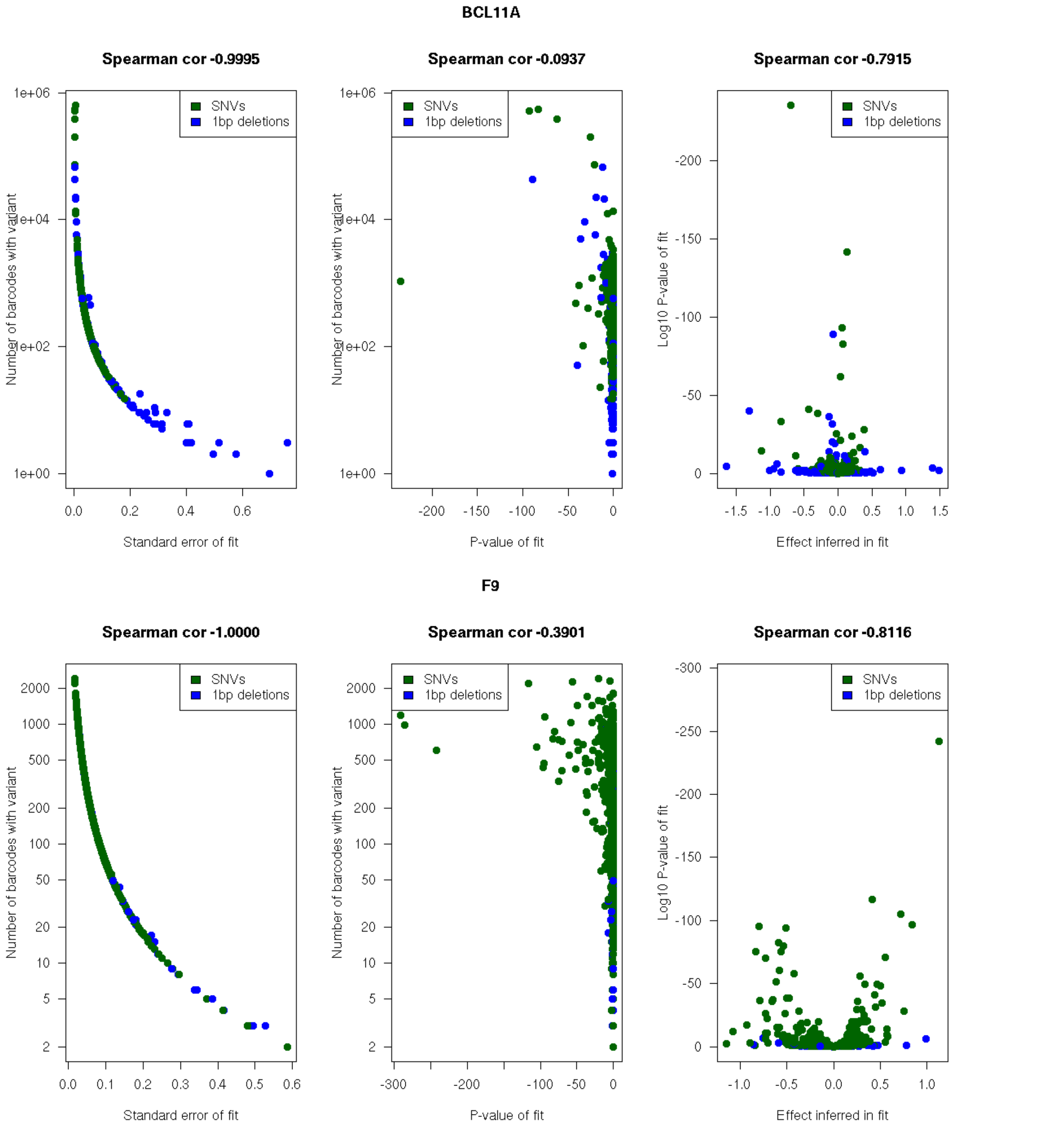

#### FOXE1

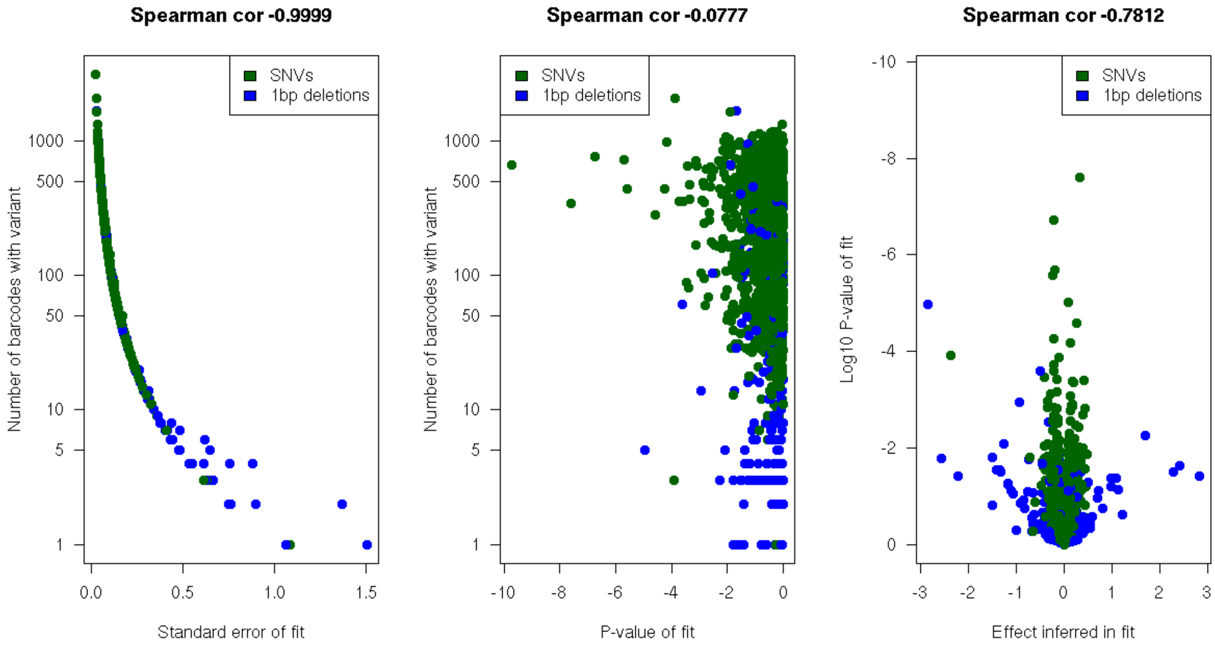
Spearman cor -0.9999
SNVs
1bp deletions
Number of barcodes with variant
1000
500
100
50
10
5
1
0.0
0.5
1.0
1.5
Standard error of fit
Spearman cor -0.0777
SNVs
1bp deletions
Number of barcodes with variant
1000
500
100
50
10
5
1
-10
-8
-6
-4
-2
0
P-value of fit
Spearman cor -0.7812
SNVs
1bp deletions
Log10 P-value of fit
-10
-8
-6
-4
-2
0
-3
-2
-1
0
1
2
3
Effect inferred in fit

## GP1BB

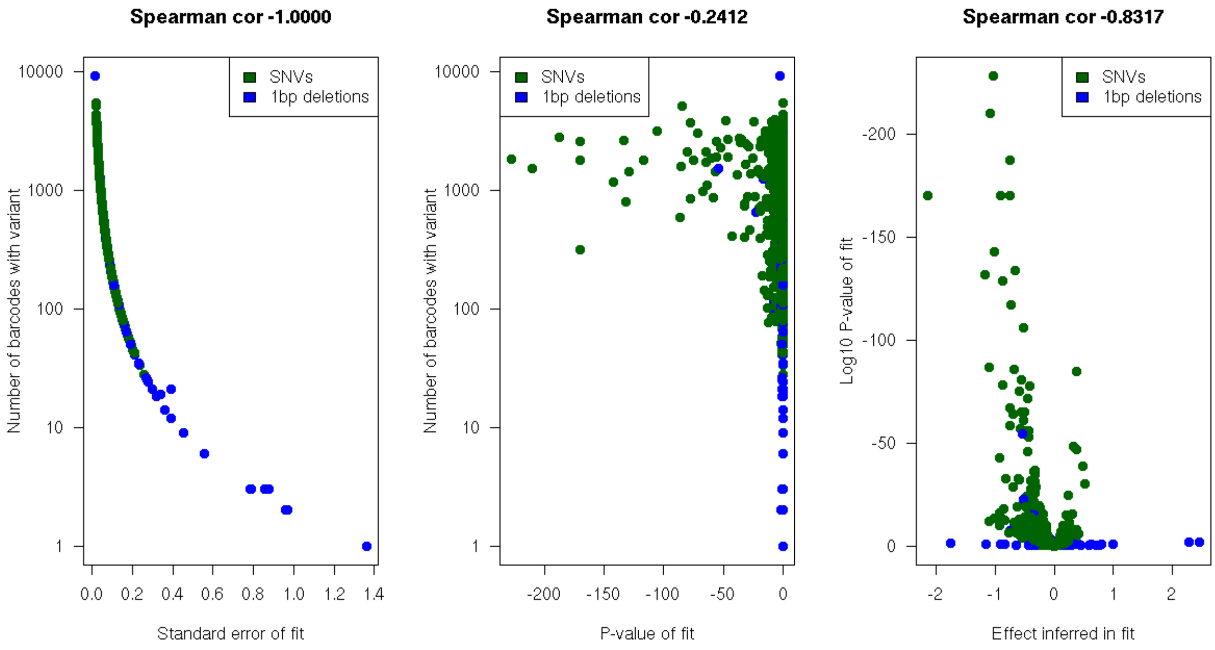
Spearman cor -1.0000
SNVs
1bp deletions
Number of barcodes with variant
10000
1000
100
10
1
0.0
0.2
0.4
0.6
0.8
1.0
1.2
1.4
Standard error of fit
Spearman cor -0.2412
SNVs
1bp deletions
Number of barcodes with variant
10000
1000
100
10
1
-200
-150
-100
-50
0
P-value of fit
Spearman cor -0.8317
SNVs
1bp deletions
Log10 P-value of fit
-200
-150
-100
-50
0
-2
-1
0
1
2
Effect inferred in fit

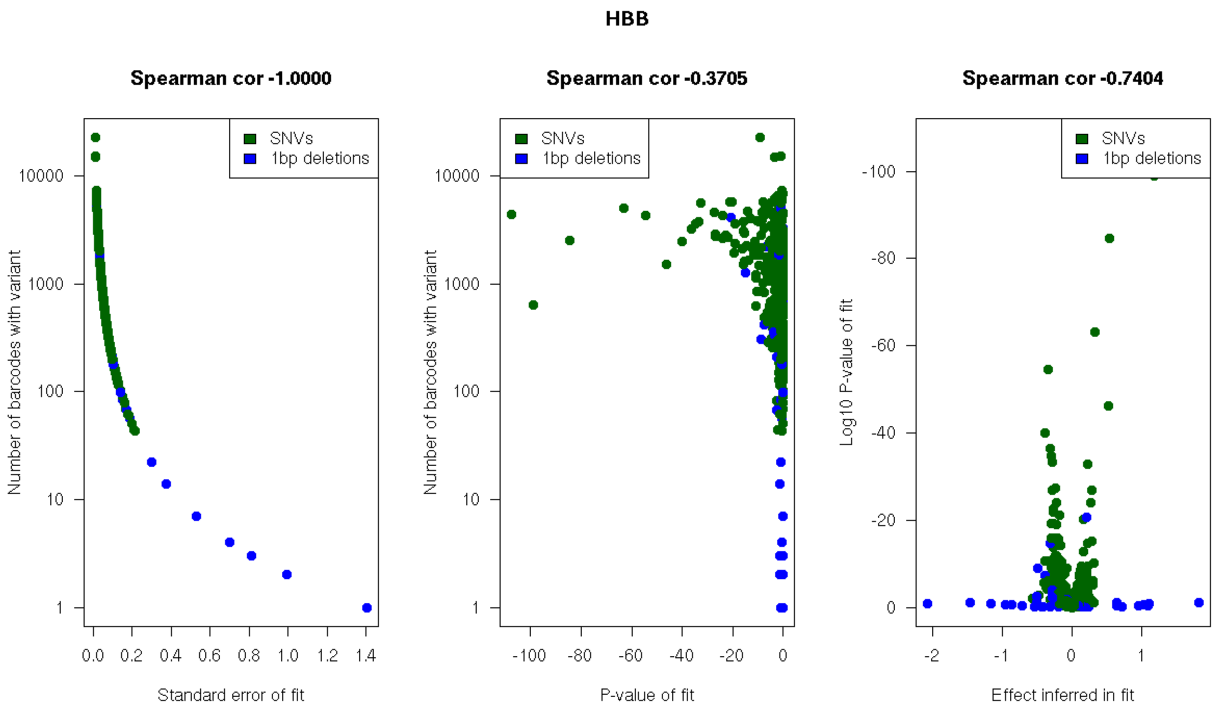
HBB
Spearman cor -1.0000
Spearman cor -0.3705
Spearman cor -0.7404
SNVs
1bp deletions
Number of barcodes with variant
Log10 P-value of fit
Standard error of fit
P-value of fit
Effect inferred in fit

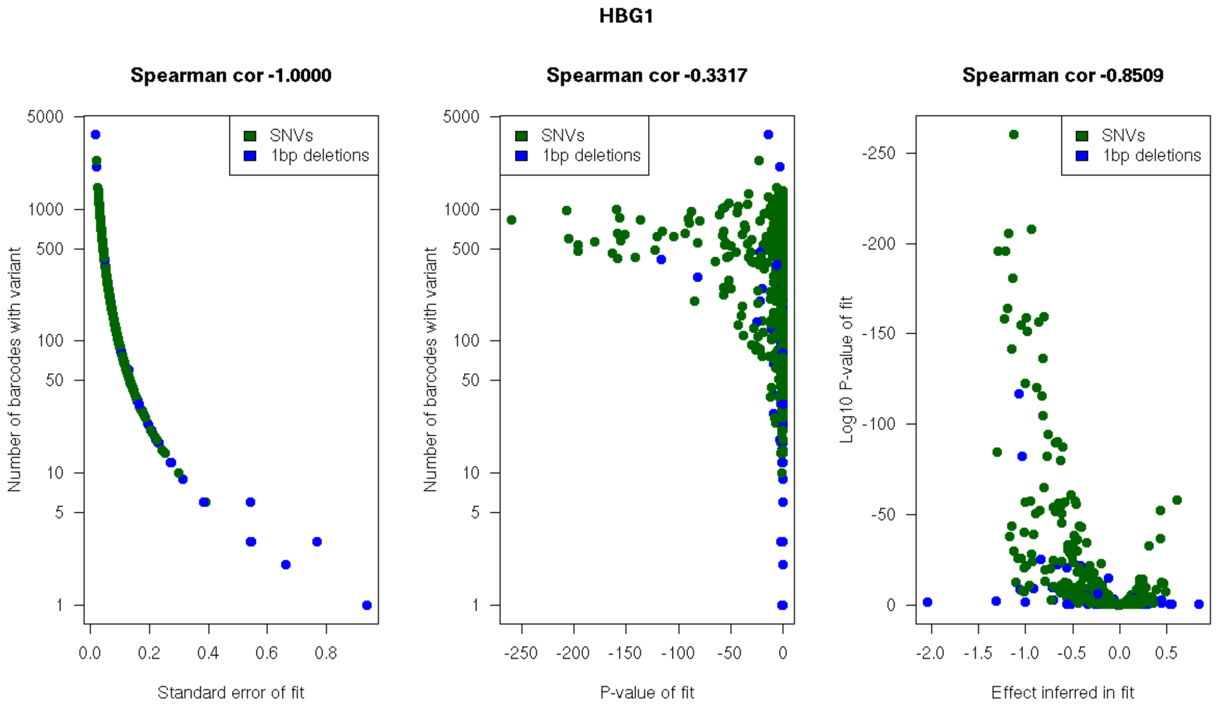
HBG1
Spearman cor -1.0000
Spearman cor -0.3317
Spearman cor -0.8509
SNVs
1bp deletions
Number of barcodes with variant
Log10 P-value of fit
Standard error of fit
P-value of fit
Effect inferred in fit

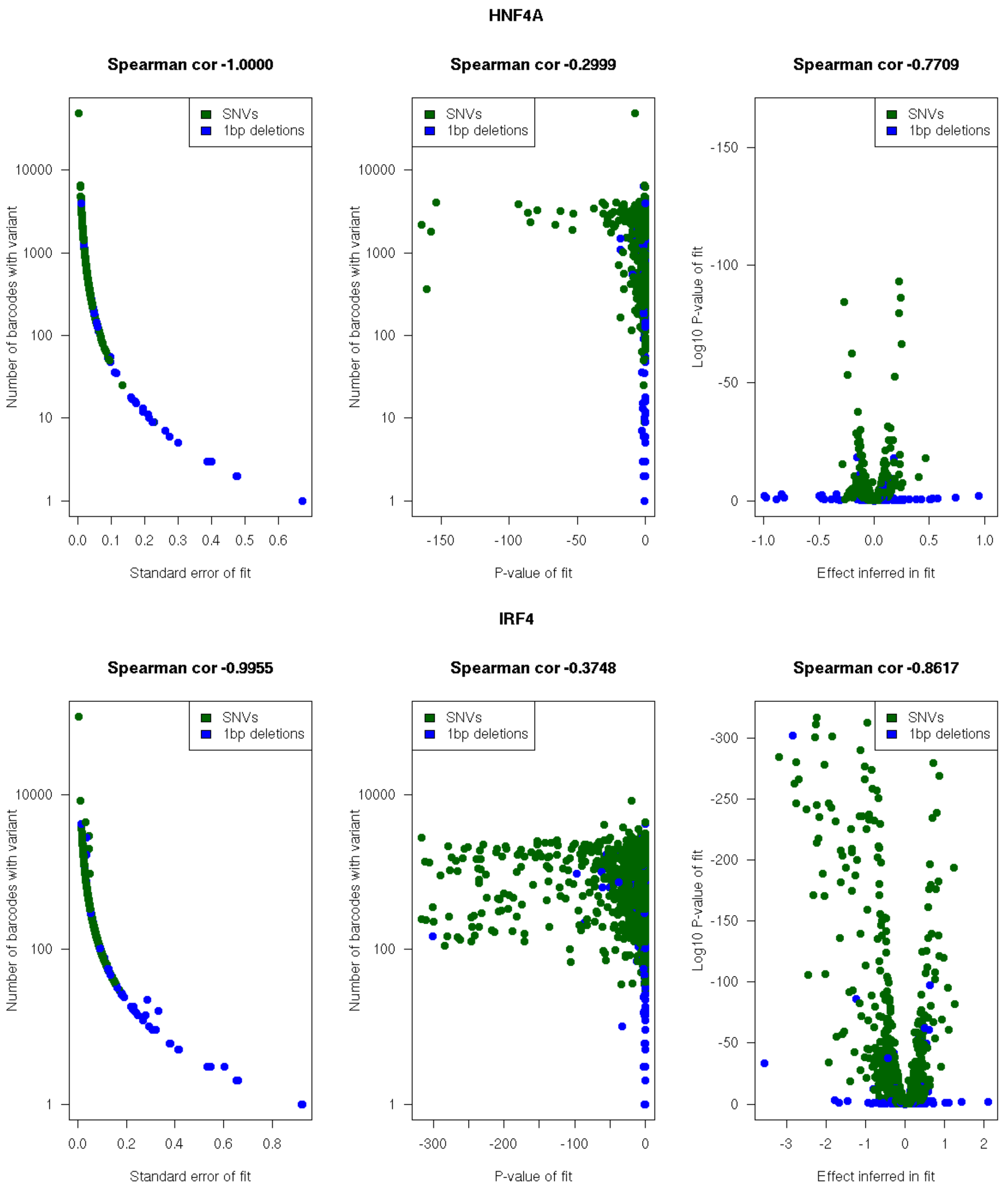
HNF4A
Spearman cor -1.0000
Spearman cor -0.2999
Spearman cor -0.7709
SNVs
1bp deletions
Number of barcodes with variant
Standard error of fit
P-value of fit
Log10 P-value of fit
Effect inferred in fit
IRF4
Spearman cor -0.9955
Spearman cor -0.3748
Spearman cor -0.8617

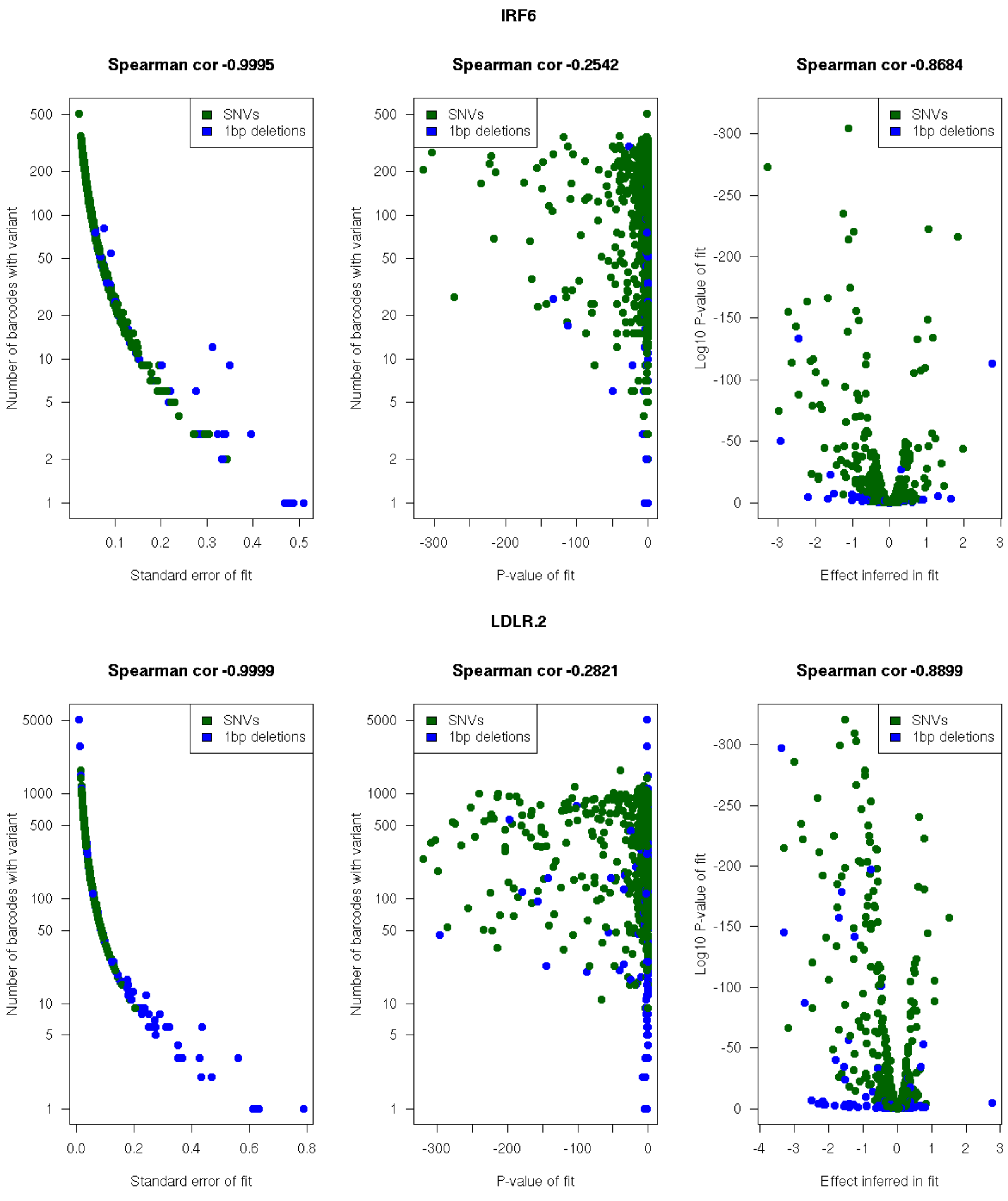
IRF6
Spearman cor -0.9995
Spearman cor -0.2542
Spearman cor -0.8684
SNVs
1bp deletions
Number of barcodes with variant
Log10 P-value of fit
Standard error of fit
P-value of fit
Effect inferred in fit
LDLR.2
Spearman cor -0.9999
Spearman cor -0.2821
Spearman cor -0.8899

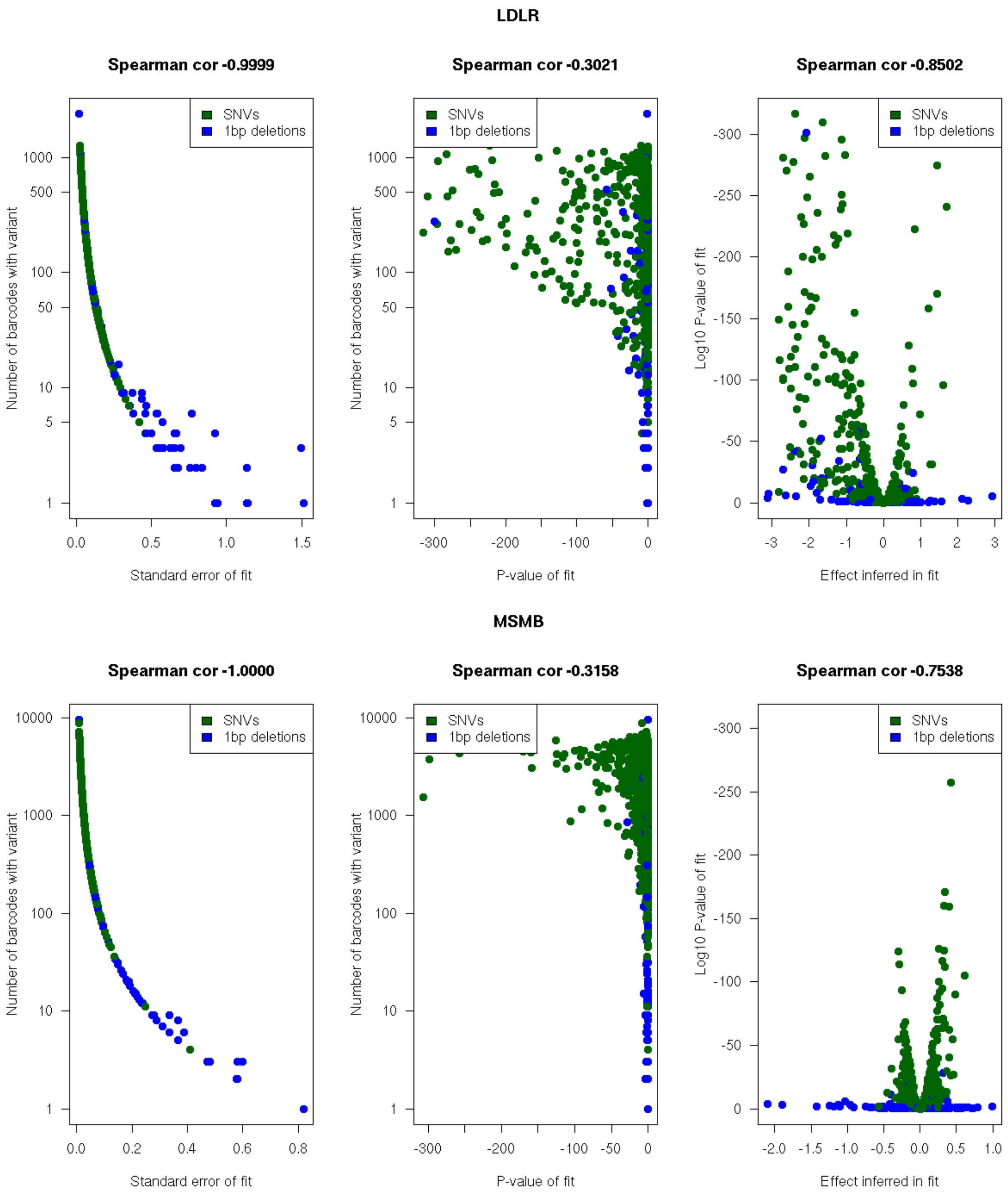
LDLR
Spearman cor -0.9999
Spearman cor -0.3021
Spearman cor -0.8502
SNVs
1bp deletions
Number of barcodes with variant
Log10 P-value of fit
Standard error of fit
P-value of fit
Effect inferred in fit
MSMB
Spearman cor -1.0000
Spearman cor -0.3158
Spearman cor -0.7538
SNVs
1bp deletions
Number of barcodes with variant
Log10 P-value of fit
Standard error of fit
P-value of fit
Effect inferred in fit

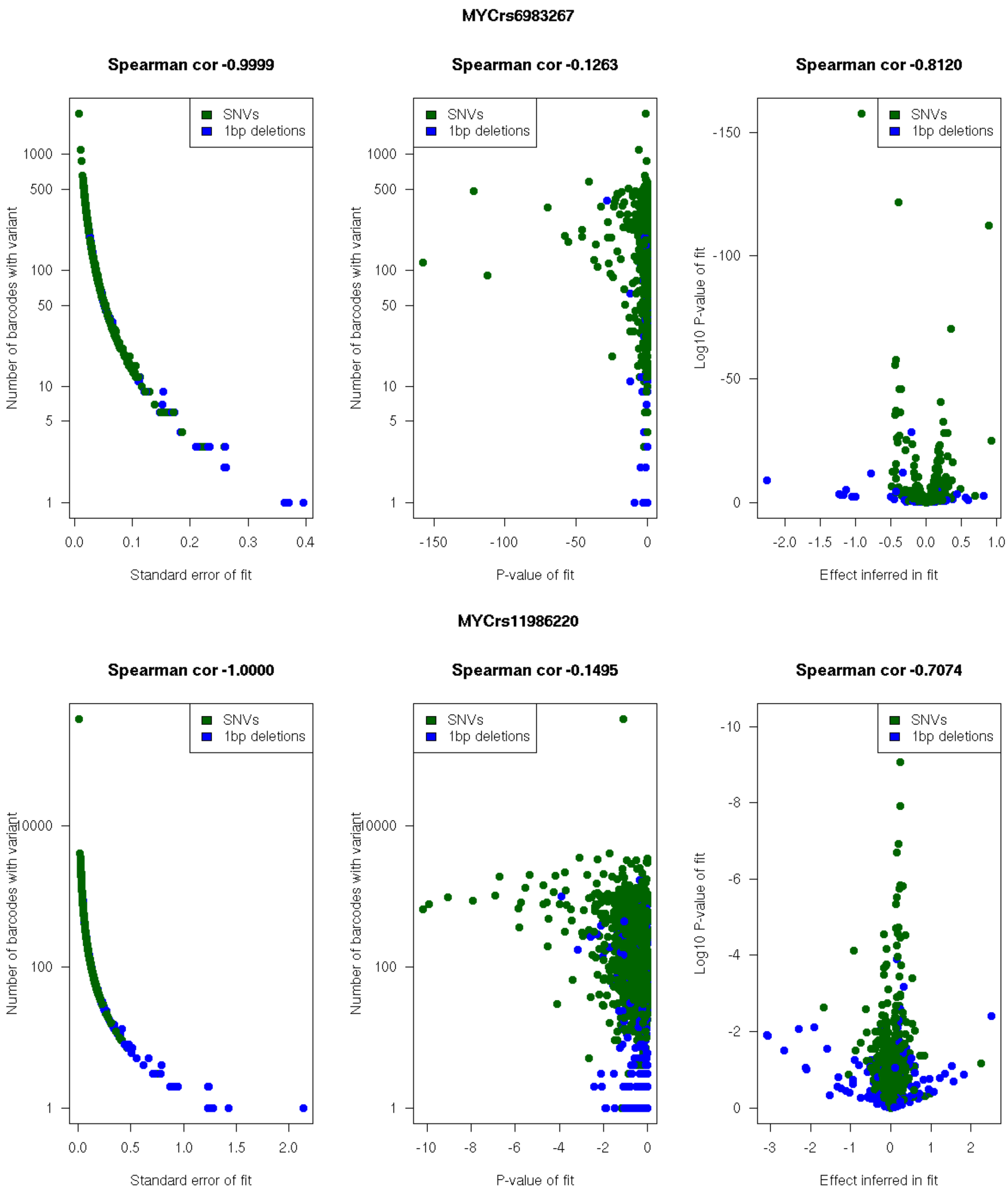
MYCrs6983267
Spearman cor -0.9999
Spearman cor -0.1263
Spearman cor -0.8120
SNVs
1bp deletions
Number of barcodes with variant
Standard error of fit
P-value of fit
Log10 P-value of fit
Effect inferred in fit
MYCrs11986220
Spearman cor -1.0000
Spearman cor -0.1495
Spearman cor -0.7074

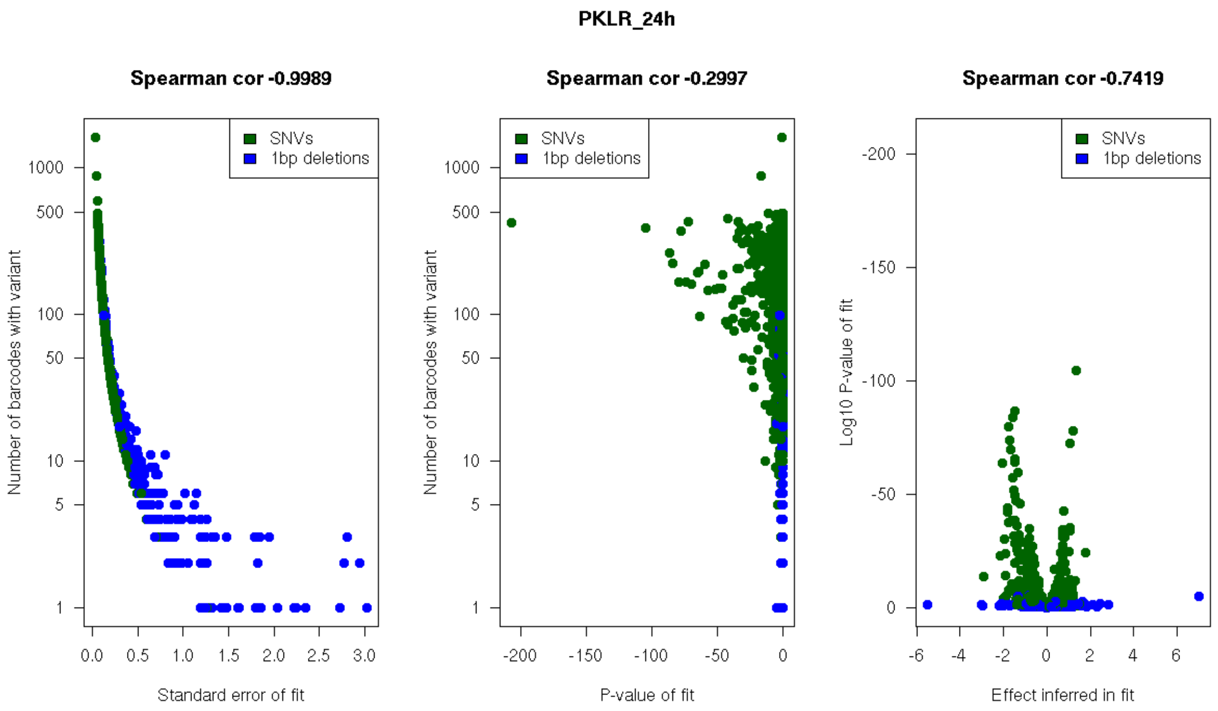
PKLR_24h
Spearman cor -0.9989
Spearman cor -0.2997
Spearman cor -0.7419
SNVs
1bp deletions
Number of barcodes with variant
Log10 P-value of fit
Standard error of fit
P-value of fit
Effect inferred in fit

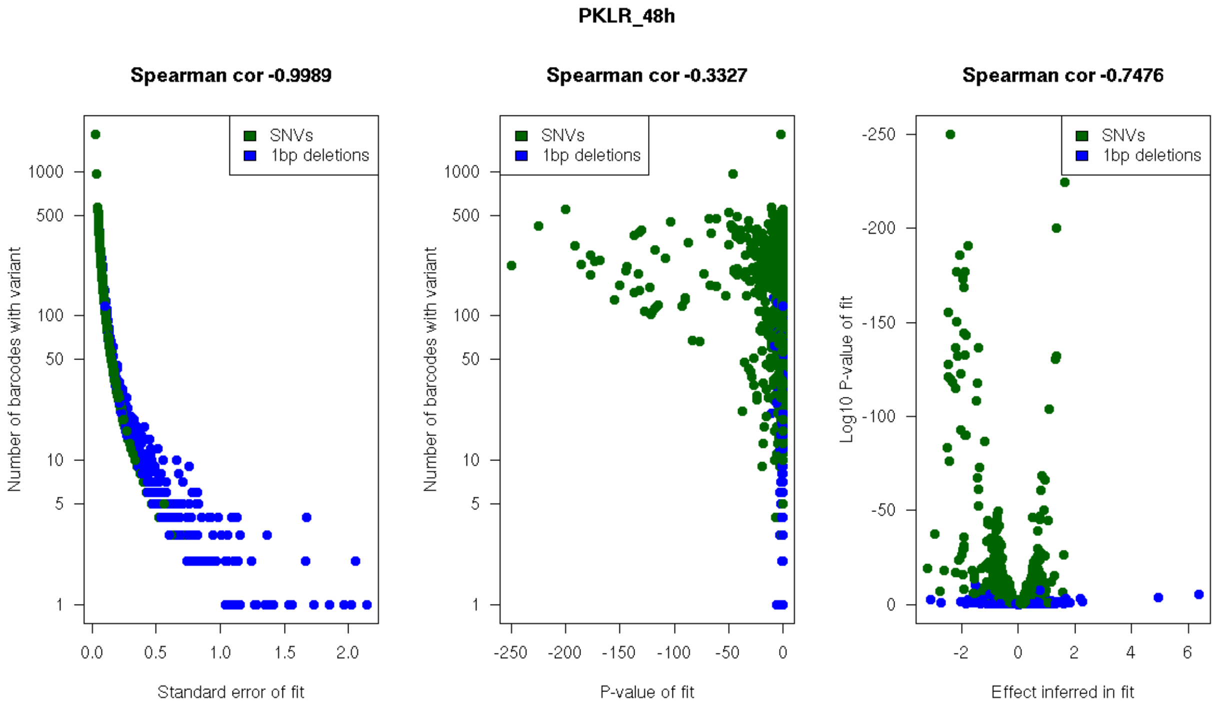
PKLR_48h
Spearman cor -0.9989
Spearman cor -0.3327
Spearman cor -0.7476
SNVs
1bp deletions
Number of barcodes with variant
Log10 P-value of fit
Standard error of fit
P-value of fit
Effect inferred in fit

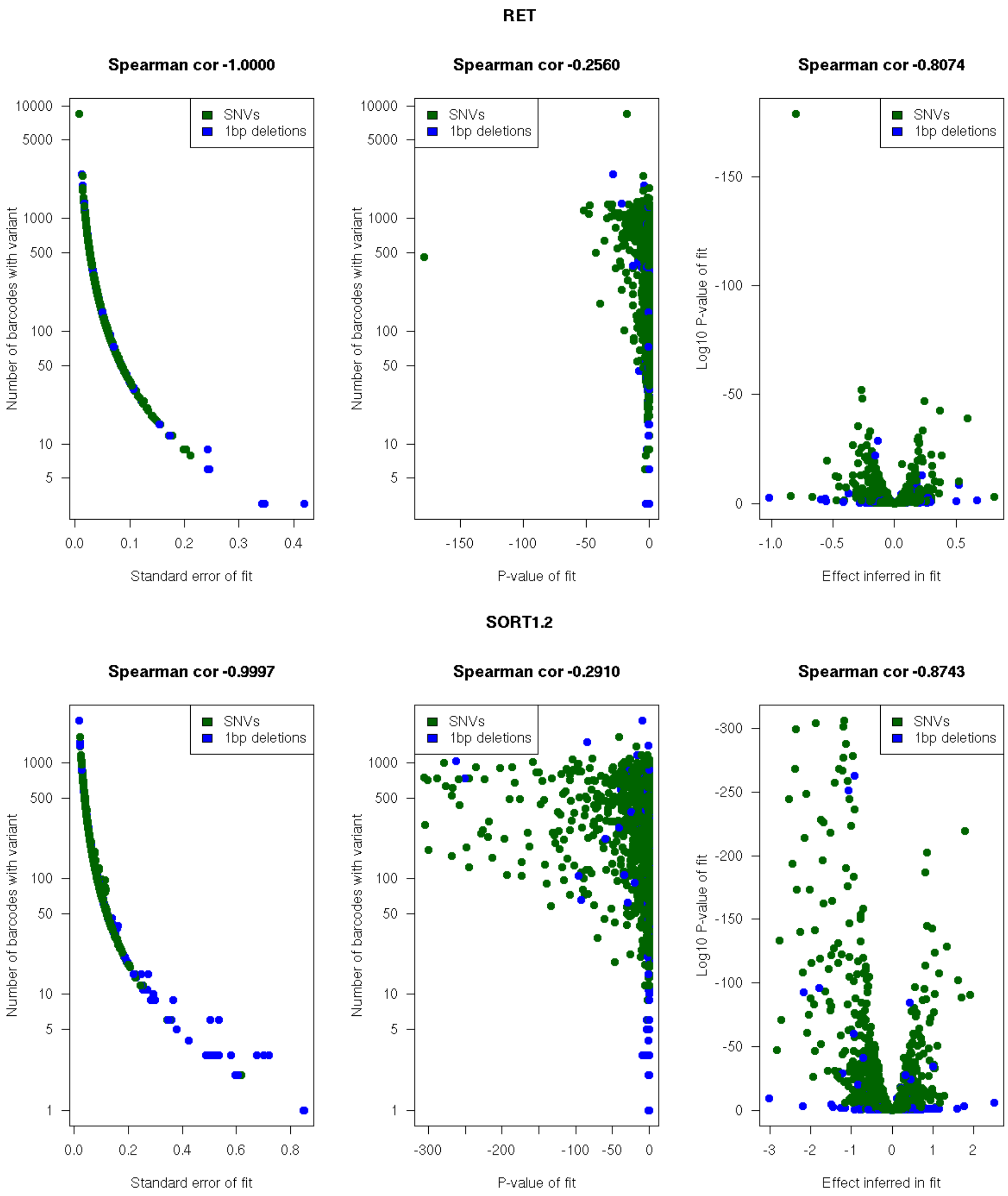
RET
Spearman cor -1.0000
Spearman cor -0.2560
Spearman cor -0.8074
SNVs
1bp deletions
Number of barcodes with variant
Standard error of fit
P-value of fit
Log10 P-value of fit
Effect inferred in fit
SORT1.2
Spearman cor -0.9997
Spearman cor -0.2910
Spearman cor -0.8743
SNVs
1bp deletions
Number of barcodes with variant
Standard error of fit
P-value of fit
Log10 P-value of fit
Effect inferred in fit

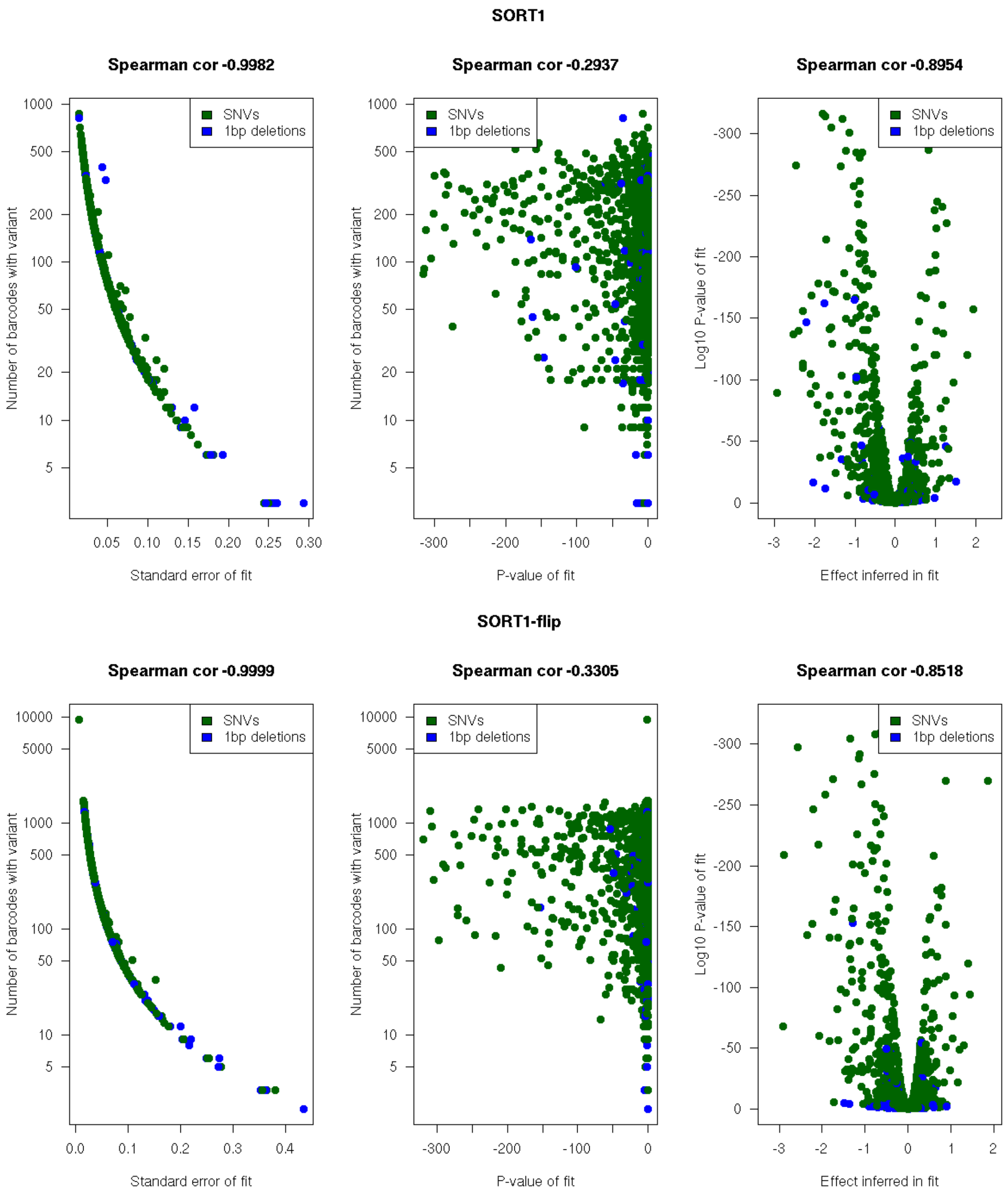
SORT1
Spearman cor -0.9982
Spearman cor -0.2937
Spearman cor -0.8954
SNVs
1bp deletions
Number of barcodes with variant
Log10 P-value of fit
Standard error of fit
P-value of fit
Effect inferred in fit
SORT1-flip
Spearman cor -0.9999
Spearman cor -0.3305
Spearman cor -0.8518
SNVs
1bp deletions
Number of barcodes with variant
Log10 P-value of fit
Standard error of fit
P-value of fit
Effect inferred in fit

**TCF7L2**

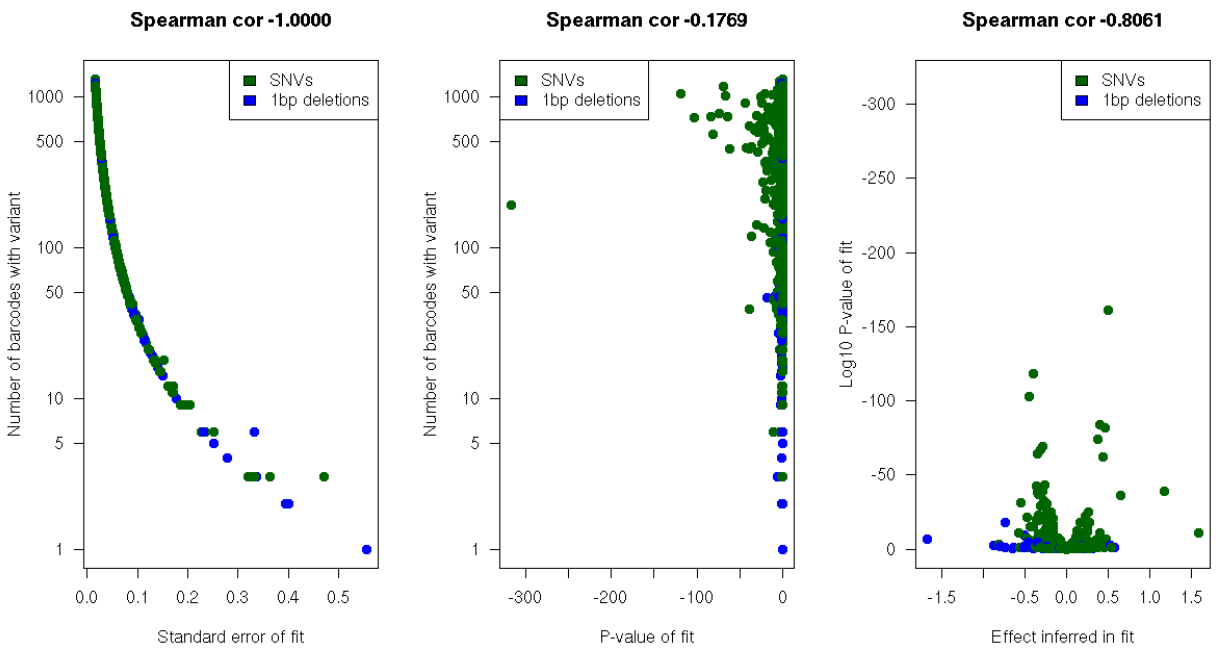
Spearman cor -1.0000
Spearman cor -0.1769
Spearman cor -0.8061
SNVs
1bp deletions
Number of barcodes with variant
Standard error of fit
P-value of fit
Log10 P-value of fit
Effect inferred in fit

**TERT**

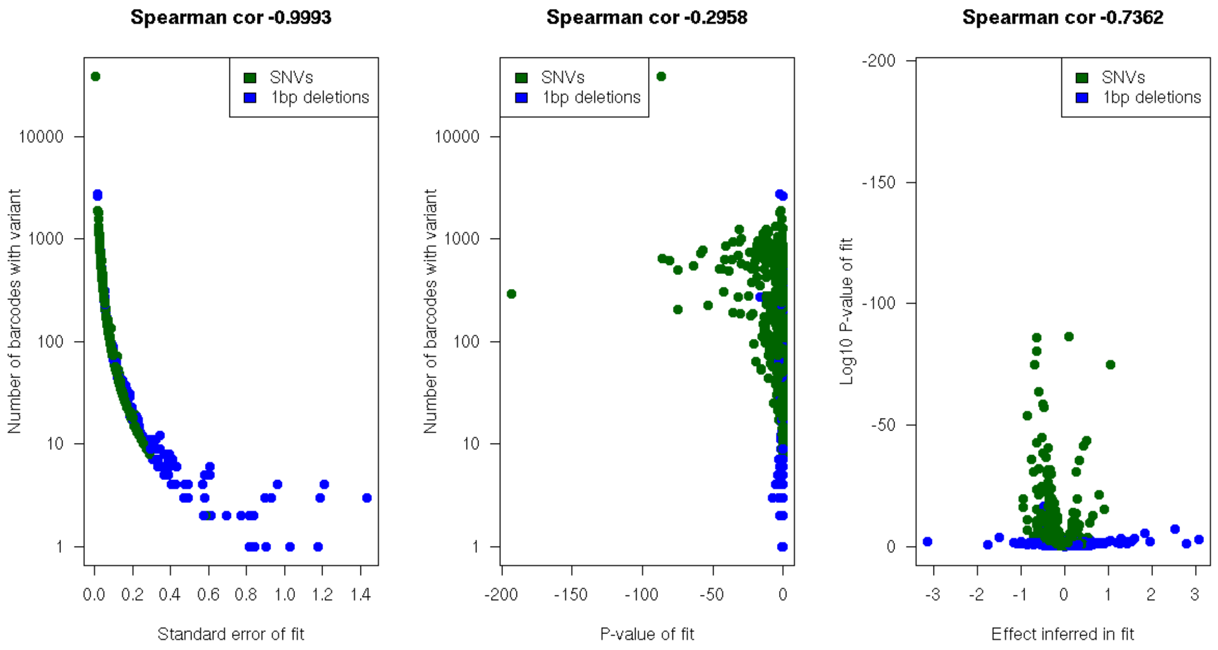
Spearman cor -0.9993
Spearman cor -0.2958
Spearman cor -0.7362
SNVs
1bp deletions
Number of barcodes with variant
Standard error of fit
P-value of fit
Log10 P-value of fit
Effect inferred in fit

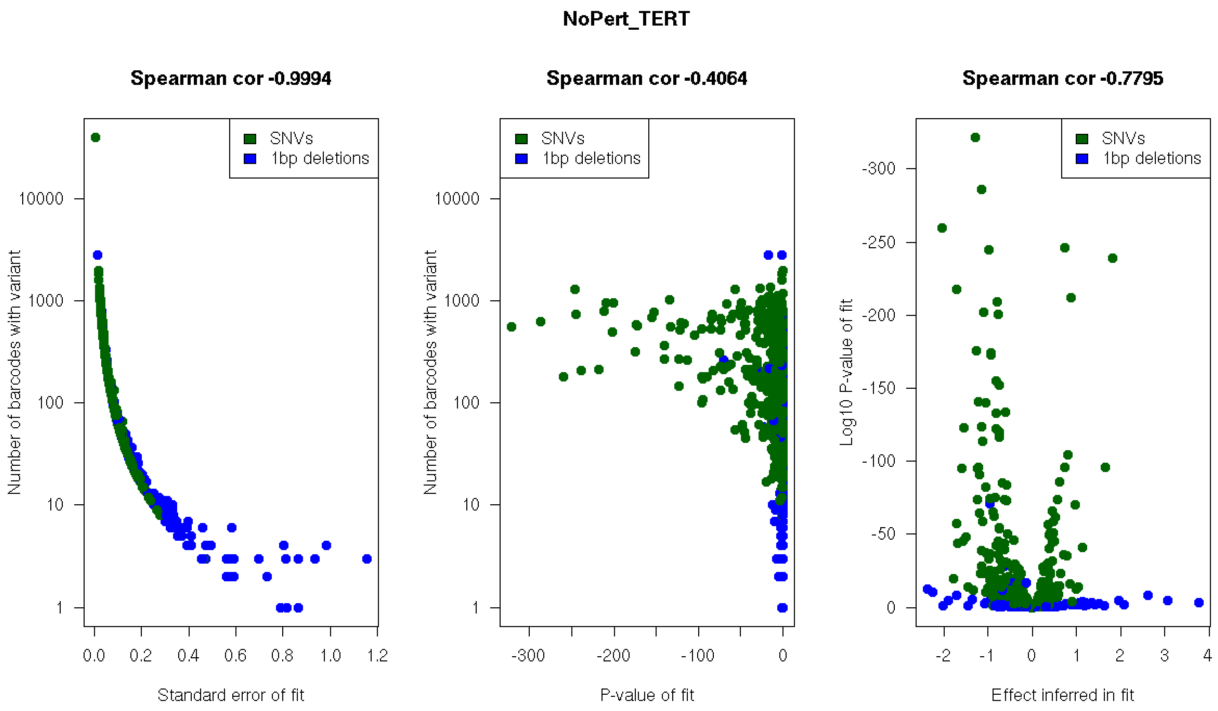
NoPert_TERT
Spearman cor -0.9994
Spearman cor -0.4064
Spearman cor -0.7795
SNVs
1bp deletions
Number of barcodes with variant
Log10 P-value of fit
Standard error of fit
P-value of fit
Effect inferred in fit

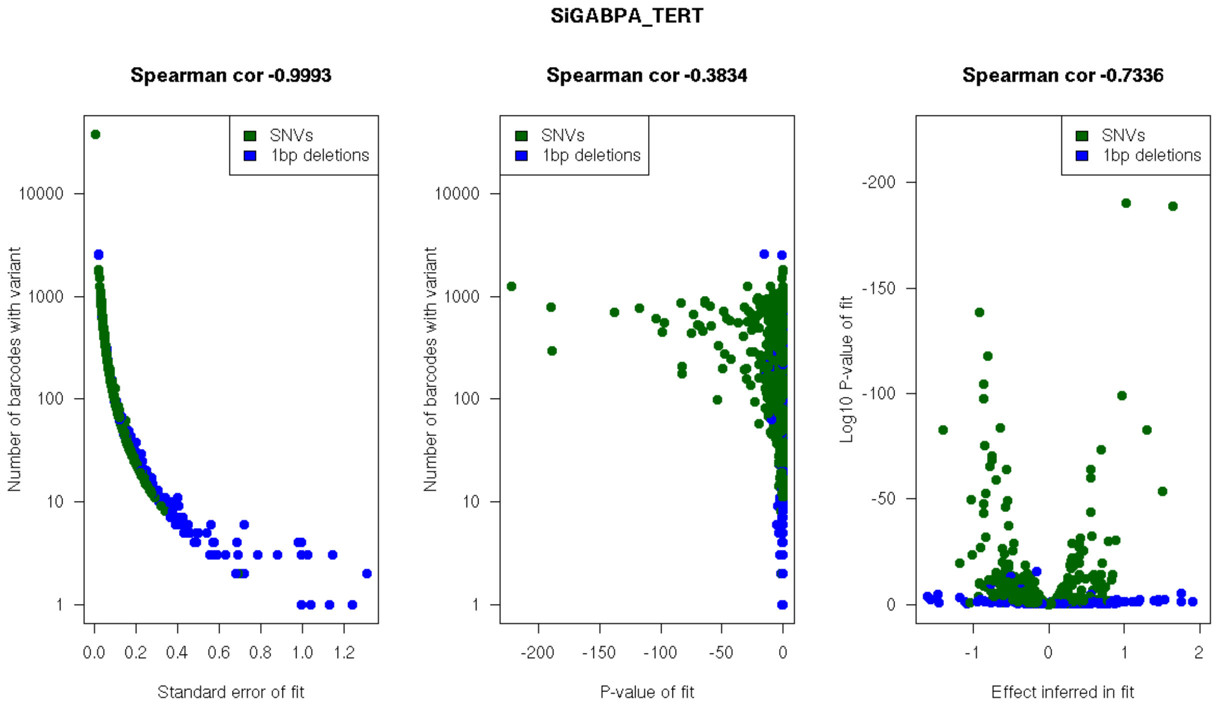
SiGABPA_TERT
Spearman cor -0.9993
Spearman cor -0.3834
Spearman cor -0.7336
SNVs
1bp deletions
Number of barcodes with variant
Log10 P-value of fit
Standard error of fit
P-value of fit
Effect inferred in fit

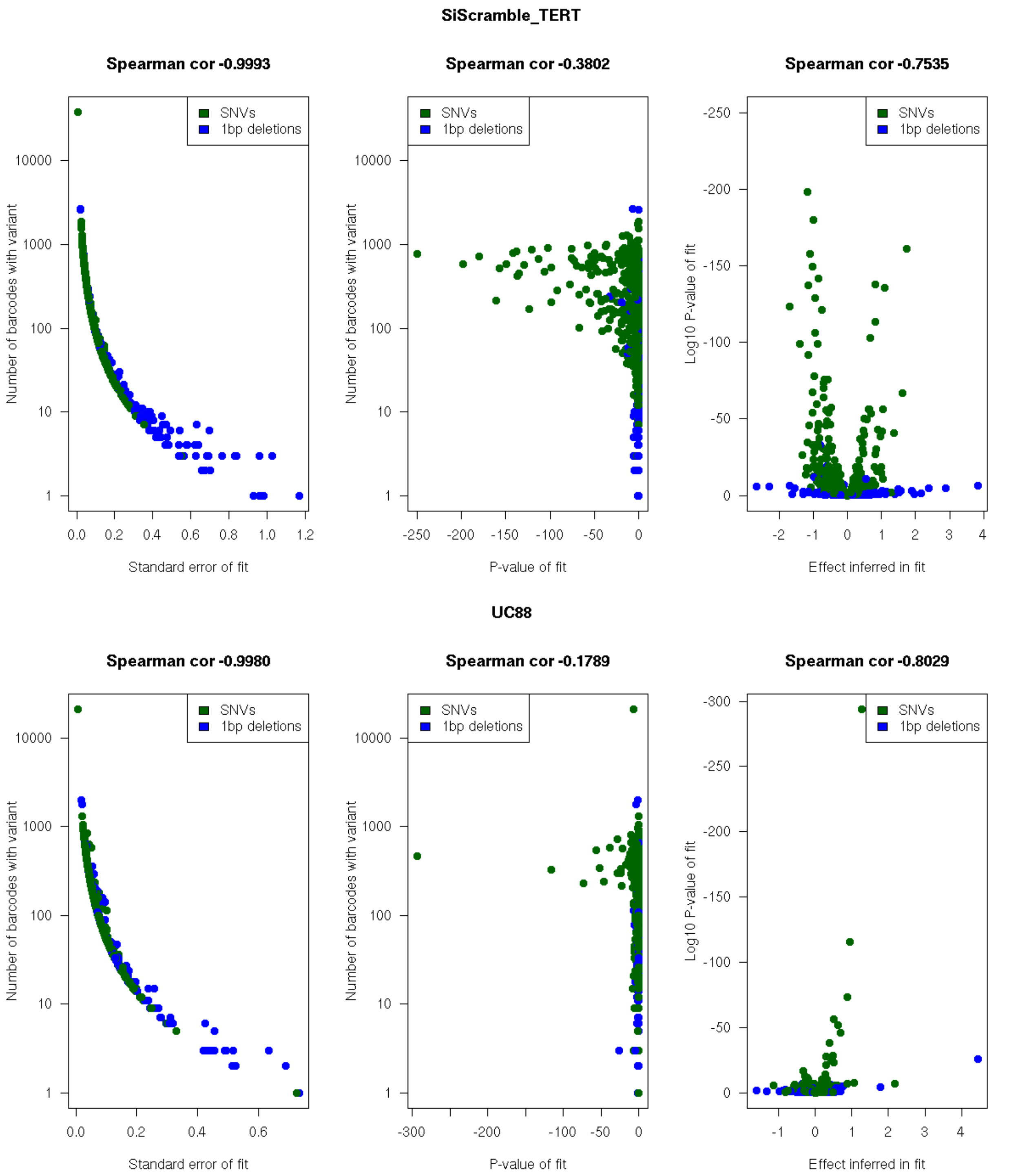

SiScramble_TERT
Spearman cor -0.9993
Spearman cor -0.3802
Spearman cor -0.7535
SNVs
1bp deletions
Number of barcodes with variant
Standard error of fit
P-value of fit
Log10 P-value of fit
Effect inferred in fit
UC88
Spearman cor -0.9980
Spearman cor -0.1789
Spearman cor -0.8029

ZFAND3

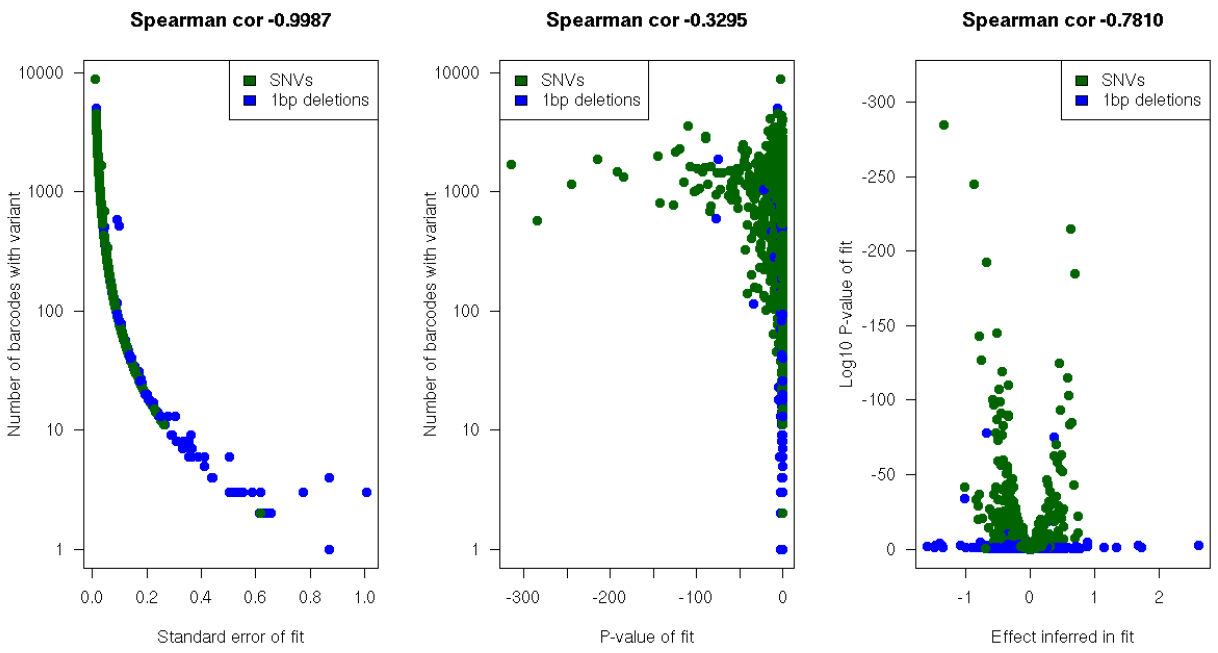

ZRSh13

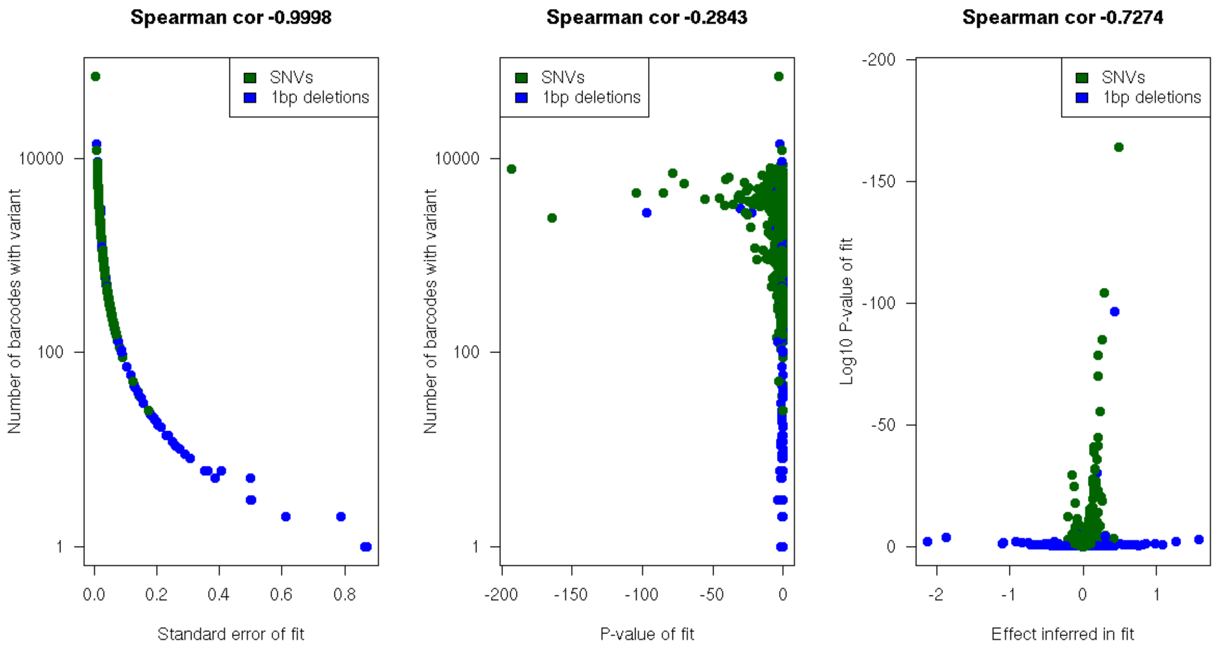

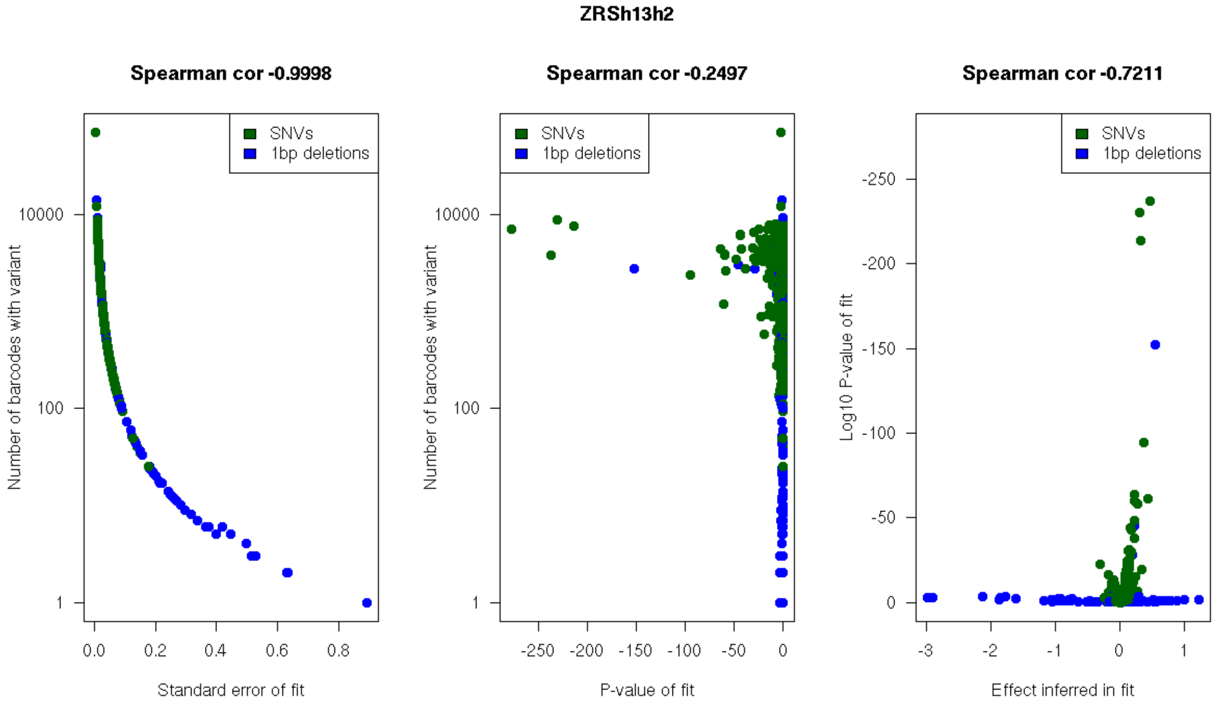

**Quality of fit measures for variant effects determined by 29 linear regression models.** For the 29 experiments (combining three technical replicates each), the left panel shows the standard error of the variant effect fit versus the number of tags/barcodes per variant, the middle panel shows the p-value ($log_{10}$ transformed) from the fit variant effects versus the number tag/barcodes per variant and the right panel shows a volcano plot (inferred variant effect vs. the $log_{10}$ transformed p-value).

#### *Supplementary Figure 5*

**A**

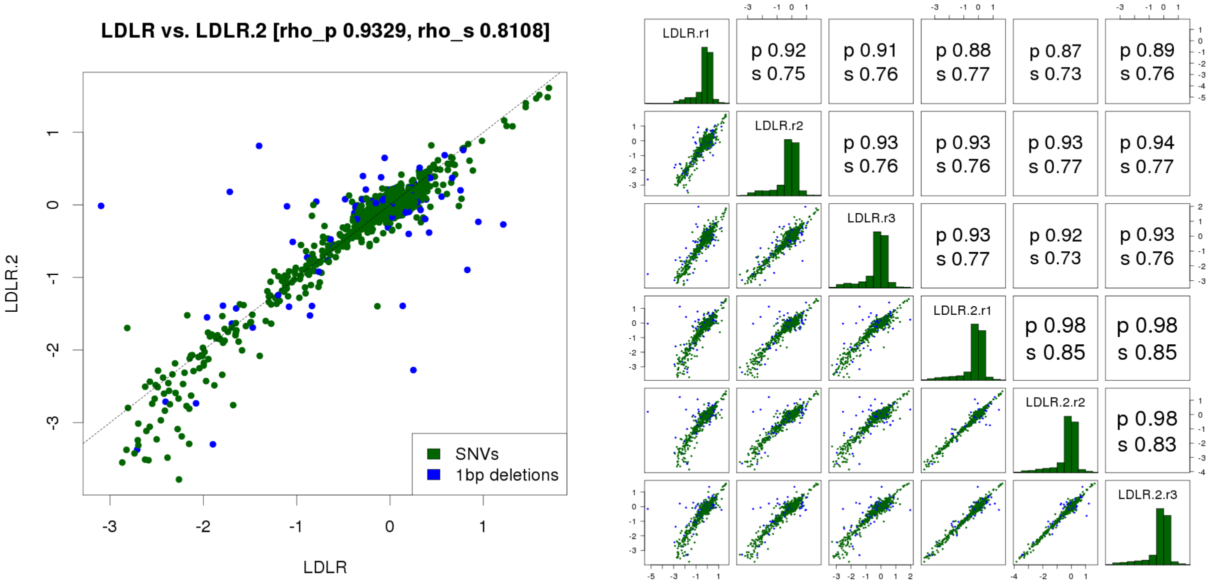

**B**

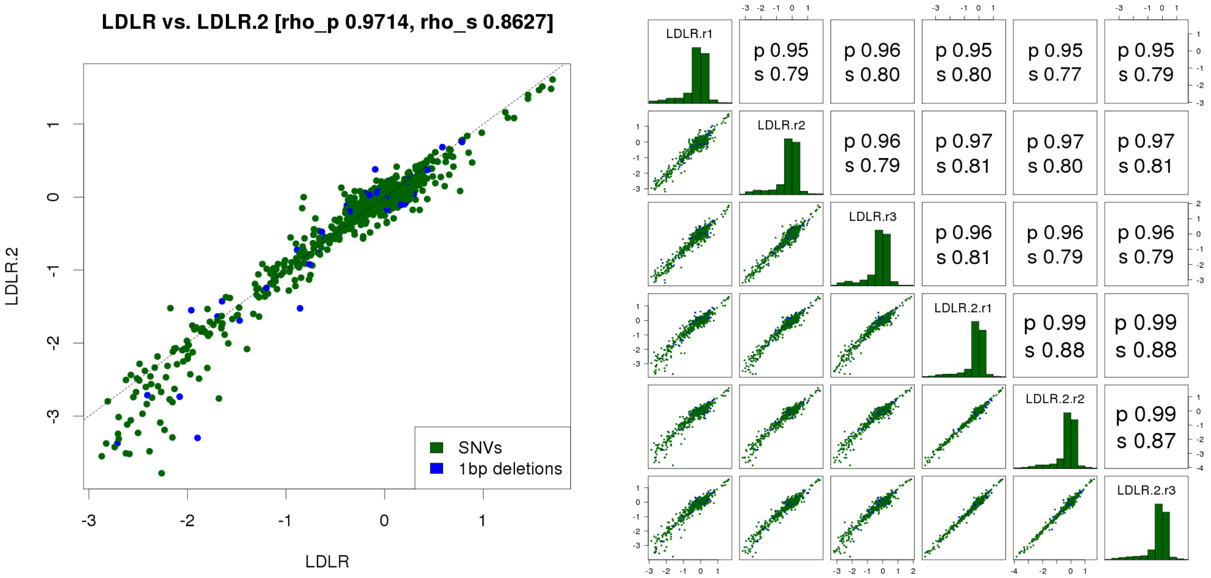

**Biological replicate correlation for LDLR.** Transfection and biological replicate correlation in LDLR experiments, before **(A)** and after **(B)** filtering for a minimum of 10 tags per variant.

***Supplementary Figure 6***

**A**

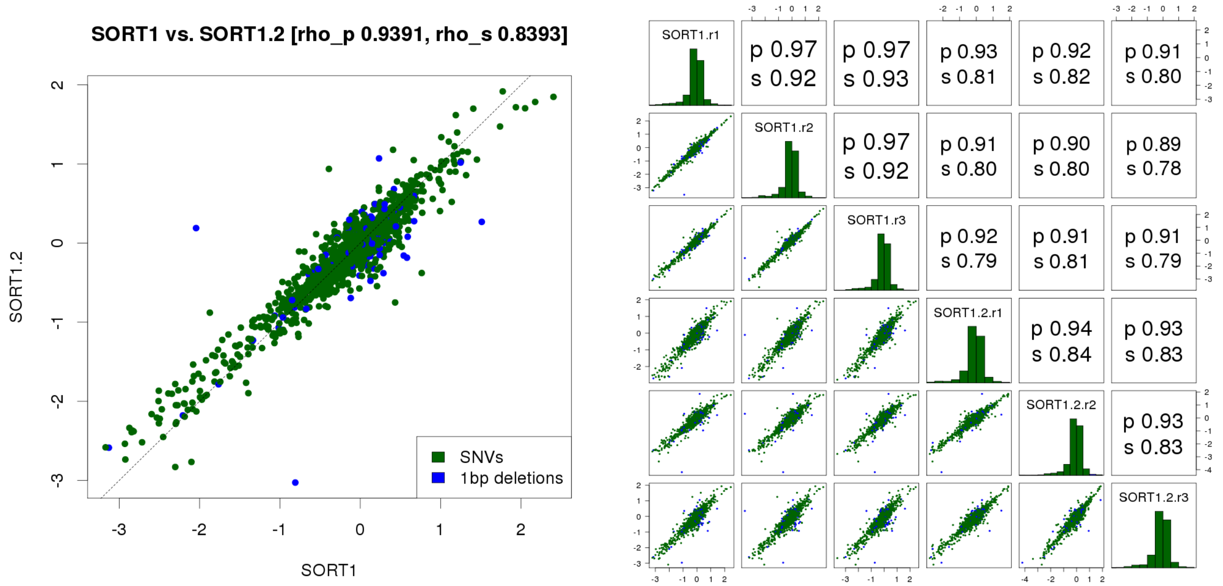

**B**

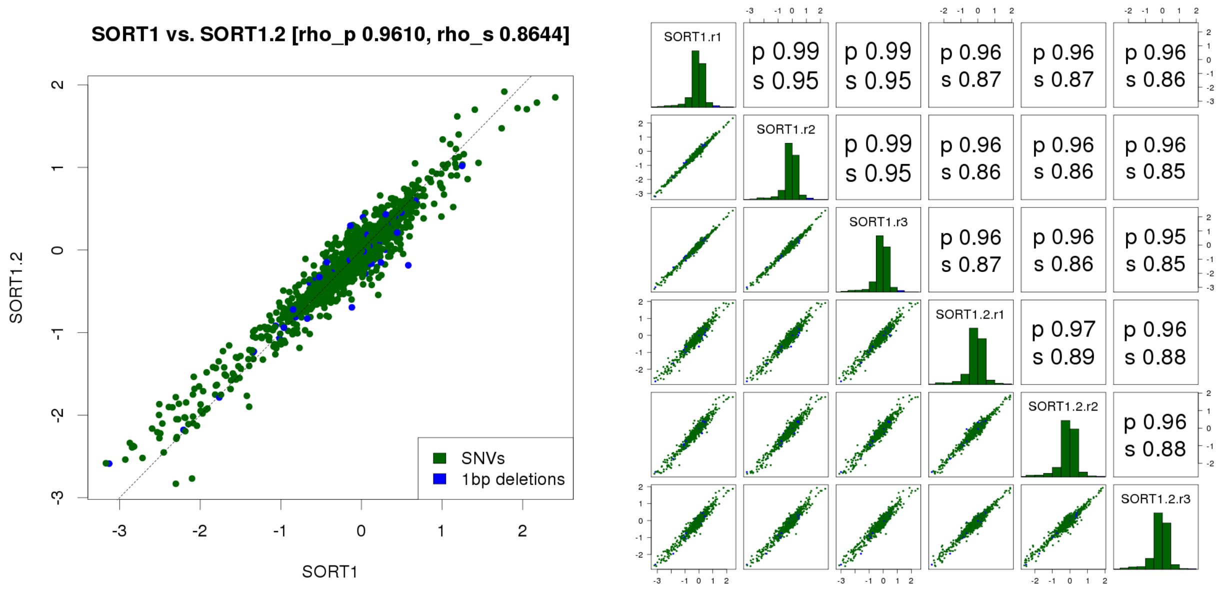

**Biological replicate correlation for SORT1.** Transfection and biological replicate correlation in SORT experiments, before **(A)** and after **(B)** filtering for a minimum of 10 tags per variant.

*Supplementary Figure 7*

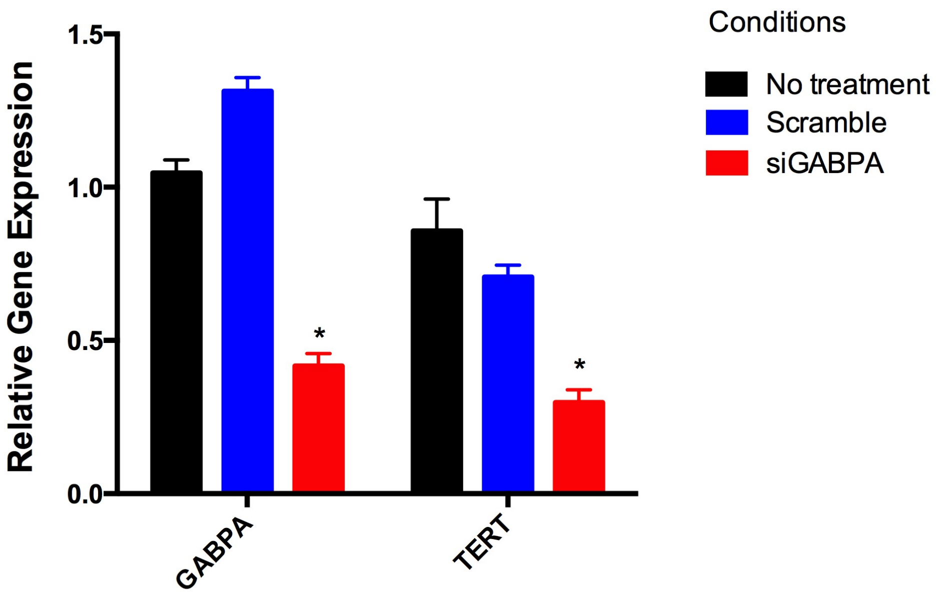

**Quantitative RT-PCR validation results of TERT-GBM short-interfering RNA experiments.** To measure siRNA knockdown efficiency, qPCR was performed using the Ambion Power SYBR Green Cells-to-Ct kit to measure mRNA abundance of *GABPA* and *TERT* with primer sequences previously used (see Methods). Relative expression levels were calculated using the deltaCT method against the housekeeping gene *GUSB*.

*Supplementary Figure 8*

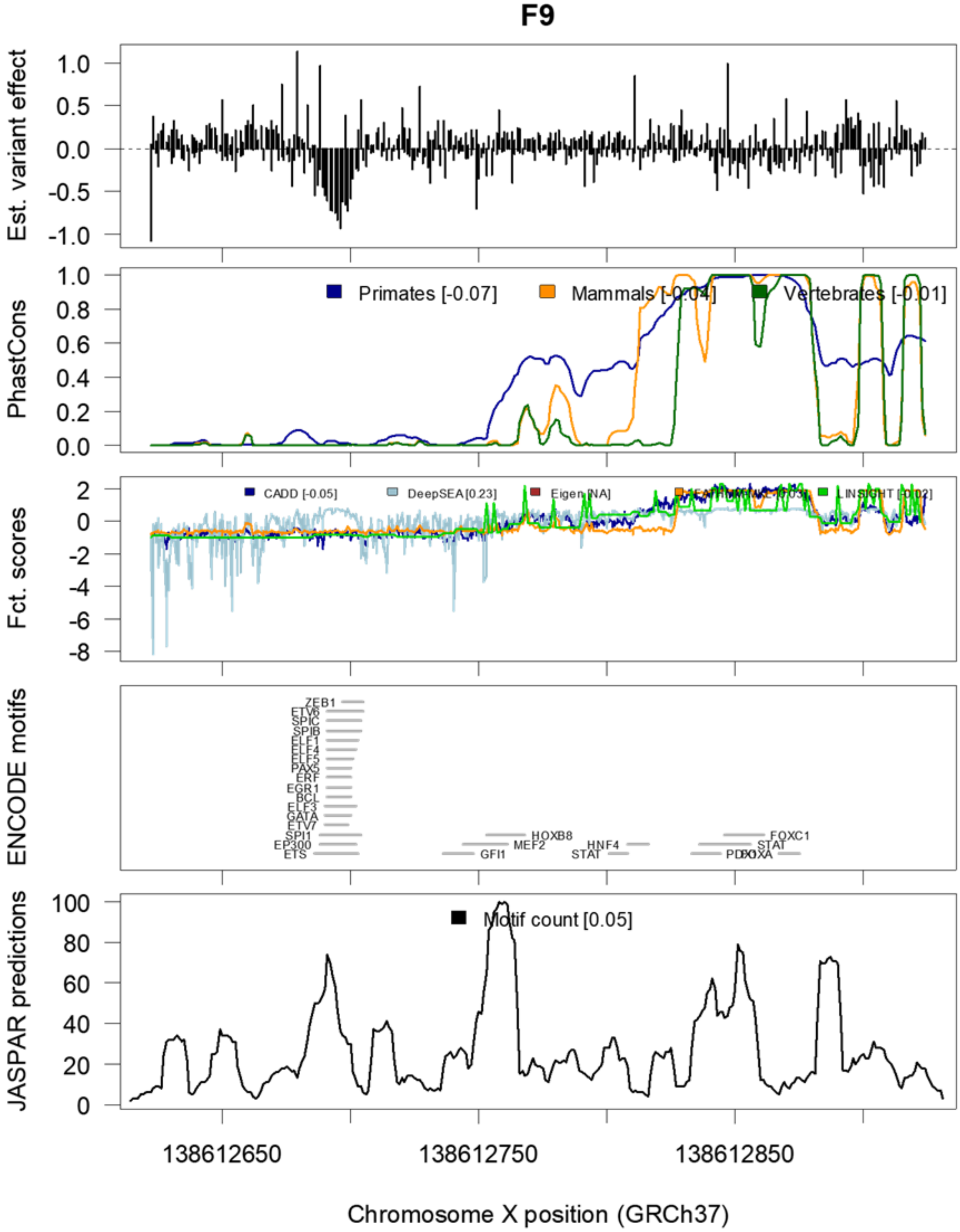

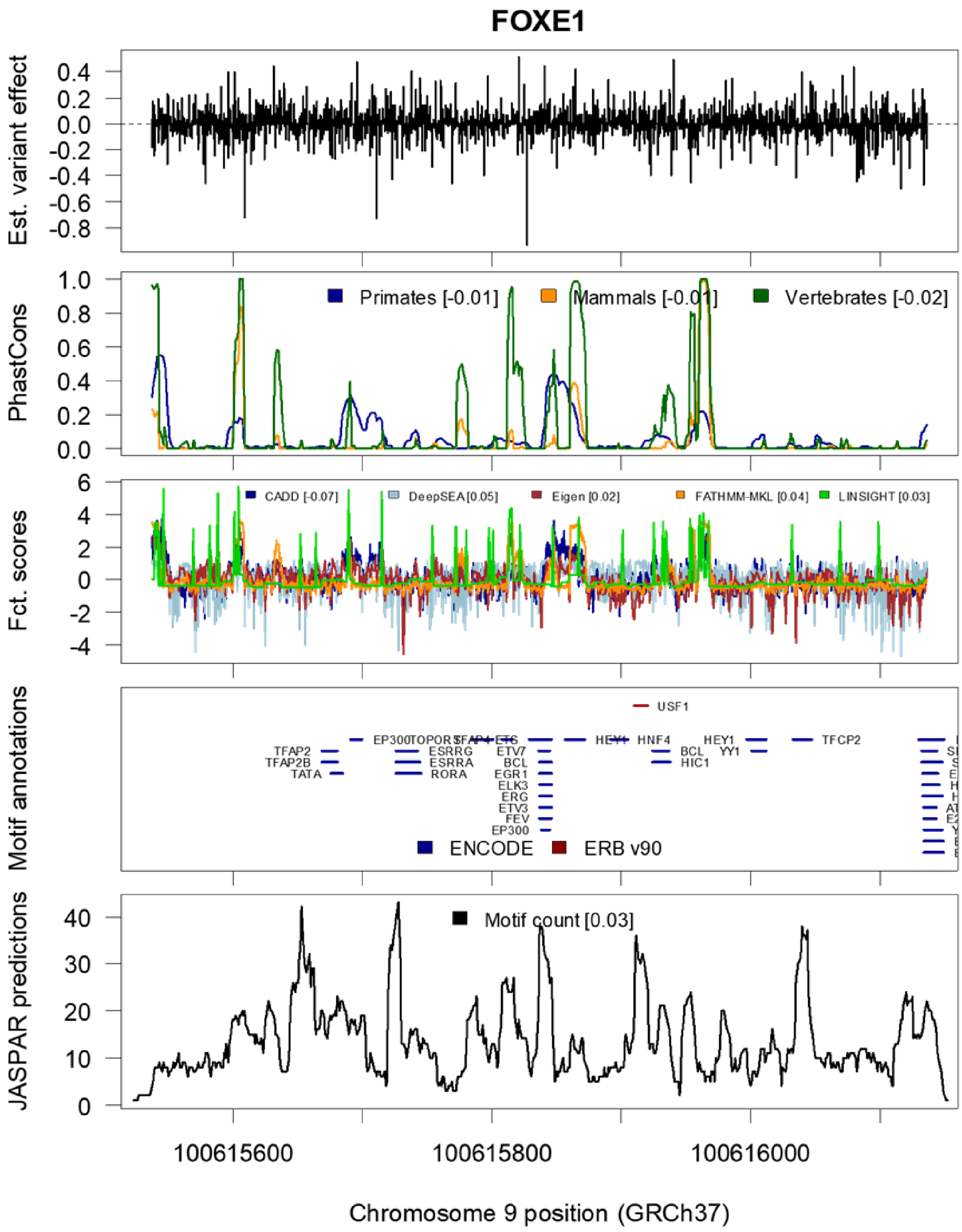

FOXE1
Est. variant effect
PhastCons
Primates [-0.01]
Mammals [-0.01]
Vertebrates [-0.02]
Fct. scores
CADD [-0.07]
DeepSEA [0.05]
Eigen [0.02]
FATHMM-MKL [0.04]
LINSIGHT [0.03]
Motif annotations
USF1
TFAP2
TFAP2B
TATA
EP300
TOPORS
SFAP4
ETS
ESRRG
ESRRA
RORA
ETV7
BCL
EGR1
ELK3
ERG
ETV3
FEV
EP300
HEY1
HNF4
BCL
HIC1
HEY1
YY1
TFCP2
ENCODE
ERB v90
JASPAR predictions
Motif count [0.03]
100615600
100615800
100616000
Chromosome 9 position (GRCh37)

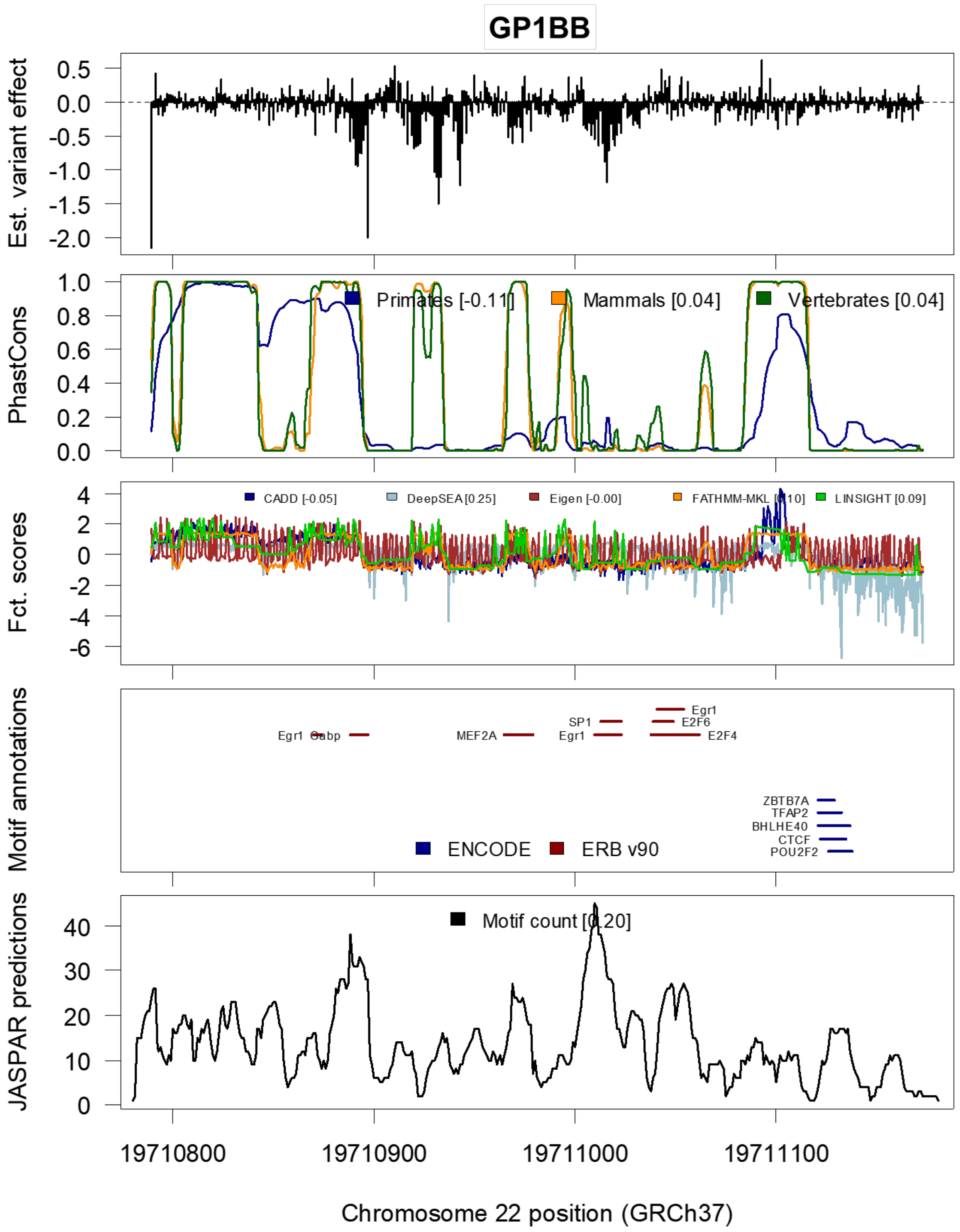
GP1BB
Est. variant effect
PhastCons
Primates [-0.11]
Mammals [0.04]
Vertebrates [0.04]
Fct. scores
CADD [-0.05]
DeepSEA [0.25]
Eigen [-0.00]
FATHMM-MKL [0.10]
LINSIGHT [0.09]
Motif annotations
Egr1
Gabp
MEF2A
SP1
Egr1
Egr1
E2F6
E2F4
ZBTB7A
TFAP2
BHLHE40
CTCF
POU2F2
ENCODE
ERB v90
JASPAR predictions
Motif count [0.20]
19710800
19710900
19711000
19711100
Chromosome 22 position (GRCh37)

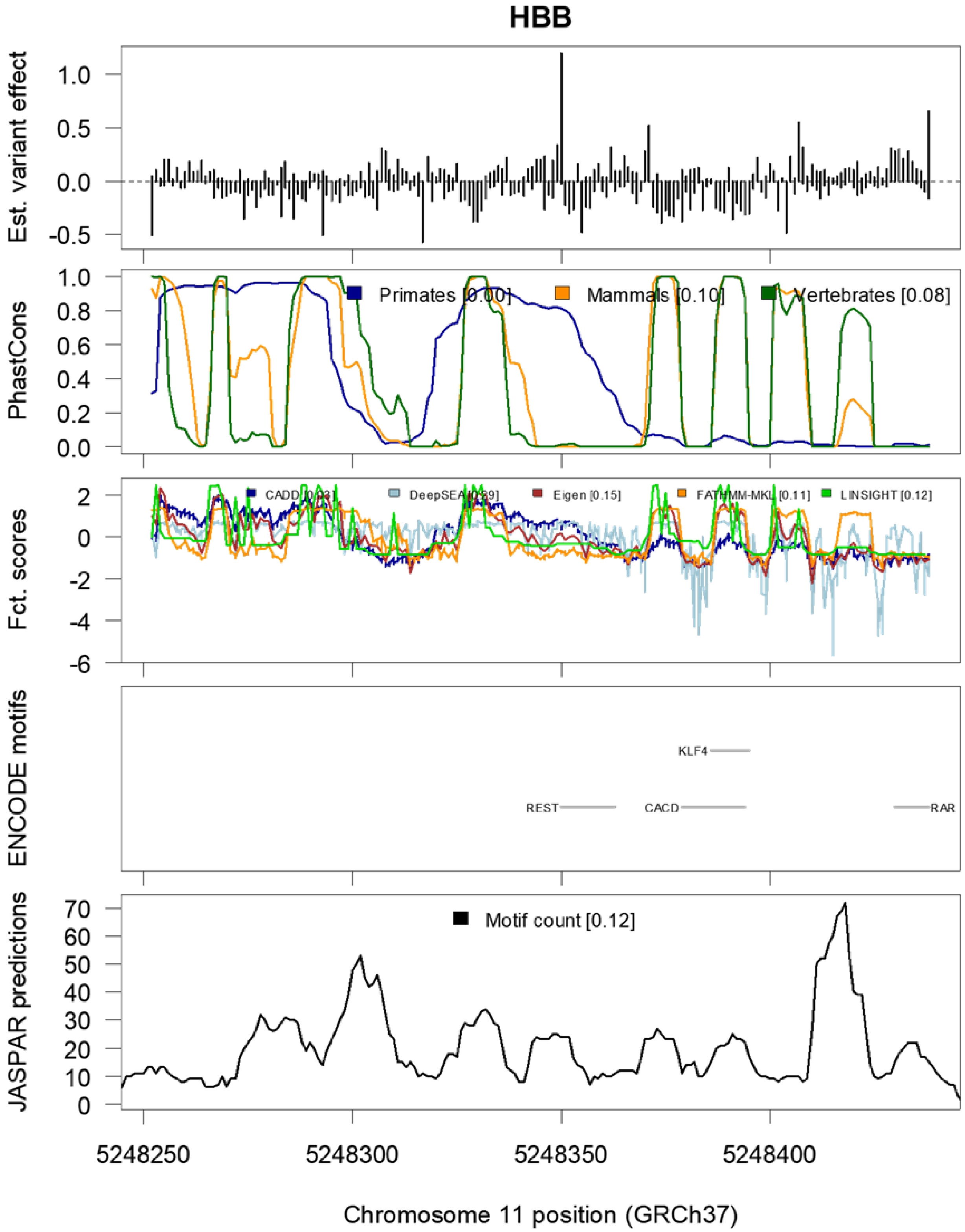
HBB
Est. variant effect
1.0
0.5
0.0
-0.5
PhastCons
Primates [0.00]
Mammals [0.10]
Vertebrates [0.08]
Fct. scores
CADD [0.03]
DeepSEA [0.09]
Eigen [0.15]
FATHMM-MKL [0.11]
LINSIGHT [0.12]
ENCODE motifs
KLF4
REST
CACD
RAR
JASPAR predictions
Motif count [0.12]
5248250
5248300
5248350
5248400
Chromosome 11 position (GRCh37)

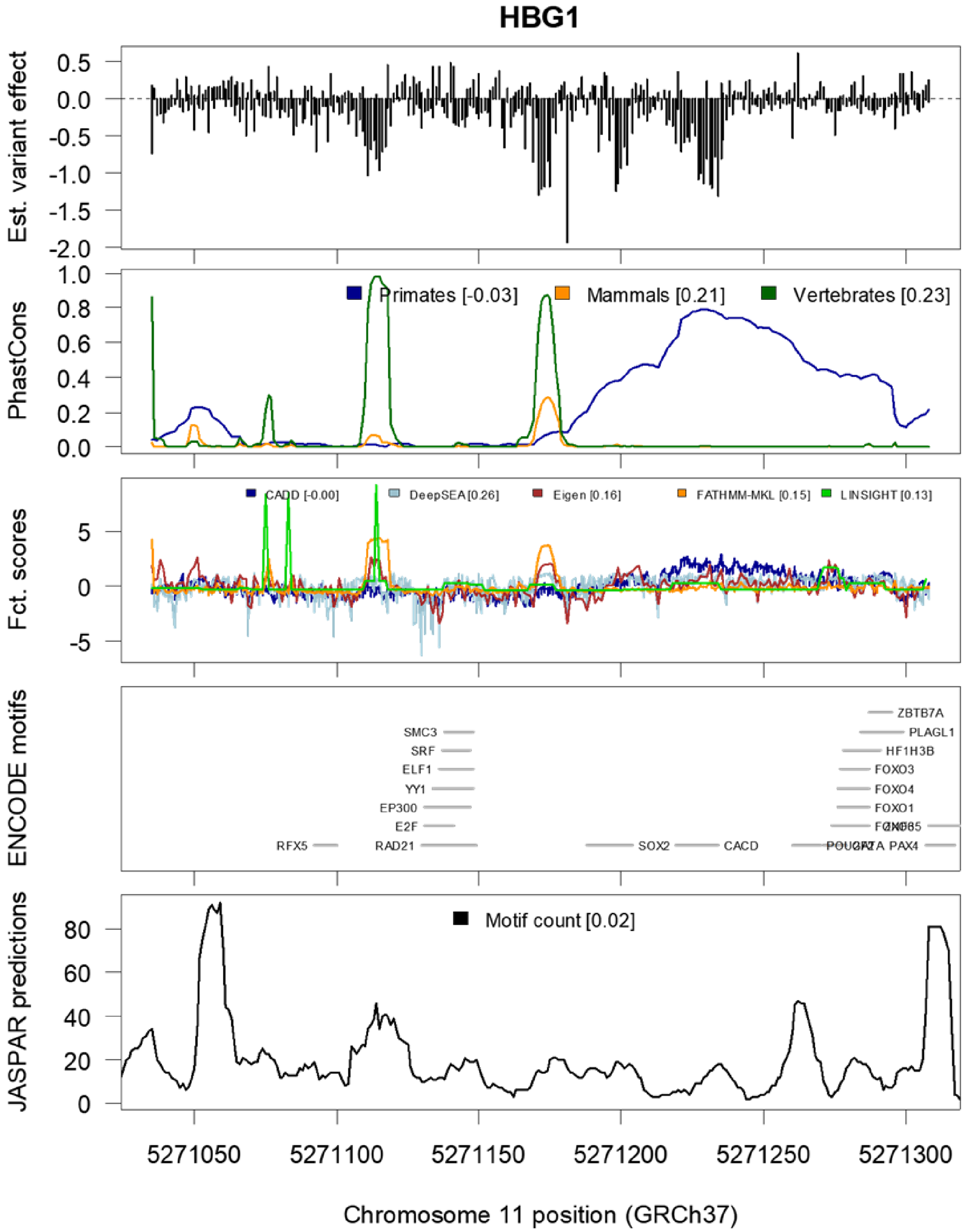

HBG1
Est. variant effect
PhastCons
Primates [-0.03]
Mammals [0.21]
Vertebrates [0.23]
Fct. scores
CADD [-0.00]
DeepSEA [0.26]
Eigen [0.16]
FATHMM-MKL [0.15]
LINSIGHT [0.13]
ENCODE motifs
JASPAR predictions
Motif count [0.02]
5271050
5271100
5271150
5271200
5271250
5271300
Chromosome 11 position (GRCh37)

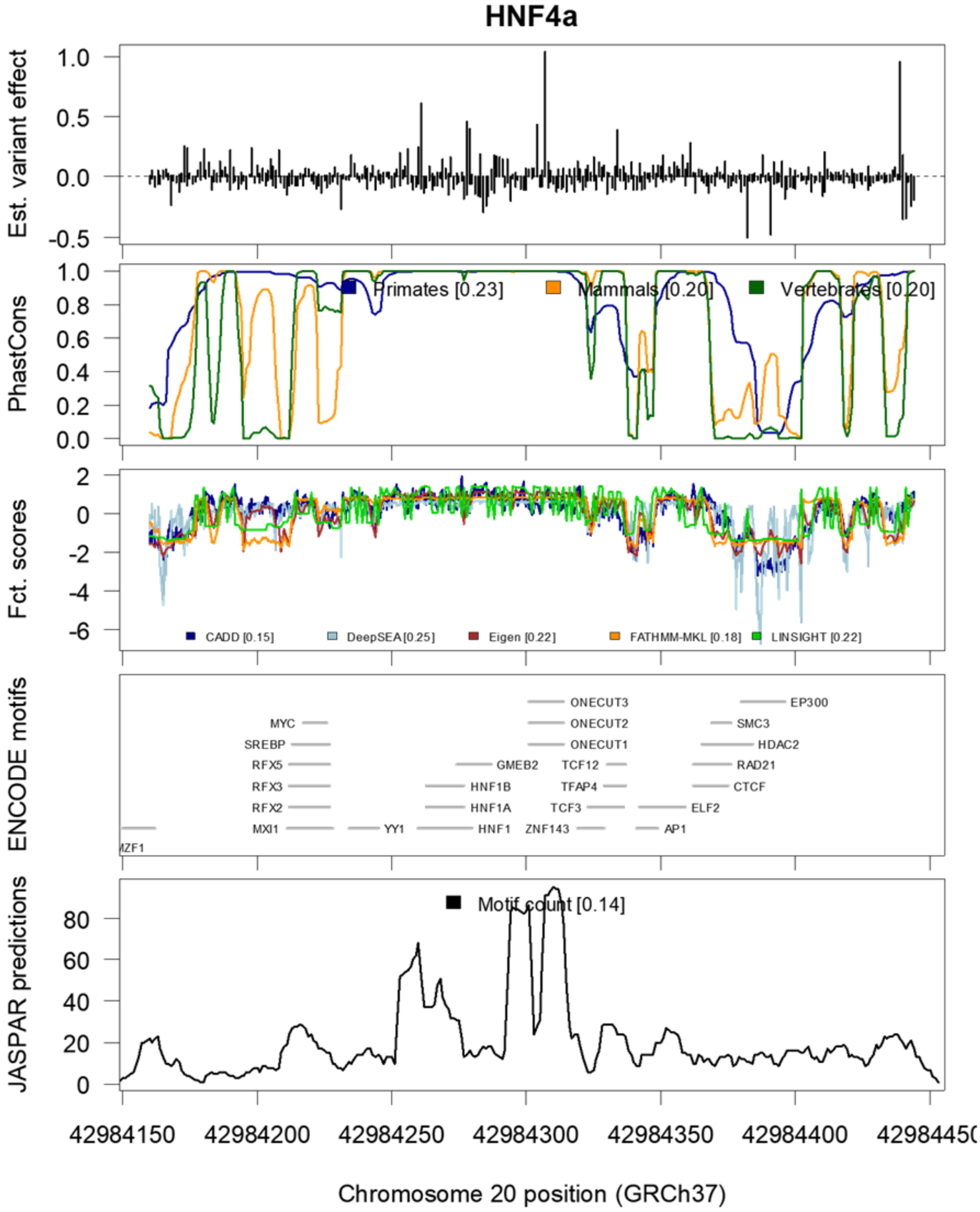
HNF4a
Est. variant effect
PhastCons
Primates [0.23]
Mammals [0.20]
Vertebrates [0.20]
Fct. scores
CADD [0.15]
DeepSEA [0.25]
Eigen [0.22]
FATHMM-MKL [0.18]
LINSIGHT [0.22]
ENCODE motifs
ONECUT3
EP300
MYC
ONECUT2
SMC3
SREBP
ONECUT1
HDAC2
RFX5
GMEB2
TCF12
RAD21
RFX3
HNF1B
TFAP4
CTCF
RFX2
HNF1A
TCF3
ELF2
MXI1
YY1
HNF1
ZNF143
AP1
JASPAR predictions
Motif count [0.14]
Chromosome 20 position (GRCh37)

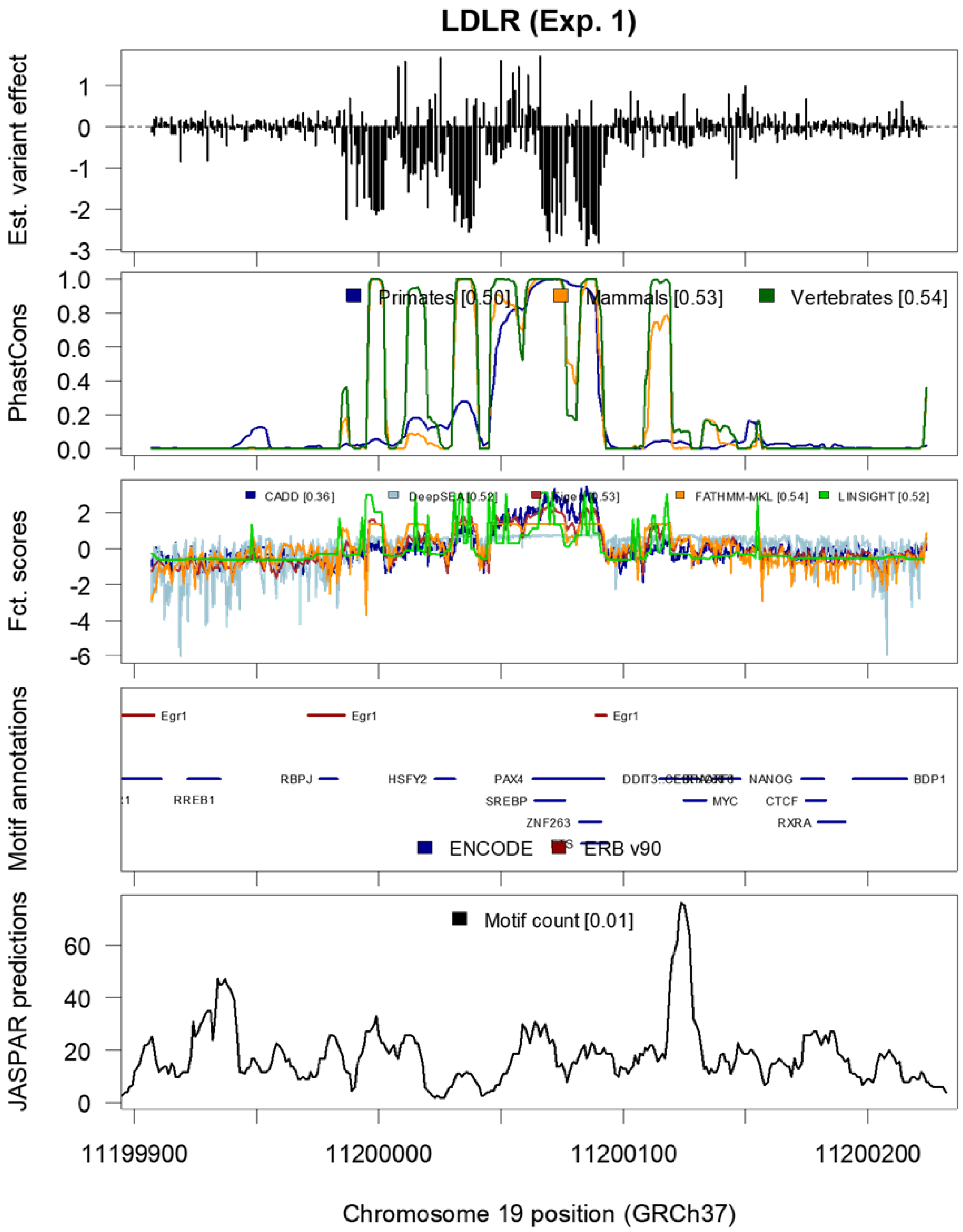
LDLR (Exp. 1)
Est. variant effect
PhastCons
Primates [0.50]
Mammals [0.53]
Vertebrates [0.54]
Fct. scores
CADD [0.36]
FATHMM-MKL [0.54]
LINSIGHT [0.52]
Motif annotations
Egr1
RBPJ
HSFY2
PAX4
SREBP
ZNF263
RREB1
MYC
NANOG
CTCF
RXRA
BDP1
ENCODE
ERB v90
JASPAR predictions
Motif count [0.01]
11199900
11200000
11200100
11200200
Chromosome 19 position (GRCh37)

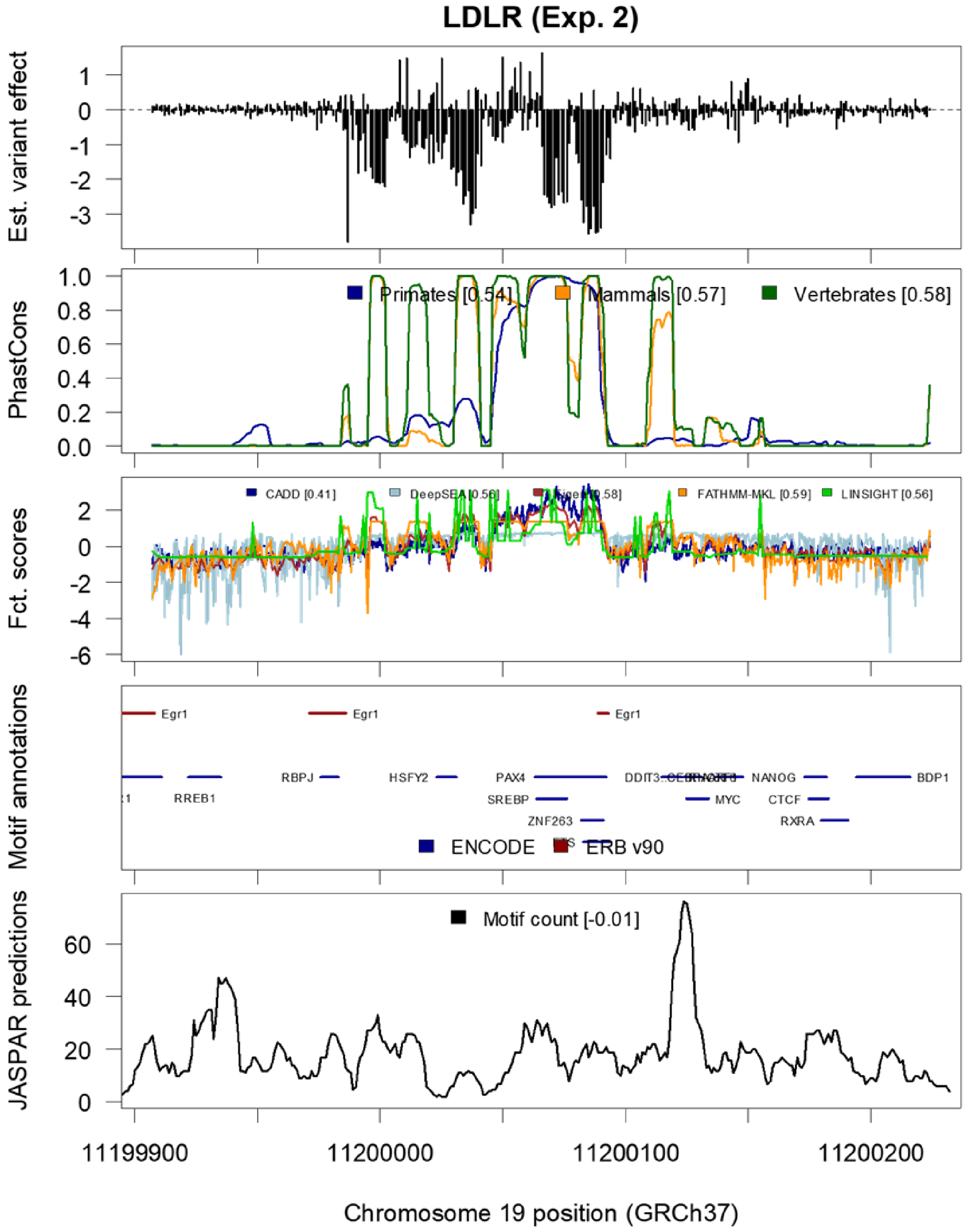

LDLR (Exp. 2)
Est. variant effect
PhastCons
Primates [0.54]
Mammals [0.57]
Vertebrates [0.58]
Fct. scores
CADD [0.41]
FATHMM-MKL [0.59]
LINSIGHT [0.56]
Motif annotations
Egr1
RBPJ
HSFY2
PAX4
NANOG
BDP1
RREB1
SREBP
MYC
CTCF
ZNF263
RXRA
ENCODE
ERB v90
JASPAR predictions
Motif count [-0.01]
11199900
11200000
11200100
11200200
Chromosome 19 position (GRCh37)

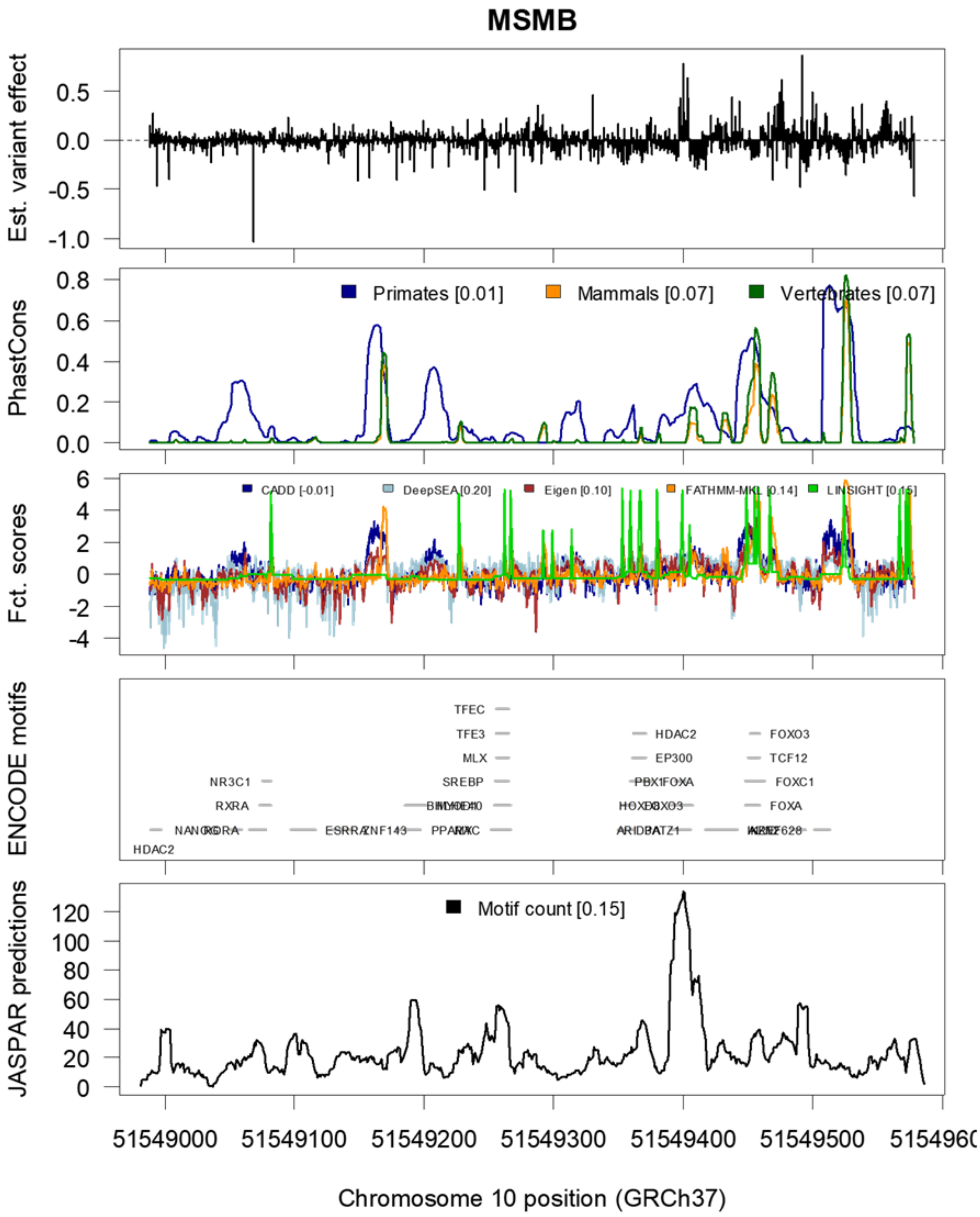

MSMB
Est. variant effect
PhastCons
Primates [0.01]
Mammals [0.07]
Vertebrates [0.07]
Fct. scores
CADD [-0.01]
DeepSEA [0.20]
Eigen [0.10]
FATHMM-MKL [0.14]
LINSIGHT [0.15]
ENCODE motifs
JASPAR predictions
Motif count [0.15]
51549000
51549100
51549200
51549300
51549400
51549500
Chromosome 10 position (GRCh37)

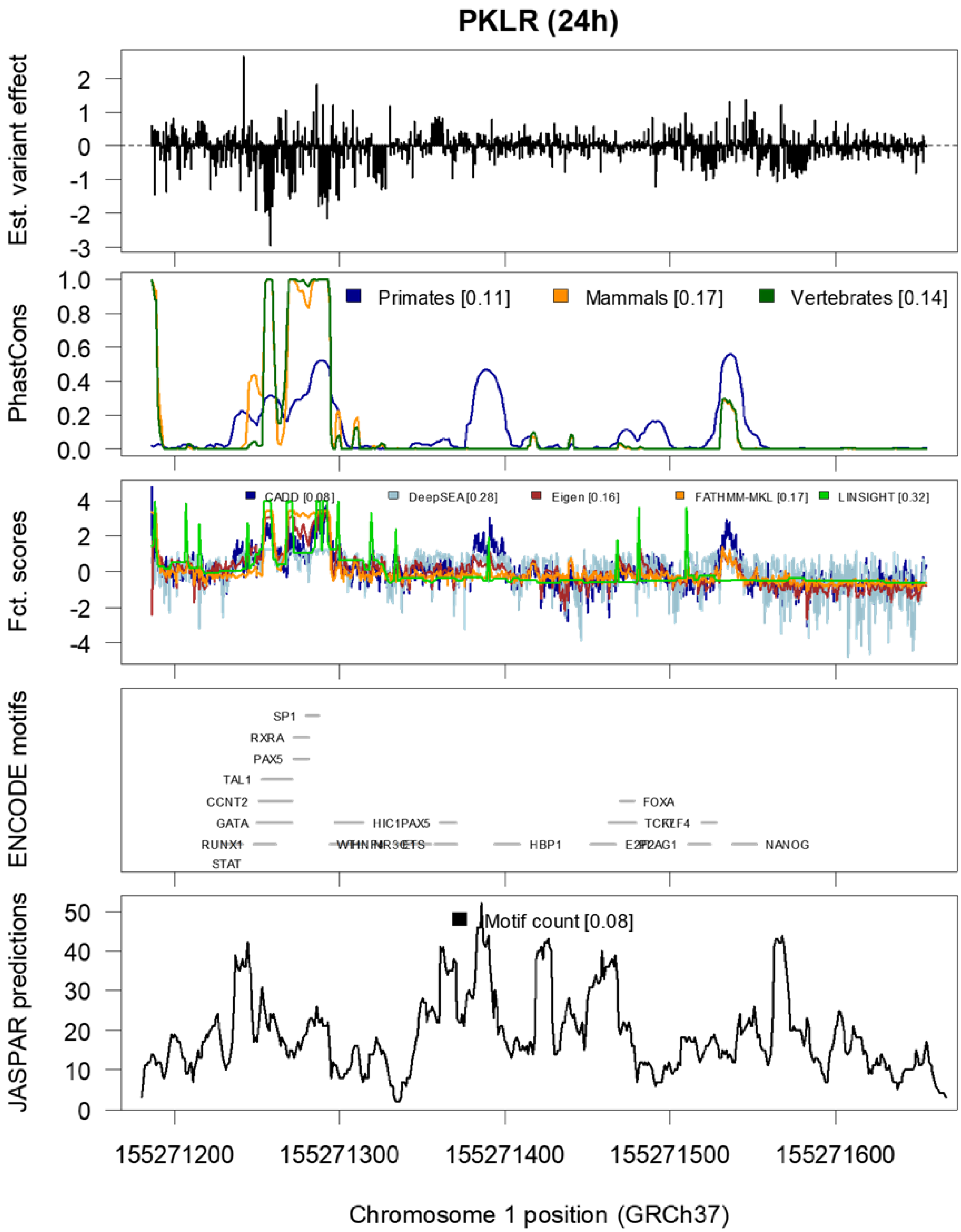
PKLR (24h)
Est. variant effect
PhastCons
Primates [0.11]
Mammals [0.17]
Vertebrates [0.14]
Fct. scores
CADD [0.08]
DeepSEA [0.28]
Eigen [0.16]
FATHMM-MKL [0.17]
LINSIGHT [0.32]
ENCODE motifs
SP1
RXRA
PAX5
TAL1
CCNT2
GATA
RUNX1
STAT
HIC1PAX5
HBP1
FOXA
NANOG
JASPAR predictions
Motif count [0.08]
155271200
155271300
155271400
155271500
155271600
Chromosome 1 position (GRCh37)

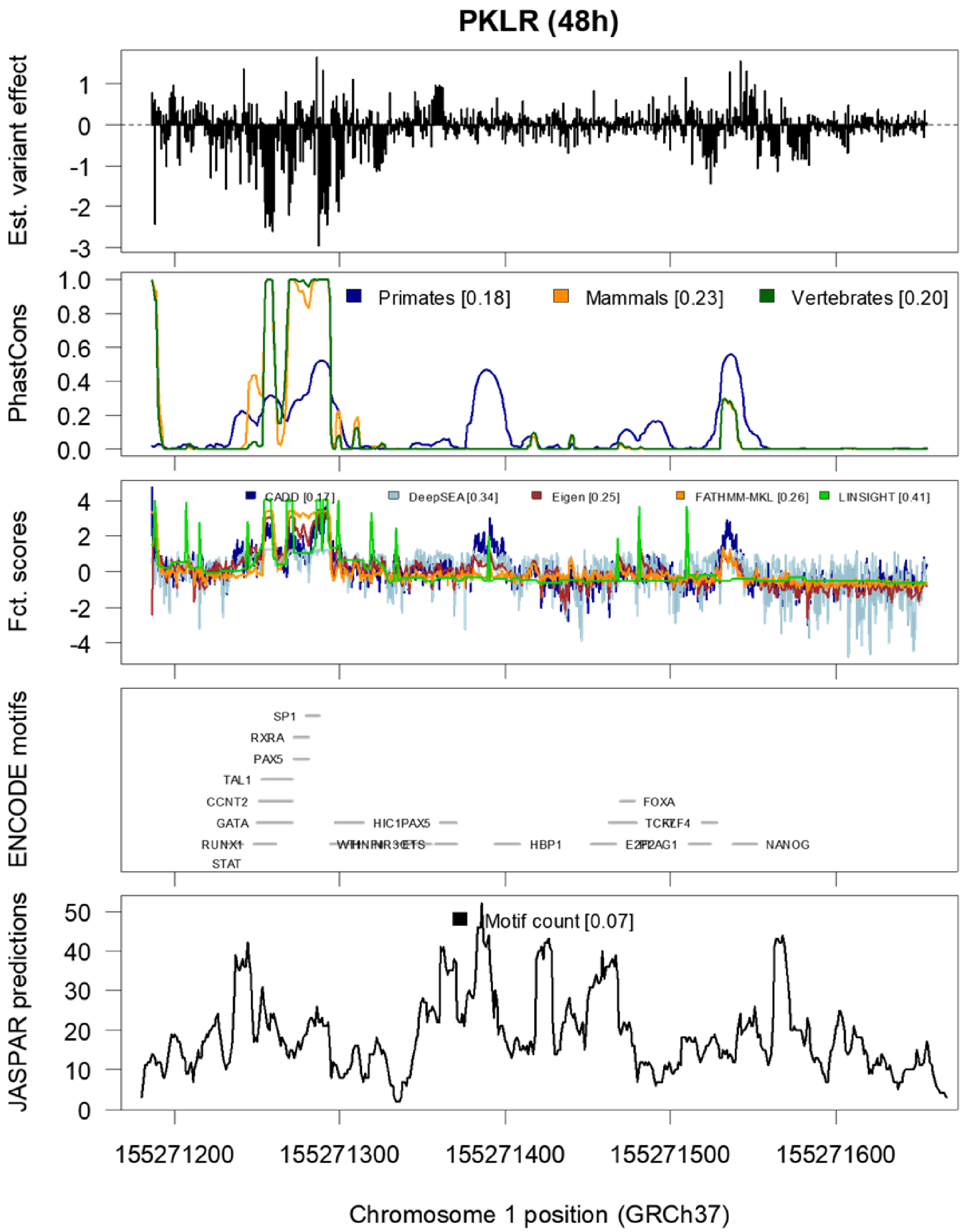
PKLR (48h)
Est. variant effect
PhastCons
Primates [0.18]
Mammals [0.23]
Vertebrates [0.20]
Fct. scores
CADD [0.17]
DeepSEA [0.34]
Eigen [0.25]
FATHMM-MKL [0.26]
LINSIGHT [0.41]
ENCODE motifs
SP1
RXRA
PAX5
TAL1
CCNT2
GATA
RUNX1
STAT
HIC1
PAX5
HBP1
FOXA
NANOG
JASPAR predictions
Motif count [0.07]
155271200
155271300
155271400
155271500
155271600
Chromosome 1 position (GRCh37)

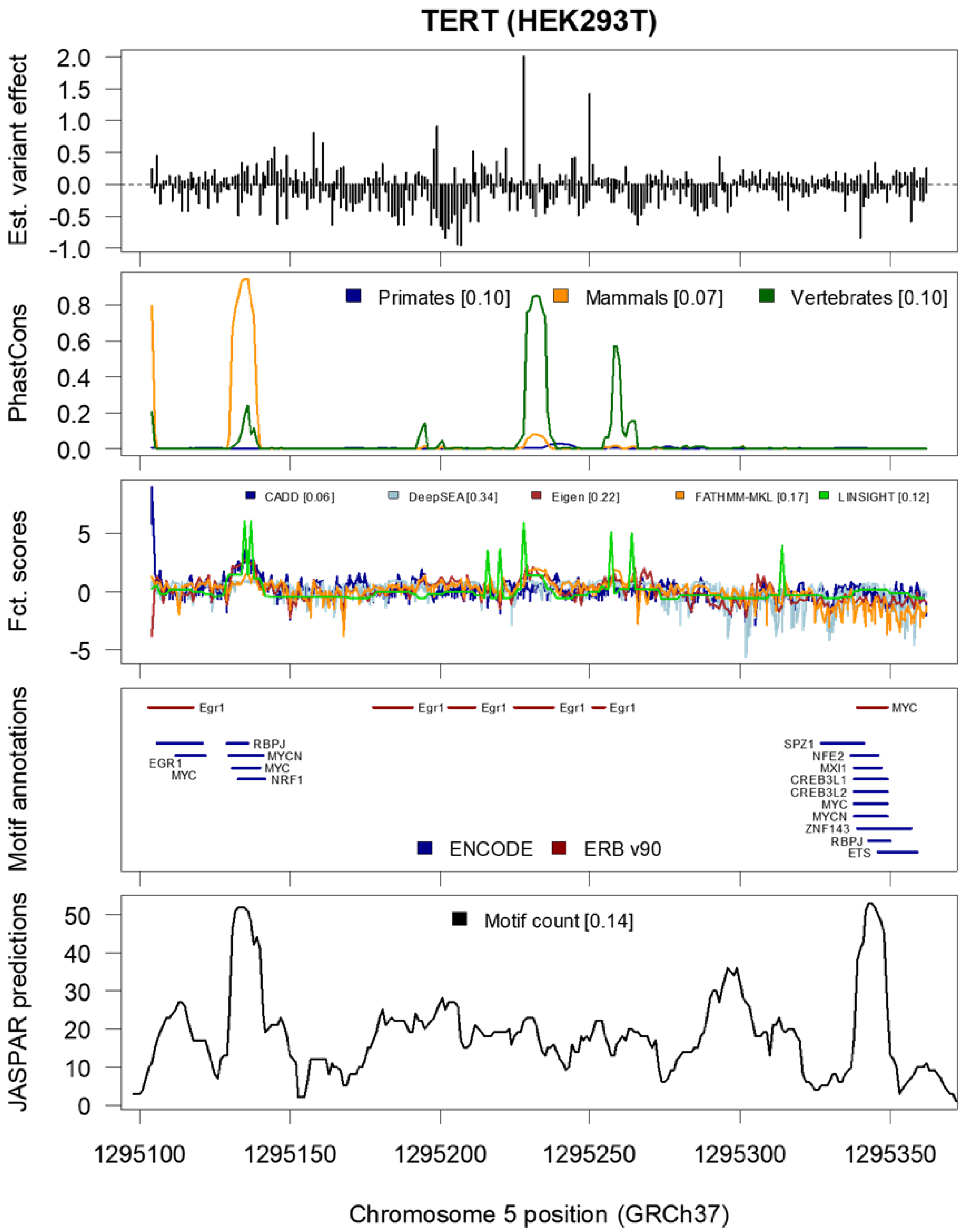
TERT (HEK293T)
Est. variant effect
2.0
1.5
1.0
0.5
0.0
-0.5
-1.0
PhastCons
Primates [0.10]
Mammals [0.07]
Vertebrates [0.10]
0.8
0.6
0.4
0.2
0.0
Fct. scores
CADD [0.06]
DeepSEA [0.34]
Eigen [0.22]
FATHMM-MKL [0.17]
LINSIGHT [0.12]
5
0
-5
Motif annotations
Egr1
Egr1
Egr1
Egr1
Egr1
MYC
RBPJ
MYCN
MYC
NRF1
EGR1
MYC
SPZ1
NFE2
MXI1
CREB3L1
CREB3L2
MYC
MYCN
ZNF143
RBPJ
ETS
ENCODE
ERB v90
JASPAR predictions
Motif count [0.14]
50
40
30
20
10
0
1295100
1295150
1295200
1295250
1295300
1295350
Chromosome 5 position (GRCh37)

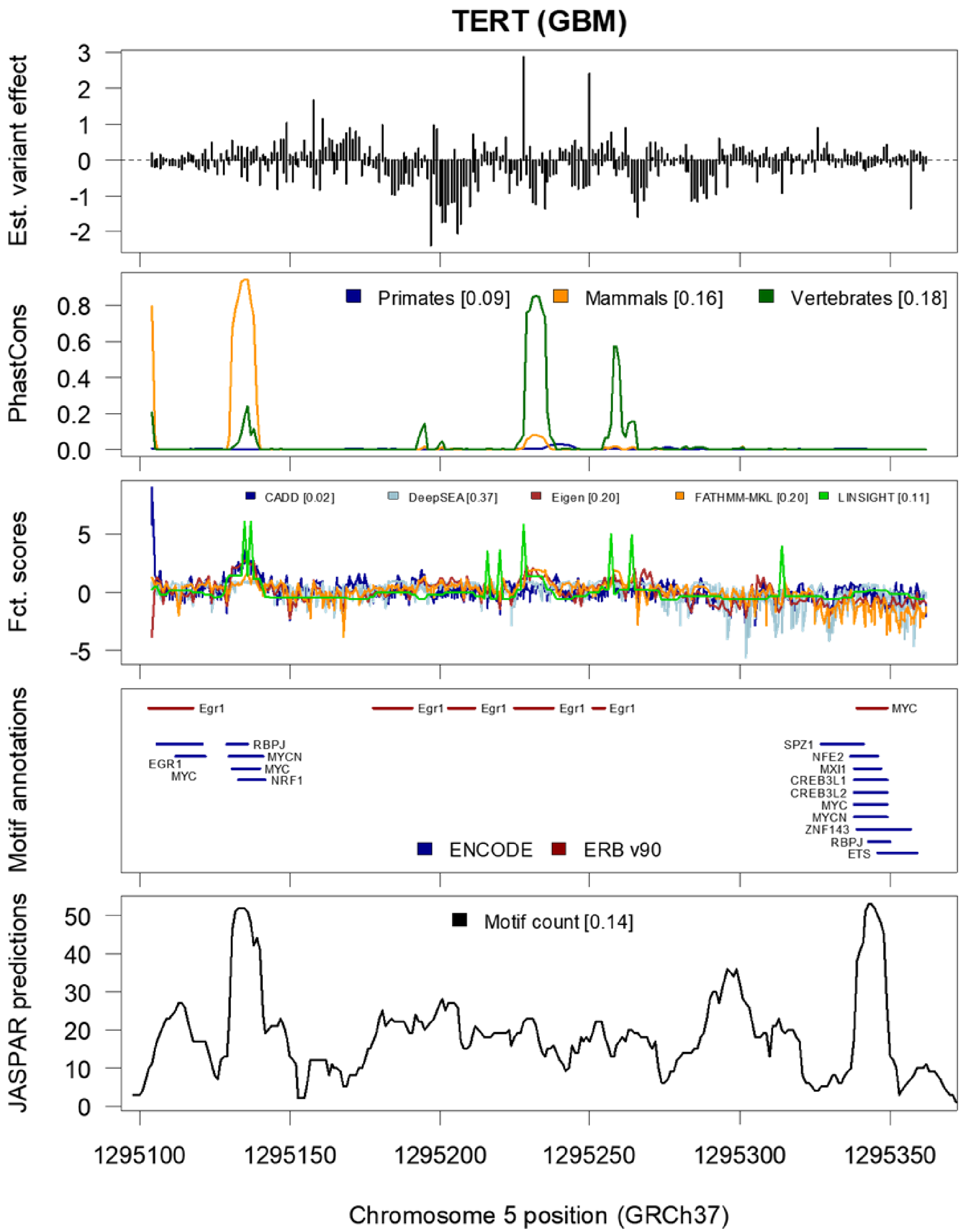
TERT (GBM)
Est. variant effect
PhastCons
Primates [0.09]
Mammals [0.16]
Vertebrates [0.18]
Fct. scores
CADD [0.02]
DeepSEA [0.37]
Eigen [0.20]
FATHMM-MKL [0.20]
LINSIGHT [0.11]
Motif annotations
Egr1
MYC
RBPJ
MYCN
EGR1
MYC
NRF1
SPZ1
NFE2
MXI1
CREB3L1
CREB3L2
MYC
MYCN
ZNF143
RBPJ
ETS
ENCODE
ERB v90
JASPAR predictions
Motif count [0.14]
1295100
1295150
1295200
1295250
1295300
1295350
Chromosome 5 position (GRCh37)

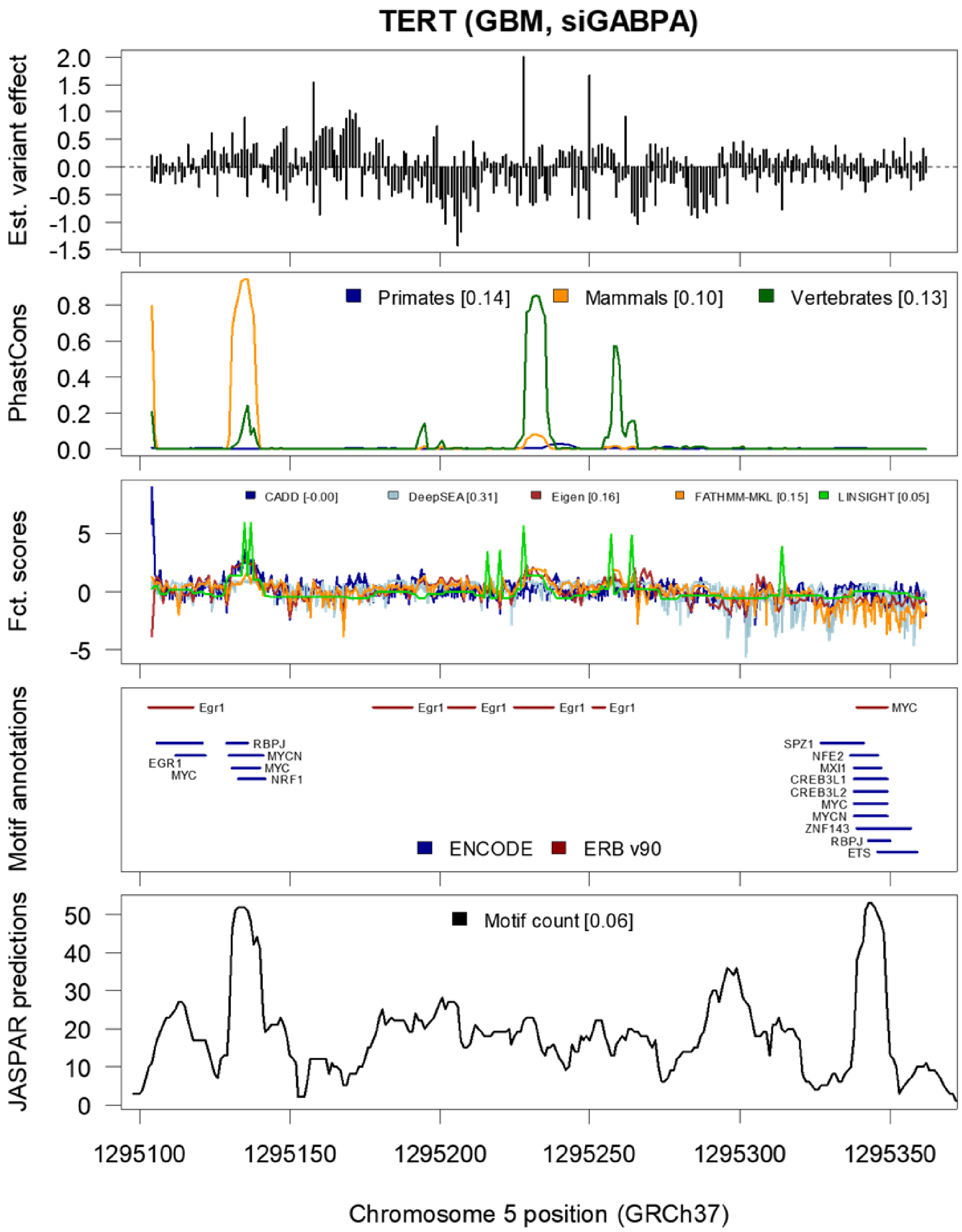
TERT (GBM, siGABPA)
Est. variant effect
Primates [0.14]
Mammals [0.10]
Vertebrates [0.13]
PhastCons
CADD [-0.00]
DeepSEA [0.31]
Eigen [0.16]
FATHMM-MKL [0.15]
LINSIGHT [0.05]
Fct. scores
Egr1
MYC
EGR1
RBPJ
MYCN
NRF1
SPZ1
NFE2
MXI1
CREB3L1
CREB3L2
ZNF143
ETS
ENCODE
ERB v90
Motif annotations
Motif count [0.06]
JASPAR predictions
1295100
1295150
1295200
1295250
1295300
1295350
Chromosome 5 position (GRCh37)

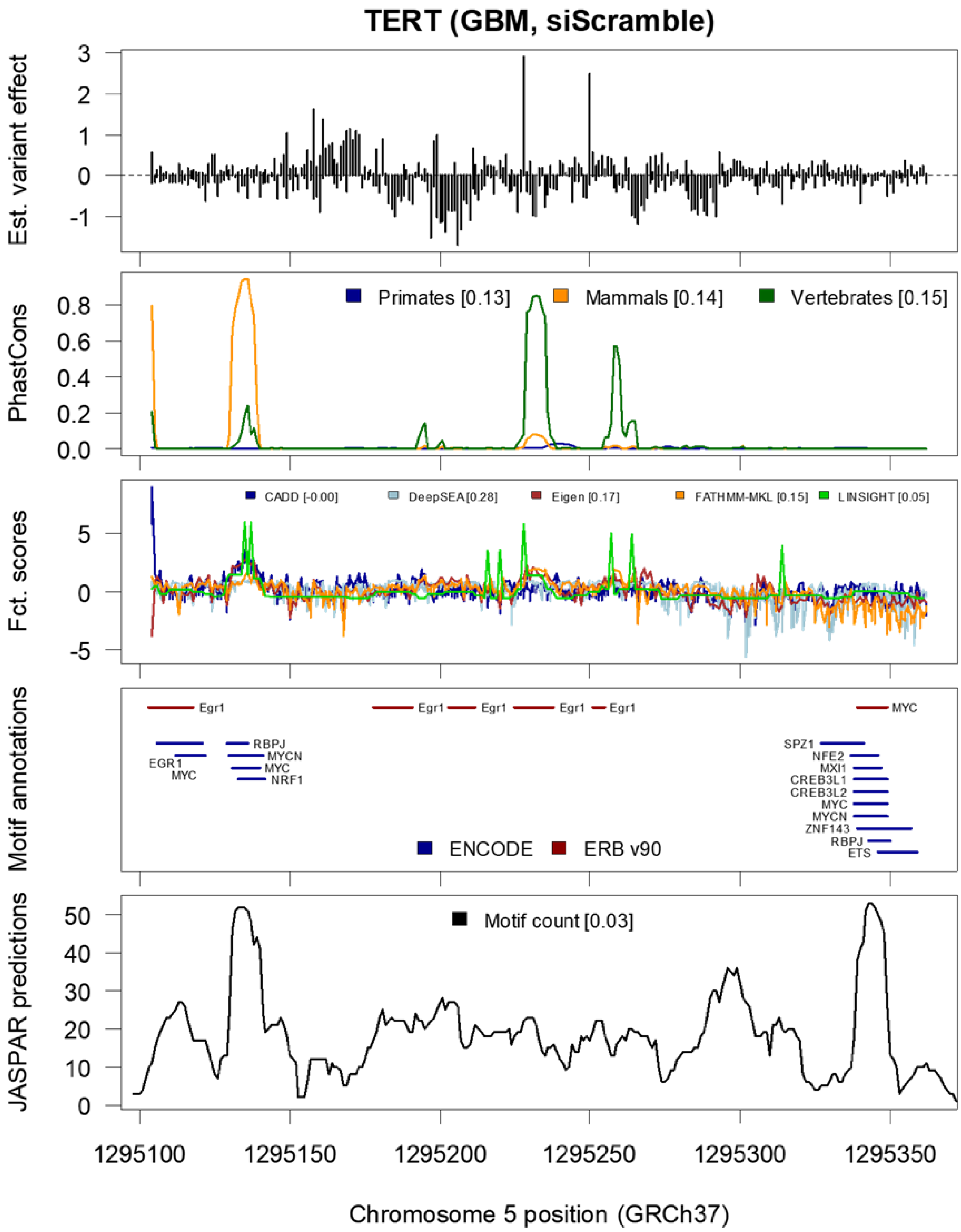

TERT (GBM, siScramble)
Est. variant effect
Primates [0.13]
Mammals [0.14]
Vertebrates [0.15]
PhastCons
CADD [-0.00]
DeepSEA [0.28]
Eigen [0.17]
FATHMM-MKL [0.15]
LINSIGHT [0.05]
Fct. scores
Egr1
MYC
RBPJ
MYCN
EGR1
NRF1
SPZ1
NFE2
MXI1
CREB3L1
CREB3L2
ZNF143
ETS
ENCODE
ERB v90
Motif annotations
Motif count [0.03]
JASPAR predictions
1295100
1295150
1295200
1295250
1295300
1295350
Chromosome 5 position (GRCh37)

**Overlay of inferred variant effects for SNVs from promoter elements with available annotation data.** Motif predictions overlayed with biochemical evidence from ChIP-seq experiments were obtained from Ensembl Regulatory Build (ERB)[33] and ENCODE[3] annotations. JASPAR motif predictions were obtained from the JASPAR 2018[6] release.

***Supplementary Figure 9***

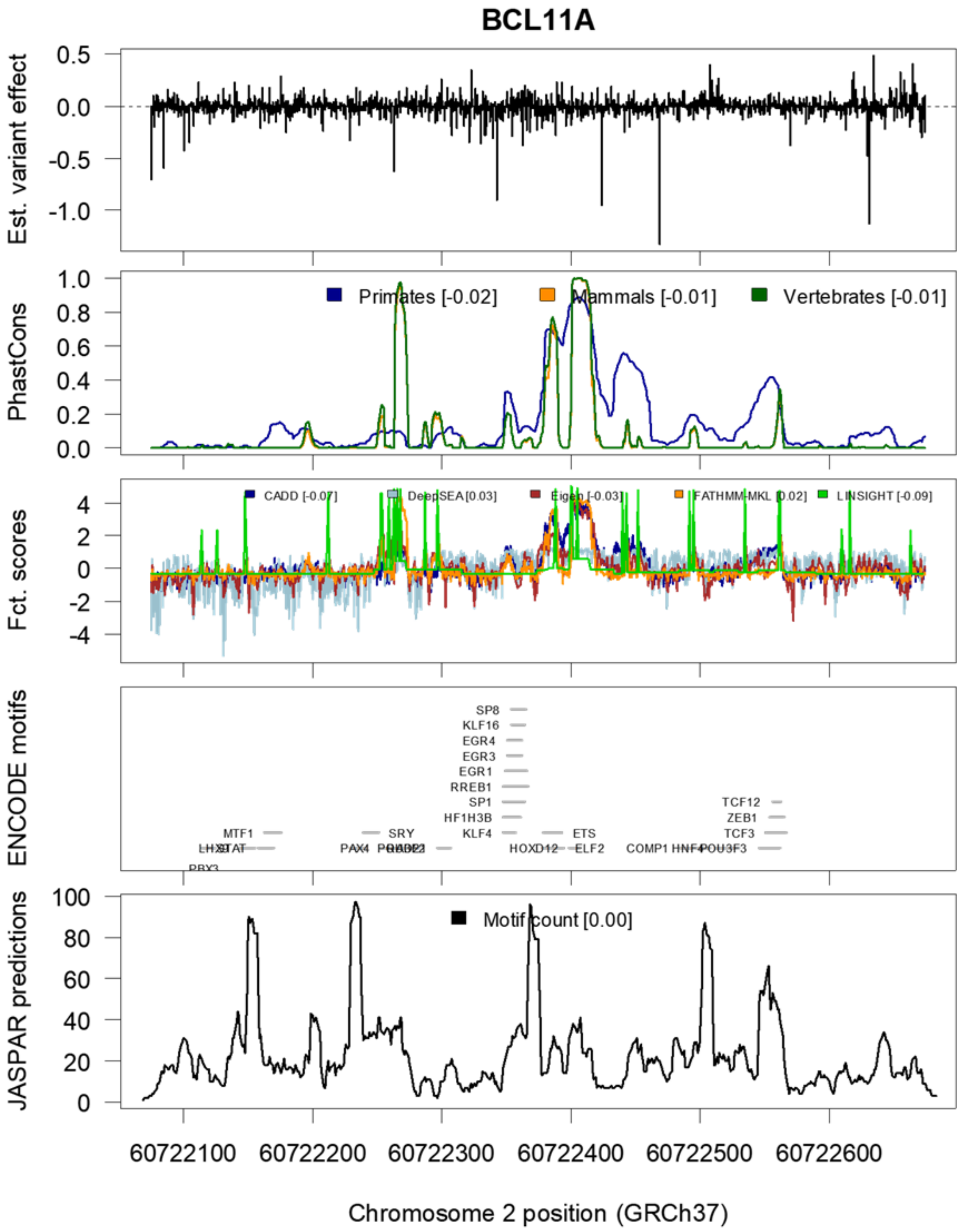

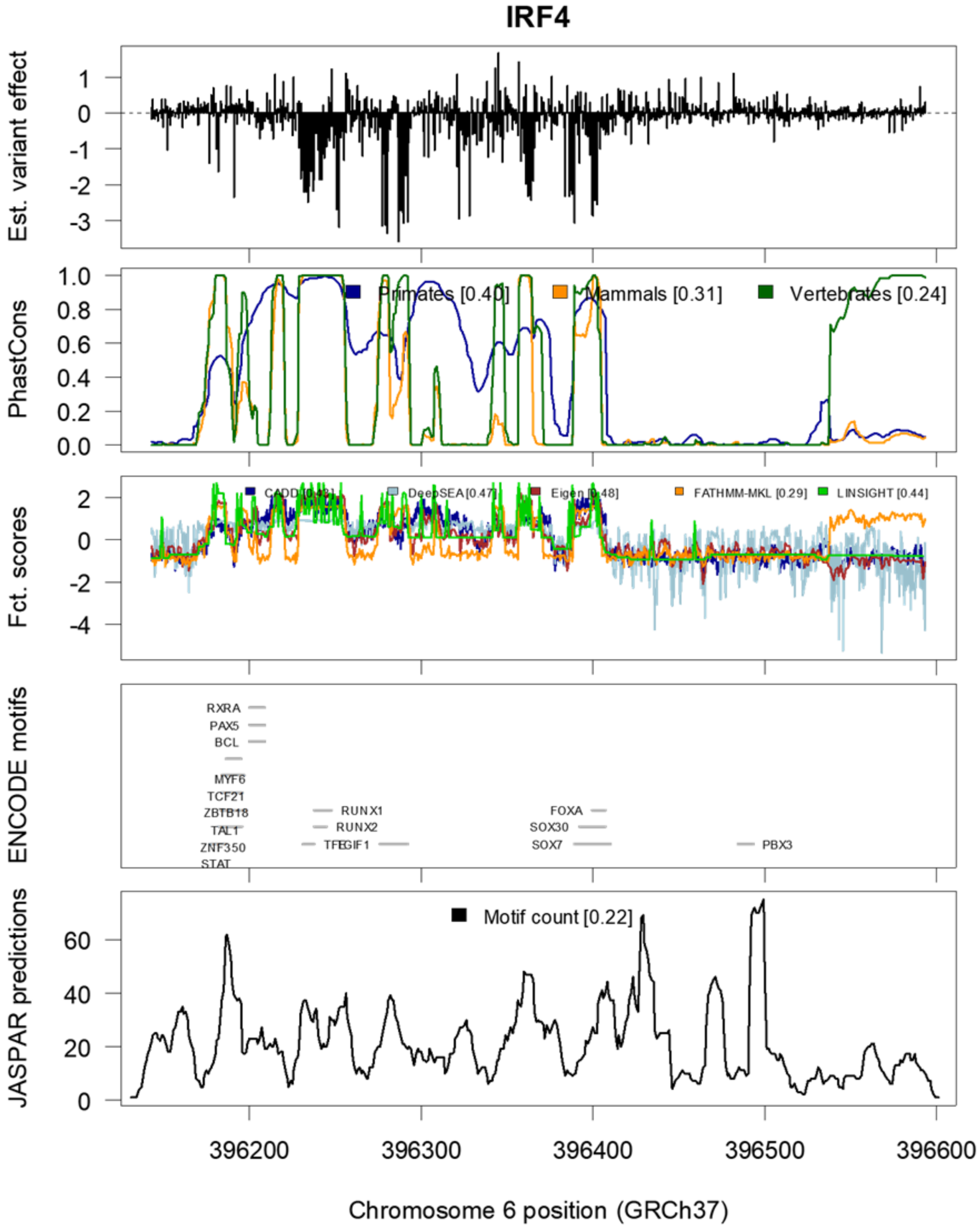

IRF4
Est. variant effect
PhastCons
Primates [0.40]
Mammals [0.31]
Vertebrates [0.24]
Fct. scores
CADD [0.43]
DeepSEA [0.47]
Eigen [0.48]
FATHMM-MKL [0.29]
LINSIGHT [0.44]
ENCODE motifs
RXRA
PAX5
BCL
MYF6
TCF21
ZBTB18
TAL1
ZNF350
STAT
RUNX1
RUNX2
TFEGIF1
FOXA
SOX30
SOX7
PBX3
JASPAR predictions
Motif count [0.22]
396200
396300
396400
396500
396600
Chromosome 6 position (GRCh37)

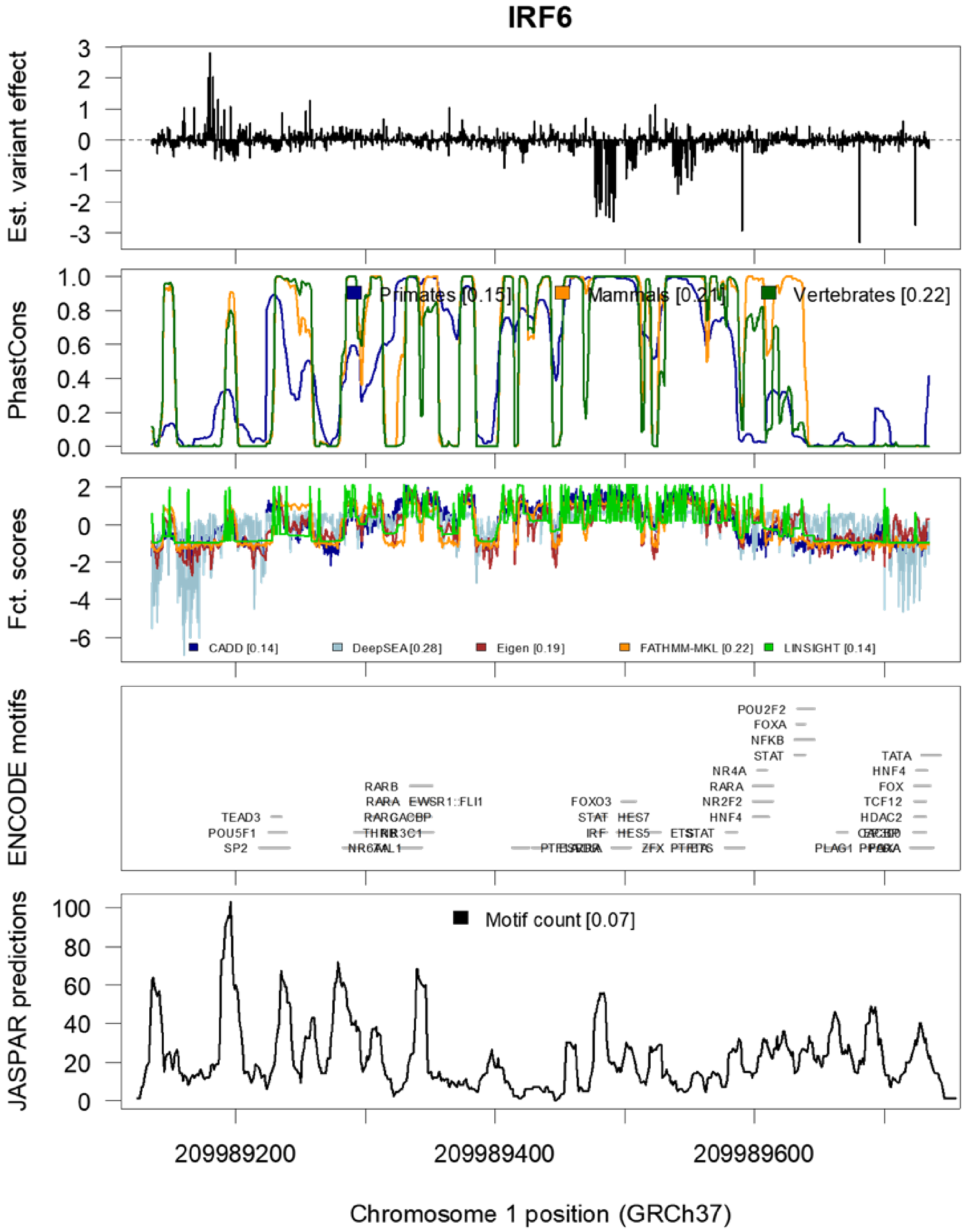
IRF6
Est. variant effect
PhastCons
Primates [0.15]
Mammals [0.21]
Vertebrates [0.22]
Fct. scores
CADD [0.14]
DeepSEA [0.28]
Eigen [0.19]
FATHMM-MKL [0.22]
LINSIGHT [0.14]
ENCODE motifs
JASPAR predictions
Motif count [0.07]
209989200
209989400
209989600
Chromosome 1 position (GRCh37)

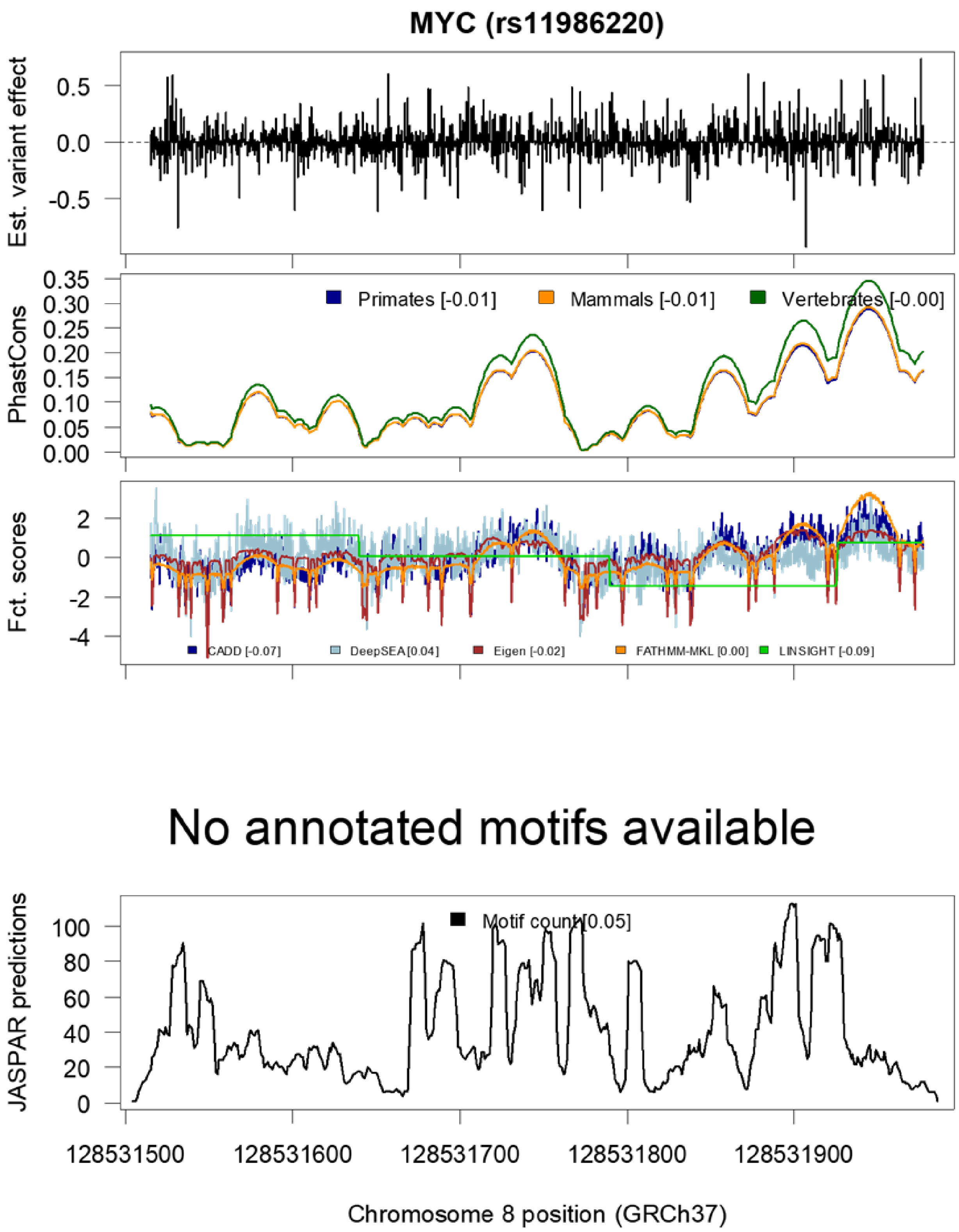
MYC (rs11986220)
Est. variant effect
0.5
0.0
-0.5
PhastCons
0.35
0.30
0.25
0.20
0.15
0.10
0.05
0.00
Primates [-0.01]
Mammals [-0.01]
Vertebrates [-0.00]
Fct. scores
2
0
-2
-4
CADD [-0.07]
DeepSEA [0.04]
Eigen [-0.02]
FATHMM-MKL [0.00]
LINSIGHT [-0.09]
No annotated motifs available
JASPAR predictions
100
80
60
40
20
0
Motif count [0.05]
128531500
128531600
128531700
128531800
128531900
Chromosome 8 position (GRCh37)

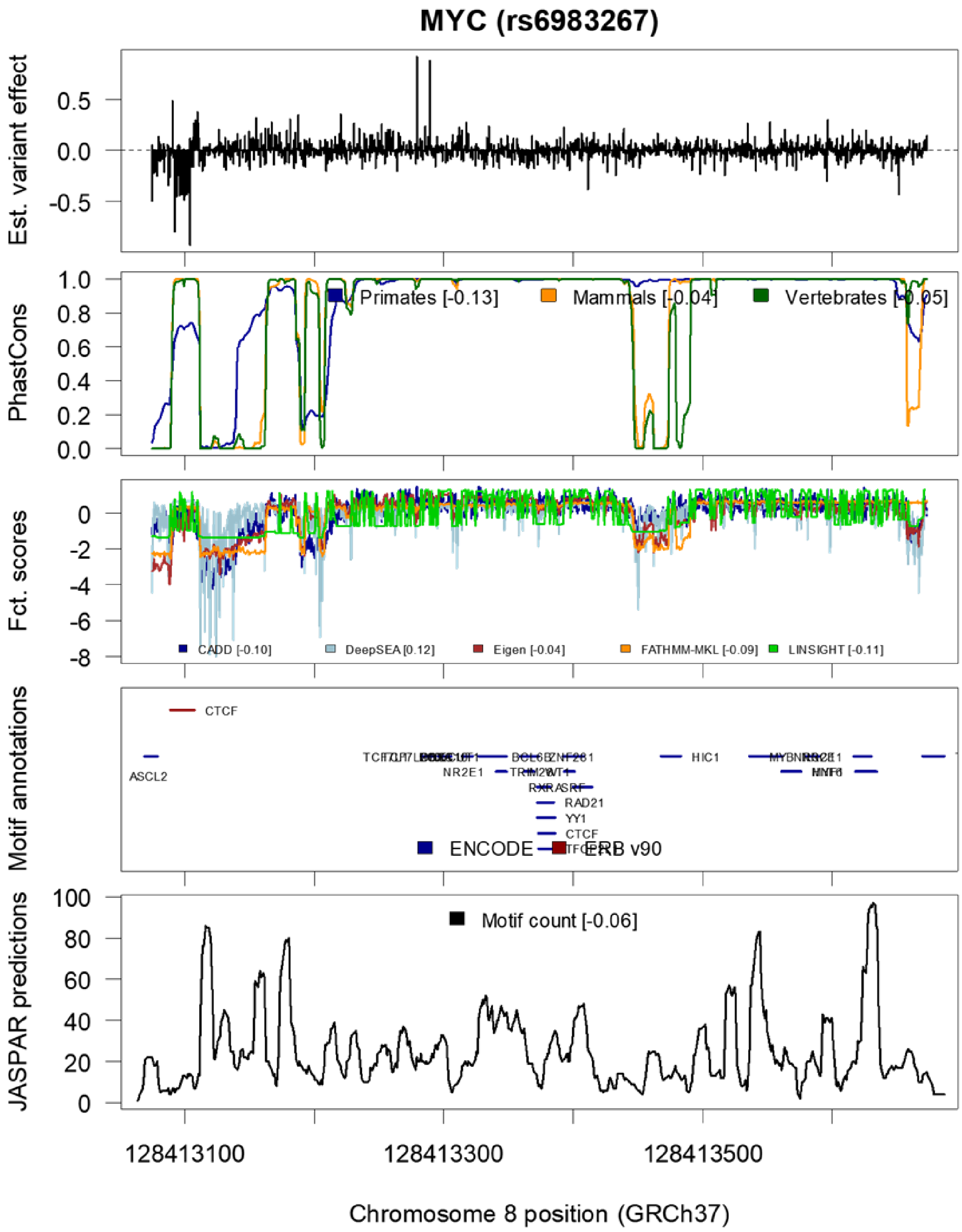
MYC (rs6983267)
Est. variant effect
0.5
0.0
-0.5
PhastCons
1.0
0.8
0.6
0.4
0.2
0.0
Primates [-0.13]
Mammals [-0.04]
Vertebrates [-0.05]
Fct. scores
0
-2
-4
-6
-8
CADD [-0.10]
DeepSEA [0.12]
Eigen [-0.04]
FATHMM-MKL [-0.09]
LINSIGHT [-0.11]
Motif annotations
CTCF
ASCL2
NR2E1
HIC1
RAD21
YY1
CTCF
ENCODE
ERB v90
JASPAR predictions
100
80
60
40
20
0
Motif count [-0.06]
128413100
128413300
128413500
Chromosome 8 position (GRCh37)

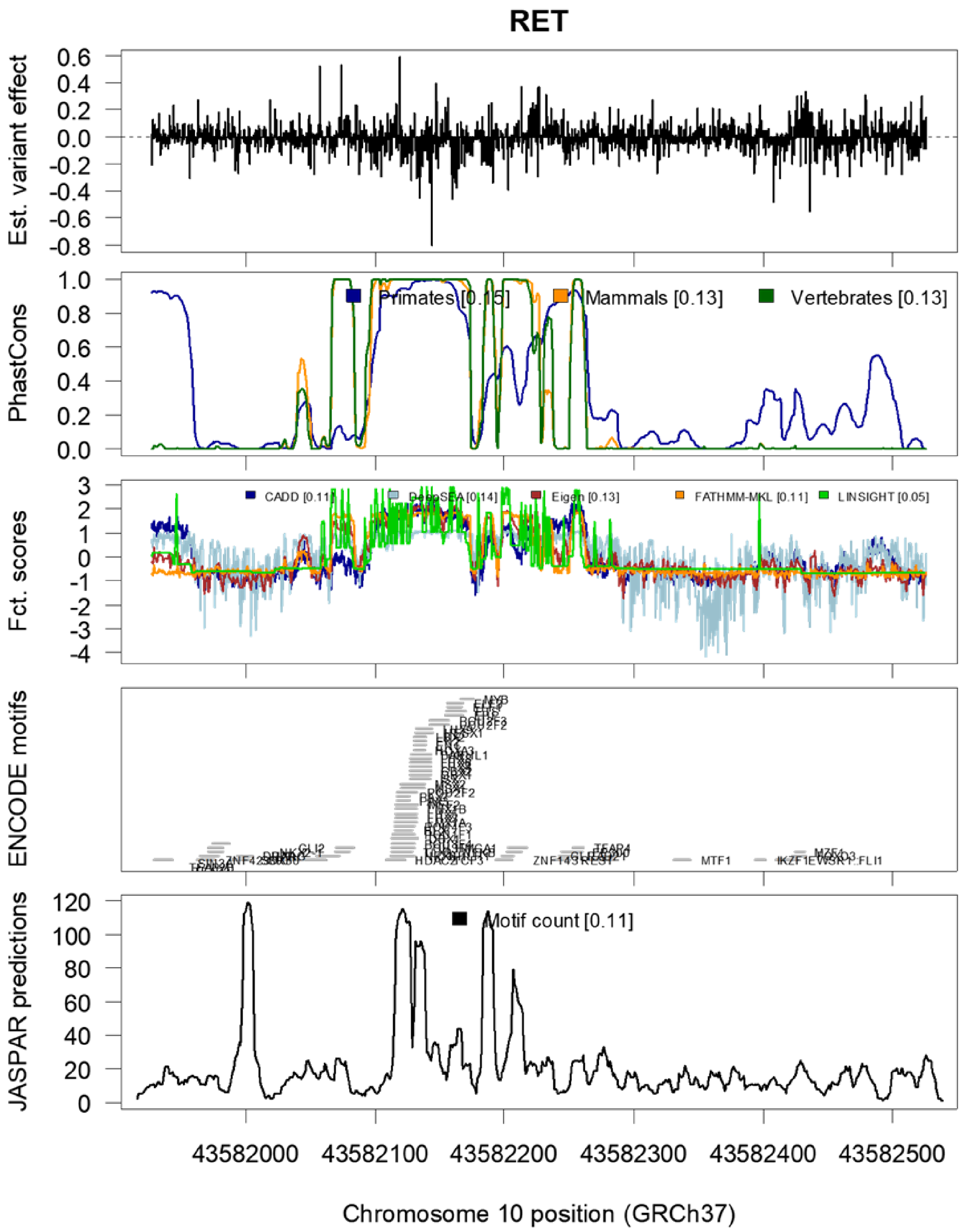

RET
Est. variant effect
PhastCons
Primates [0.15]
Mammals [0.13]
Vertebrates [0.13]
Fct. scores
CADD [0.11]
DeepSEA [0.14]
Eigen [0.13]
FATHMM-MKL [0.11]
LINSIGHT [0.05]
ENCODE motifs
JASPAR predictions
Motif count [0.11]
43582000
43582100
43582200
43582300
43582400
43582500
Chromosome 10 position (GRCh37)

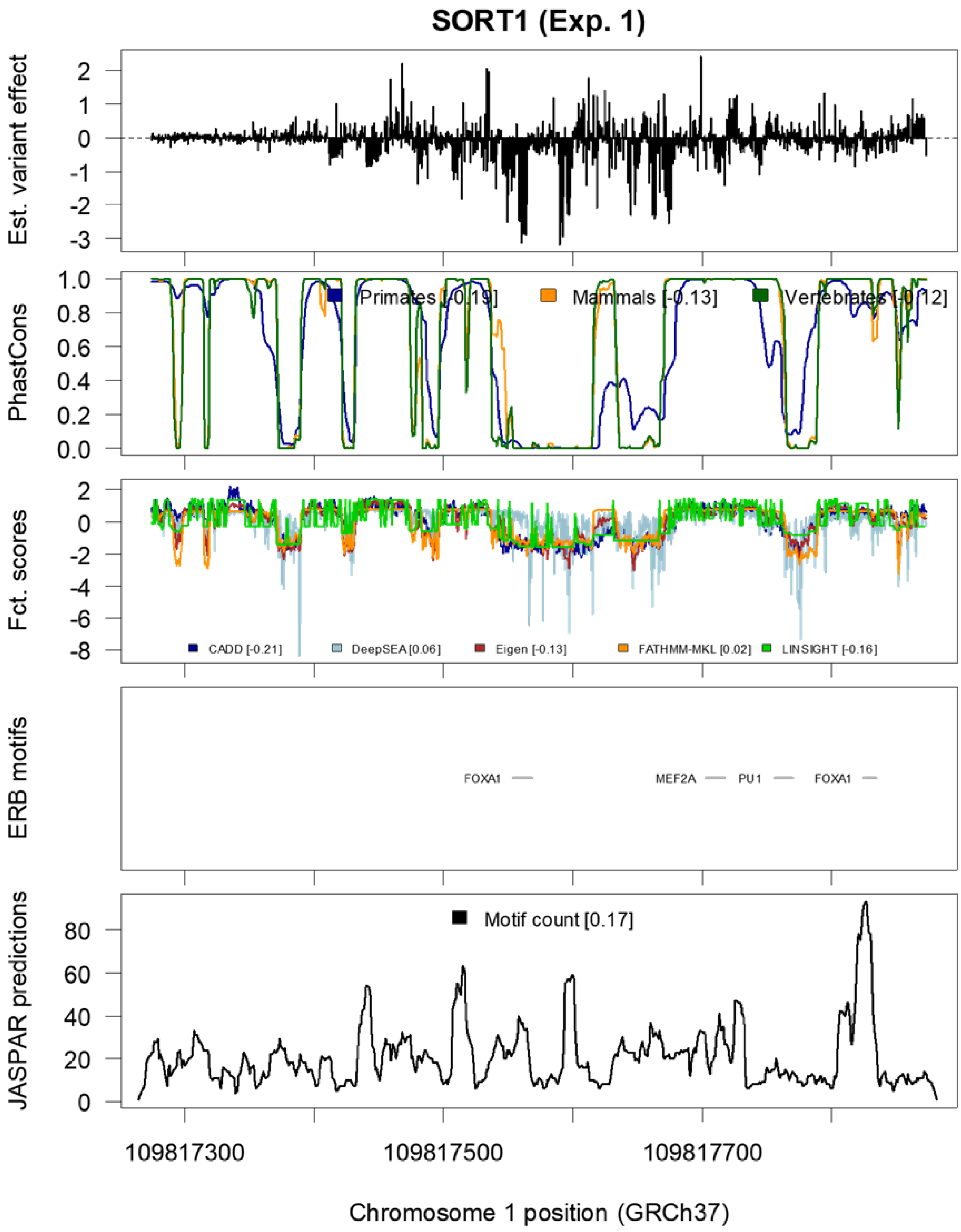
SORT1 (Exp. 1)
Est. variant effect
PhastCons
Primates [-0.19]
Mammals [-0.13]
Vertebrates [-0.12]
Fct. scores
CADD [-0.21]
DeepSEA [0.06]
Eigen [-0.13]
FATHMM-MKL [0.02]
LINSIGHT [-0.16]
ERB motifs
FOXA1
MEF2A
PU1
FOXA1
JASPAR predictions
Motif count [0.17]
109817300
109817500
109817700
Chromosome 1 position (GRCh37)

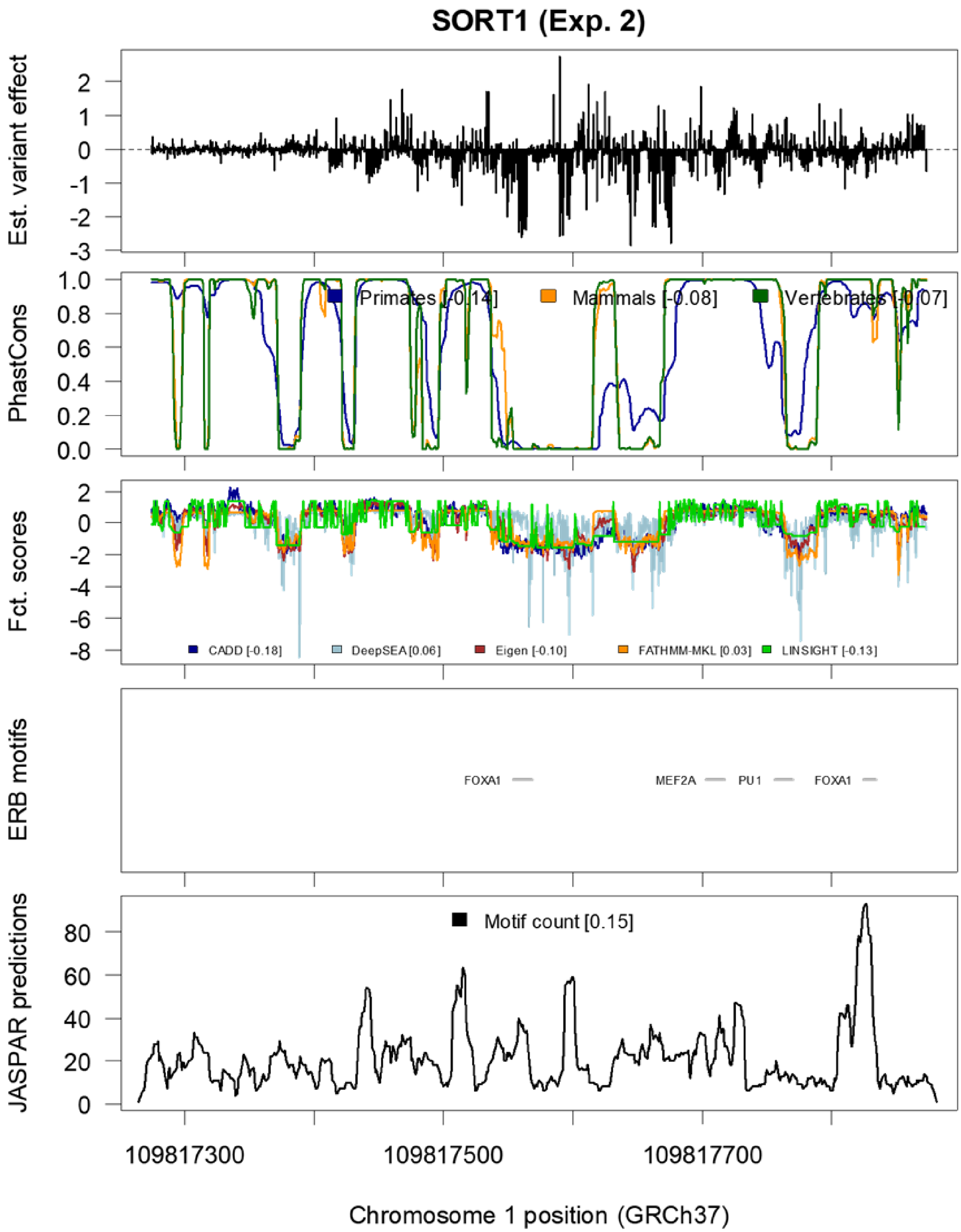

SORT1 (Exp. 2)
Est. variant effect
PhastCons
Primates [-0.14]
Mammals [-0.08]
Vertebrates [-0.07]
Fct. scores
CADD [-0.18]
DeepSEA [0.06]
Eigen [-0.10]
FATHMM-MKL [0.03]
LINSIGHT [-0.13]
ERB motifs
FOXA1
MEF2A
PU1
FOXA1
JASPAR predictions
Motif count [0.15]
109817300
109817500
109817700
Chromosome 1 position (GRCh37)

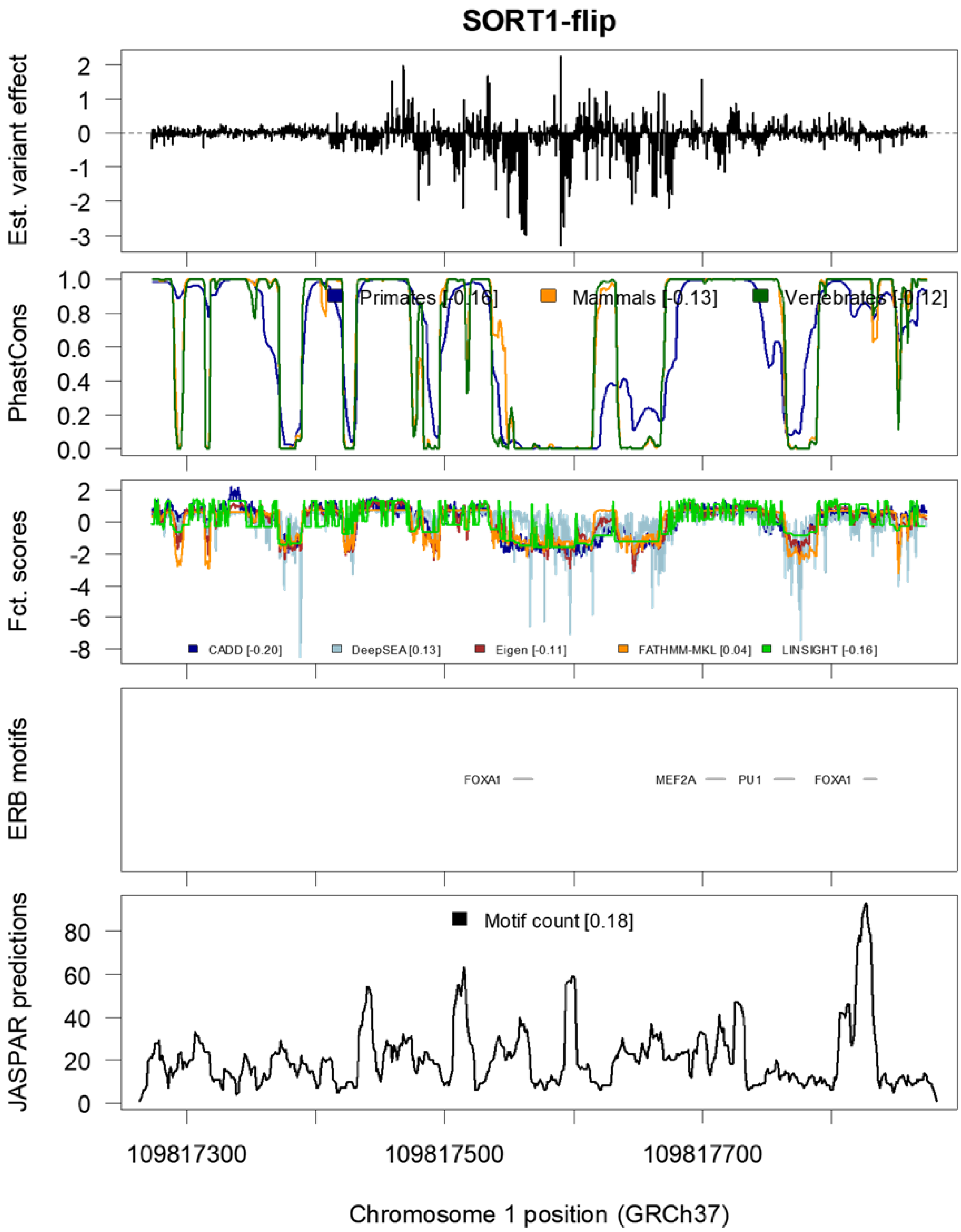
SORT1-flip
Est. variant effect
PhastCons
Primates [-0.16]
Mammals [-0.13]
Vertebrates [-0.12]
Fct. scores
CADD [-0.20]
DeepSEA [0.13]
Eigen [-0.11]
FATHMM-MKL [0.04]
LINSIGHT [-0.16]
ERB motifs
FOXA1
MEF2A
PU1
FOXA1
JASPAR predictions
Motif count [0.18]
109817300
109817500
109817700
Chromosome 1 position (GRCh37)

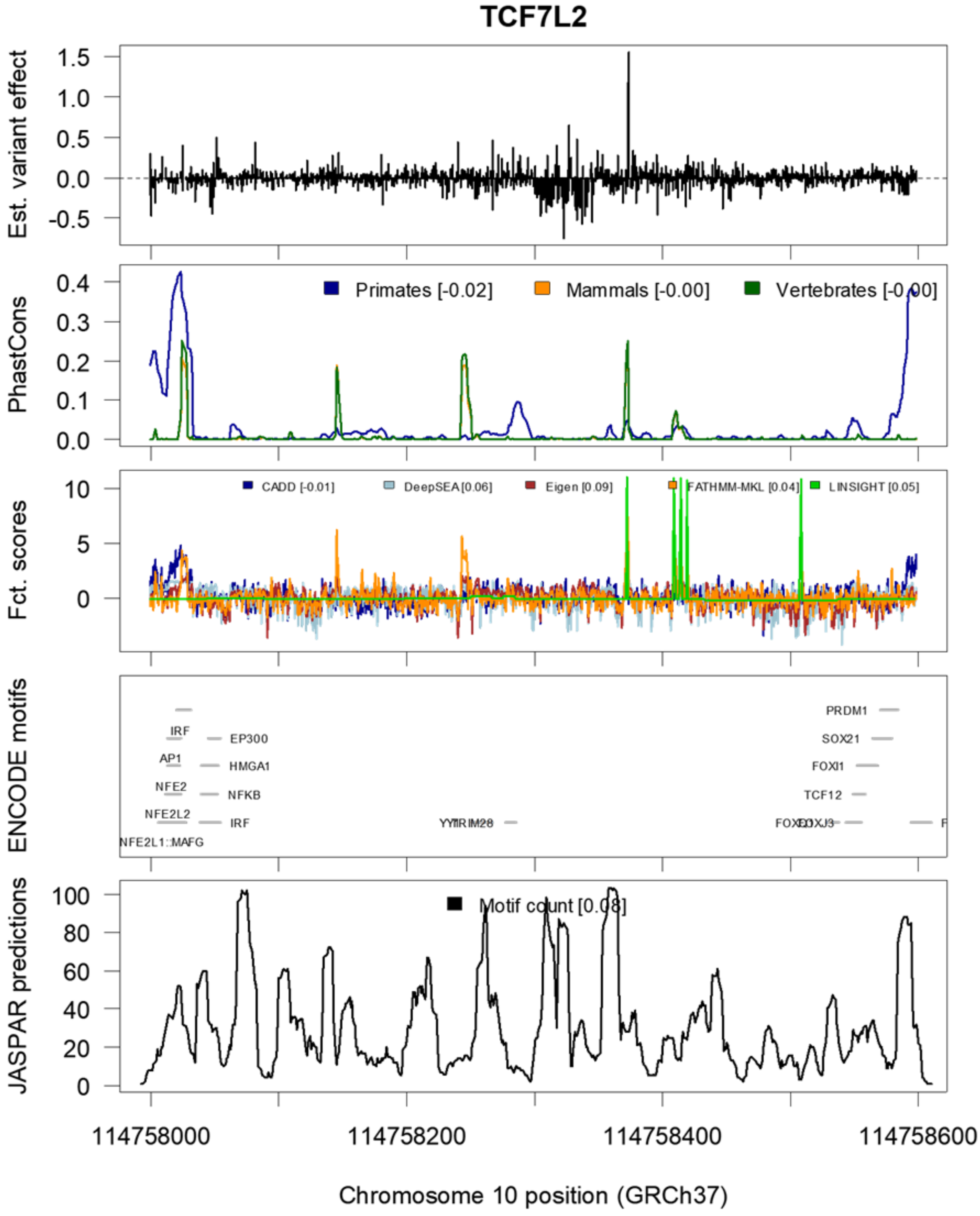
TCF7L2
Est. variant effect
PhastCons
Primates [-0.02]
Mammals [-0.00]
Vertebrates [-0.00]
Fct. scores
CADD [-0.01]
DeepSEA [0.06]
Eigen [0.09]
FATHMM-MKL [0.04]
LINSIGHT [0.05]
ENCODE motifs
IRF
AP1
NFE2
NFE2L2
NFE2L1::MAFG
EP300
HMGA1
NFKB
IRF
PRDM1
SOX21
FOXI1
TCF12
JASPAR predictions
Motif count [0.08]
114758000
114758200
114758400
114758600
Chromosome 10 position (GRCh37)

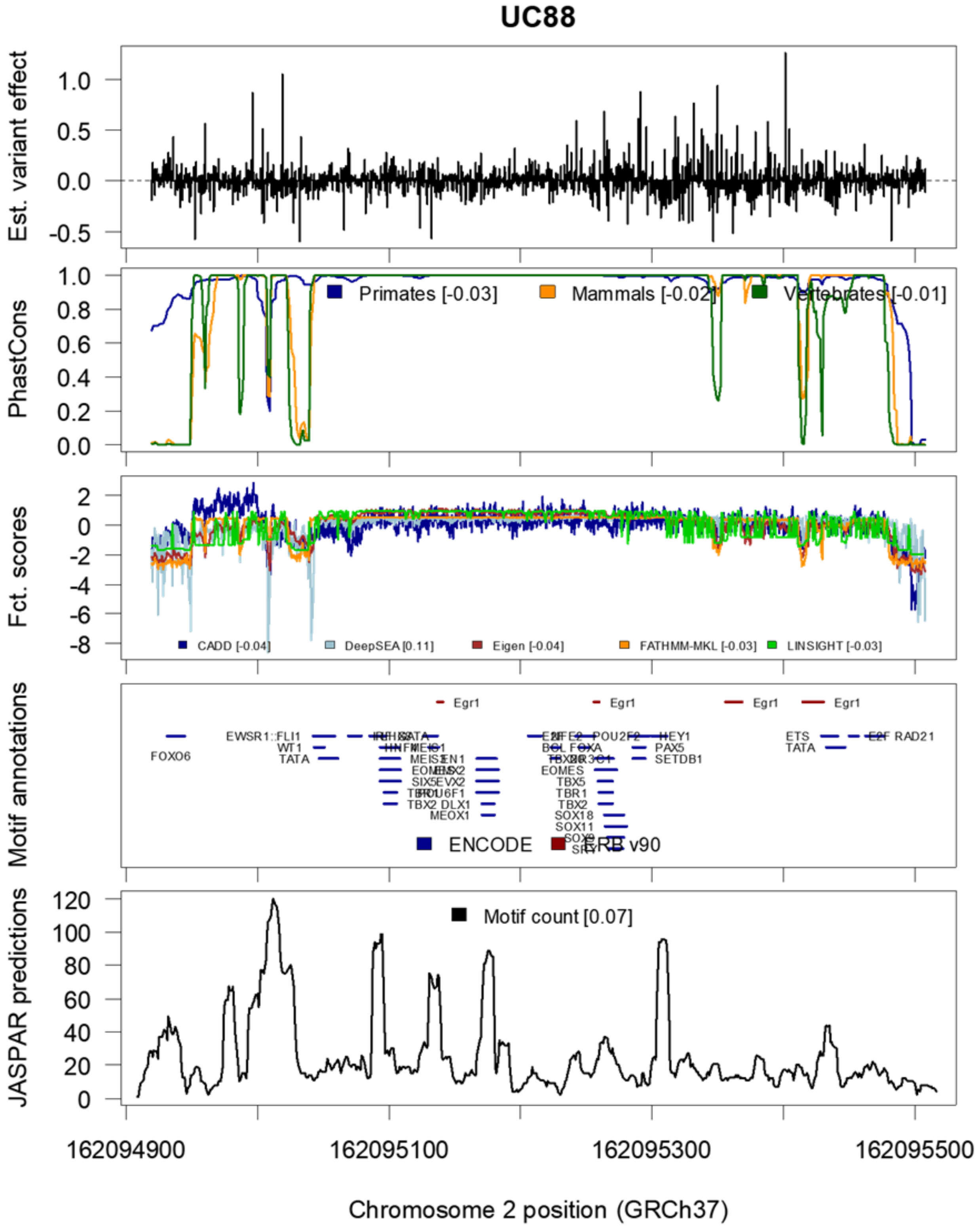
UC88
Est. variant effect
PhastCons
Primates [-0.03]
Mammals [-0.02]
Vertebrates [-0.01]
Fct. scores
CADD [-0.04]
DeepSEA [0.11]
Eigen [-0.04]
FATHMM-MKL [-0.03]
LINSIGHT [-0.03]
Motif annotations
ENCODE
ERB v90
JASPAR predictions
Motif count [0.07]
162094900
162095100
162095300
162095500
Chromosome 2 position (GRCh37)

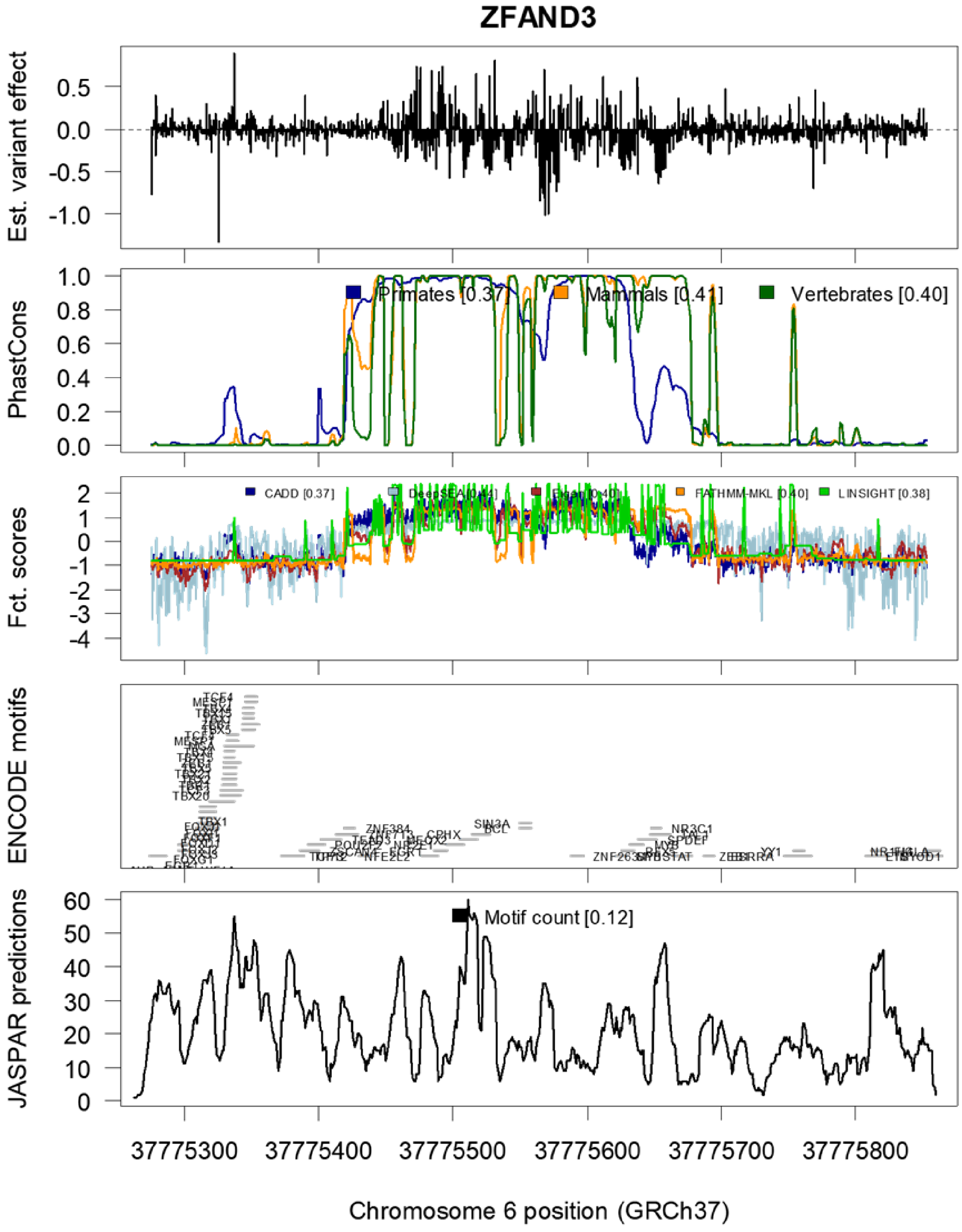
ZFAND3
Est. variant effect
PhastCons
Primates [0.37]
Mammals [0.41]
Vertebrates [0.40]
Fct. scores
CADD [0.37]
LINSIGHT [0.38]
ENCODE motifs
JASPAR predictions
Motif count [0.12]
37775300
37775400
37775500
37775600
37775700
37775800
Chromosome 6 position (GRCh37)

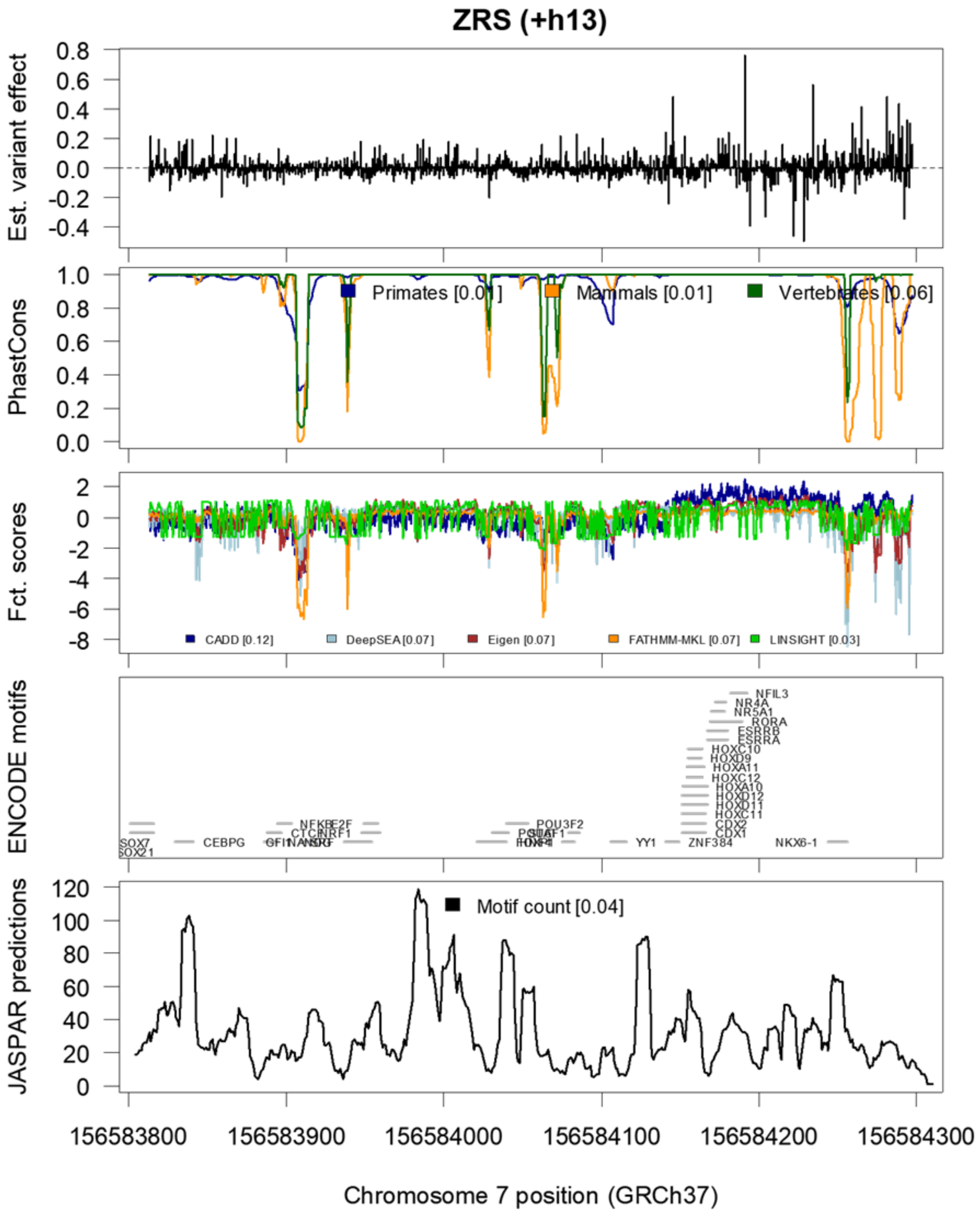
ZRS (+h13)
Est. variant effect
PhastCons
Primates [0.01]
Mammals [0.01]
Vertebrates [0.06]
Fct. scores
CADD [0.12]
DeepSEA [0.07]
Eigen [0.07]
FATHMM-MKL [0.07]
LINSIGHT [0.03]
ENCODE motifs
NFIL3
NR4A
NR5A1
RORA
ESRRB
ESRRA
HOXC10
HOXD9
HOXA11
HOXC12
HOXA10
HOXD12
HOXD11
HOXC11
CDX2
CDX1
ZNF384
NKX6-1
YY1
POU3F2
CEBPG
SOX7
SOX21
JASPAR predictions
Motif count [0.04]
156583800
156583900
156584000
156584100
156584200
156584300
Chromosome 7 position (GRCh37)

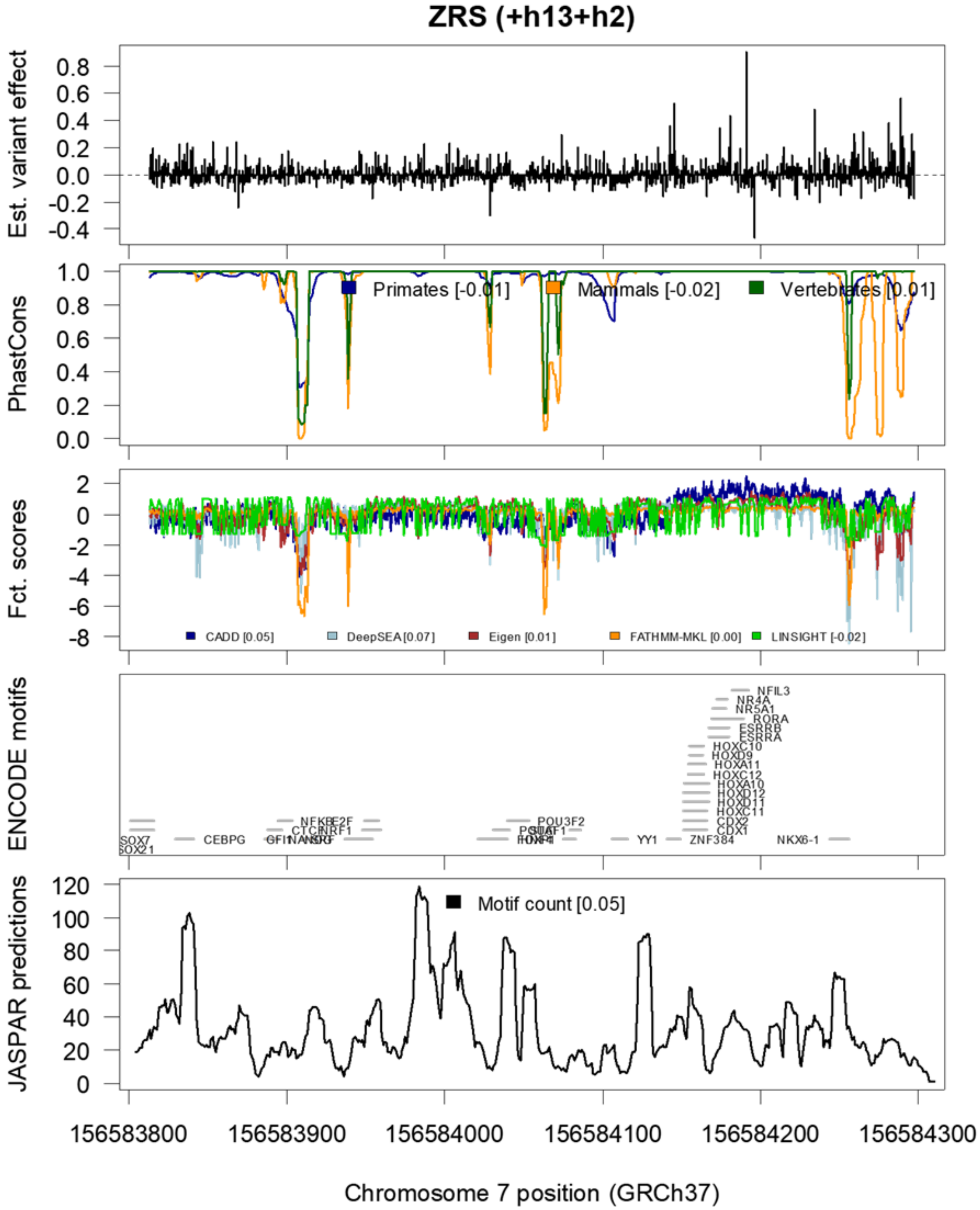

ZRS (+h13+h2)
Est. variant effect
0.8
0.6
0.4
0.2
0.0
-0.2
-0.4
PhastCons
1.0
0.8
0.6
0.4
0.2
0.0
Primates [-0.01]
Mammals [-0.02]
Vertebrates [0.01]
Fct. scores
2
0
-2
-4
-6
-8
CADD [0.05]
DeepSEA [0.07]
Eigen [0.01]
FATHMM-MKL [0.00]
LINSIGHT [-0.02]
ENCODE motifs
NFIL3
NR4A
NR5A1
RORA
ESRRB
ESRRA
HOXC10
HOXD9
HOXA11
HOXC12
HOXA10
HOXD12
HOXD11
HOXC11
CDX2
CDX1
ZNF384
NKX6-1
YY1
POU3F2
CEBPG
SOX7
SOX21
JASPAR predictions
120
100
80
60
40
20
0
Motif count [0.05]
156583800
156583900
156584000
156584100
156584200
156584300
Chromosome 7 position (GRCh37)

**Overlay of inferred variant effects for SNVs with available annotation data for all enhancer elements.** Motif predictions overlayed with biochemical evidence from ChIP-seq experiments were obtained from Ensembl Regulatory Build (ERB)[33] and ENCODE[3] annotations. JASPAR motif predictions were obtained from the JASPAR 2018[6] release.

### Supplementary Tables

#### *Supplementary Table 1*

**Tested promoter sequences.** Note that the construct length of MSMB is 2 bp longer than the described reference genome coordinate range, due to an insertion polymorphism present in the construct. Vector sequences for pGL4.11b and pGL4.11c are available for download from our OSF project 75B2M.

| Name | Genomic coordinates (GRCh37/GRCh38) | Transcript | Associated Phenotype | Luciferase vector | MPRA vector | Cell line | Transf. time (hr) | Fold Ch. (Wild type) | Fold Ch. (MPRA) | Construct size (bp) |
|---|---|---|---|---|---|---|---|---|---|---|
| *F9* | X:138,612,622-138,612,924<br>X:139,530,463-139,530,765 | NM_000133.3 | Hemophilia B | pGL4.11b | pGL4.11c | HepG2 | 24 | 2.6 | 2.1 | 303 |
| *FOXE1* | 9:100,615,537-100,616,136<br>9:97,853,255-97,853,854 | NM_004473.3 | Thyroid cancer | pGL4.11b | pGL4.11c | HeLa | 24 | 4.2 | 2.5 | 600 |
| *GP1BB* | 22:19,710,789-19,711,173<br>22:19,723,266-19,723,650 | NM_000407.4 | Bernard-Soulier Syndrome | pGL4.11b | pGL4.11c | HEL 92.1.7 | 24 | 22.1 | 12.3 | 385 |
| *HBB* | 11:5,248,252-5,248,438<br>11:5,227,022-5,227,208 | NM_000518.4 | Thalassemia | pGL4.11b | pGL4.11c | HEL 92.1.7 | 24 | 14.3 | 8.4 | 187 |
| *HBG1* | 11:5,271,035-5,271,308<br>11:5,249,805-5,250,078 | NM_000559.2 | Hereditary persistence of fetal hemoglobin | pGL4.11b | pGL4.11c | HEL 92.1.7 | 24 | 118.1 | 41.8 | 274 |
| *HNF4A (P2)* | 20:42,984,160-42,984,444<br>20:44,355,520-44,355,804 | NM_175914.4 | Maturity-onset diabetes of the young (MODY) | pGL4.11b | pGL4.11c | HEK293T | 24 | 2.8 | 1.3 | 285 |
| *LDLR* | 19:11,199,907-11,200,224<br>19:11,089,231-11,089,548 | NM_000527.4 | Familial hypercholesterolemia | pGL4.11b | pGL4.11b | HepG2 | 24 | 110.7 | 76.6 | 318 |
| *MSMB* | 10:51,548,988-51,549,578<br>10:46,046,244-46,046,834 | NM_002443.3 | Prostate cancer | pGL4.11b | pGL4.11c | HEK293T | 24 | 8.4 | 3.4 | 593 |
| *PKLR* | 1:155,271,186-155,271,655<br>1:155,301,395-155,301,864 | NM_000298.5 | Pyruvate kinase deficiency | pGL4.11b | pGL4.11c | K562 | 48 | 29.4 | 9.6 | 470 |
| *TERT* | 5:1,295,104-1,295,362<br>5:1,294,989-1,295,247 | NM_198253.2 | Various types of cancer | pGL4.11b | pGL4.11b | HEK293T, SF7996 | 24 | 231.8,5.2 | 148.2, 2.7 | 259 |

***Supplementary Table 2***

**Tested enhancer sequences.** Note that the construct length of MYC (rs11986220) is 1 bp longer than the described reference genome coordinate range, due to an insertion polymorphism present in the construct. Human ZRS (hZRS, a limb-specific enhancer of the Sonic hedgehog gene) was co-transfected with HOXD13 and HOXD13+HAND2. UC88 is an ultraconserved element, which has not been previously associated with a phenotype. Vector sequences for pGL3c, pGL4.23c/d, and pGL4Zc are available for download from our OSF project 75B2M. The TATA-pGL4m(EV087) vector was previously described by Galli A et al. in PLoS Genetics 2010.

| Name | Genomic coordinates (GRCh37/GRCh38) | Associated Phenotype | Luciferase vector | MPRA vector | Cell line | Transf. time (hr) | Fold Ch. (Wild type) | Fold Ch. (MPRA) | Construct (bp) |
|---|---|---|---|---|---|---|---|---|---|
| *BCL11A +58* | 2:60,722,075-60,722,674<br>2:60,494,940-60,495,539 | Sickle cell disease | pGL4.23 | pGL4.23d | HEL 92.1.7 | 24 | 2.5 | 1.7 | 600 |
| *IRF4* | 6:396,143-396,593<br>6:396,143-396,593 | Human pigmentation | pGL4.23 | pGL4.23d | SK-MEL-28 | 24 | 44.5 | 16.3 | 451 |
| *IRF6* | 1:209,989,135-209,989,735<br>1:209,815,790-209,816,390 | Cleft lip | pGL4.23 | pGL4.23c | HaCaT | 24 | 17.0 | 16.7 | 600 |
| *MYC (rs 6983267)* | 8:128,413,074-128,413,673<br>8:127,400,829-127,401,428 | Various types of cancer | pGL4.23 | pGL4.23c | HEK293T | 32, 20nM LiCl added after 24hr | 2.8 | 0.7 | 600 |
| *MYC (rs 11986220)* | 8:128,531,515-128,531,977<br>8:127,519,270-127,519,732 | Various types of cancer | pGL4.23 | pGL4.23d | LNCaP + 100nM DHT | 24 | 5.5 | 3.2 | 464 |
| *RET* | 10:43,581,927-43,582,526<br>10:43,086,479-43,087,078 | Hirschsprung | pGL3 | pGL3c | Neuro-2a | 24 | 2.0 | 0.9 | 600 |
| *SORT1* | 1:109,817,274-109,817,873<br>1:109,274,652-109,275,251 | Plasma low-density lipoprotein cholesterol & myocardial infarction | pGL4.23 | pGL4.23c | HepG2 | 24 | 235.3 | 202.2 | 600 |
| *TCF7L2* | 10:114,757,999-114,758,598<br>10:112,998,240-112,998,839 | Type 2 diabetes | pGL4.23 | pGL4.23d | MIN6 | 24 | 9.0 | 2.7 | 600 |
| *UC88* | 2:162,094,919-162,095,508<br>2:161,238,408-161,238,997 | - | pGL4.23 | pGL4.23c | Neuro-2a | 24 | 9.3 | 5.4 | 590 |
| *ZFAND3* | 6:37,775,275-37,775,853<br>6:37,807,499-37,808,077 | Type 2 diabetes | pGL4.23 | pGL4.23c | MIN6 | 24 | 14.3 | 7.3 | 579 |
| *ZRS* | 7:156,583,813-156,584,297<br>7:156,791,119-156,791,603 | Limb malformations | TATA-pGL4m (EV087) | pGL4Zc | NIH/3T3 (with HOXD13/ HOXD13+HAND2) | 24 | 4.2/ 2.0 | 3.7/ 2.6 | 485 |

*Supplementary Table 3*

**Saturation mutagenesis tag-variant assignments.** Twenty four tag assignments were obtained by saturation mutagenesis. A minimum read coverage of three was required for reads aligning across the construct length before a tag was considered in analysis.

| Assignment | Construct length | Associated tags | Fraction WT inserts | Variants called per insert | Mutation rate | SNV cov. | 1bp-del cov | Ave #tags per SNV | Ave #tags per 1bp-del |
|---|---|---|---|---|---|---|---|---|---|
| BCL11A | 600 | 263,000 | 0.03% | 6.00 | 1.00% | 100.00% | 61.00% | 809.1 | 40.9 |
| F9 | 303 | 134,665 | 43.22% | 0.96 | 0.32% | 99.56% | 54.13% | 132.7 | 20.7 |
| FOXE1 | 600 | 148,459 | 19.61% | 2.00 | 0.33% | 100.00% | 56.50% | 154.9 | 21.2 |
| GP1BB | 385 | 485,856 | 31.82% | 1.25 | 0.33% | 99.91% | 66.23% | 488.4 | 89.8 |
| HBB | 187 | 2,010,638 | 55.62% | 0.62 | 0.33% | 100.00% | 71.12% | 2064.4 | 414.5 |
| HBG1 | 274 | 150,499 | 43.36% | 0.92 | 0.34% | 100.00% | 55.11% | 155.7 | 31.4 |
| HNF4A | 285 | 555,705 | 40.47% | 0.98 | 0.35% | 100.00% | 63.16% | 595.0 | 105.5 |
| IRF4 | 451 | 342,770 | 25.43% | 1.66 | 0.37% | 100.00% | 63.64% | 396.8 | 57.7 |
| IRF6 | 601 | 33,174 | 14.03% | 2.33 | 0.39% | 99.45% | 31.45% | 41.6 | 2.9 |
| LDLR.2 | 318 | 159,867 | 39.27% | 1.04 | 0.33% | 100.00% | 56.92% | 154.5 | 45.6 |
| LDLR | 318 | 96,145 | 17.78% | 1.99 | 0.62% | 100.00% | 58.80% | 185.3 | 35.6 |
| MSMB | 593 | 738,972 | 15.72% | 2.07 | 0.35% | 100.00% | 67.12% | 817.3 | 109.3 |
| MYC (rs11986220) | 464 | 541,564 | 2.45% | 2.52 | 0.54% | 100.00% | 55.39% | 884.1 | 73.7 |
| MYC (rs6983267) | 600 | 70,234 | 16.34% | 2.05 | 0.34% | 99.67% | 43.33% | 75.4 | 8.8 |
| PKLR | 470 | 119,675 | 22.88% | 1.75 | 0.37% | 99.93% | 45.74% | 125.5 | 12.7 |
| RET | 600 | 257,630 | 29.19% | 1.51 | 0.25% | 99.94% | 55.50% | 199.6 | 41.5 |
| SORT1.2 | 600 | 132,222 | 15.78% | 2.07 | 0.34% | 99.94% | 49.83% | 139.5 | 27.3 |
| SORT1-flip | 600 | 192,602 | 21.12% | 1.73 | 0.29% | 99.61% | 50.83% | 169.8 | 31.7 |
| SORT1 | 600 | 45,509 | 12.78% | 2.29 | 0.38% | 99.72% | 41.67% | 53.9 | 9.0 |
| TCF7L2 | 600 | 120,996 | 19.03% | 1.85 | 0.31% | 99.06% | 42.33% | 119.8 | 9.9 |
| TERT | 259 | 276,048 | 19.22% | 1.76 | 0.68% | 100.00% | 57.53% | 549.3 | 103.9 |
| UC88 | 590 | 87,948 | 11.16% | 2.44 | 0.41% | 99.77% | 52.71% | 111.1 | 17.2 |
| ZFAND3 | 579 | 183,352 | 6.52% | 3.00 | 0.52% | 99.94% | 63.21% | 297.5 | 38.4 |
| ZRS | 485 | 850,422 | 16.94% | 2.07 | 0.43% | 100.00% | 66.19% | 1130.2 | 192.9 |

*Supplementary Table 4*

**Obtained RNA and DNA tag counts across 29 experiments with each 3 transfection replicates.** MYC (rs11986220) was abbreviated to MYCs1. MYC (rs6983267) was abbreviated to MYCs2. TERT-GBM-SiGABPA and TERT-GBM-SiScramble were abbreviated to TERT-GAa and TERT-GSc, respectively.

| Exp. | Tags | DNA | RNA | Assign. tags | [%] | DNA assigned | [%] | RNA assigned | [%] | WT tags | [%] | Variants | Var/ tag |
|---|---|---|---|---|---|---|---|---|---|---|---|---|---|
| BCL11A-1 | 298,966 | 5,639,266 | 12,078,937 | 219,172 | 73.3 | 4,782,063 | 84.8 | 10,265,384 | 85.0 | 63 | 0.0 | 1,295,570 | 5.6 |
| BCL11A-2 | 285,273 | 5,482,187 | 7,401,487 | 218,289 | 76.5 | 4,660,218 | 85.0 | 6,302,212 | 85.1 | 64 | 0.0 | 1,290,502 | 5.9 |
| BCL11A-3 | 298,554 | 4,098,878 | 18,055,670 | 218,565 | 73.2 | 3,466,623 | 84.6 | 15,372,758 | 85.1 | 65 | 0.0 | 1,291,930 | 5.9 |
| F9-1 | 342,802 | 4,373,175 | 2,377,127 | 110,714 | 32.3 | 2,941,983 | 67.3 | 1,532,333 | 64.5 | 48,303 | 43.6 | 104,215 | 0.9 |
| F9-2 | 393,783 | 5,913,265 | 3,343,343 | 114,478 | 29.1 | 3,895,338 | 65.9 | 2,142,187 | 64.1 | 50,414 | 44.0 | 106,735 | 0.9 |
| F9-3 | 399,823 | 5,288,624 | 3,885,413 | 114,669 | 28.7 | 3,467,454 | 65.6 | 2,489,575 | 64.1 | 50,479 | 44.0 | 106,929 | 0.9 |
| FOXE1-1 | 371,106 | 5,844,884 | 12,244,334 | 104,919 | 28.3 | 5,071,439 | 86.8 | 10,307,674 | 84.2 | 21,761 | 20.7 | 202,298 | 1.9 |
| FOXE1-2 | 403,744 | 6,652,522 | 14,494,525 | 107,339 | 26.6 | 5,746,967 | 86.4 | 12,188,942 | 84.1 | 22,141 | 20.6 | 207,434 | 1.9 |
| FOXE1-3 | 343,043 | 5,769,389 | 9,841,232 | 101,484 | 29.6 | 5,048,490 | 87.5 | 8,271,983 | 84.1 | 21,082 | 20.8 | 195,486 | 1.9 |
| GP1BB-1 | 486,707 | 6,643,587 | 43,000,079 | 355,564 | 73.1 | 6,072,911 | 91.4 | 39,803,462 | 92.6 | 119,228 | 33.5 | 424,334 | 1.2 |
| GP1BB-2 | 578,012 | 6,367,523 | 83,229,576 | 380,180 | 65.8 | 5,752,248 | 90.3 | 77,001,869 | 92.5 | 126,862 | 33.4 | 455,869 | 1.2 |
| GP1BB-3 | 523,517 | 6,606,471 | 54,650,491 | 366,222 | 70.0 | 6,013,969 | 91.0 | 50,533,028 | 92.5 | 122,381 | 33.4 | 438,316 | 1.2 |
| HBB-1 | 695,586 | 2,525,201 | 6,773,900 | 472,421 | 67.9 | 1,900,539 | 75.3 | 4,963,191 | 73.3 | 262,233 | 55.5 | 294,795 | 0.6 |
| HBB-2 | 791,164 | 3,308,597 | 9,589,115 | 528,865 | 66.8 | 2,472,343 | 74.7 | 6,979,796 | 72.8 | 293,299 | 55.5 | 330,805 | 0.6 |
| HBB-3 | 762,560 | 3,072,583 | 9,994,200 | 511,302 | 67.1 | 2,299,623 | 74.8 | 7,295,451 | 73.0 | 283,360 | 55.4 | 319,728 | 0.6 |
| HBG1-1 | 185,967 | 5,162,824 | 14,791,792 | 120,433 | 64.8 | 4,949,744 | 95.9 | 14,271,858 | 96.5 | 53,434 | 44.4 | 107,176 | 0.9 |
| HBG1-2 | 175,408 | 5,018,528 | 12,030,988 | 120,273 | 68.6 | 4,820,630 | 96.1 | 11,613,726 | 96.5 | 53,323 | 44.3 | 107,094 | 0.9 |
| HBG1-3 | 184,771 | 4,015,818 | 16,150,944 | 120,736 | 65.3 | 3,837,946 | 95.6 | 15,589,891 | 96.5 | 53,489 | 44.3 | 107,672 | 0.9 |
| HNF4A-1 | 543,584 | 10,113,379 | 13,852,568 | 463,549 | 85.3 | 9,618,823 | 95.1 | 13,191,109 | 95.2 | 190,726 | 41.1 | 446,023 | 1.0 |
| HNF4A-2 | 542,733 | 8,186,687 | 14,864,839 | 462,195 | 85.2 | 7,774,829 | 95.0 | 14,159,802 | 95.3 | 190,130 | 41.1 | 444,659 | 1.0 |
| HNF4A-3 | 561,090 | 9,046,256 | 19,619,664 | 464,283 | 82.7 | 8,580,599 | 94.9 | 18,684,171 | 95.2 | 191,001 | 41.1 | 446,731 | 1.0 |
| IRF4-1 | 728,932 | 32,932,153 | 27,880,905 | 284,578 | 39.0 | 30,206,386 | 91.7 | 25,292,566 | 90.7 | 75,644 | 26.6 | 450,535 | 1.6 |
| IRF4-2 | 567,407 | 29,656,398 | 15,481,379 | 277,818 | 49.0 | 27,343,569 | 92.2 | 14,048,886 | 90.7 | 74,623 | 26.9 | 435,912 | 1.6 |
| IRF4-3 | 740,887 | 31,370,721 | 29,406,932 | 282,446 | 38.1 | 28,740,624 | 91.6 | 26,627,825 | 90.5 | 75,361 | 26.7 | 445,782 | 1.6 |
| IRF6-1 | 151,722 | 9,739,620 | 8,560,920 | 25,571 | 16.9 | 9,387,362 | 96.4 | 8,135,409 | 95.0 | 3,200 | 12.5 | 60,112 | 2.4 |
| IRF6-2 | 119,932 | 7,757,534 | 5,329,576 | 25,366 | 21.2 | 7,495,059 | 96.6 | 5,057,007 | 94.9 | 3,175 | 12.5 | 59,593 | 2.3 |
| IRF6-3 | 172,660 | 8,680,655 | 14,173,746 | 25,715 | 14.9 | 8,324,169 | 95.9 | 13,503,071 | 95.3 | 3,222 | 12.5 | 60,449 | 2.4 |
| LDLR-1 | 130,525 | 5,638,805 | 1,273,404 | 49,591 | 38.0 | 5,105,608 | 90.5 | 1,064,829 | 83.6 | 9,909 | 20.0 | 90,699 | 1.8 |
| LDLR-2 | 161,476 | 4,213,346 | 2,399,123 | 54,118 | 33.5 | 3,761,940 | 89.3 | 2,017,701 | 84.1 | 10,524 | 19.4 | 100,845 | 1.9 |
| LDLR-3 | 161,006 | 4,749,497 | 2,363,067 | 53,263 | 33.1 | 4,256,408 | 89.6 | 1,986,608 | 84.1 | 10,462 | 19.6 | 98,735 | 1.9 |
| LDLR.2-1 | 191,367 | 4,867,064 | 17,645,358 | 126,173 | 65.9 | 4,508,376 | 92.6 | 16,491,468 | 93.5 | 51,876 | 41.1 | 122,565 | 1.0 |
| LDLR.2-2 | 190,807 | 4,788,780 | 18,280,817 | 126,106 | 66.1 | 4,435,204 | 92.6 | 17,086,851 | 93.5 | 51,828 | 41.1 | 122,517 | 1.0 |
| LDLR.2-3 | 193,991 | 5,181,072 | 17,957,299 | 126,228 | 65.1 | 4,800,137 | 92.6 | 16,776,656 | 93.4 | 51,907 | 41.1 | 122,565 | 1.0 |
| MSMB-1 | 685,961 | 6,701,186 | 17,507,026 | 601,035 | 87.6 | 6,224,222 | 92.9 | 16,299,420 | 93.1 | 97,346 | 16.2 | 1,221,342 | 2.0 |
| MSMB-2 | 682,883 | 7,320,007 | 13,849,695 | 601,313 | 88.1 | 6,805,746 | 93.0 | 12,888,944 | 93.1 | 97,407 | 16.2 | 1,221,803 | 2.0 |
| MSMB-3 | 670,037 | 6,390,696 | 11,781,755 | 594,914 | 88.8 | 5,944,966 | 93.0 | 10,973,851 | 93.1 | 96,308 | 16.2 | 1,208,546 | 2.0 |

| | | | | | | | | | | | | | |
|---|---|---|---|---|---|---|---|---|---|---|---|---|---|
| MYCs1-1 | 553,764 | 6,429,193 | 6,814,583 | 113,598 | 20.5 | 1,731,164 | 26.9 | 1,545,201 | 22.7 | 2,878 | 2.5 | 268,226 | 2.4 |
| MYCs1-2 | 687,851 | 8,899,471 | 6,461,177 | 140,778 | 20.5 | 2,369,217 | 26.6 | 1,463,349 | 22.6 | 3,683 | 2.6 | 331,779 | 2.4 |
| MYCs1-3 | 657,970 | 8,930,292 | 26,961,629 | 113,323 | 17.2 | 2,363,234 | 26.5 | 6,025,793 | 22.3 | 2,933 | 2.6 | 266,889 | 2.4 |
| MYCs2-1 | 242,405 | 10,383,834 | 25,278,391 | 47,311 | 19.5 | 9,761,909 | 94.0 | 23,938,321 | 94.7 | 7,589 | 16.0 | 94,555 | 2.0 |
| MYCs2-2 | 238,102 | 10,450,639 | 24,253,590 | 47,256 | 19.8 | 9,828,501 | 94.0 | 22,978,218 | 94.7 | 7,569 | 16.0 | 94,471 | 2.0 |
| MYCs2-3 | 230,792 | 10,097,657 | 22,738,159 | 47,230 | 20.5 | 9,500,218 | 94.1 | 21,544,683 | 94.8 | 7,565 | 16.0 | 94,496 | 2.0 |
| PKLR-24h-1 | 173,084 | 60,070,760 | 2,163,171 | 46,646 | 26.9 | 58,491,803 | 97.4 | 1,955,698 | 90.4 | 11,769 | 25.2 | 71,122 | 1.5 |
| PKLR-24h-2 | 138,614 | 60,223,777 | 1,429,279 | 45,523 | 32.8 | 58,768,966 | 97.6 | 1,288,776 | 90.2 | 11,463 | 25.2 | 69,411 | 1.5 |
| PKLR-24h-3 | 149,197 | 63,764,262 | 1,519,080 | 44,389 | 29.8 | 62,203,474 | 97.6 | 1,359,753 | 89.5 | 11,326 | 25.5 | 67,361 | 1.5 |
| PKLR-48h-1 | 136,270 | 7,455,307 | 4,969,789 | 49,614 | 36.4 | 7,208,398 | 96.7 | 4,723,095 | 95.0 | 12,465 | 25.1 | 75,773 | 1.5 |
| PKLR-48h-2 | 154,324 | 8,986,733 | 6,822,420 | 50,683 | 32.8 | 8,680,506 | 96.6 | 6,504,490 | 95.3 | 12,624 | 24.9 | 77,809 | 1.5 |
| PKLR-48h-3 | 191,654 | 8,862,707 | 7,912,466 | 53,256 | 27.8 | 8,510,594 | 96.0 | 7,457,785 | 94.3 | 13,132 | 24.7 | 82,482 | 1.5 |
| RET-1 | 212,407 | 6,490,718 | 5,332,490 | 162,442 | 76.5 | 5,777,363 | 89.0 | 4,740,932 | 88.9 | 46,024 | 28.3 | 247,543 | 1.5 |
| RET-2 | 209,619 | 5,497,713 | 9,949,970 | 163,221 | 77.9 | 4,891,505 | 89.0 | 8,864,017 | 89.1 | 46,268 | 28.3 | 248,715 | 1.5 |
| RET-3 | 228,711 | 7,153,233 | 9,256,893 | 163,304 | 71.4 | 6,351,594 | 88.8 | 8,224,281 | 88.8 | 46,292 | 28.3 | 248,789 | 1.5 |
| SORT1.2-1 | 361,601 | 18,633,183 | 6,148,782 | 103,019 | 28.5 | 14,099,010 | 75.7 | 4,443,731 | 72.3 | 17,317 | 16.8 | 204,598 | 2.0 |
| SORT1.2-2 | 377,463 | 15,970,828 | 7,364,456 | 103,869 | 27.5 | 12,028,676 | 75.3 | 5,314,027 | 72.2 | 17,426 | 16.8 | 206,638 | 2.0 |
| SORT1.2-3 | 268,226 | 10,332,218 | 5,286,779 | 102,015 | 38.0 | 7,838,126 | 75.9 | 3,903,519 | 73.8 | 17,227 | 16.9 | 202,268 | 2.0 |
| SORT1-1 | 746,577 | 16,673,908 | 31,367,939 | 39,578 | 5.3 | 14,474,306 | 86.8 | 24,750,248 | 78.9 | 5,307 | 13.4 | 88,124 | 2.2 |
| SORT1-2 | 723,742 | 18,962,383 | 19,747,191 | 39,511 | 5.5 | 16,517,700 | 87.1 | 15,807,222 | 80.0 | 5,300 | 13.4 | 87,955 | 2.2 |
| SORT1-3 | 748,314 | 20,320,267 | 17,953,683 | 39,516 | 5.3 | 17,683,609 | 87.0 | 14,046,965 | 78.2 | 5,309 | 13.4 | 87,964 | 2.2 |
| SORT1-flip-1 | 331,589 | 12,166,621 | 29,066,644 | 140,787 | 42.5 | 9,953,413 | 81.8 | 24,014,577 | 82.6 | 31,099 | 22.1 | 235,726 | 1.7 |
| SORT1-flip-2 | 313,075 | 11,542,062 | 25,124,427 | 140,650 | 44.9 | 9,455,478 | 81.9 | 20,763,311 | 82.6 | 31,109 | 22.1 | 235,379 | 1.7 |
| SORT1-flip-3 | 317,684 | 11,916,102 | 25,362,908 | 140,795 | 44.3 | 9,759,092 | 81.9 | 20,955,947 | 82.6 | 31,113 | 22.1 | 235,724 | 1.7 |
| TCF7L2-1 | 374,223 | 15,108,800 | 23,655,355 | 100,157 | 26.8 | 14,202,439 | 94.0 | 22,260,177 | 94.1 | 18,273 | 18.2 | 185,613 | 1.9 |
| TCF7L2-2 | 417,767 | 14,956,164 | 32,468,628 | 100,350 | 24.0 | 14,006,822 | 93.7 | 30,546,904 | 94.1 | 18,328 | 18.3 | 185,912 | 1.9 |
| TCF7L2-3 | 368,615 | 14,520,864 | 23,374,207 | 100,134 | 27.2 | 13,646,053 | 94.0 | 21,988,603 | 94.1 | 18,278 | 18.3 | 185,497 | 1.9 |
| TERT-HEK-1 | 335,270 | 11,248,490 | 11,378,602 | 58,565 | 17.5 | 7,294,186 | 64.8 | 7,127,589 | 62.6 | 13,188 | 22.5 | 94,368 | 1.6 |
| TERT-HEK-2 | 171,124 | 2,560,571 | 3,994,303 | 54,373 | 31.8 | 1,678,719 | 65.6 | 2,574,708 | 64.5 | 12,301 | 22.6 | 87,278 | 1.6 |
| TERT-HEK-3 | 183,808 | 3,252,096 | 3,789,458 | 55,035 | 29.9 | 2,132,221 | 65.6 | 2,403,315 | 63.4 | 12,438 | 22.6 | 88,395 | 1.6 |
| TERT-GBM-1 | 355,220 | 13,418,248 | 19,211,451 | 58,007 | 16.3 | 8,720,079 | 65.0 | 12,333,250 | 64.2 | 13,065 | 22.5 | 93,285 | 1.6 |
| TERT-GBM-2 | 267,946 | 11,585,462 | 8,259,377 | 56,500 | 21.1 | 7,598,877 | 65.6 | 5,308,817 | 64.3 | 12,796 | 22.6 | 90,455 | 1.6 |
| TERT-GBM-3 | 321,270 | 10,799,638 | 15,808,970 | 57,477 | 17.9 | 7,012,582 | 64.9 | 10,147,049 | 64.2 | 12,968 | 22.6 | 92,378 | 1.6 |
| TERT-GAa-1 | 330,418 | 13,997,031 | 12,720,466 | 54,526 | 16.5 | 9,157,350 | 65.4 | 8,074,884 | 63.5 | 12,293 | 22.5 | 87,683 | 1.6 |
| TERT-GAa-2 | 273,034 | 13,438,634 | 7,480,710 | 52,841 | 19.4 | 8,872,520 | 66.0 | 4,759,939 | 63.6 | 11,896 | 22.5 | 84,944 | 1.6 |
| TERT-GAa-3 | 330,409 | 13,973,219 | 11,210,584 | 54,503 | 16.5 | 9,142,186 | 65.4 | 7,061,935 | 63.0 | 12,263 | 22.5 | 87,683 | 1.6 |
| TERT-GSc-1 | 274,950 | 15,911,173 | 6,513,278 | 54,440 | 19.8 | 10,480,550 | 65.9 | 4,149,292 | 63.7 | 12,280 | 22.6 | 87,358 | 1.6 |
| TERT-GSc-2 | 313,078 | 15,921,742 | 9,269,517 | 55,079 | 17.6 | 10,438,703 | 65.6 | 5,878,186 | 63.4 | 12,447 | 22.6 | 88,329 | 1.6 |
| TERT-GSc-3 | 319,046 | 16,567,050 | 9,177,995 | 54,956 | 17.2 | 10,870,900 | 65.6 | 5,814,748 | 63.4 | 12,434 | 22.6 | 88,139 | 1.6 |
| UC88-1 | 189,033 | 10,320,132 | 12,162,659 | 58,087 | 30.7 | 8,982,208 | 87.0 | 10,610,535 | 87.2 | 6,939 | 11.9 | 135,643 | 2.3 |
| UC88-2 | 282,779 | 13,218,404 | 25,762,258 | 58,905 | 20.8 | 11,430,470 | 86.5 | 22,389,889 | 86.9 | 7,019 | 11.9 | 137,570 | 2.3 |
| UC88-3 | 130,950 | 6,816,894 | 6,412,974 | 57,784 | 44.1 | 5,958,971 | 87.4 | 5,621,456 | 87.7 | 6,883 | 11.9 | 134,912 | 2.3 |
| ZFAND3-1 | 762,485 | 13,009,004 | 9,895,152 | 146,235 | 19.2 | 5,678,835 | 43.7 | 4,176,057 | 42.2 | 10,071 | 6.9 | 427,345 | 2.9 |
| ZFAND3-2 | 730,676 | 15,242,143 | 8,146,864 | 144,184 | 19.7 | 6,739,354 | 44.2 | 3,433,906 | 42.2 | 9,927 | 6.9 | 421,275 | 2.9 |
| ZFAND3-3 | 739,848 | 15,163,717 | 7,536,228 | 145,076 | 19.6 | 6,669,983 | 44.0 | 3,179,378 | 42.2 | 9,994 | 6.9 | 423,932 | 2.9 |

| | | | | | | | | | | | | |
|---|---|---|---|---|---|---|---|---|---|---|---|---|
| ZRS-h13-1 | 753,735 | 5,879,814 | 15,487,602 | 614,082 | 81.5 | 5,123,141 | 87.1 | 13,538,056 | 87.4 | 106,980 | 17.4 | 1,247,544 | 2.0 |
| ZRS-h13-2 | 697,780 | 5,652,278 | 5,038,577 | 581,126 | 83.3 | 4,948,648 | 87.6 | 4,404,719 | 87.4 | 101,040 | 17.4 | 1,181,566 | 2.0 |
| ZRS-h13-3 | 764,586 | 6,731,343 | 15,815,800 | 620,257 | 81.1 | 5,866,656 | 87.2 | 13,817,121 | 87.4 | 108,107 | 17.4 | 1,259,238 | 2.0 |
| ZRS-h13h2-1 | 714,593 | 4,064,366 | 13,354,202 | 590,306 | 82.6 | 3,544,054 | 87.2 | 11,695,065 | 87.6 | 102,860 | 17.4 | 1,199,781 | 2.0 |
| ZRS-h13h2-2 | 719,151 | 4,510,062 | 11,064,121 | 594,407 | 82.7 | 3,935,117 | 87.3 | 9,680,563 | 87.5 | 103,505 | 17.4 | 1,208,072 | 2.0 |
| ZRS-h13h2-3 | 736,156 | 4,822,109 | 15,527,744 | 604,186 | 82.1 | 4,203,127 | 87.2 | 13,583,482 | 87.5 | 105,191 | 17.4 | 1,228,001 | 2.0 |

*Supplementary Table 5*

**Correlation of SORT1 and LDLR with different minimum required tags.** Pearson and Spearman correlation coefficients between LDLR and SORT1 replicate experiments requiring different minimum number of tags readouts per substitution or 1 bp deletion variant included.

| LDLR | Pearson | Spearman | SORT1 | Pearson | Spearman |
|---|---|---|---|---|---|
| *All* | 0.93292 | 0.81082 | *All* | 0.93912 | 0.83928 |
| *Min 10* | 0.97144 | 0.86273 | *Min 10* | 0.96099 | 0.86436 |
| *Min 20* | 0.97484 | 0.86045 | *Min 20* | 0.96796 | 0.87945 |
| *Min 30* | 0.97664 | 0.86358 | *Min 30* | 0.97311 | 0.89615 |
| *Min 40* | 0.97827 | 0.87610 | *Min 40* | 0.97626 | 0.90280 |
| *Min 50* | 0.98004 | 0.87905 | *Min 50* | 0.97793 | 0.90508 |

*Supplementary Table 6*

**Variant coverage with different thresholds of required tags.** Single nucleotide coverage and 1 bp deletion coverage obtained after combining transfection replicates, tabulated for different minimum numbers of tags. MYC (rs11986220) is abbreviated to MYCs1 and MYC (rs6983267) abbreviated to MYCs2. TERT-GBM-SiGABPA and TERT-GBM-SiScramble-2 are abbreviated to TERT-GAa and TERT-GSc, respectively.

| Combined Exp. | Len. | All | | | | Min 10 Obs | | | | Min 20 Obs | | | |
|---|---|---|---|---|---|---|---|---|---|---|---|---|---|
| | | SNVs | | 1bpDel | | SNVs | | 1bpDel | | SNVs | | 1bpDel | |
| *BCL11A* | 600 | 1799 | 99.94% | 263 | 43.83% | 1798 | 99.89% | 187 | 31.17% | 1794 | 99.67% | 167 | 27.83% |
| *F9* | 303 | 904 | 99.45% | 80 | 26.40% | 888 | 97.69% | 64 | 21.12% | 852 | 93.73% | 52 | 17.16% |
| *FOXE1* | 600 | 1800 | 100.00% | 248 | 41.33% | 1793 | 99.61% | 130 | 21.67% | 1768 | 98.22% | 111 | 18.50% |
| *GP1BB* | 385 | 1154 | 99.91% | 114 | 29.61% | 1154 | 99.91% | 85 | 22.08% | 1154 | 99.91% | 79 | 20.52% |
| *HBB* | 187 | 561 | 100.00% | 62 | 33.16% | 561 | 100.00% | 35 | 18.72% | 561 | 100.00% | 34 | 18.18% |
| *HBG1* | 274 | 822 | 100.00% | 85 | 31.02% | 820 | 99.76% | 68 | 24.82% | 811 | 98.66% | 61 | 22.26% |
| *HNF4A* | 285 | 855 | 100.00% | 122 | 42.81% | 854 | 99.88% | 77 | 27.02% | 854 | 99.88% | 69 | 24.21% |
| *IRF4* | 451 | 1353 | 100.00% | 157 | 34.81% | 1353 | 100.00% | 105 | 23.28% | 1353 | 100.00% | 91 | 20.18% |
| *IRF6* | 601 | 1785 | 99.00% | 121 | 20.13% | 1676 | 92.96% | 63 | 10.48% | 1497 | 83.03% | 41 | 6.82% |
| *LDLR* | 318 | 954 | 100.00% | 129 | 40.57% | 945 | 99.06% | 59 | 18.55% | 924 | 96.86% | 49 | 15.41% |
| *LDLR.2* | 318 | 954 | 100.00% | 139 | 43.71% | 950 | 99.58% | 74 | 23.27% | 941 | 98.64% | 57 | 17.92% |
| *MSMB* | 593 | 1779 | 100.00% | 247 | 41.65% | 1778 | 99.94% | 140 | 23.61% | 1777 | 99.89% | 124 | 20.91% |
| *MYCs1* | 464 | 1392 | 100.00% | 293 | 63.15% | 1375 | 98.78% | 183 | 39.44% | 1339 | 96.19% | 159 | 34.27% |
| *MYCs2* | 600 | 1792 | 99.56% | 158 | 26.33% | 1715 | 95.28% | 94 | 15.67% | 1578 | 87.67% | 77 | 12.83% |
| *PKLR-24h* | 470 | 1407 | 99.79% | 369 | 78.51% | 1369 | 97.09% | 134 | 28.51% | 1279 | 90.71% | 83 | 17.66% |
| *PKLR-48h* | 470 | 1407 | 99.79% | 387 | 82.34% | 1379 | 97.80% | 154 | 32.77% | 1307 | 92.70% | 94 | 20.00% |
| *RET* | 600 | 1797 | 99.83% | 165 | 27.50% | 1776 | 98.67% | 135 | 22.50% | 1752 | 97.33% | 127 | 21.17% |
| *SORT1* | 600 | 1790 | 99.44% | 137 | 22.83% | 1710 | 95.00% | 96 | 16.00% | 1572 | 87.33% | 76 | 12.67% |
| *SORT1.2* | 600 | 1798 | 99.89% | 200 | 33.33% | 1787 | 99.28% | 131 | 21.83% | 1760 | 97.78% | 114 | 19.00% |
| *SORT1-flip* | 600 | 1791 | 99.50% | 183 | 30.50% | 1755 | 97.50% | 129 | 21.50% | 1709 | 94.94% | 116 | 19.33% |
| *TCF7L2* | 600 | 1770 | 98.33% | 140 | 23.33% | 1700 | 94.44% | 101 | 16.83% | 1650 | 91.67% | 87 | 14.50% |
| *TERT-GAa* | 259 | 777 | 100.00% | 197 | 76.06% | 769 | 98.97% | 100 | 38.61% | 747 | 96.14% | 77 | 29.73% |
| *TERT-GBM* | 259 | 777 | 100.00% | 196 | 75.68% | 770 | 99.10% | 108 | 41.70% | 751 | 96.65% | 78 | 30.12% |
| *TERT-GSc* | 259 | 777 | 100.00% | 197 | 76.06% | 769 | 98.97% | 97 | 37.45% | 750 | 96.53% | 76 | 29.34% |
| *TERT-HEK* | 259 | 777 | 100.00% | 198 | 76.45% | 770 | 99.10% | 95 | 36.68% | 753 | 96.91% | 72 | 27.80% |
| *UC88* | 590 | 1761 | 99.49% | 203 | 34.41% | 1706 | 96.38% | 134 | 22.71% | 1642 | 92.77% | 112 | 18.98% |
| *ZFAND3* | 579 | 1736 | 99.94% | 276 | 47.67% | 1735 | 99.88% | 158 | 27.29% | 1725 | 99.31% | 140 | 24.18% |
| *ZRS-h13* | 485 | 1455 | 100.00% | 206 | 42.47% | 1455 | 100.00% | 116 | 23.92% | 1455 | 100.00% | 106 | 21.86% |
| *ZRS-h13h2* | 485 | 1455 | 100.00% | 207 | 42.68% | 1455 | 100.00% | 117 | 24.12% | 1455 | 100.00% | 106 | 21.86% |
| **Average** | | | 99.79% | | 44.43% | | 98.43% | | 25.29% | | 95.97% | | 20.87% |
| **Minimum** | | | 98.33% | | 20.13% | | 92.96% | | 10.48% | | 83.03% | | 6.82% |
| **Maximum** | | | 100.00% | | 82.34% | | 100.00% | | 41.70% | | 100.00% | | 34.27% |

| Comb. Exp. | Len. | Min 30 Obs | | | | Min 40 Obs | | | | Min 50 Obs | | | |
|---|---|---|---|---|---|---|---|---|---|---|---|---|---|
| | | SNVs | | 1bpDel | | SNVs | | 1bpDel | | SNVs | | 1bpDel | |
| *BCL11A* | 600 | 1784 | 99.11% | 153 | 25.50% | 1772 | 98.44% | 145 | 24.17% | 1750 | 97.22% | 141 | 23.50% |
| *F9* | 303 | 821 | 90.32% | 43 | 14.19% | 790 | 86.91% | 41 | 13.53% | 768 | 84.49% | 36 | 11.88% |
| *FOXE1* | 600 | 1729 | 96.06% | 101 | 16.83% | 1685 | 93.61% | 93 | 15.50% | 1621 | 90.06% | 86 | 14.33% |
| *GP1BB* | 385 | 1153 | 99.83% | 74 | 19.22% | 1152 | 99.74% | 72 | 18.70% | 1145 | 99.13% | 70 | 18.18% |
| *HBB* | 187 | 561 | 100.00% | 33 | 17.65% | 561 | 100.00% | 33 | 17.65% | 559 | 99.64% | 33 | 17.65% |
| *HBG1* | 274 | 802 | 97.57% | 55 | 20.07% | 783 | 95.26% | 47 | 17.15% | 760 | 92.46% | 46 | 16.79% |
| *HNF4A* | 285 | 853 | 99.77% | 69 | 24.21% | 853 | 99.77% | 67 | 23.51% | 853 | 99.77% | 66 | 23.16% |
| *IRF4* | 451 | 1353 | 100.00% | 84 | 18.63% | 1344 | 99.33% | 81 | 17.96% | 1337 | 98.82% | 80 | 17.74% |
| *IRF6* | 601 | 1317 | 73.04% | 32 | 5.32% | 1158 | 64.23% | 23 | 3.83% | 1054 | 58.46% | 13 | 2.16% |
| *LDLR* | 318 | 892 | 93.50% | 45 | 14.15% | 843 | 88.36% | 43 | 13.52% | 787 | 82.49% | 40 | 12.58% |
| *LDLR.2* | 318 | 929 | 97.38% | 50 | 15.72% | 906 | 94.97% | 47 | 14.78% | 879 | 92.14% | 44 | 13.84% |
| *MSMB* | 593 | 1777 | 99.89% | 117 | 19.73% | 1776 | 99.83% | 114 | 19.22% | 1774 | 99.72% | 113 | 19.06% |
| *MYCs1* | 464 | 1267 | 91.02% | 142 | 30.60% | 1204 | 86.49% | 126 | 27.16% | 1132 | 81.32% | 117 | 25.22% |
| *MYCs2* | 600 | 1462 | 81.22% | 67 | 11.17% | 1330 | 73.89% | 51 | 8.50% | 1227 | 68.17% | 48 | 8.00% |
| *PKLR-24h* | 470 | 1170 | 82.98% | 59 | 12.55% | 1082 | 76.74% | 48 | 10.21% | 1007 | 71.42% | 36 | 7.66% |
| *PKLR-48h* | 470 | 1205 | 85.46% | 66 | 14.04% | 1126 | 79.86% | 53 | 11.28% | 1049 | 74.40% | 48 | 10.21% |
| *RET* | 600 | 1726 | 95.89% | 119 | 19.83% | 1665 | 92.50% | 110 | 18.33% | 1612 | 89.56% | 101 | 16.83% |
| *SORT1* | 600 | 1416 | 78.67% | 66 | 11.00% | 1280 | 71.11% | 58 | 9.67% | 1178 | 65.44% | 54 | 9.00% |
| *SORT1.2* | 600 | 1712 | 95.11% | 103 | 17.17% | 1665 | 92.50% | 94 | 15.67% | 1617 | 89.83% | 83 | 13.83% |
| *SORT1-flip* | 600 | 1661 | 92.28% | 102 | 17.00% | 1593 | 88.50% | 95 | 15.83% | 1539 | 85.50% | 90 | 15.00% |
| *TCF7L2* | 600 | 1605 | 89.17% | 76 | 12.67% | 1527 | 84.83% | 64 | 10.67% | 1458 | 81.00% | 58 | 9.67% |
| *TERT-GAa* | 259 | 714 | 91.89% | 61 | 23.55% | 676 | 87.00% | 57 | 22.01% | 631 | 81.21% | 50 | 19.31% |
| *TERT-GBM* | 259 | 721 | 92.79% | 63 | 24.32% | 682 | 87.77% | 59 | 22.78% | 646 | 83.14% | 52 | 20.08% |
| *TERT-GSc* | 259 | 716 | 92.15% | 62 | 23.94% | 675 | 86.87% | 58 | 22.39% | 636 | 81.85% | 48 | 18.53% |
| *TERT-HEK* | 259 | 718 | 92.41% | 62 | 23.94% | 682 | 87.77% | 56 | 21.62% | 639 | 82.24% | 49 | 18.92% |
| *UC88* | 590 | 1562 | 88.25% | 98 | 16.61% | 1453 | 82.09% | 82 | 13.90% | 1357 | 76.67% | 75 | 12.71% |
| *ZFAND3* | 579 | 1715 | 98.73% | 129 | 22.28% | 1689 | 97.24% | 117 | 20.21% | 1664 | 95.80% | 101 | 17.44% |
| *ZRS-h13* | 485 | 1454 | 99.93% | 100 | 20.62% | 1452 | 99.79% | 97 | 20.00% | 1451 | 99.73% | 94 | 19.38% |
| *ZRS-h13h2* | 485 | 1454 | 99.93% | 99 | 20.41% | 1453 | 99.86% | 97 | 20.00% | 1450 | 99.66% | 95 | 19.59% |
| **Average** | | | 92.91% | | 18.38% | | 89.49% | | 16.89% | | 86.25% | | 15.59% |
| **Minimum** | | | 73.04% | | 5.32% | | 64.23% | | 3.83% | | 58.46% | | 2.16% |
| **Maximum** | | | 100.00% | | 30.60% | | 100.00% | | 27.16% | | 99.77% | | 25.22% |

##### *Supplementary Table 7*

**Pearson correlation across different transfection replicates.** Correlation from the three transfection replicates of fitted SNV and 1 bp deletion variant effects divided by their standard deviation (requiring at least 10 tags in each replicate). MYC (rs11986220) is abbreviated to MYCs1 and MYC (rs6983267) abbreviated to MYCs2. TERT-HEK293T, TER-GBM, TERT-GBM-SiGABPA and TERT-GBM-SiScramble-2 are abbreviated to TERT-H, TERT-G, TERT-GAa and TERT-GSc, respectively. The left side considers all estimates obtained, the right side tries to exclude nonsignificant allele effects by requiring a lenient p-value threshold (<0.01) in at least one of the replicates. We use the reduction in included alleles as a proxy for the proportion of alleles with regulatory activity for each element (Frac.). We note that this estimate is confounded by other factors of experimental reproducibility, as can be seen from the variance observed for elements with multiple experiments (LDLR, PKLR, SORT1, TERT).

| | **Min. 10 tags** | | | | **Min. 10 tags, excl. non-significant effects** | | | | |
|---|---|---|---|---|---|---|---|---|---|
| **Name** | **Rep 1 v. 2** | **Rep 1 v. 3** | **Rep 2 v. 3** | **Mean** | **Rep 1 v. 2** | **Rep 1 v. 3** | **Rep 2 v. 3** | **Mean** | **Frac.** |
| BCL11A | 0.38 (1937) | 0.42 (1938) | 0.35 (1938) | 0.38 | 0.70 (107) | 0.69 (108) | 0.66 (95) | 0.68 | 5% |
| F9 | 0.88 (863) | 0.87 (861) | 0.90 (865) | 0.88 | 0.95 (196) | 0.95 (215) | 0.96 (210) | 0.95 | 21% |
| FOXE1 | 0.17 (1827) | 0.12 (1823) | 0.18 (1827) | 0.16 | 0.46 (65) | 0.23 (63) | 0.38 (52) | 0.36 | 3% |
| GP1BB | 0.88 (1226) | 0.89 (1225) | 0.88 (1227) | 0.88 | 0.94 (268) | 0.95 (249) | 0.94 (276) | 0.94 | 21% |
| HBB | 0.78 (594) | 0.75 (594) | 0.77 (594) | 0.77 | 0.89 (119) | 0.89 (115) | 0.89 (119) | 0.89 | 19% |
| HBG1 | 0.92 (856) | 0.92 (856) | 0.91 (858) | 0.92 | 0.96 (239) | 0.95 (244) | 0.94 (246) | 0.95 | 27% |
| HNF4A | 0.88 (922) | 0.89 (922) | 0.88 (922) | 0.89 | 0.95 (192) | 0.96 (182) | 0.96 (171) | 0.96 | 19% |
| IRF4 | 0.99 (1437) | 0.99 (1438) | 0.99 (1437) | 0.99 | 0.99 (789) | 0.99 (793) | 0.99 (777) | 0.99 | 52% |
| IRF6 | 0.96 (1347) | 0.97 (1351) | 0.96 (1347) | 0.96 | 0.98 (340) | 0.98 (390) | 0.98 (371) | 0.98 | 19% |
| LDLR | 0.97 (930) | 0.97 (928) | 0.98 (936) | 0.98 | 0.98 (354) | 0.98 (352) | 0.99 (369) | 0.98 | 34% |
| LDLR.2 | 0.99 (979) | 0.99 (979) | 0.99 (979) | 0.99 | 1.00 (455) | 1.00 (454) | 1.00 (448) | 1.00 | 42% |
| MSMB | 0.89 (1893) | 0.88 (1894) | 0.88 (1893) | 0.88 | 0.96 (477) | 0.95 (466) | 0.95 (469) | 0.95 | 23% |
| MYCs1 | 0.33 (1385) | 0.26 (1355) | 0.35 (1376) | 0.31 | 0.58 (83) | 0.61 (64) | 0.69 (77) | 0.63 | 5% |
| MYCs2 | 0.75 (1529) | 0.76 (1529) | 0.76 (1529) | 0.75 | 0.92 (136) | 0.93 (133) | 0.93 (138) | 0.93 | 7% |
| PKLR 24h | 0.87 (1220) | 0.85 (1208) | 0.86 (1209) | 0.86 | 0.94 (328) | 0.93 (326) | 0.93 (322) | 0.93 | 20% |
| PKLR 48h | 0.91 (1246) | 0.91 (1255) | 0.92 (1267) | 0.91 | 0.95 (366) | 0.95 (383) | 0.96 (386) | 0.95 | 22% |
| RET | 0.70 (1845) | 0.68 (1845) | 0.75 (1845) | 0.71 | 0.87 (306) | 0.85 (308) | 0.89 (325) | 0.87 | 16% |
| SORT1 | 0.99 (1483) | 0.99 (1482) | 0.99 (1483) | 0.99 | 1.00 (753) | 1.00 (756) | 1.00 (756) | 1.00 | 39% |
| SORT1.2 | 0.98 (1815) | 0.97 (1812) | 0.98 (1815) | 0.98 | 0.99 (793) | 0.98 (773) | 0.98 (790) | 0.99 | 39% |
| SORT1-flip | 0.99 (1763) | 0.99 (1763) | 0.99 (1763) | 0.99 | 1.00 (780) | 0.99 (754) | 0.99 (778) | 0.99 | 39% |
| TCF7L2 | 0.75 (1680) | 0.77 (1681) | 0.75 (1682) | 0.76 | 0.91 (221) | 0.92 (230) | 0.91 (215) | 0.91 | 12% |
| TERT-G | 0.97 (778) | 0.97 (785) | 0.96 (780) | 0.97 | 0.98 (341) | 0.98 (343) | 0.98 (341) | 0.98 | 35% |
| TERT-GAa | 0.91 (769) | 0.93 (779) | 0.91 (771) | 0.92 | 0.96 (239) | 0.96 (247) | 0.96 (244) | 0.96 | 25% |
| TERT-GSc | 0.95 (778) | 0.95 (777) | 0.95 (780) | 0.95 | 0.97 (281) | 0.97 (279) | 0.97 (273) | 0.97 | 29% |
| TERT-H | 0.87 (777) | 0.87 (781) | 0.81 (775) | 0.85 | 0.93 (216) | 0.93 (228) | 0.92 (164) | 0.93 | 21% |
| UC88 | 0.67 (1660) | 0.59 (1659) | 0.66 (1661) | 0.64 | 0.91 (140) | 0.88 (113) | 0.92 (134) | 0.90 | 7% |

| | | | | | | | | | |
|---|---|---|---|---|---|---|---|---|---|
| ZFAND3 | 0.89 (1842) | 0.90 (1841) | 0.89 (1841) | 0.89 | 0.96 (429) | 0.96 (474) | 0.96 (440) | 0.96 | 22% |
| ZRSh13 | 0.73 (1553) | 0.80 (1554) | 0.75 (1553) | 0.76 | 0.90 (194) | 0.93 (199) | 0.91 (190) | 0.91 | 12% |
| ZRSh13h2 | 0.74 (1553) | 0.77 (1552) | 0.78 (1552) | 0.76 | 0.90 (164) | 0.92 (180) | 0.93 (178) | 0.92 | 11% |

***Supplementary Table 8***

**Number of mutations (Mut.) and sites studied for a representative experiment of each element.** The number of mutations with a significant p-value ($< 10^{-5}$) is reported from the model combining transfection replicates. Counts are provided for reporter expression increasing (↑) and decreasing (↓) mutations. The overlap with single nucleotide variants present in the gnomAD r2.1 database is reported including singleton variants (gnAD columns) and excluding singleton variants (no sgl columns). Of the 31,243 mutations, 28,937 are single nucleotide variants. MYC (rs11986220) is abbreviated to MYCs1 and MYC (rs6983267) abbreviated to MYCs2.

| **Experiment** | **Mut.** | **Sites** | **Sig. ($10^{-5}$)** | ↑ | ↓ | **2-fold** | ↑ | ↓ | **gnAD** | **Sig. ($10^{-5}$)** | ↑ | ↓ | **gnAD (no sgl)** | **Sig. ($10^{-5}$)** | ↑ | ↓ |
|---|---|---|---|---|---|---|---|---|---|---|---|---|---|---|---|---|
| *BCL11A* | 1985 | 600 | 66 | 35 | 31 | 2 | 0 | 2 | 47 | 3 | 2 | 1 | 14 | 0 | 0 | 0 |
| *F9* | 952 | 303 | 142 | 76 | 66 | 2 | 1 | 1 | 11 | 2 | 1 | 1 | 4 | 0 | 0 | 0 |
| *FOXE1* | 1923 | 600 | 6 | 3 | 3 | 0 | 0 | 0 | 45 | 0 | 0 | 0 | 26 | 0 | 0 | 0 |
| *GP1BB* | 1239 | 385 | 184 | 33 | 151 | 11 | 0 | 11 | 43 | 5 | 3 | 2 | 11 | 2 | 1 | 1 |
| *HBB* | 596 | 187 | 90 | 33 | 57 | 1 | 1 | 0 | 18 | 7 | 0 | 7 | 8 | 4 | 0 | 4 |
| *HBG1* | 888 | 274 | 181 | 24 | 157 | 27 | 0 | 27 | 13 | 3 | 0 | 3 | 5 | 2 | 0 | 2 |
| *HNF4a* | 931 | 285 | 119 | 63 | 56 | 1 | 1 | 0 | 22 | 3 | 1 | 2 | 11 | 1 | 0 | 1 |
| *IRF4* | 1458 | 451 | 678 | 267 | 411 | 135 | 9 | 126 | 39 | 20 | 7 | 13 | 20 | 9 | 4 | 5 |
| *IRF6* | 1739 | 599 | 306 | 112 | 194 | 52 | 14 | 38 | 40 | 8 | 0 | 8 | 20 | 5 | 0 | 5 |
| *LDLR.2* | 1024 | 318 | 411 | 103 | 308 | 136 | 9 | 127 | 17 | 5 | 2 | 3 | 5 | 1 | 1 | 0 |
| *MSMB* | 1912 | 591 | 369 | 178 | 191 | 1 | 0 | 1 | 36 | 13 | 9 | 4 | 19 | 7 | 7 | 0 |
| *MYCs1* | 1554 | 463 | 11 | 11 | 0 | 0 | 0 | 0 | 22 | 0 | 0 | 0 | 12 | 0 | 0 | 0 |
| *MYCs2* | 1809 | 600 | 85 | 47 | 38 | 0 | 0 | 0 | 41 | 2 | 1 | 1 | 13 | 2 | 1 | 1 |
| *PKLR (48h)* | 1533 | 470 | 277 | 88 | 189 | 78 | 10 | 68 | 28 | 9 | 4 | 5 | 13 | 3 | 3 | 0 |
| *RET* | 1911 | 600 | 165 | 67 | 98 | 0 | 0 | 0 | 60 | 9 | 4 | 5 | 28 | 6 | 4 | 2 |
| *SORT1.1* | 1806 | 600 | 789 | 268 | 521 | 144 | 29 | 115 | 55 | 27 | 8 | 19 | 27 | 11 | 3 | 8 |
| *TCF7L2* | 1801 | 595 | 130 | 43 | 87 | 2 | 2 | 0 | 42 | 3 | 3 | 0 | 19 | 1 | 1 | 0 |
| *TERT (GBM)* | 878 | 259 | 293 | 99 | 194 | 48 | 6 | 42 | 24 | 13 | 6 | 7 | 10 | 4 | 3 | 1 |
| *UC88* | 1840 | 590 | 69 | 41 | 28 | 2 | 2 | 0 | 40 | 2 | 2 | 0 | 12 | 0 | 0 | 0 |
| *ZFAND3* | 1893 | 579 | 328 | 104 | 224 | 3 | 0 | 3 | 40 | 12 | 2 | 10 | 22 | 6 | 1 | 5 |
| *ZRS (+h13)* | 1571 | 485 | 131 | 96 | 35 | 0 | 0 | 0 | 24 | 0 | 0 | 0 | 12 | 0 | 0 | 0 |
| ***Sum*** | **31243** | **9834** | **4830** | **1791** | **3039** | **645** | **84** | **561** | **707** | **146** | **55** | **91** | **311** | **64** | **29** | **35** |

*Supplementary Table 9*

**Effects of specific substitution or transition (ts) vs. transversion (tv) effects for significant readouts across a representative experiment of each element.** The number of mutations with a significant p-value (< $10^{-5}$) is reported from models combining transfection replicates. The overlap with single nucleotide variants present in the gnomAD r2.1 is reported. Elements from Supplementary Table 8 were used.

| Subst | Type | All | Sig. | Sig.20% | Sig. 2-fold | Var. | Var.Sig. | Var.Sig.2 0% | Var.Sig.2 -fold |
|---|---|---|---|---|---|---|---|---|---|
| A>- | Del | 583 | 48 | 31 | 11 | n.a. | | | |
| A>C | Tv | 2232 | 205 | 174 | 46 | 19 | 1 | 1 | 0 |
| A>G | Ts | 2332 | 518 | 220 | 45 | 73 | 18 | 9 | 4 |
| A>T | Tv | 2334 | 392 | 210 | 54 | 12 | 3 | 1 | 1 |
| C>- | Del | 625 | 64 | 41 | 13 | n.a. | | | |
| C>A | Tv | 2577 | 348 | 257 | 61 | 52 | 9 | 6 | 2 |
| C>G | Tv | 2481 | 237 | 196 | 58 | 47 | 4 | 4 | 0 |
| C>T | Ts | 2607 | 520 | 246 | 49 | 172 | 48 | 20 | 3 |
| G>- | Del | 589 | 67 | 50 | 7 | n.a. | | | |
| G>A | Ts | 2552 | 543 | 265 | 44 | 141 | 30 | 15 | 1 |
| G>C | Tv | 2437 | 247 | 219 | 59 | 58 | 4 | 4 | 1 |
| G>T | Tv | 2538 | 343 | 255 | 52 | 37 | 7 | 5 | 0 |
| T>- | Del | 509 | 50 | 34 | 8 | n.a. | | | |
| T>A | Tv | 2327 | 404 | 229 | 37 | 8 | 2 | 1 | 0 |
| T>C | Ts | 2333 | 609 | 262 | 44 | 69 | 19 | 10 | 1 |
| T>G | Tv | 2187 | 235 | 206 | 47 | 19 | 1 | 0 | 0 |
| **Total** | **Dels** | **2306** | **229** | **156** | **39** | **n.a.** | | | |
| **Total** | **SNVs** | **28937** | **4601** | **2739** | **596** | **707** | **146** | **76** | **13** |
| **Total** | **Tv** | **19113** | **2411** | **1746** | **414** | **252** | **31** | **22** | **4** |
| **Total** | **Ts** | **9824** | **2190** | **993** | **182** | **455** | **115** | **54** | **9** |

***Supplementary Table 10***

**Comparison with previously described LDLR mutations showing effect on promoter activity or being implicated in disease.** This table has been modified from Khamis *et al.* 2015[48]. We are reporting $\log_2$ fold changes as well as percentages for our LDLR and LDLR.2 experiments. A reported fold change is considered "no effect" (noEff), if the associated p-Value is higher than $10^{-5}$. In all other cases, we report a variant as enhancing (Enh) or repressing (Repr), depending on the sign of the $\log_2$ fold change. Familial hypercholesterolemia is abbreviated with FH. FP1 stands for Footprint. SP1 refers to a binding site of the transcription factor SP1. SRE stands for sterol-dependent regulatory element. SREBP1 and SREBP1 are SRE repeat 1 and 2, respectively.

| **Coordinates [GRCh37, GRCh38]** | **Variant in NM_000527.4 [Khamis A et al. 2015]** | **Location** | **Promoter activity (% of WT)** | **This experiment (Exp 1/2)** | **Reference (PMID)** |
|---|---|---|---|---|---|
| 19:11199957 19:11089281 | c.-268G>T | 1 bp from FP1 | In FH and normal (polymorphism) | 0.06 (104%, noEff) -0.07 (95%, noEff) | 10484771 |
| 19:11200008 19:11089332 | c.-217C>T | 2 bp from FP1 | 160% Luciferase | 1.45 (273%, Enh) 1.40 (263%, Enh) | 10484771 |
| 19:11200010 19:11089334 | c.-215A>G | 4 bp from FP1 | 100% Luciferase [results depending on Serum level] | -0.78 (58%, Repr) -0.81 (57%, Repr) | 22881376 |
| 19:11200017 19:11089341 | c.-208A>T | Between SREBP1 and FP1 | 100% Luciferase | -1.28 (41%, Repr) -1.06 (47%, Repr) | 21538688 |
| 19:11200019 19:11089343 | c.-206C>T | Between SREBP1 and FP1 | Not tested | -0.84 (55%, Repr) -0.81 (57%, Repr) | 12052488 |
| 19:11200037 19:11089361 | c.-188C>T | SREBP1 | Not tested | -2.41 (18%, Repr) -2.84 (13%, Repr) | 16250003 |
| 19:11200069 19:11089393 | c.-156C>T | SREBP2 | Not tested | -2.52 (17%, Repr) -2.67 (15%, Repr) | 14974088 |
| 19:11200072 19:11089396 | c.-153C>T | SREBP2 | Not tested, considered pathogenic by location in conserved region (SREBP2) | -1.22 (42%, Repr) -1.36 (38%, Repr) | 15303010, 3030558, 10484771 |
| 19:11200073 19:11089397 | c.-152C>T | SREBP2 | 40% Luciferase | -1.33 (39%, Repr) -1.60 (32%, Repr) | 10484771 |
| 19:11200079 19:11089403 | c.-146C>A | Between SREBP2 and SP1 | Not tested | -0.48 (71%, noEff) -0.33 (79%, Repr) | 9259195 |

| | | | | | |
|---|---|---|---|---|---|
| 19:11200083 19:11089407 | c.-142C>T | SP1 | 20% Luciferase | -2.28 (20%, Repr) - 3.09 (11%, Repr) | 11792717 |
| 19:11200085 19:11089409 | c.-140C>G | SP1 | 7% Luciferase | -2.15 (22%, Repr) - 3.29 (10%, Repr) | 21538688 |
| 19:11200085 19:11089409 | c.-140C>T | SP1 | 6% Luciferase | -2.87 (13%, Repr) - 3.55 (8%, Repr) | 21538688 |
| 19:11200086 19:11089410 | c.-139C>A | SP1 | Not tested | -2.70 (15%, Repr) - 3.24 (10%, Repr) | 16250003 |
| 19:11200086 19:11089410 | c.-139C>G | SP1 | 26% Luciferase | -2.70 (15%, Repr) - 3.31 (10%, Repr) | 17625505 |
| 19:11200087 19:11089411 | c.-138T>C | SP1 | 24% LDLR activity | -2.36 (19%, Repr) - 2.75 (14%, Repr) | 8589690 |
| 19:11200087 19:11089411 | c.-138delT | SP1 | 25% Luciferase | - - | 14616764 |
| 19:11200088 19:11089412 | c.-137C>T | SP1 | Not tested | -2.59 (16%, Repr) - 3.52 (8%, Repr) | 1301956 |
| 19:11200089 19:11089413 | c.-136C>G | SP1 | 12% Luciferase | -2.57 (16%, Repr) - 3.06 (11%, Repr) | 21538688 |
| 19:11200089 19:11089413 | c.-136C>T | SP1 | 5% CAT assay | -2.62 (16%, Repr) - 3.51 (8%, Repr) | 7937987 |
| 19:11200090 19:11089414 | c.-135C>G | SP1 | Pathogenic, LIPOchip assay | -2.70 (15%, Repr) - 3.01 (12%, Repr) | 1301956 |
| 19:11200104 19:11089428 | c.-121T>C | Between TATA box and SP1 | 72% Luciferase | -0.56 (67%, Repr) - 0.63 (64%, Repr) | 20236128 |
| 19:11200105 19:11089429 | c.-120C>T | Between TATA box and SP1 | 3% Luciferase / 10% Luciferase | -0.14 (90%, noEff) - 0.17 (88%, Repr) | 15303010 |
| 19:11200124 19:11089448 | c.-101T>C | TATA BOX | 64% Luciferase | -0.32 (80%, Repr) - 0.31 (80%, Repr) | 22881376 |
| 19:11200137 19:11089461 | c.-88G>A | 5′UTR | 100% Luciferase | -0.23 (85%, Repr) - 0.05 (96%, noEff) | 21538688 |
| 19:11200157 19:11089481 | c.-68A>C | 5′UTR | Not tested | -0.16 (89%, noEff) - 0.04 (97%, noEff) | 15556094 |
| 19:11200189 19:11089513 | c.-36T>G | 5′UTR | 100% Luciferase | 0.10 (107%, noEff) 0.18 (113%, noEff) | 21538688 |
| 19:11200202 19:11089526 | c.-23A>C | 5′UTR | Not tested | - 0.09 (106%, noEff) | 15241806 |
| 19:11200211 19:11089535 | c.-14C>A | 5′UTR | Not tested | 0.18 (113%, noEff) -0.05 (96%, noEff) | 9259195 |
| 19:11200212 19:11089536 | c.-13A>G | 5′UTR | 100% Luciferase | -0.03 (97%, noEff) 0.09 (106%, noEff) | 20828696 |
| 19:11200220 19:11089544 | c.-5C>T | 5′UTR | Not tested | 0.01 (100%, noEff) -0.07 (95%, noEff) | 16250003 |

***Supplementary Table 11***

**Variants that are significantly different between the knockdown of the scrambled siRNA and siGABPA (non-overlapping 95% confidence intervals).** In addition the motif of the Ets-family with the highest change in the position weight matrix (PWM) is reported where available. For the majority of significantly different variants, a member of the Ets-family transcription factors is annotated or matched using PWMs in JASPAR 2018[6]. The variants NM_198253.2:c.-146C>A and c.-124C>A create similar ETV/GABPA binding motifs to the previously known variants at the same positions, as observed in previous studies[49]. A new ETV motif is created by c.-79C>A, c.-78delT and c.-77C>T, and a novel GABPA/ETS1 motif is created by c.-189A>G, c.-189A>T or c.-126C>T. Finally, two of the three less common activating variants identified in cancer studies[50–52], c.-45G>A and c.-57A>C, create an ETS motif. Following Jaspar IDs (including motif name) of the Ets-family are used: MA0062.1 (GABPA), MA0098.3 (ETS1), MA0764.1 (ETV4), MA0765.1 (ETV5), MA0645.1 (ETV6), MA0076.2 (ELK4).

| Coordinates [GRCh37, GRCh38] | Variants (NM_198253.2) | $Log_2$ variant effect scramble siRNA | $Log_2$ variant effect siGABPA | Motif | PWM score change |
|---|---|---|---|---|---|
| 5:1295293 5:1295178 | c.-189A>G | 0.56 | 0.25 | GABPA | 4.95 |
| 5:1295293 5:1295178 | c.-189A>T | 0.25 | -0.06 | GABPA | 2.95 |
| 5:1295292 5:1295177 | c.-188G>C | -1.01 | -0.30 | GABPA | -2.11 |
| 5:1295267 5:1295152 | c.-163G>A | -0.71 | -0.53 | | |
| 5:1295264 5:1295149 | c.-160C>T | -0.46 | -0.28 | | |
| 5:1295250 5:1295135 | c.-146C>T | 2.47 | 1.65 | GABPA | 5.17 |
| 5:1295248 5:1295133 | c.-144C>T | -0.36 | -0.18 | GABPA | -5.17 |
| 5:1295245 5:1295130 | c.-141G>A | 0.33 | 0.10 | GABPA | -2.59 |
| 5:1295238 5:1295123 | c.-134G>A | -0.32 | -0.15 | | |
| 5:1295237 5:1295122 | c.-133C>T | 0.24 | 0.02 | | |
| 5:1295232 5:1295117 | c.-128C>A | -1.01 | -0.64 | | |
| 5:1295232 5:1295117 | c.-128C>T | -0.75 | -0.56 | | |
| 5:1295231 5:1295116 | c.-127C>T | -0.63 | -0.46 | GABPA | -4.39 |

| | | | | | |
|---|---|---|---|---|---|
| 5:1295230 51295115 | c.-126C>T | 0.35 | 0.16 | GABPA | 1.42 |
| 5:1295228 5:1295113 | c.-124C>A | 1.75 | 1.31 | ETV4 | 5.86 |
| 5:1295228 5:1295113 | c.-124C>T | 2.9 | 1.99 | ETV6 | 7.70 |
| 5:1295225 5:1295110 | c.-121C>G | -0.21 | 0.03 | ETV6 | -15.03 |
| 5:1295224 5:1295109 | c.-120G>A | -0.23 | -0.04 | ETS1 | -13.71 |
| 5:1295215 5:1295100 | c.-111C>T | 0.46 | 0.26 | | |
| 5:1295211 5:1295096 | c.-107C>T | 0.65 | 0.46 | | |
| 5:1295210 5:1295095 | c.-106C>T | -1.11 | -0.7 | | |
| 5:1295207 5:1295092 | c.-103delC | -0.8 | -0.51 | ELK4 | -0.89 |
| 5:1295207 5:1295092 | c.-103C>T | -1.18 | -0.86 | ELK4 | 0.00 |
| 5:1295206 5:1295091 | c.-102C>A | -1.7 | -1.41 | ELK4 | -8.70 |
| 5:1295206 5:1295091 | c.-102C>T | -1.14 | -0.85 | ETV6 | -0.65 |
| 5:1295204 5:1295089 | c.-100C>A | -0.87 | -0.4 | ETV6 | -15.03 |
| 5:1295204 5:1295089 | c.-100C>T | -0.56 | -0.23 | ELK4 | -14.06 |
| 5:1295203 5:1295088 | c.-99T>A | -0.86 | -0.48 | ELK4 | -14.06 |
| 5:1295203 5:1295088 | c.-99T>C | -0.69 | -0.29 | ETV6 | -7.70 |
| 5:1295202 5:1295087 | c.-98T>A | -1.39 | -1.02 | ETV6 | -15.03 |
| 5:1295202 5:1295087 | c.-98T>C | -1.02 | -0.57 | ELK4 | -14.06 |
| 5:1295201 5:1295086 | c.-97delC | -0.98 | -0.39 | ELK4 | -2.00 |
| 5:1295201 5:1295086 | c.-97C>A | -1.13 | -0.62 | ELK4 | -11.29 |
| 5:1295201 5:1295086 | c.-97C>T | -0.85 | -0.39 | ETV5 | -14.40 |
| 5:1295200 5:1295085 | c.-96C>T | -1.15 | -0.58 | ETV5 | -14.40 |
| 5:1295199 5:1295084 | c.-95T>A | -1.03 | -0.55 | ELK4 | -14.06 |

| | | | | | |
|---|---|---|---|---|---|
| 5:1295199<br>5:1295084 | c.-95T>C | -0.49 | -0.24 | ETV4 | -3.83 |
| 5:1295197<br>5:1295082 | c.-93T>A | -0.88 | -0.36 | ETV6 | -15.03 |
| 5:1295191<br>5:1295076 | c.-87G>A | -0.32 | -0.17 | | |
| 5:1295188<br>5:1295073 | c.-84C>T | -0.47 | -0.22 | | |
| 5:1295187<br>5:1295072 | c.-83C>T | -0.51 | -0.28 | | |
| 5:1295187<br>5:1295072 | c.-83C>A | -0.63 | -0.25 | | |
| 5:1295185<br>5:1295070 | c.-81C>A | -1.01 | -0,41 | | |
| 5:1295185<br>5:1295070 | c.-81C>T | -0.66 | -0.29 | | |
| 5:1295184<br>5:1295069 | c.-80C>T | -0.6 | -0.25 | ETV6 | -0.65 |
| 5:1295183<br>5:1295068 | c.-79C>A | -0.17 | 0.19 | ETV6 | 4.84 |
| 5:1295182<br>5:1295067 | c.-78delT | 0.36 | -1.19 | ETV6 | 1.29 |
| 5:1295181<br>5:1295066 | c.-77C>T | 0.89 | 0.48 | ETV6 | 7.70 |
| 5:1295171<br>5:1295056 | c.-67C>T | 0.82 | 0.56 | | |
| 5:1295169<br>5:1295054 | c.-65C>T | 1.08 | 0.88 | | |
| 5:1295164<br>5:1295049 | c.-60T>C | 0.68 | 0.55 | GABPA | -5.17 |
| 5:1295161<br>5:1295046 | c.-57A>C | 1.37 | 0.69 | ETS1 | 13.71 |
| 5:1295159<br>5:1295044 | c.-55G>A | -0.52 | -0.31 | ETS1 | -6.92 |
| 5:1295149<br>5:1295034 | c.-45G>A | 0.03 | -0.13 | ETV5 | 0.83 |
| 5:1295146<br>5:1295031 | c.-42C>A | -0.09 | 0.38 | ETV5 | -5.72 |
| 5:1295140<br>5:1295025 | c.-36C>G | -0.22 | 0.01 | | |
| 5:1295138<br>5:1295023 | c.-34C>T | 0.12 | 0.41 | | |
| 5:1295137<br>5:1295022 | c.-33A>G | -0.05 | 0.08 | | |
| 5:1295136<br>5:1295021 | c.-32C>T | 0.18 | 0.31 | | |

| | | | |
|---|---|---|---|
| 5:1295134<br>5:1295019 | c.-30T>C | 0.14 | 0.31 |
| 5:1295133<br>5:1295018 | c.-29G>A | 0.03 | 0.25 |
| 5:1295111<br>5:1294996 | c.-7C>A | -0.17 | 0.01 |
| 5:1295104<br>5:1294989 | c.1delA | 0.55 | 0.21 |

***Supplementary Table 12***

**Pearson correlation of computational scores with measured expression effects of all SNVs with at least 10 tags.** The table reports Pearson correlation coefficients of the absolute expression effect for each element with various measures agnostic to the cell-type. These include conservation (PhyloP, PhastCons, GERP++), overlapping TFBS as predicted in JASPAR 2018 (exploring different score thresholds or factors predicted most frequently across the region; for details see Methods) and computational tools that integrate large sets of functional genomics data in combined scores (CADD, DeepSEA, Eigen, FATHMM-MKL, FunSeq2, GWAVA, LINSIGHT, ReMM). In addition, we compared a subset of experiments (10/21) to sequence-based models (deltaSVM) publically available only for specific cell-types (HEK293T, HeLa S3, HepG2, K562, and LNCaP). The table lists correlations with absolute expression effects in all rows other than "deltaSVM", where directionality of the expression alteration was considered. In cases where an annotation is based on positions rather than alleles, we assumed the same value for all substitutions at each position. MYC (rs11986220) is abbreviated to MYCs1 and MYC (rs6983267) abbreviated to MYCs2. TERT-GBM is abbreviated as TERT-G.

| Name | BCL11A | F9 | FOXE1 | GP1BA | HBB | HBG1 | HNF4a | IRF4 | IRF6 | LDLR.2 | MSMB | MYCs1 | MYCs2 | PKLR_48h | RET | SORT1.1 | TCF7L2 | TERT-G | UC88 | ZFAND3 | ZRSh13 |
|---|---|---|---|---|---|---|---|---|---|---|---|---|---|---|---|---|---|---|---|---|---|
| Type | enh. | prom. | prom. | prom. | prom. | prom. | prom. | enh. | enh. | prom. | prom. | enh. | enh. | prom. | enh. | enh. | enh. | prom. | UC | enh. | enh. |
| CADD | -0.04 | -0.09 | -0.07 | -0.09 | 0.00 | 0.10 | 0.10 | 0.42 | 0.18 | 0.57 | 0.01 | -0.07 | -0.06 | 0.39 | 0.14 | -0.28 | -0.01 | -0.02 | -0.02 | 0.40 | 0.15 |
| FunSeq | 0.00 | -0.03 | 0.05 | 0.19 | 0.13 | 0.13 | 0.08 | 0.35 | 0.12 | 0.36 | 0.23 | 0.02 | 0.01 | 0.47 | 0.11 | 0.07 | 0.09 | 0.15 | 0.06 | 0.25 | -0.08 |
| Eigen | -0.03 | -0.03 | 0.00 | 0.03 | 0.15 | 0.27 | 0.14 | 0.52 | 0.18 | 0.63 | 0.11 | -0.05 | -0.05 | 0.51 | 0.18 | -0.23 | 0.09 | 0.23 | -0.01 | 0.44 | 0.06 |
| FATHMM | 0.00 | -0.01 | 0.01 | 0.11 | 0.09 | 0.40 | 0.10 | 0.37 | 0.18 | 0.55 | 0.12 | 0.03 | -0.01 | 0.54 | 0.18 | -0.14 | 0.06 | 0.20 | 0.01 | 0.44 | 0.07 |
| DeepSEA | 0.02 | 0.17 | 0.01 | 0.19 | 0.21 | 0.25 | 0.13 | 0.32 | 0.12 | 0.31 | 0.17 | 0.02 | 0.09 | 0.33 | 0.11 | -0.05 | 0.09 | 0.31 | 0.04 | 0.33 | 0.04 |
| LINSIGHT | -0.01 | -0.08 | -0.02 | 0.10 | 0.13 | 0.09 | 0.14 | 0.46 | 0.14 | 0.51 | 0.02 | -0.06 | -0.10 | 0.57 | 0.19 | -0.21 | -0.03 | 0.11 | -0.02 | 0.38 | 0.04 |
| ReMM | -0.01 | -0.04 | -0.01 | 0.09 | 0.10 | 0.05 | 0.07 | 0.23 | 0.09 | 0.37 | -0.05 | -0.03 | -0.01 | 0.20 | 0.11 | -0.20 | -0.02 | 0.11 | 0.02 | 0.24 | -0.01 |
| GWAVA_U | 0.02 | 0.05 | -0.03 | 0.22 | -0.10 | -0.01 | 0.01 | -0.18 | -0.02 | 0.11 | 0.04 | -0.03 | -0.15 | 0.32 | 0.01 | -0.11 | -0.02 | -0.22 | 0.05 | 0.16 | -0.19 |
| GWAVA_T | -0.04 | 0.14 | -0.03 | 0.23 | -0.03 | 0.03 | 0.11 | 0.25 | -0.03 | 0.11 | 0.07 | 0.06 | -0.13 | 0.18 | 0.03 | 0.02 | 0.14 | -0.23 | -0.05 | 0.22 | -0.02 |
| GWAVA_R | 0.02 | -0.10 | -0.05 | 0.24 | -0.11 | 0.04 | 0.08 | 0.15 | -0.06 | 0.07 | 0.24 | -0.03 | -0.01 | 0.26 | 0.10 | 0.16 | 0.04 | -0.03 | 0.05 | 0.07 | -0.15 |
| deltaSVM | | 0.44 | 0.04 | | | | 0.07 | | | 0.26 | 0.23 | 0.07 | 0.23 | 0.48 | | 0.55 | | 0.27 | | | |
| deltaSVM (abs) | | 0.25 | 0.01 | | | | 0.03 | | | 0.28 | 0.14 | 0.13 | 0.20 | 0.42 | | 0.40 | | 0.21 | | | |
| JASPAR_All | 0.00 | 0.05 | 0.03 | 0.20 | 0.12 | 0.02 | 0.14 | 0.22 | 0.07 | -0.01 | 0.15 | 0.05 | -0.06 | 0.07 | 0.11 | 0.17 | 0.08 | 0.14 | 0.07 | 0.12 | 0.04 |
| JASPAR_90 | 0.02 | 0.05 | 0.02 | 0.19 | 0.14 | 0.03 | 0.14 | 0.23 | 0.08 | -0.04 | 0.16 | 0.05 | -0.05 | 0.09 | 0.11 | 0.17 | 0.07 | 0.14 | 0.08 | 0.11 | 0.04 |
| JASPAR_75 | 0.02 | 0.08 | 0.01 | 0.21 | 0.14 | 0.03 | 0.16 | 0.24 | 0.08 | -0.07 | 0.14 | 0.06 | -0.04 | 0.14 | 0.10 | 0.16 | 0.08 | 0.15 | 0.09 | 0.11 | 0.04 |
| JASPAR_50 | 0.04 | 0.11 | 0.01 | 0.18 | 0.16 | 0.08 | 0.12 | 0.25 | 0.10 | 0.02 | 0.14 | 0.06 | -0.03 | 0.21 | 0.11 | 0.18 | 0.08 | 0.18 | 0.11 | 0.12 | 0.04 |
| JASPAR_25 | 0.05 | 0.17 | 0.00 | 0.18 | 0.08 | 0.03 | 0.12 | 0.24 | 0.10 | 0.16 | 0.17 | 0.07 | 0.00 | 0.27 | 0.07 | 0.20 | 0.06 | 0.11 | 0.14 | 0.13 | 0.03 |
| JASPAR_10 | 0.05 | 0.18 | -0.01 | 0.18 | 0.08 | 0.04 | 0.13 | 0.28 | 0.06 | 0.32 | 0.14 | 0.05 | 0.08 | 0.22 | 0.06 | 0.14 | 0.08 | 0.14 | 0.12 | 0.07 | 0.08 |
| JASPAR_freqTF | -0.03 | -0.23 | 0.02 | -0.14 | 0.13 | 0.00 | 0.00 | 0.00 | -0.06 | 0.04 | -0.02 | 0.04 | -0.05 | 0.21 | -0.02 | -0.02 | 0.13 | 0.15 | 0.11 | 0.08 | 0.00 |
| priPhCons | -0.02 | -0.13 | 0.02 | -0.14 | -0.05 | 0.11 | 0.12 | 0.37 | 0.17 | 0.51 | 0.03 | 0.03 | -0.09 | 0.30 | 0.14 | -0.28 | -0.02 | 0.01 | 0.03 | 0.39 | 0.00 |
| mamPhCons | -0.01 | -0.05 | -0.01 | 0.10 | 0.11 | 0.34 | 0.10 | 0.51 | 0.15 | 0.64 | 0.11 | 0.03 | 0.02 | 0.56 | 0.18 | -0.21 | 0.04 | -0.05 | 0.02 | 0.44 | 0.00 |
| verPhCons | -0.01 | -0.01 | -0.01 | 0.09 | 0.09 | 0.39 | 0.10 | 0.34 | 0.18 | 0.62 | 0.12 | 0.03 | 0.02 | 0.54 | 0.18 | -0.20 | 0.04 | 0.12 | 0.02 | 0.45 | 0.06 |
| priPhyloP | -0.06 | 0.00 | -0.01 | 0.08 | 0.11 | 0.00 | 0.04 | 0.14 | 0.07 | 0.28 | 0.00 | -0.04 | 0.00 | 0.11 | 0.05 | -0.13 | 0.01 | 0.16 | 0.04 | 0.12 | 0.01 |
| mamPhyloP | -0.01 | 0.00 | 0.01 | 0.13 | 0.18 | 0.18 | 0.11 | 0.35 | 0.17 | 0.45 | 0.02 | -0.04 | 0.02 | 0.36 | 0.11 | -0.14 | 0.04 | 0.10 | -0.01 | 0.27 | 0.06 |
| verPhyloP | -0.01 | 0.02 | 0.00 | 0.10 | 0.14 | 0.20 | 0.11 | 0.37 | 0.21 | 0.51 | 0.02 | -0.04 | -0.02 | 0.38 | 0.12 | -0.15 | 0.04 | 0.11 | -0.05 | 0.26 | 0.07 |
| GerpN | -0.01 | -0.13 | 0.02 | 0.03 | 0.09 | -0.12 | 0.11 | 0.29 | 0.04 | 0.46 | 0.26 | 0.00 | -0.04 | 0.26 | 0.13 | -0.31 | 0.14 | 0.17 | -0.03 | 0.32 | 0.09 |

***Supplementary Table 13***

**Spearman correlation of computational scores with measured expression effects of all SNVs with at least 10 tags.** The table reports Spearman correlation coefficients of the absolute expression effect for each element with various measures agnostic to the cell-type. These include conservation (PhyloP, PhastCons, GERP++), overlapping TFBS as predicted in JASPAR 2018 (exploring different score thresholds or factors predicted most frequently across the region; for details see Methods) and computational tools that integrate large sets of functional genomics data in combined scores (CADD, DeepSEA, Eigen, FATHMM-MKL, FunSeq2, GWAVA, LINSIGHT, ReMM). In addition, we compared a subset of experiments (10/21) to sequence-based models (deltaSVM) publically available only for specific cell-types (HEK293T, HeLa S3, HepG2, K562, and LNCaP). The table lists correlations with absolute expression effects in all rows other than "deltaSVM", where directionality of the expression alteration was considered. In cases where an annotation is based on positions rather than alleles, we assumed the same value for all substitutions at each position. MYC (rs11986220) is abbreviated to MYCs1 and MYC (rs6983267) abbreviated to MYCs2. TERT-GBM is abbreviated as TERT-G.

| Name | BCL11A | F9 | FOXE1 | GP1BA | HBB | HBG1 | HNF4a | IRF4 | IRF6 | LDLR.2 | MSMB | MYCs1 | MYCs2 | PKLR_48h | RET | SORT1.1 | TCF7L2 | TERT-G | UC88 | ZFAND3 | ZRSh13 |
|---|---|---|---|---|---|---|---|---|---|---|---|---|---|---|---|---|---|---|---|---|---|
| Type | enh. | prom. | prom. | prom. | prom. | prom. | prom. | enh. | enh. | prom. | prom. | enh. | enh. | prom. | enh. | enh. | enh. | prom. | UC | enh. | enh. |
| CADD | -0.07 | 0.05 | -0.07 | -0.05 | 0.03 | 0.00 | 0.15 | 0.43 | 0.14 | 0.41 | -0.01 | -0.07 | -0.10 | 0.17 | 0.11 | -0.21 | -0.01 | 0.02 | -0.04 | 0.37 | 0.12 |
| FunSeq | 0.00 | 0.02 | 0.04 | 0.13 | 0.11 | 0.11 | 0.10 | 0.40 | 0.12 | 0.40 | 0.24 | 0.02 | 0.02 | 0.39 | 0.07 | 0.28 | 0.12 | 0.21 | 0.06 | 0.23 | 0.00 |
| Eigen | -0.03 | 0.03 | 0.02 | 0.00 | 0.15 | 0.16 | 0.22 | 0.48 | 0.19 | 0.58 | 0.10 | -0.02 | -0.04 | 0.25 | 0.13 | -0.13 | 0.09 | 0.20 | -0.04 | 0.40 | 0.07 |
| FATHMM | 0.02 | 0.03 | 0.04 | 0.10 | 0.11 | 0.15 | 0.18 | 0.29 | 0.22 | 0.59 | 0.14 | 0.00 | -0.09 | 0.26 | 0.11 | 0.02 | 0.04 | 0.20 | -0.03 | 0.40 | 0.07 |
| DeepSEA | 0.03 | 0.23 | 0.05 | 0.25 | 0.29 | 0.26 | 0.25 | 0.47 | 0.28 | 0.56 | 0.20 | 0.04 | 0.12 | 0.34 | 0.14 | 0.06 | 0.06 | 0.37 | 0.11 | 0.44 | 0.07 |
| LINSIGHT | -0.09 | 0.02 | 0.03 | 0.09 | 0.12 | 0.13 | 0.22 | 0.44 | 0.14 | 0.56 | 0.15 | -0.09 | -0.11 | 0.41 | 0.05 | -0.16 | 0.05 | 0.11 | -0.03 | 0.38 | 0.03 |
| ReMM | -0.02 | 0.04 | -0.01 | 0.07 | 0.11 | 0.00 | 0.14 | 0.30 | 0.19 | 0.52 | -0.09 | -0.04 | -0.01 | 0.19 | 0.14 | -0.16 | -0.06 | 0.00 | -0.07 | 0.30 | 0.05 |
| GWAVA_U | 0.00 | 0.10 | -0.04 | 0.28 | 0.03 | -0.09 | -0.04 | -0.07 | 0.06 | 0.06 | 0.01 | -0.03 | -0.11 | 0.32 | 0.01 | -0.20 | -0.03 | -0.23 | 0.07 | 0.16 | -0.15 |
| GWAVA_T | -0.01 | 0.14 | -0.03 | 0.23 | 0.09 | -0.08 | 0.18 | 0.23 | 0.03 | 0.00 | 0.08 | 0.06 | -0.10 | 0.18 | 0.01 | 0.09 | 0.16 | -0.28 | -0.05 | 0.24 | 0.02 |
| GWAVA_R | -0.02 | 0.04 | -0.08 | 0.28 | 0.05 | -0.03 | 0.09 | 0.24 | -0.01 | 0.04 | 0.27 | -0.02 | 0.02 | 0.32 | 0.10 | 0.16 | 0.05 | -0.04 | 0.04 | 0.09 | -0.15 |
| deltaSVM | | 0.43 | 0.04 | | | | 0.09 | | | 0.21 | 0.20 | 0.13 | 0.12 | 0.39 | | 0.53 | | 0.26 | | | |
| deltaSVM (abs) | | 0.20 | 0.01 | | | | 0.03 | | | 0.15 | 0.11 | 0.06 | 0.08 | 0.21 | | 0.28 | | 0.12 | | | |
| JASPAR_A | -0.01 | 0.11 | 0.01 | 0.28 | 0.01 | -0.02 | 0.11 | 0.14 | 0.06 | -0.02 | 0.23 | 0.06 | -0.09 | 0.03 | 0.13 | 0.20 | 0.13 | 0.05 | 0.04 | 0.15 | 0.00 |
| JASPAR_90 | 0.00 | 0.12 | 0.01 | 0.27 | 0.02 | -0.01 | 0.10 | 0.13 | 0.05 | -0.05 | 0.23 | 0.06 | -0.08 | 0.05 | 0.14 | 0.20 | 0.12 | 0.05 | 0.05 | 0.14 | 0.00 |
| JASPAR_75 | 0.00 | 0.17 | 0.00 | 0.30 | 0.01 | -0.01 | 0.11 | 0.13 | 0.07 | -0.04 | 0.21 | 0.06 | -0.09 | 0.11 | 0.15 | 0.22 | 0.13 | 0.06 | 0.05 | 0.13 | 0.00 |
| JASPAR_50 | 0.01 | 0.23 | 0.00 | 0.28 | 0.04 | 0.03 | 0.09 | 0.18 | 0.06 | 0.06 | 0.21 | 0.06 | -0.07 | 0.14 | 0.15 | 0.27 | 0.13 | 0.08 | 0.05 | 0.13 | 0.00 |
| JASPAR_25 | 0.01 | 0.39 | 0.00 | 0.29 | 0.05 | 0.00 | 0.08 | 0.24 | 0.06 | 0.18 | 0.22 | 0.05 | -0.02 | 0.21 | 0.16 | 0.33 | 0.15 | -0.02 | 0.04 | 0.11 | -0.01 |
| JASPAR_10 | 0.01 | 0.52 | 0.00 | 0.33 | 0.05 | 0.01 | 0.10 | 0.26 | 0.02 | 0.39 | 0.17 | 0.02 | 0.07 | 0.15 | 0.17 | 0.29 | 0.15 | 0.07 | 0.01 | 0.04 | 0.05 |
| JASPAR_TF | 0.00 | 0.24 | 0.01 | -0.14 | 0.10 | -0.08 | -0.02 | 0.01 | 0.01 | -0.04 | -0.02 | 0.03 | -0.04 | 0.08 | -0.02 | -0.04 | 0.15 | 0.14 | 0.10 | 0.14 | -0.02 |
| priPhCons | -0.02 | 0.07 | -0.01 | -0.11 | 0.00 | -0.03 | 0.23 | 0.40 | 0.15 | 0.54 | 0.01 | -0.01 | -0.13 | 0.18 | 0.15 | -0.19 | -0.02 | 0.09 | -0.03 | 0.37 | 0.01 |
| mamPhCons | -0.01 | 0.04 | -0.01 | 0.04 | 0.10 | 0.21 | 0.20 | 0.31 | 0.21 | 0.57 | 0.07 | -0.01 | -0.04 | 0.23 | 0.13 | -0.13 | 0.00 | 0.16 | -0.02 | 0.41 | 0.01 |
| verPhCons | -0.01 | 0.01 | -0.02 | 0.04 | 0.08 | 0.23 | 0.20 | 0.24 | 0.22 | 0.58 | 0.07 | 0.00 | -0.05 | 0.20 | 0.13 | -0.12 | 0.00 | 0.18 | -0.01 | 0.40 | 0.06 |
| priPhyloP | -0.01 | 0.01 | 0.02 | 0.13 | 0.11 | 0.03 | 0.02 | 0.23 | 0.07 | 0.35 | 0.05 | -0.01 | 0.05 | 0.06 | 0.04 | -0.12 | 0.02 | 0.12 | 0.02 | 0.18 | 0.04 |
| mamPhyloP | -0.01 | 0.03 | 0.03 | 0.05 | 0.19 | 0.12 | 0.15 | 0.24 | 0.17 | 0.44 | 0.00 | -0.01 | 0.01 | 0.17 | 0.11 | -0.11 | 0.03 | 0.14 | -0.02 | 0.27 | 0.06 |
| verPhyloP | -0.01 | 0.05 | 0.01 | 0.06 | 0.14 | 0.13 | 0.14 | 0.24 | 0.20 | 0.47 | 0.00 | -0.01 | -0.02 | 0.16 | 0.11 | -0.11 | 0.03 | 0.14 | -0.05 | 0.27 | 0.07 |
| GerpN | -0.01 | 0.02 | 0.00 | 0.01 | 0.05 | -0.03 | 0.22 | 0.47 | 0.04 | 0.59 | 0.31 | 0.00 | -0.10 | 0.28 | 0.11 | -0.28 | 0.17 | 0.13 | -0.10 | 0.37 | 0.07 |

***Supplementary Table 14***

**Previously described variants of promoters.** Clinical variants of promoters (HBB, HBG1, PKLR, F9, GP1BB, TERT, HNF4A, MSMB, and FOXE1) measured in this MPRA study, as well as additional described variants in Supplementary Note 1. Corresponding RefSeq transcripts of the HGVS annotation are listed in Supplementary Table 1. Fold changes are reported as "non significant" (n.s), if the associated p-value is higher than $10^{-5}$. Variants that were not available from our experiments are marked as “not covered” (n.c.). We are including effects for the PKLR_24h and PKLR_48h (PKLR_24h/PKLR_48h) as well as TERT-GBM and TERT-HEK experiments (TERT-GBM/TERT-HEK). Previous reported phenotypes in literature are summarized in the Phenotype column.

| Coordinates [GRCh37, GRCh38] | Variants | Expression effect [$log_2$] | Promoter activity (ratio WT) | Phenotype |
|---|---|---|---|---|
| X:138612869<br>X:139530710 | F9:c.-55G>C | n.s. | (0.84) | Hemophilia B Leyden |
| X:138612874<br>X:139530715 | F9:c.-50T>G | n.s. | (0.94) | Hemophilia B Leyden |
| X:138612875<br>X:139530716 | F9:c.-49T>A | -0.14 | 0.91 | Hemophilia B Leyden |
| X:138612889<br>X:139530730 | F9:c.-35G>A | n.s. | (0.8) | Hemophilia B Leyden |
| X:138612889<br>X:139530730 | F9:c.-35G>C | n.c. | n.c | Hemophilia B Leyden |
| X:138612890<br>X:139530731 | F9:c.-34A>T | n.s. | (1.00) | Hemophilia B Leyden |
| X:138612902<br>X:139530743 | F9:c.-22T>C | -0.15 | 0.9 | Hemophilia B Leyden |
| X:138612907<br>X:139530748 | F9:c.-17A>G | n.s. | (0.91) | Hemophilia B Leyden |
| X:138612907<br>X:139530748 | F9:c.-17A>C | n.s. | (0.89) | Hemophilia B Leyden |
| 9:100615914<br>9:97853632 | FOXE1:c.-283G>A, rs1867277 | n.s. | 0.94 | thyroid cancer-associated |
| 22:19710933<br>22:19723410 | GP1BB:c.-160C>G | -0.93 | 0.52 | Bernard-Soulier Syndrome |
| 11:5248402<br>11:5227172 | HBB:c.-151C>T | -0.28 | 0.82 | beta+ thal |
| 11:5248394<br>11:5227164 | HBB:c.-143C>G | n.s. | (0.90) | beta+ thal |

| | | | | |
|---|---|---|---|---|
| 11:5248393 11:5227163 | HBB:c.-142C>T | -0.23 | 0.85 | beta+ thal |
| 11:5248391 11:5227161 | HBB:c.-140C>T | -0.31 | 0.81 | beta+ thal |
| 11:5248389 11:5227159 | HBB:c.-138C>A | n.s. | (0.86) | beta+ thal |
| 11:5248389 11:5227159 | HBB:c.-138C>T | -0.29 | 0.82 | beta+ thal |
| 11:5248389 11:5227159 | HBB:c.-138C>G | n.s. | (0.89) | beta0 thal |
| 11:5248388 11:5227158 | HBB:c.-137C>A | -0.22 | 0.86 | beta+ thal |
| 11:5248388 11:5227158 | HBB:c.-137C>G | -0.25 | 0.84 | beta+ thal |
| 11:5248388 11:5227158 | HBB:c.-137C>T | -0.28 | 0.82 | beta+ thal |
| 11:5248387 11:5227157 | HBB:c.-136C>A | -0.18 | 0.88 | beta+ thal |
| 11:5248387 11:5227157 | HBB:c.-136C>G | n.s. | (0.85) | beta+ thal |
| 11:5248378 11:5227148 | HBB:c.-127G>C | n.s. | (0.89) | beta+ thal |
| 11:5248373 11:5227143 | HBB:c.-122T>A | -0.29 | 0.82 | beta+ thal |
| 11:5248372 11:5227142 | HBB:c.-121C>T | -0.25 | 0.84 | beta0 thal |
| 11:5248351 11:5227121 | HBB:c.-100G>A | -0.22 | 0.86 | beta+ thal |
| 11:5248350 11:5227120 | HBB:c.-99C>G | 1.19 | 2.28 | no previously reported phenotype |
| 11:5248333 11:5227103 | HBB:c.-82C>T | n.s. | (0.96) | beta0 thal |
| 11:5248333 11:5227103 | HBB:c.-82C>A | n.s. | (0.97) | beta+ thal |
| 11:5248332 11:5227102 | HBB:c.-81A>C | n.s. | (0.89) | beta+ thal |
| 11:5248332 11:5227102 | HBB:c.-81A>G | -0.15 | 0.90 | beta+ thal |
| 11:5248331 11:5227101 | HBB:c.-80T>C | -0.23 | 0.85 | beta0 thal |
| 11:5248331 11:5227101 | HBB:c.-80T>G | n.s. | (0.84) | beta0 thal |
| 11:5248331 11:5227101 | HBB:c.-80T>A | -0.28 | 0.82 | beta+ thal |
| 11:5248330 | HBB:c.-79A>G | -0.38 | 0.77 | beta+ thal |

| | | | | |
|---|---|---|---|---|
| 11:5227100 | | | | |
| 11:5248329<br>11:5227099 | HBB:c.-78A>C | -0.38 | 0.77 | beta+ thal |
| 11:5248329<br>11:5227099 | HBB:c.-78A>G | -0.34 | 0.79 | beta+ thal |
| 11:5248327<br>11:5227097 | HBB:c.-76A>C | n.s. | (0.88) | beta0 thal |
| 11:5248326<br>11:5227096 | HBB:c.-75G>C | n.s. | (0.97) | beta0 thal |
| 11:5248301<br>11:5227071 | HBB:c.-50A>C | n.s. | (0.94) | beta+ thal |
| 11:5248280<br>11:5227050 | HBB:c.-29G>A | n.s. | (0.91) | beta+ thal |
| 11:5248269<br>11:5227039 | HBB:c.-18C>G | n.s. | (1.01) | beta+ thal |
| 11:5271298<br>11:5250068 | HBG1:c.-264C>T | n.s. | (0.98) | HPFH |
| 11:5271289<br>11:5250059 | HBG1:c.-255C>T | -0.20 | 0.87 | HPFH |
| 11:5271289<br>11:5250059 | HBG1:c.-255C>G | n.s. | (1.05) | HPFH |
| 11:5271285<br>11:5250055 | HBG1:c.-251T>C | 0.31 | 1.24 | HPFH |
| 11:5271284<br>11:5250054 | HBG1:c.-250C>T | n.s. | (0.88) | HPFH |
| 11:5271283<br>11:5250053 | HBG1:c.-249C>T | n.s. | (0.98) | HPFH |
| 11:5271282<br>11:5250052 | HBG1:c.-248C>G | n.s. | (0.97) | HPFH |
| 11:5271262<br>11:5250032 | HBG1:c.-228T>C | 0.61 | 1.53 | HPFH |
| 11:5271245<br>11:5250015 | HBG1:c.-211C>T | 0.25 | 1.19 | HPFH |
| 11:5271204<br>11:5249974 | HBG1:c.-170G>A | n.s. | (1.09) | HPFH |
| 11:5271201<br>11:5249971 | HBG1:c.-167C>T | -0.63 | 0.65 | HPFH |
| 11:5271201<br>11:5249971 | HBG1:c.-167C>G | n.s. | (1.00) | HPFH |
| 11:5271201<br>11:5249971 | HBG1:c.-167C>A | -0.30 | 0.81 | HPFH |

| | | | | |
|---|---|---|---|---|
| 11:5271197<br>11:5249967 | HBG1:c.-163A>C | n.s. | (1.12) | HPFH |
| 20:42984253<br>20:44355613 | HNF4A:c.-192C>G | 0.20 | 1.15 | monogenic B cell diabetes |
| 20:42984264<br>20:44355624 | HNF4A:c.-181G>A | n.s. | (0.98) | monogenic B cell diabetes |
| 20:42984276<br>20:44355636 | HNF4A:c.-169C>T | n.s. | (0.99) | monogenic B cell diabetes |
| 20:42984299<br>20:44355659 | HNF4A:c.-146T>C | -0.06 | 0.96 | monogenic B cell diabetes |
| 20:42984307<br>20:44355667 | HNF4A:c.-138T>C | 1.04 | 2.06 | no previous reported phenotype |
| 20:42984309<br>20:44355669 | HNF4A:c.-136A>G | n.s. | (0.02) | monogenic B cell diabetes |
| 20:42984439<br>20:44355799 | HNF4A:c.-6T>G | 0.96 | 1.95 | no previous reported phenotype |
| 10:51549496<br>10:46046326 | MSMB:c.-89C>T, rs10993994 | 0.20 | 1.15 | prostate cancer-associated |
| 1:155271510<br>1:155301719 | PKLR:c.-324T>A | n.s./n.s. | (1.29)/(1.23) | nonfunctional polymorphisms |
| 1:155271434<br>1:155301643 | PKLR:c.-248delT | n.s./n.s. | (1.57)/(0.75) | nonfunctional polymorphisms |
| 1:155271269<br>1:155301478 | PKLR:c.-83G>C | n.s./-2.19 | (0.42)/0.22 | PK deficiency |
| 1:155271258<br>1:155301467 | PKLR:c.-72A>G | -1.77/-2.40 | 0.29/0.19 | PK deficiency |
| 5:1295349<br>5:1295234 | TERT:c.-245T>C, rs2853669 | n.s./n.s. | (0.95)/(0.99) | melanoma |
| 5:1295250<br>5:1295135 | TERT:c.-146C>T | 2.42/1.42 | 5.35/2.68 | glioblastoma |
| 5:1295242 - 1295243<br>5:1295127-1295128 | TERT:c.[-138C>T;-139C>T], rs35550267 | [0.20;n.s.]/ [n.s.;n.s.] | [1.15;(0.99)]/ [(1.04);(0.95)] | melanoma |
| 5:1295228-1295229<br>5:1295113-1295112 | TERT:c.[-124C>T;-125C>T] | [n.s.;2.86]/ [n.s.;2.00] | [(0.91);7.26]/ [(0.93);4.00] | bladder carcinoma, melanoma and glioma |
| 5:1295161<br>5:1295046 | TERT:c.-57A>C | 1.14/0.65 | 2.2/1.57 | melanoma, bladder cancer |

| | | | | |
|---|---|---|---|---|
| 5:1295158<br>5:1295043 | TERT:c.-54C>A | 1.65/0.81 | 3.14/1.75 | Bladder cancer |
| 5:1295149<br>5:1295034 | TERTc.-45:G>T | 1.03/n.s. | 2.04/(1.37) | Bladder cancer |

***Supplementary Table 15***

**Previously described variants of enhancers.** Clinical variants of enhancers (IRF4, IRF6, MYC, RET, TCF7L2, ZFAND3, SORT1, and ZRS) characterized in this MPRA study, as well as some additional described variants in the Supplementary Note 1. Fold changes are reported as "non significant" (n.s), if the associated p-value is higher than $10^{-5}$. Results from multiple experiments are separated by a dash and refer to SORT1/SORT1.2/SORT1-flip and ZRS+HOXD13/ZRS+HOXD13+HAND2, respectively. Phenotypes previously reported in literature are summarized in the Phenotype column.

| Coordinates [GRCh37, GRCh38] | Variants | Expression effect [$log_2$] | Enhancer activity (ratio WT) | Phenotype |
|---|---|---|---|---|
| 6:396321<br>6:396321 | IRF4:C>T, rs12203592 | -1.47 | 0.36 | pigmentation associated |
| 1:209989232<br>1:209815887 | IRF6:T>C, rs76145088 | -0.28 | 0.82 | no annotation |
| 1:209989270<br>1:209815925 | IRF6:C>T, rs642961 | n.s. | (1.06) | isolated cleft lip associated |
| 1:209989281<br>1:209815936 | IRF6:C>T, rs77542756 | n.s. | (1.00) | no annotation |
| 8:128413279<br>8:127401034 | MYCrs6983267:C>G | 0.92 | 1.86 | no previous reported phenotype |
| 8:128413289<br>8:127401044 | MYCrs6983267:G>T | 0.88 | 1.84 | no previous reported phenotype |
| 8:128413305<br>8:127401060 | MYCrs6983267:G>T, rs6983267 | n.s. | (0.95) | cancer related |
| 8:128531689<br>8:127519444 | MYC:T>A, rs11986220 | n.s. | (0.86) | cancer related |
| 10:43582056<br>10:43086608 | RET:T>C, rs2435357 | n.s. | (0.99) | Hirschsprung disease |
| 1:109817590<br>1:109274968 | SORT1:T>A, rs12740374 | 2.92/2.74 | 7.57/6.68 | Associated with plasma low-density lipoprotein cholesterol and myocardial infarctio. |
| 10:114758349<br>10:112998590 | TCF7L2:C>T, rs7903146 | 0.25 | 1.19 | Diabetes associated |
| 10:114758373<br>10:112998614 | TCF7L2:G>C | 1.18 | 2.27 | no previous reported phenotype |
| 6:37775652<br>6:37807876 | ZFAND3:C>T, rs58692659 | -0.51 | 0.7 | Diabetes associated |
| 7:156583831<br>7:156791137 | ZRS:T>C | n.s./n.s. | (1.04/1.04) | Preaxial polydactyly & TPT |
| 7:156583949 | ZRS:G>C | n.s./n.s. | (0.94/0.94) | Preaxial polydactyly & TPT |

| | | | | |
|---|---|---|---|---|
| 7:156791255 | | | | |
| 7:156583951<br>7:156791257 | ZRS:G>A | -0.07/n.s. | 0.95/(0.97) | Townes Brocks syndrome |
| 7:156584006<br>7:156791312 | ZRS:A>G | n.s./n.s. | (0.99/1.00) | No phenotype |
| 7:156584015<br>7:156791321 | ZRS:C>T | n.s./ n.s. | (0.99/0.98) | Polydactyly（mouse） |
| 7:156584093<br>7:156791399 | ZRS:T>C | n.s./n.s. | (1.01/0.99) | Polydactyly (cat) |
| 7:156584095<br>7:156791401 | ZRS:T>C | n.s. /n.s. | (1.01/1.00 ) | Polydactyly (cat) |
| 7:156584107<br>7:156791413 | ZRS:A>C | n.s./n.s. | (1.04/0.99) | Preaxial polydactyly & TPT |
| 7:156584153<br>7:156791459 | ZRS:T>C | 0.21 /0.12 | 1.16 /1.09 | Polydactyly of the four extremities and bilateral tibial deficiency |
| 7:156584163<br>7:156791469 | ZRS:A>T | n.s./n.s. | (1.01/0.98) | Polydactyly (mouse) |
| 7:156584164<br>7:156791470 | ZRS:T>C | 0.06/n.s. | 1.04/(1.03) | Polydactyly (mouse) |
| 7:156584166<br>7:156791472 | ZRS:C>A | n.s./n.s. | (0.91/0.9) | Polydactyly |
| 7:156584166<br>7:156791472 | ZRS:C>G | n.s./n.s. | (0.99/0.99) | Werner mesomelic syndrome |
| 7:156584166<br>7:156791472 | ZRS:C>T | n.s./n.s | (0.99/1.01) | Werner mesomelic syndrome |
| 7:156584168<br>7:156791474 | ZRS:G>A | n.s./-0.06 | (0.99)/0.96 | Werner mesomelic syndrome |
| 7:156584174<br>7:156791480 | ZRS:G>A | n.s. /n.s. | (1.02/1.01) | Polydactyly & TPT |
| 7:156584236<br>7:156791542 | ZRS:A>C | n.s./n.s. | (0.97/0.99) | Polydactyly |
| 7:156584241<br>7:156791547 | ZRS:A>G | 0.08/n.s. | 1.06 /(1.04) | Polydactyly |
| 7:156584265<br>7:156791571 | ZRS:T>A | n.s./n.s. | (0.99/0.99) | Polydactyly |
| 7:156584273<br>7:156791579 | ZRS:C>T | 0.13/n.s. | 1.09 /(1.04) | Polydactyly |
| 7:156584275<br>7:156791581 | ZRS:A>G | n.s./n.s. | (0.99/1.01) | TPT |
| 7:156584283<br>7:156791589 | ZRS:G>T | n.s./n.s. | (1.00/1.01) | Polydactyly & TPT & syndactyly |
| 7:156584285<br>7:156791591 | ZRS:C>A | 0.23/0.17 | 1.17/1.13 | Polydactyly (chicken) |

#### References

1. Reijnen, M. J., Sladek, F. M., Bertina, R. M. & Reitsma, P. H. Disruption of a binding site for hepatocyte nuclear factor 4 results in hemophilia B Leyden. *Proc. Natl. Acad. Sci. U. S. A.* **89,** 6300–6303 (1992).
2. Reijnen, M. J., Peerlinck, K., Maasdam, D., Bertina, R. M. & Reitsma, P. H. Hemophilia B Leyden: substitution of thymine for guanine at position -21 results in a disruption of a hepatocyte nuclear factor 4 binding site in the factor IX promoter. *Blood* **82,** 151–158 (1993).
3. Kheradpour, P. & Kellis, M. Systematic discovery and characterization of regulatory motifs in ENCODE TF binding experiments. *Nucleic Acids Res.* **42,** 2976–2987 (2014).
4. Landa, I. *et al.* The variant rs1867277 in FOXE1 gene confers thyroid cancer susceptibility through the recruitment of USF1/USF2 transcription factors. *PLoS Genet.* **5,** e1000637 (2009).
5. Ludlow, L. B. *et al.* Identification of a mutation in a GATA binding site of the platelet glycoprotein Ibbeta promoter resulting in the Bernard-Soulier syndrome. *J. Biol. Chem.* **271,** 22076–22080 (1996).
6. Khan, A. *et al.* JASPAR 2018: update of the open-access database of transcription factor binding profiles and its web framework. *Nucleic Acids Res.* **46,** D260–D266 (2018).
7. Giardine, B. *et al.* HbVar database of human hemoglobin variants and thalassemia mutations: 2007 update. *Hum. Mutat.* **28,** 206 (2007).
8. Moi, P. *et al.* A novel silent beta-thalassemia mutation in the distal CACCC box affects the binding and responsiveness to EKLF. *Br. J. Haematol.* **126,** 881–884 (2004).
9. Sgourou, A. *et al.* Thalassaemia mutations within the 5'UTR of the human beta-globin gene disrupt transcription. *Br. J. Haematol.* **124,** 828–835 (2004).
10. Gumucio, D. L. *et al.* Nuclear proteins that bind the human gamma-globin gene promoter: alterations in binding produced by point mutations associated with hereditary persistence of fetal hemoglobin. *Mol. Cell. Biol.* **8,** 5310–5322 (1988).
11. Sykes, K. & Kaufman, R. A naturally occurring gamma globin gene mutation enhances SP1 binding activity. *Mol. Cell. Biol.* **10,** 95–102 (1990).
12. Thomas, H. *et al.* A distant upstream promoter of the HNF-4alpha gene connects the transcription factors involved in maturity-onset diabetes of the young. *Hum. Mol. Genet.* **10,** 2089–2097 (2001).
13. Wirsing, A. *et al.* Novel monogenic diabetes mutations in the P2 promoter of the

HNF4A gene are associated with impaired function in vitro. *Diabet. Med.* **27,** 631–635 (2010).

14. Thomas, G. *et al.* Multiple loci identified in a genome-wide association study of prostate cancer. *Nat. Genet.* **40,** 310–315 (2008).
15. Eeles, R. A. *et al.* Multiple newly identified loci associated with prostate cancer susceptibility. *Nat. Genet.* **40,** 316–321 (2008).
16. Kote-Jarai, Z. *et al.* Multiple novel prostate cancer predisposition loci confirmed by an international study: the PRACTICAL Consortium. *Cancer Epidemiol. Biomarkers Prev.* **17,** 2052–2061 (2008).
17. Zheng, S. L. *et al.* Genetic variants and family history predict prostate cancer similar to prostate-specific antigen. *Clin. Cancer Res.* **15,** 1105–1111 (2009).
18. Chang, B.-L. *et al.* Fine mapping association study and functional analysis implicate a SNP in MSMB at 10q11 as a causal variant for prostate cancer risk. *Hum. Mol. Genet.* **18,** 1368–1375 (2009).
19. Buckland, P. R. *et al.* Strong bias in the location of functional promoter polymorphisms. *Hum. Mutat.* **26,** 214–223 (2005).
20. Manco, L. *et al.* A new PKLR gene mutation in the R-type promoter region affects the gene transcription causing pyruvate kinase deficiency. *Br. J. Haematol.* **110,** 993–997 (2000).
21. van Wijk, R. *et al.* Disruption of a novel regulatory element in the erythroid-specific promoter of the human PKLR gene causes severe pyruvate kinase deficiency. *Blood* **101,** 1596–1602 (2003).
22. Bauer, D. E. *et al.* An erythroid enhancer of BCL11A subject to genetic variation determines fetal hemoglobin level. *Science* **342,** 253–257 (2013).
23. Canver, M. C. *et al.* BCL11A enhancer dissection by Cas9-mediated in situ saturating mutagenesis. *Nature* **527,** 192–197 (2015).
24. Vinjamur, D. S. & Bauer, D. E. Growing and Genetically Manipulating Human Umbilical Cord Blood-Derived Erythroid Progenitor (HUDEP) Cell Lines. *Methods Mol. Biol.* **1698,** 275–284 (2018).
25. Praetorius, C. *et al.* A polymorphism in IRF4 affects human pigmentation through a tyrosinase-dependent MITF/TFAP2A pathway. *Cell* **155,** 1022–1033 (2013).
26. Rahimov, F. *et al.* Disruption of an AP-2alpha binding site in an IRF6 enhancer is associated with cleft lip. *Nat. Genet.* **40,** 1341–1347 (2008).
27. Yeager, M. *et al.* Genome-wide association study of prostate cancer identifies a second risk locus at 8q24. *Nat. Genet.* **39,** 645–649 (2007).
28. Haiman, C. A. *et al.* A common genetic risk factor for colorectal and prostate cancer.

*Nat. Genet.* **39,** 954–956 (2007).

29. Tomlinson, I. *et al.* A genome-wide association scan of tag SNPs identifies a susceptibility variant for colorectal cancer at 8q24.21. *Nat. Genet.* **39,** 984–988 (2007).
30. Pomerantz, M. M. *et al.* The 8q24 cancer risk variant rs6983267 shows long-range interaction with MYC in colorectal cancer. *Nat. Genet.* **41,** 882–884 (2009).
31. Ahmadiyeh, N. *et al.* 8q24 prostate, breast, and colon cancer risk loci show tissue-specific long-range interaction with MYC. *Proc. Natl. Acad. Sci. U. S. A.* **107,** 9742–9746 (2010).
32. Jia, L. *et al.* Functional enhancers at the gene-poor 8q24 cancer-linked locus. *PLoS Genet.* **5,** e1000597 (2009).
33. Zerbino, D. R., Wilder, S. P., Johnson, N., Juettemann, T. & Flicek, P. R. The ensembl regulatory build. *Genome Biol.* **16,** 56 (2015).
34. Emison, E. S. *et al.* A common sex-dependent mutation in a RET enhancer underlies Hirschsprung disease risk. *Nature* **434,** 857–863 (2005).
35. Lyssenko, V. *et al.* Mechanisms by which common variants in the TCF7L2 gene increase risk of type 2 diabetes. *J. Clin. Invest.* **117,** 2155–2163 (2007).
36. Bejerano, G. *et al.* Ultraconserved elements in the human genome. *Science* **304,** 1321–1325 (2004).
37. Visel, A. *et al.* Ultraconservation identifies a small subset of extremely constrained developmental enhancers. *Nat. Genet.* **40,** 158–160 (2008).
38. Visel, A., Minovitsky, S., Dubchak, I. & Pennacchio, L. A. VISTA Enhancer Browser--a database of tissue-specific human enhancers. *Nucleic Acids Res.* **35,** D88–92 (2007).
39. Pasquali, L. *et al.* Pancreatic islet enhancer clusters enriched in type 2 diabetes risk-associated variants. *Nat. Genet.* **46,** 136–143 (2014).
40. VanderMeer, J. E. & Ahituv, N. cis-regulatory mutations are a genetic cause of human limb malformations. *Dev. Dyn.* **240,** 920–930 (2011).
41. Gurnett, C. A. *et al.* Two novel point mutations in the long-range SHH enhancer in three families with triphalangeal thumb and preaxial polydactyly. *Am. J. Med. Genet. A* **143A,** 27–32 (2007).
42. Al-Qattan, M. M., Al Abdulkareem, I., Al Haidan, Y. & Al Balwi, M. A novel mutation in the SHH long-range regulator (ZRS) is associated with preaxial polydactyly, triphalangeal thumb, and severe radial ray deficiency. *Am. J. Med. Genet. A* **158A,** 2610–2615 (2012).
43. Farooq, M. *et al.* Preaxial polydactyly/triphalangeal thumb is associated with changed

transcription factor-binding affinity in a family with a novel point mutation in the long-range cis-regulatory element ZRS. *Eur. J. Hum. Genet.* **18,** 733–736 (2010).

44. Vanlerberghe, C. *et al.* Intrafamilial variability of ZRS-associated syndrome: characterization of a mosaic ZRS mutation by pyrosequencing. *Clin. Genet.* **88,** 479–483 (2015).

45. Wieczorek, D. *et al.* A specific mutation in the distant sonic hedgehog (SHH) cis-regulator (ZRS) causes Werner mesomelic syndrome (WMS) while complete ZRS duplications underlie Haas type polysyndactyly and preaxial polydactyly (PPD) with or without triphalangeal thumb. *Hum. Mutat.* **31,** 81–89 (2010).

46. VanderMeer, J. E. *et al.* A novel ZRS mutation leads to preaxial polydactyly type 2 in a heterozygous form and Werner mesomelic syndrome in a homozygous form. *Hum. Mutat.* **35,** 945–948 (2014).

47. Maas, S. A., Suzuki, T. & Fallon, J. F. Identification of spontaneous mutations within the long-range limb-specific Sonic hedgehog enhancer (ZRS) that alter Sonic hedgehog expression in the chicken limb mutants oligozeugodactyly and silkie breed. *Dev. Dyn.* **240,** 1212–1222 (2011).

48. Khamis, A. *et al.* Functional analysis of four LDLR 5'UTR and promoter variants in patients with familial hypercholesterolaemia. *Eur. J. Hum. Genet.* **23,** 790–795 (2015).

49. Bell, R. J. A. *et al.* Cancer. The transcription factor GABP selectively binds and activates the mutant TERT promoter in cancer. *Science* **348,** 1036–1039 (2015).

50. Horn, S. *et al.* TERT promoter mutations in familial and sporadic melanoma. *Science* **339,** 959–961 (2013).

51. Hurst, C. D., Platt, F. M. & Knowles, M. A. Comprehensive mutation analysis of the TERT promoter in bladder cancer and detection of mutations in voided urine. *Eur. Urol.* **65,** 367–369 (2014).

52. Huang, D.-S. *et al.* Recurrent TERT promoter mutations identified in a large-scale study of multiple tumour types are associated with increased TERT expression and telomerase activation. *Eur. J. Cancer* **51,** 969–976 (2015).
